# Adenovirus 14p1 induced changes in miRNA expression increases lung immunopathogenesis

**DOI:** 10.1101/2022.09.03.506485

**Authors:** Eric R. McIndoo, Ethan Wood, Gina Kuffel, Michael J. Zilliox, Jay R. Radke

## Abstract

Adenovirus is a frequent cause of mild, usually self-limited infections in infants and young children. Severe infections occur in immunocompromised patients but are rarely observed in healthy, immunocompetent adults. However, there have been outbreaks around the world of infections with different adenoviral (Ad) serotypes that have resulted in acute lung injury (ALI) and acute respiratory distress syndrome (ARDS) in some of those infected. Ad14p1, the predominant circulating strain of Ad14 worldwide is one such serotype. The explanations for the severity of illness caused by Ad14p1 infection in immunocompetent patients is unknown. Previously, we have shown that A549 cells infected with Ad14 repress macrophage pro29 inflammatory responses whereas cells infected with Ad14p1 fail to repress macrophages and, instead, can increase pro-inflammatory responses. Micro-RNAs (miRNA) are small noncoding RNAs that regulate gene expression at the posttranscriptional level. Adenoviral infection has been shown to modulate host miRNA expression, and we hypothesized that differences in miRNA expression between Ad14 and Ad14p1 infected cells might impact pathogenesis. A549 cells were infected with either Ad14 or Ad14p1 and total RNA samples were collected at 6, 12, 24, 36 and 48 post infection for miRNA sequencing. Cluster analysis revealed that there were 3 temporal changes in miRNA expression profiles following infection. Differential expression analysis showed 8-23 differentially expressed miRNA between Ad14 and Ad14p1 from 6 to 36hpi. However, at 48hpi there were 98 differentially expressed miRNAs in Ad14p1 infected cells compared to those infected by Ad14. Pathway enrichment analysis showed that the differentially expressed miRNA might explain the increased pathogenesis of Ad14p1caused by strain-related loss of modulation of cytokine expression. Overall, the data suggest a role for viral regulation of host miRNA expression in pathogenesis by regulating host inflammatory responses through the delivery of deregulated miRNAs by virally infected cell corpses to macrophages.

**Author Summary:** Acute respiratory distress syndrome (ARDS) is a severe inflammatory disease in the lungs, and both the onset and resolution of ARDS appears to be regulated primarily by alveolar macrophages. Emergent strains of human adenovirus (Ad) can induce ARDS in healthy immunocompetent people, whereas most, common Ad infections go unnoticed or causes minimal symptoms. Why emergent strains of Ad, such as Ad14p1, are more likely to induce acute lung injury (ALI) and ARDS is unknown. Cells that die from wild type Ad14 infection have been shown to repress alveolar macrophage inflammatory responses, whereas cells dying from Ad14p1 infection enhance inflammatory responses of alveolar macrophages. Here, we explored whether virus induced changes in the expression of cellular small regulatory RNAs could explain the differential effects of virally infected cells on alveolar macrophage inflammatory responses. The data show that there are differences in small RNA expression between Ad14 and Ad14p1 infected cells at late times after infection, when Ad14p1 infected cells lose expression of the small RNAs that repress pro-inflammatory cellular signaling pathways in macrophages. Our study provides insights into a mechanism that could drive the increased pathogenesis of some outbreak strains of adenovirus.

## Introduction

Ambros and Ruvkun discovered the first microRNA (miRNA), *lin-4*, in *C. elegans* in 1993[1,2]. miRNAs are small non-protein coding RNAs that can post-transcriptionally regulate gene expression, have been found in all animal models and show a high degree of conservation across species[3–5]. miRNAs are transcribed from DNA into primary miRNAs and processed into precursor miRNAs and finally mature miRNAs[6,7]. Mature miRNAs average 22-25 nucleotides in length and regulate gene expression by interacting with mRNAs primarily in the 3’ UTR but can also interact with the 5’ UTR mRNA coding sequence and gene promoters to suppress mRNA expression[6,8]. Mature miRNA associated with Argonaute to form the RNA-induced silencing complex (RISC)[9]. In the RISC, nucleotides 2-8 of the miRNA (seed sequence) bind to complementary sequences of the target mRNA through Watson-Crick base pairing resulting in the cleavage of the mRNA[10]. This provides the cell with a mechanism to fine tune gene expression post-transcriptionally. While the majority of miRNA reside intracellularly, miRNA can be excreted by cells into the blood via exosomes or can be shed through other membrane vesicles such as apoptotic bodies providing a way for communication with neighboring cells[11–20]. Secreted miRNA can be biomarkers for many types of cancer, sepsis, nervous system disorders, traumatic brain injury and infectious diseases[21–23].

miRNAs can play a role in regulating the immune response to viral infection and may be able to be used as biomarkers to indicate severe infection[24]. Rhinovirus, respiratory syncytial virus, human metapneumovirus, SARS-CoV and SARS-CoV-2, influenza A and adenovirus (Ad) infection results in differential miRNA expression in infected cells and in the blood of infected individuals[25–35]. miRNA expression in cells infected with respiratory viruses could regulate host inflammatory responses and could thereby determine the altered pathogenesis of emerging viral strains[36].

Respiratory Ad infection usually results in mild, self-limited infections in immunocompetent individuals. However, outbreaks of emergent strains of Ad have resulted in severe and sometimes fatal infections in otherwise healthy people[37]. Ad14p1 is one such emergent strain, which first emerged in the U.S. and subsequently throughout the world resulting in acute lung injury (ALI) and acute respiratory distress syndrome (ARDS)[38–44]. Ad14 is a member of the B2 subgroup of adenovirus and uses Desmoglein-2 as a receptor to infect respiratory epithelial cells[45]. Ad14 and Ad14p1 are 99.7% genetically identical[46]. We have shown that Syrian hamsters are fully permissive for Ad14/Ad14p1 infection and that hamster infection with Ad14p1 results in lung pathology that is consistent with ALI and early stages of ARDS[47]. In vitro infection of A549 cells with prototype Ad14 results in dying cells that repress human alveolar macrophage pro-inflammatory responses. In contrast, infection with pathogenic Ad14p1 strain results in dying cells that enhance pro-inflammatory responses of human alveolar macrophages[48,49]. These results suggest that differential modulation of the host innate immune response to Ad14/Ad14p1 infections may determine whether infection results in resolution of viral pathogenesis or progression to acute lung injury and ARDS[49–52]. The objective of the current study was to determine whether Ad14p1 infection leads to differential miRNA expression that could explain the disparate inflammatory responses of alveolar macrophages to cells dying from Ad14 vs Ad14p1 infection.

Here we show that both Ad14 and Ad14p1 infection of A549 cells results in temporal changes in miRNA expression during the course of HAdV infection. Globally, both Ad14 and Ad14p1 have a similar effect of miRNA expression at early times after infection. However, the miRNA patterns are markedly different at late times after infection, when viral replication results in cell death. miRNA target and pathway enrichment analyses suggest that the differentially expressed miRNAs may account for virus strain-specific differences in immunopathogenesis. These data provide one explanation for the divergent inflammatory responses of alveolar macrophages to cells dying from Ad14 vs. Ad14p1 infection.

## Results

### Cellular miRNA Expression in A549 cells during Ad14 and Ad14p1 infection

To exaimine the effects of Ad14/Ad14p1 infection on cellular miRNAs, A549 cells were infected with either Ad14 or Ad14p1 at an MOI of 10 plaque forming units (PFU)/cell. Small RNA sequencing libraries were made from total RNA extracted from cells at varying points post infection that represent the full infectious cycle from early infection through full cytopathic effect (CPE). As shown in table 1, at 6 hours post infection (hpi), there was little change in the percent of small RNA reads that map to known human miRNAs, with less than one percent of the total reads mapping to Ad14/Ad14p1 viral miRNAs (mivaRNA) that are encoded in the VA RNA gene. Beginning at 12hpi, the percent of small RNAs that map to cellular miRNAs dropped, as mivaRNA expression increased. At 24hpi and later, approximately 38% of the reads mapped to cellular miRNAs, and ~20% of the reads mapped to mivaRNAs. This is consistent with previous observations in Ad2 and Ad3 infected tissue culture cells[25,26].

**Table 1.** Summary of cellular miRNA and HAdV14/p1 mivaRNA expression

|  | Mock | 6hpi | 12hpi | 24hpi | 36hpi | 48hpi |
| --- | --- | --- | --- | --- | --- | --- |
| **HAdV14** |  |  |  |  |  |  |
| Total Reads<sup>a</sup> | 2,915,019 | 3,408,959 | 3,391,732 | 3,387,175 | 3,330,665 | 2,821,104 |
| miRBase (%<sup>b</sup>) | 1,794,862<br>(61.57%) | 2,198,516<br>(64.49) | 1,628,255<br>(48.01) | 1,389,443<br>(41.02) | 1,277,346<br>(38.35) | 1,376,789<br>(48.8) |
| mivaRNA (%<sup>c</sup>) | 300 (0.013) | 26,676<br>(0.78) | 506,467<br>(14.93) | 835,406<br>(24.63) | 617,366<br>(18.54) | 419,037<br>(14.85) |
| **HAdV14p1** |  |  |  |  |  |  |
| Total Reads |  | 2,803,990 | 3,267,888 | 4,990,604 | 4,160,615 | 3,310,553 |
| miRBase (%<sup>b</sup>) |  | 1,755,693<br>(62.61) | 1,533,510<br>(46.93) | 1,738,608<br>(34.84) | 1,507,889<br>(36.24) | 1,242,118<br>(37.52) |
| mivaRNA (%<sup>c</sup>) |  | 14,835<br>(0.53) | 258,305<br>(7.90) | 1,032,127<br>(20.68) | 891,497<br>(21.42) | 640,156<br>(19.34) |

<sup>a</sup> Cumulative from 4 replicate infections and after discarded reads
<sup>b</sup> Percentage of total reads that mapped to miRbase V22
<sup>c</sup> Percentage of total reads that aligned to VA RNA, which map predominantly to 5' and 3' mivaRNAs

### Ad14 and Ad14p1 deregulate cellular miRNA expression

While other studies have shown that Ad infection can deregulate cellular miRNA expression, we wanted to determine whether there are differences in miRNA expression following infection with the non-pathogenic Ad14 compared to the pathogenic Ad14p1 strain. To understand if Ad14 and Ad14p1 infection have the same effect on cellular miRNA expression, principle component analysis (PCA) and heatmap with hierarchical clustering were used to indentify patterns in the large complex datasets. Both Ad14 and Ad14p1 infection caused disregulation of cellular miRNAs starting at 6hpi (Fig 1a, light and dark blue dots) and to a similar degree at 12hpi. At 24hpi deregulation increased the principle component (PC) 2 direction). Overall through 24hpi, Ad14 and Ad14p1 deregulation of miRNAs was similar. At 36hpi, strain (Fig 1a, Ad14 light green and Ad14p1 teal) dependent effects on cellular miRNA expression could be detected (in the PC2 direction) and were even more apparent (a shift in the PC1 direction) at 48hpi (Fig 1a, Ad14 red and Ad14p1 brown). At that point infection had proceeded to full CPE – virus-induced cytopathic effect of all cells in the culture. The deregulation seen at 48hpi was distinct from the deregualtion seen at earlier time points. Overall infection resulted in both temporal and viral strain-specific deregulation of cellular miRNAs. Heatmap clustering of the top 75 expressed miRNAs (Fig 1b) showed both time related and viral strain-specific differences between Ad14 and Ad14p1 infection on miRNA expression profiles, with the largest differences seen at 48hpi.

**Fig 1.**
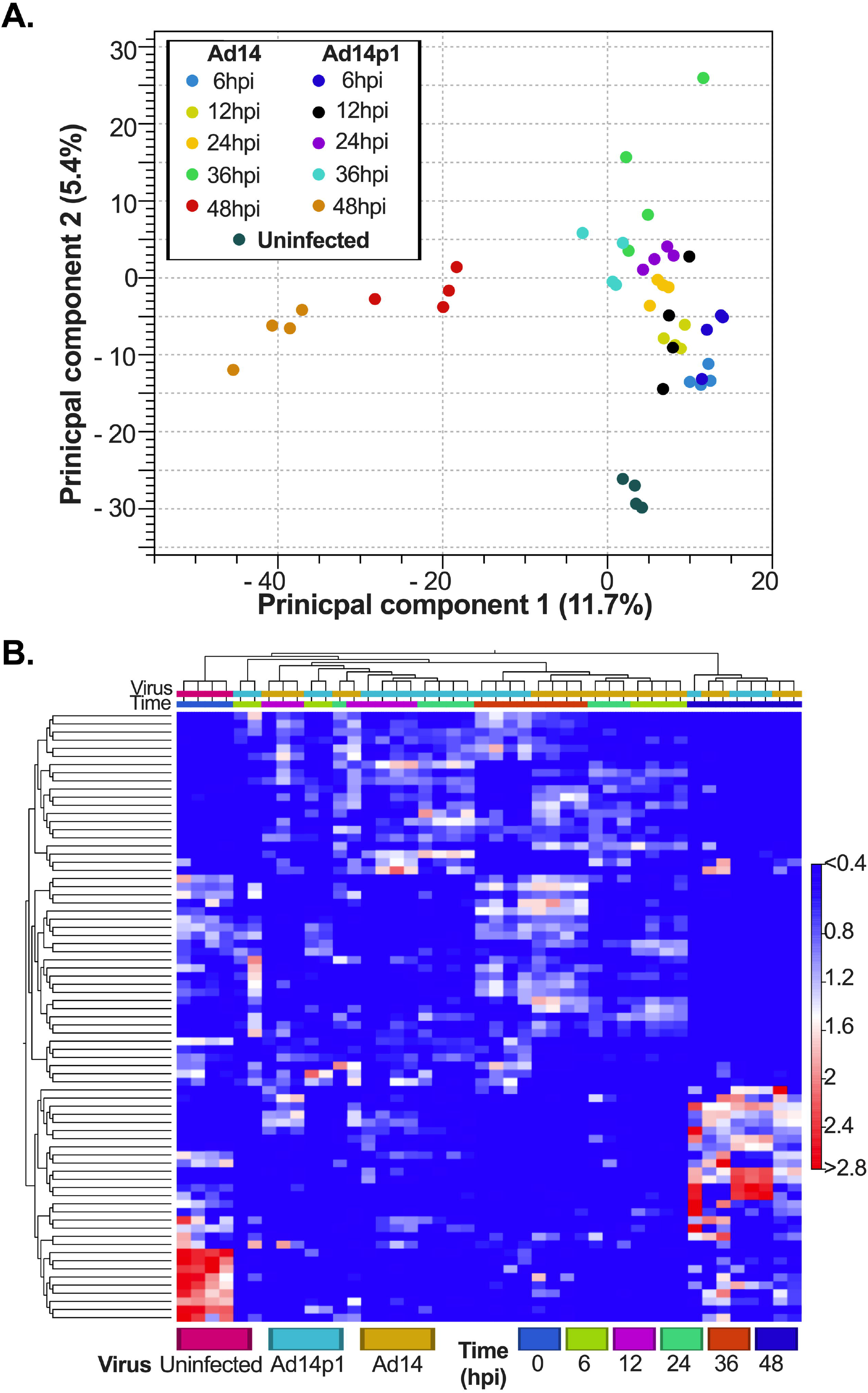
Unsupervised analysis of miRNA expression during Ad14 and Ad14p1 infection. A549 cells were infected with Ad14 or Ad14p1 at 10 PFU per cell, and miRNAseq was performed from total RNA at the indiated times. **A**. PCA plot of cellular miRNA expression during Ad14 and Ad14p1 infection. **B**. Heatmap with Euclidean distance clustering of miRNA expression of the top 75 features.

### Differential miRNA expression during Ad14 and Ad14p1 infection

Based on the unsupervised analysis, differential expression analysis was performed on miRNA from both Ad14 and Ad14p1 viral infections at each time point against uninfected cells and compared with each other. miRNAs that had a false discovery rate (FDR) adjusted p-value that was <0.05 were considered significant (Table 2 and S1 Table). Infections with Ad14 or Ad14p1 both up-regulated and down-regulated miRNA expression compared to uninfected cells. At 6hpi, both viruses had more up-regulated than down-regulated miRNA. The number of up-regulated miRNA fell at 12hpi and the ratio of up-regulated to down-regulated miRNA was nearly equal. From 24hpi to 48hpi, the number of total deregulated miRNA increased in both infections, with some differences. In Ad14 infected cells, the total number of up-regulated and down-regulated miRNAs were very similar at all time points. In Ad14p1 infected cells the number of deregulated miRNA decreased again at 36hpi before a dramatic increase at 48hpi that was far higher than in Ad14 infected cells, consistent with the PCA performed.

**Table 2:**
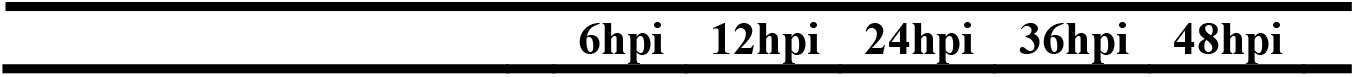
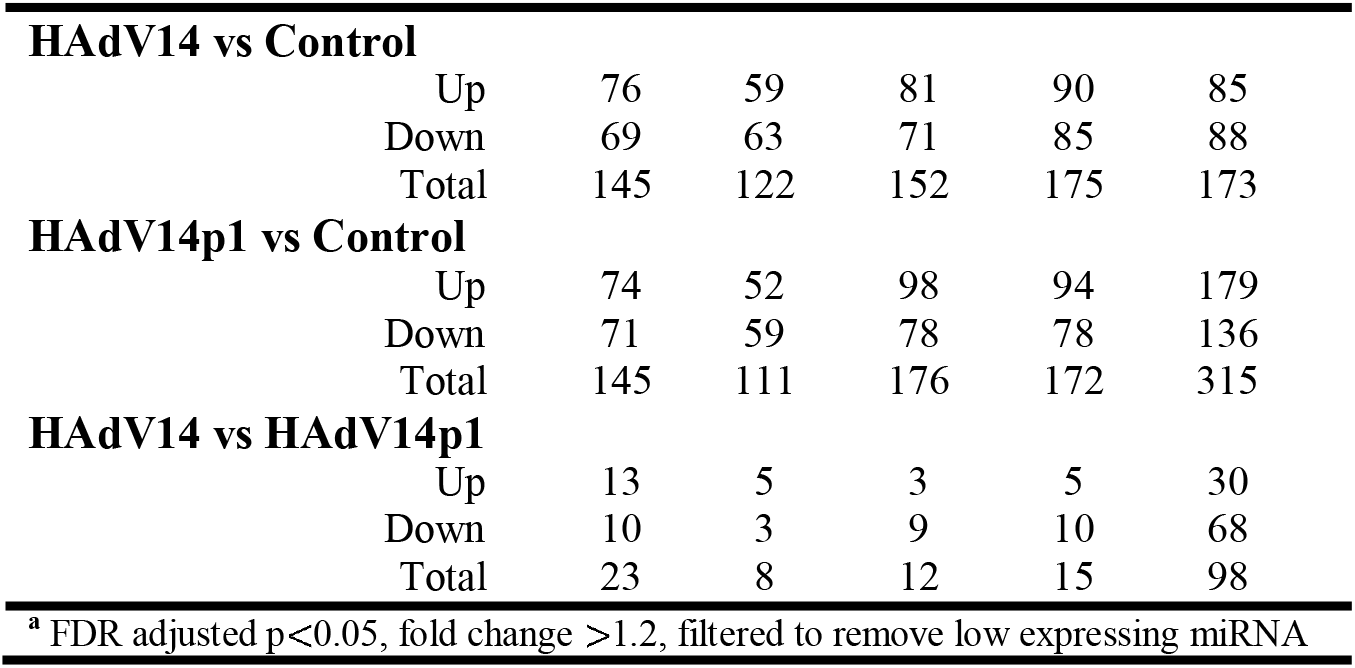
Number of differentially^**a**^ expressed miRNA during infection

| 6hpi | 12hpi | 24hpi | 36hpi | 48hpi |
| --- | --- | --- | --- | --- |

|  |  |  |  |  |  |
| --- | --- | --- | --- | --- | --- |
| **HAdV14 vs Control** |  |  |  |  |  |
| Up | 76 | 59 | 81 | 90 | 85 |
| Down | 69 | 63 | 71 | 85 | 88 |
| Total | 145 | 122 | 152 | 175 | 173 |
| **HAdV14p1 vs Control** |  |  |  |  |  |
| Up | 74 | 52 | 98 | 94 | 179 |
| Down | 71 | 59 | 78 | 78 | 136 |
| Total | 145 | 111 | 176 | 172 | 315 |
| **HAdV14 vs HAdV14p1** |  |  |  |  |  |
| Up | 13 | 5 | 3 | 5 | 30 |
| Down | 10 | 3 | 9 | 10 | 68 |
| Total | 23 | 8 | 12 | 15 | 98 |
| <sup>a</sup> FDR adjusted p<0.05, fold change >1.2, filtered to remove low expressing miRNA |  |  |  |  |  |

The heatmap analysis showed that both Ad14 and Ad14p1 miRNA profiles between 6 and 36hpi were similar, while at 48hpi they were not. Venn diagram analysis was used to understand how similar the miRNA expression profiles were in Ad14 and Ad14p1 infected cells. As seen in figure 2, approximately 50% of the differentially expressed miRNA in Ad14 and Ad14p1 infected cells were shared (overlap of yellow [Ad14] and blue [Ad14p1] circles). Differential miRNA expression between Ad14 and Ad14p1 infected cells (Fig 2, pink circles) showed that, between 6 to 36 hpi, only 8-23 miRNA were differentially expressed in either strain. At 48 hpi, there were 98 differentially expressed miRNA in Ad14 infected cells compared to Ad14p1 infected cells. 55 of the 98 miRNA were differentially regulated in Ad14 vs Ad14p1 and Ad14p1 vs uninfected cells. 5 were differentially regulated in Ad14 vs Ad14p1 and Ad14 vs uninfected cells. 30 were differentially regulated in all comparisons and 8 were only differentially regulated in Ad14 vs Ad14p1 infected cells.

**Fig 2.**
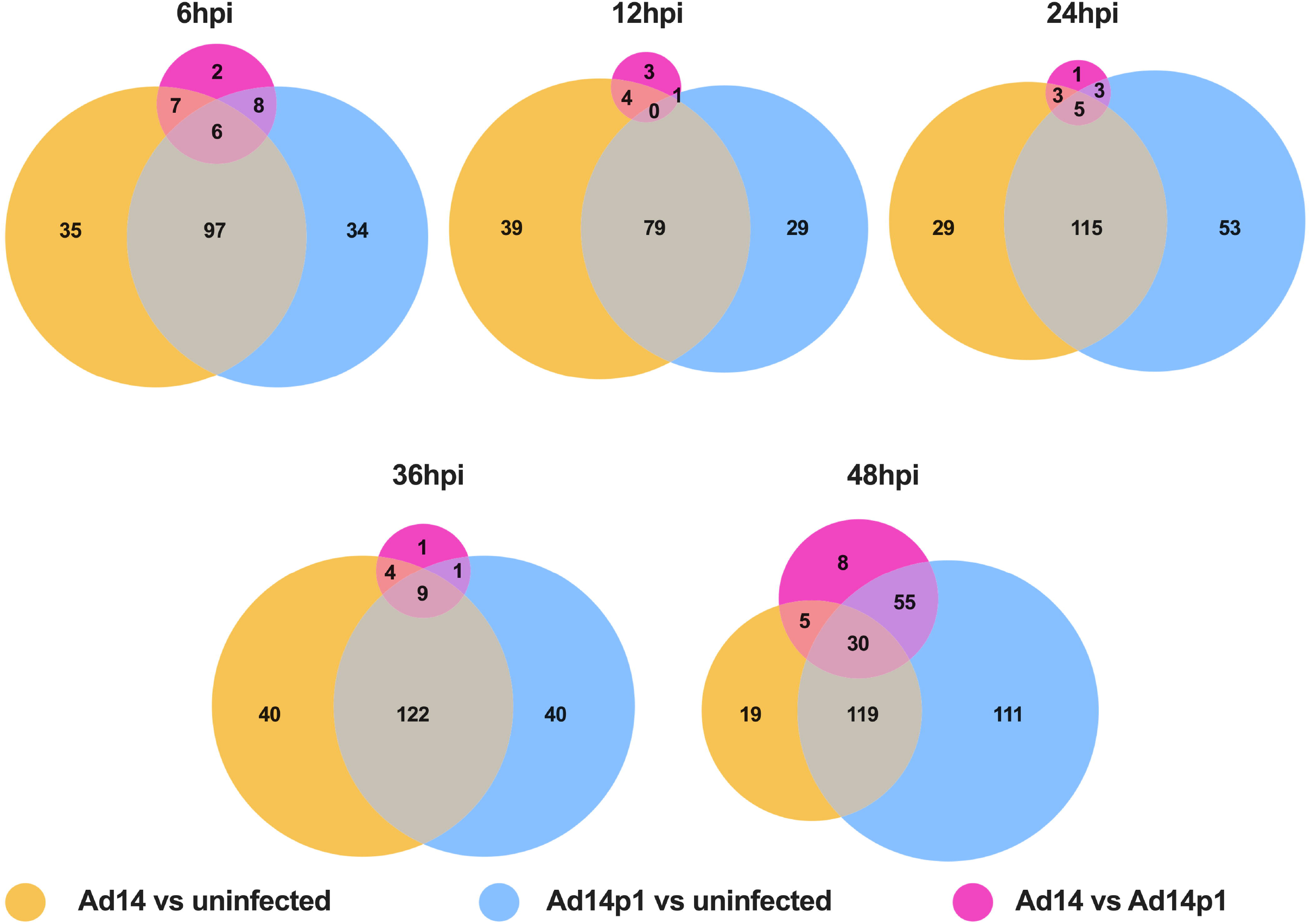
Ad14 and Ad14p1 deregulation of miRNA expression. Venn diagrams of miRNA expression over time in Ad14 and Ad14p1 infected A549 cells. Comparisons are Ad14 vs uninfected cells (yellow circles), Ad14p1 vs uninfected cells (blue circles) and Ad14 vs Ad14p1 infected cells (pink circles). Differential expression was determined by an FDR-adjusted p value <0.05 and absolute fold change >1.2 with N=4.

### Ad14 differentially expressed miRNA target cell signalling pathways

To begin to understand whether macrophage immunosuppression caused by Ad14 corpses was associated with their miRNA content, we focused on the 98 miRNA that were differentially expressed in Ad14 vs Ad14p1 corpses (S1 Table). To select for differentially expressed miRNAs in Ad14 CPE corpses compared to Ad14p1 CPE corpses that have the potential for a biological effect, a threshold was set for a mean expression of the miRNAs >1000. This resulted in detecting 10 miRNAs that were enriched in Ad14 CPE corpses vs Ad14p1 CPE coprses (Table 3). To understand the potential cellular effects of these miRNAs, mirPath v3 was used for KEGG and GO functional enrichment[53]. Using TarBase v7.0, the enriched Ad14 CPE miRNAs were shown to target between 249 to 2429 genes (Fig 3A and S2 Table). KEGG Pathway enrichment analysis showed that multiple miRNAs target thyroid hormone, FOXO, p53, HIPPO, PI3K-Akt and MAPK signalling pathways (Fig 3B&C and S1 Fig). GO analysis showed that TLR, MAPK, Fc-epsilon receptor, epidermal growth factor, neurotropin TRK receptor and TGF-β receptor signaling pathways are targeted by the enriched miRNAs (Fig 3D & S1 Fig). The web-based tool MIENTURNET was used to further probe network-based analysis of the enriched miRNAs[54]. In MIENTURNET, target enrichment was performed with a minimum of 2 miRNA-mRNA interactions with recommended default FDR threshold with TarBase v7.0. This resulted in 568 target genes for the 10 enriched Ad14 CPE miRNAs, of which 22 of the 568 genes are targeted by at least 2 or more of the enriched miRNAs (Figs 4A & 4B). Functional enrichment with Wikipathway, KEGG and Reactome showed multiple signaling pathways to be targeted by most of the enriched miRNAs (Fig 4C & S2 Fig).

**Table 3.** miRNAs enriched in Ad14 CPE corpses

| miRNA | Max Group Mean<sup>a</sup> | Fold Increase<sup>b</sup> | FDR p-value<sup>c</sup> |
| --- | --- | --- | --- |
| hsa-miR-27a-3p | 71095.25 | 1.348751 | 0.012899 |
| hsa-miR-21-5p | 67880.5 | 1.31362 | 0.025758 |
| hsa-let-7a-5p | 27083.75 | 1.353496 | 0.01777 |
| hsa-miR-22-3p | 19373.75 | 1.558081 | 0.000147 |
| hsa-miR-181a-5p | 18607.75 | 1.547176 | 0.000477 |
| hsa-miR-25-3p | 17419.5 | 1.38644 | 0.010636 |
| hsa-miR-31-5p | 2736.25 | 1.450643 | 0.003482 |
| hsa-miR-28-3p | 2501 | 1.445523 | 0.005461 |
| hsa-miR-151a-5p | 1980.75 | 1.345793 | 0.025758 |
| hsa-miR-221-3p | 1947.5 | 1.414809 | 0.005475 |

<sup>a</sup> Mean read counts from 4 different infections
<sup>b</sup> Fold increase in miRNA expression caused by Ad14 vs Ad14p1 infection
<sup>c</sup> Benjamini-Hochberg test for false discovery rate

**Fig 3.**
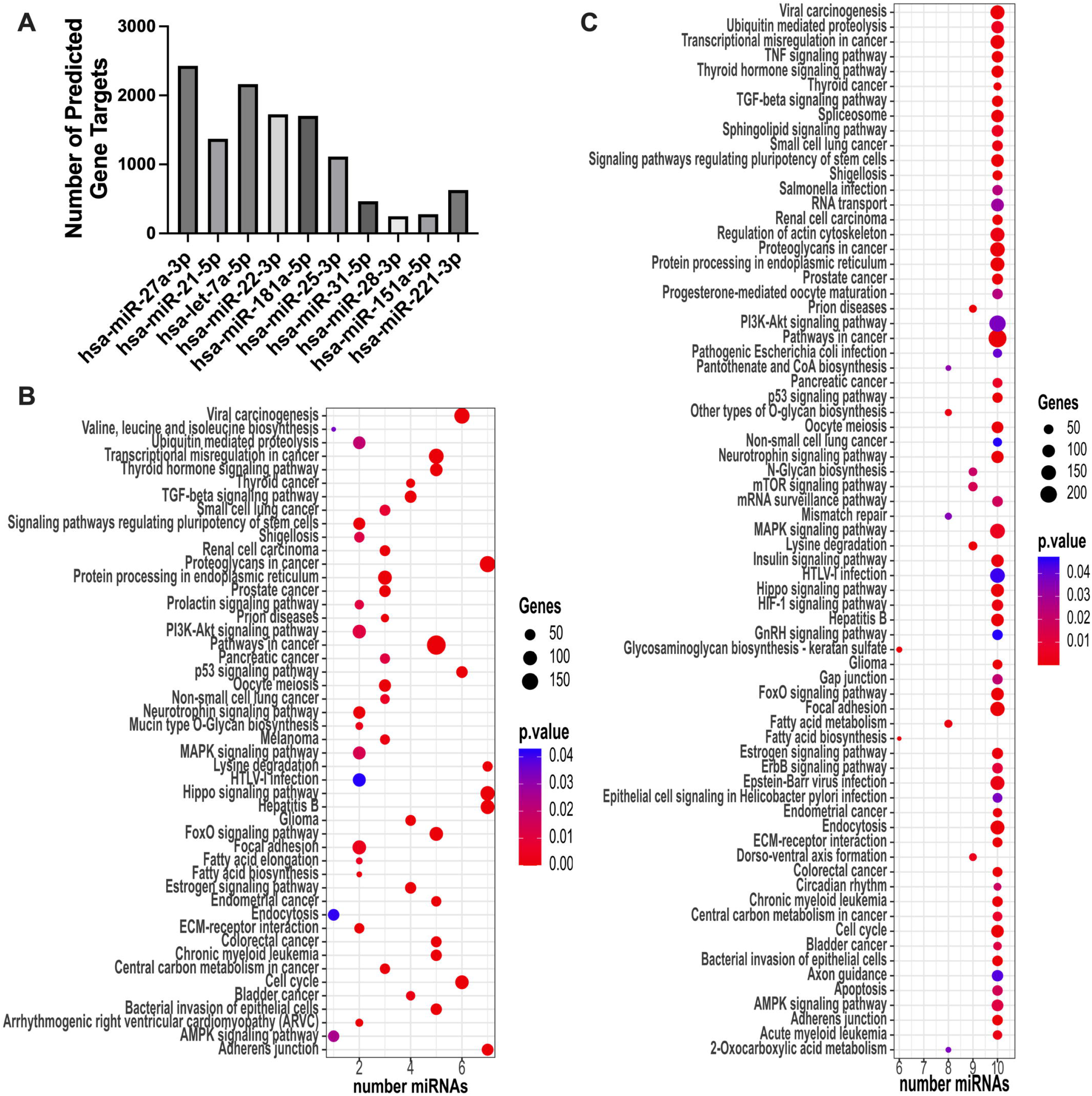
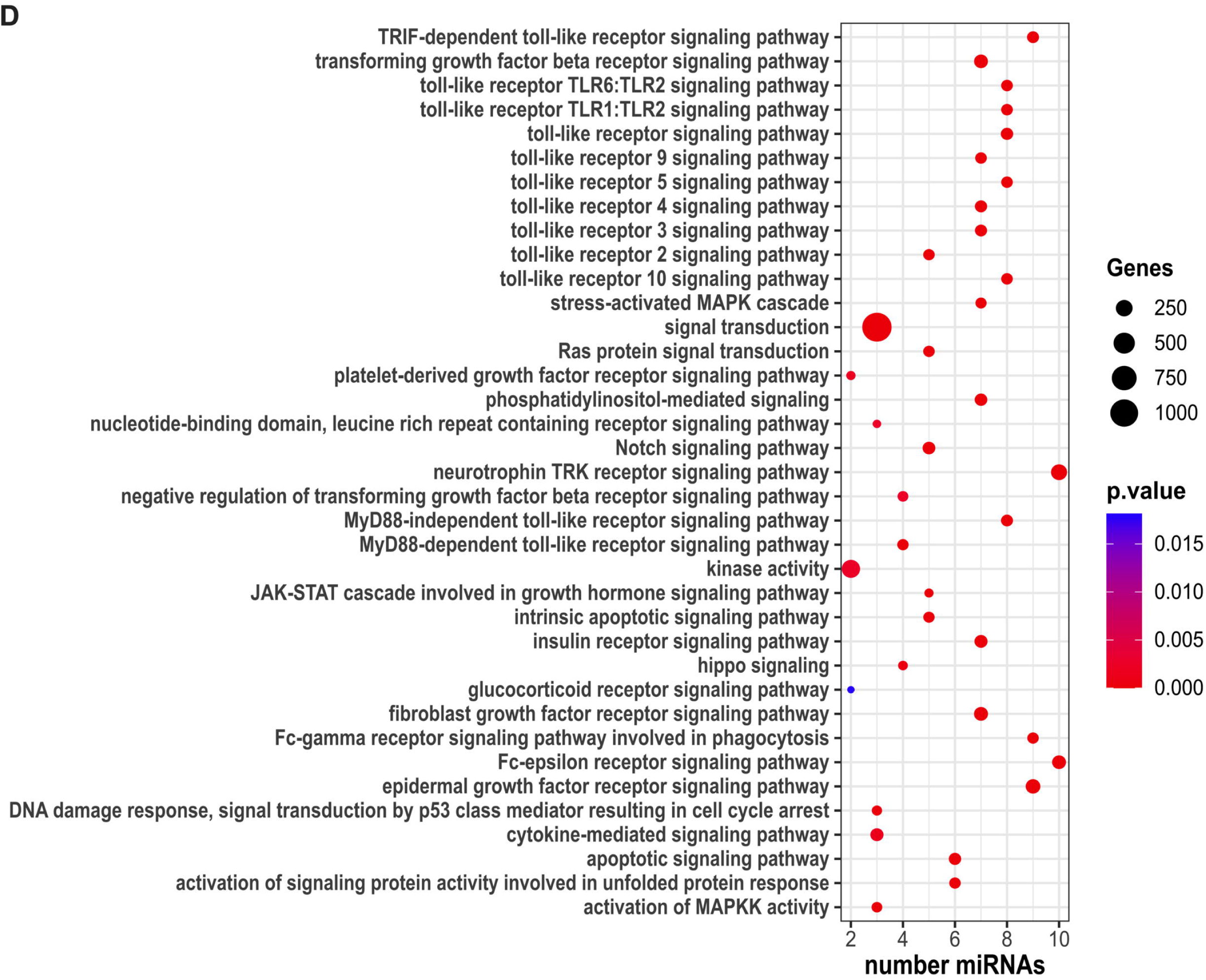
Bioinformatic analysis of Ad14 miRNA by mirPath V3. The 10 enriched miRNAs in Ad14 CPE corpses were uploaded to mirPath V3. **A**. Total number of predicted target genes per miRNA. **B**. Dot plot of KEGG Pathways Union analysis. **C**. Dot plot of KEGG Genes Union analysis. **D**. Dot plot of Signaling Pathways idenfited by GO Categories Union analysis.

**Fig 4.**
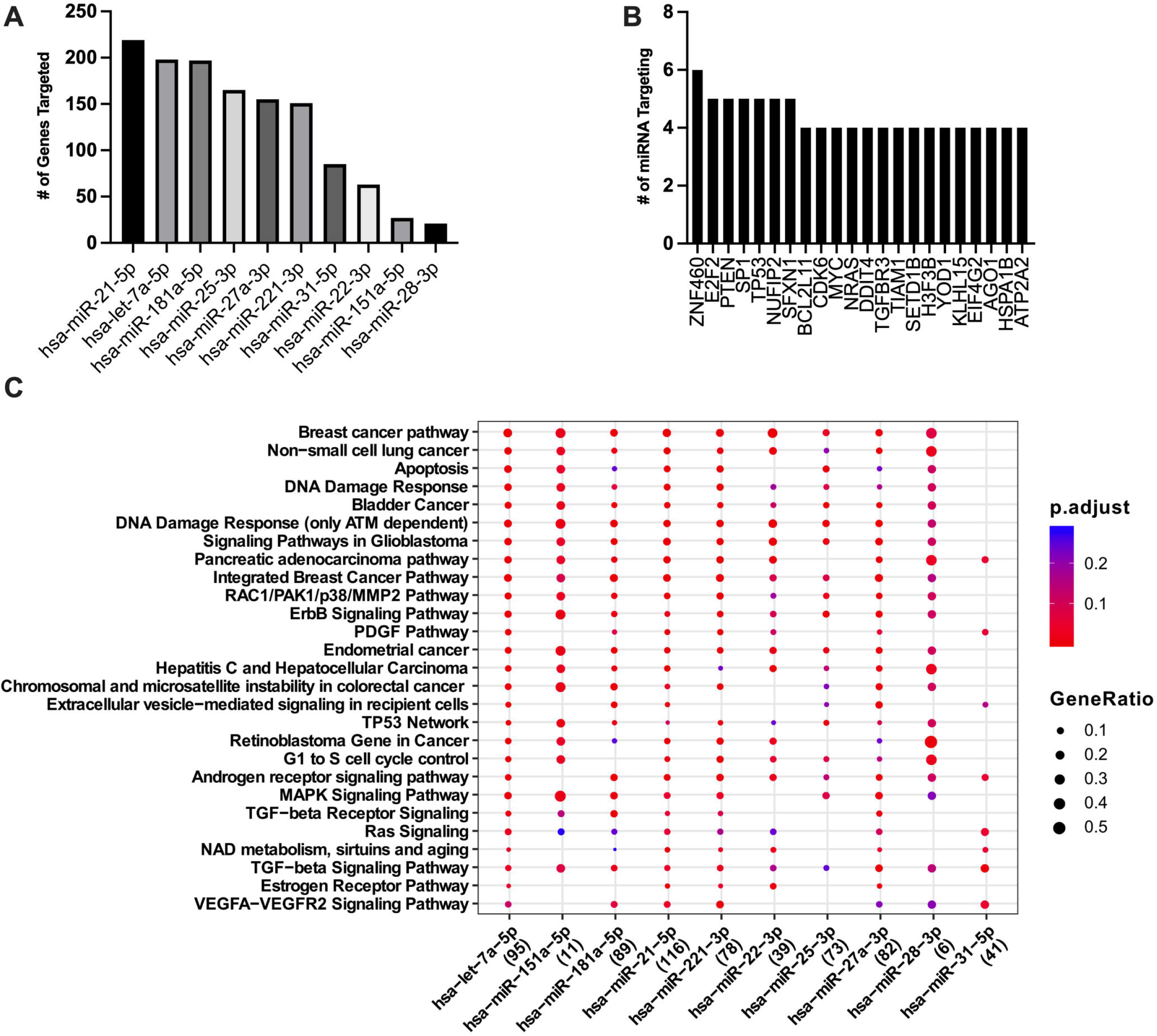
MIENTURNET target and functional enrichment of Ad14 miRNA. The 10 enriched miRNAs in Ad14 CPE corpses were uploaded to MIENTURNET. Targent enrichment was performed with miRTarBase with a minimum of 2 miRNA-RNA interactions and an FDR threshold of 1. **A**. Number of genes targeted per miRNA. **B**. Number of miRNAs targeting each predicted gene for genes that were targted by ≥ 4 miRNAs. **C**. Dot plot of WikiPathways functional enrichment. Number of genes targeted by each miRNA are shown in parentheses.

### Ad14 expressed miRNA targeting cell signaling pathways in macrophages

Our previous studies have shown that Ad14 CPE corpses are capable of repressing macrophage inflammatory responses[49]. The bioinformatics analysis performed above indicated that the enriched miRNA in Ad14 CPE corpses can target many cellular signaling pathways. Qiagen’s Ingenuity Pathway Analysis (IPA) software was used to determine which genes and signaling pathways in macrophages might be regulated by the Ad14 CPE enriched miRNAs. Initial analysis showed that the 10 miRNAs are predicted to regulate 6233 mRNAs (Fig 5A). After filtering the data for cell type (macrophages), pathways (cellular stress & injury, cytokine signaling, disease specific pathways and pathogen-influenced signaling) and confidence level (experimentally observed and highly predicted) 416 different mRNAs were predicted to be targeted for repression (Fig 5B and S3 Table). Let7a-5p is predicted to interact with the most targets (Fig 5C). Network analysis showed that there are 463 interactions for the 10 enriched miRNAs and their targets (S3 Fig).

**Fig 5.**
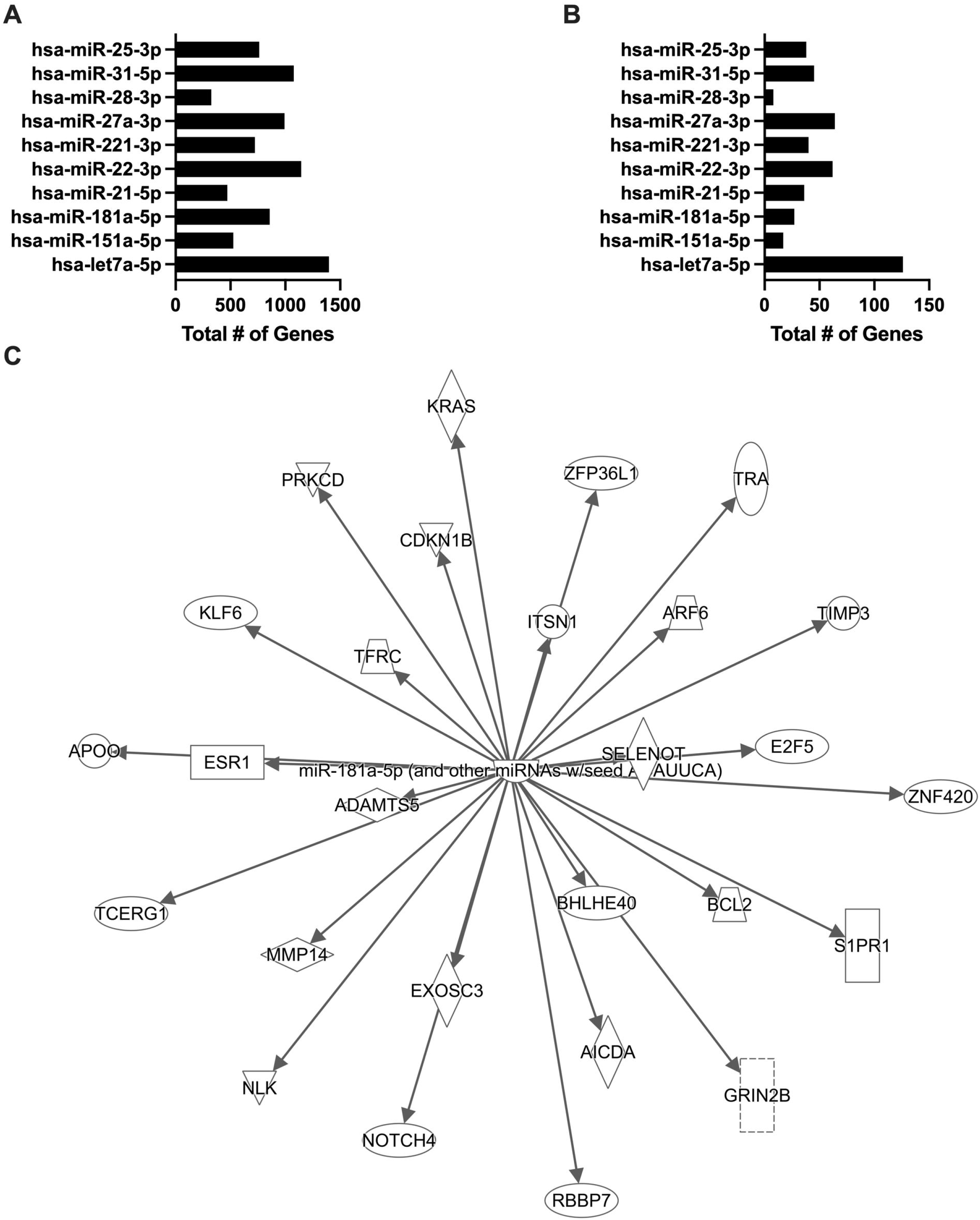
Ingenuity Pathway Analysis of Ad14 enriched miRNAs. **A&B**. IPA microRNA filter analysis was performed on the 10 Ad14 enriched miRNAs. The number of genes targeted by each miRNA is shown for unfiltered (A) and filtered (B) for cell type (macrophages), pathways (cellular stress & injury, cytokine signaling, disease specific pathways and pathogen-influenced signaling) and confidence level (experimental observed and high predicted). **C**. Visualization of the miR-181a-5p:mRNA target network from the filtered data.

### Ad14 miRNA targeted signaling pathways regulate inflammatory responses and acute lung injury

We have shown that Ad14 CPE corpses repress NF-κB dependent transcription in responder macrophages [49]. Of the 416 mRNAs predicted to be targeted in signaling pathways in macrophages, IPA showed that 21 proteins are involved in NF-κB signaling pathways and regulated by 8 of the Ad14 miRNAs (Fig 6). The targets for these miRNAs are upstream of NF-κB and collectively can repress NF-κB activation through numerous cell receptors. Other signal transductions pathways, such as mitogen activated protein kinase (p38 MAPK and ERK) pathways and the c-jun N-terminal kinase (JNK) pathway also lead to activation of NF-κB-dependent transcription and can drive expression of pro-inflammatory cytokines through both NF-κB dependent and independent transcription. IPA analysis showed that the ERK signaling pathway is regulated by 9 of 10 Ad14 miRNAs (S4 Fig) targeting proteins upstream and downstream of ERK including transcription factors that ERK activates. Likewise, 9 of the 10 miRNA target proteins in the p38 MAPK signaling pathway upstream and downstream of MAPK. MAPK11 (p38β) is directly targeted by miR-151-5p, miR-31-5p and let7a-5p, while MAPK14 (p38α) is directly targeted by miR-22-3p (S5 Fig). The JNK signaling pathway is targeted by 9 of the 10 Ad14 miRNAs, mainly upstream of MAP2K4/7 (S6 Fig). ERK, p38 and JNK pathways lead to the activation of multiple transcription factors that drive the expression of cytokines and chemokines that can be involved in the inflammatory response leading to acute lung injury (ALI). IPA showed that these transcription factors and NF-κB drive the expression of 15 cytokines and chemokines (Fig 7A). Further analysis revealed that 6 of Ad14 miRNAs can directly repress the activation of CXCL8, CXCL10, CCL2, CCL3, IL6, IL2, IL1β, IL10 and TNFα (Fig 7B). Repression of these chemokines and cytokines is predicted to decrease both ALI and ARDS.

**Fig 6.**
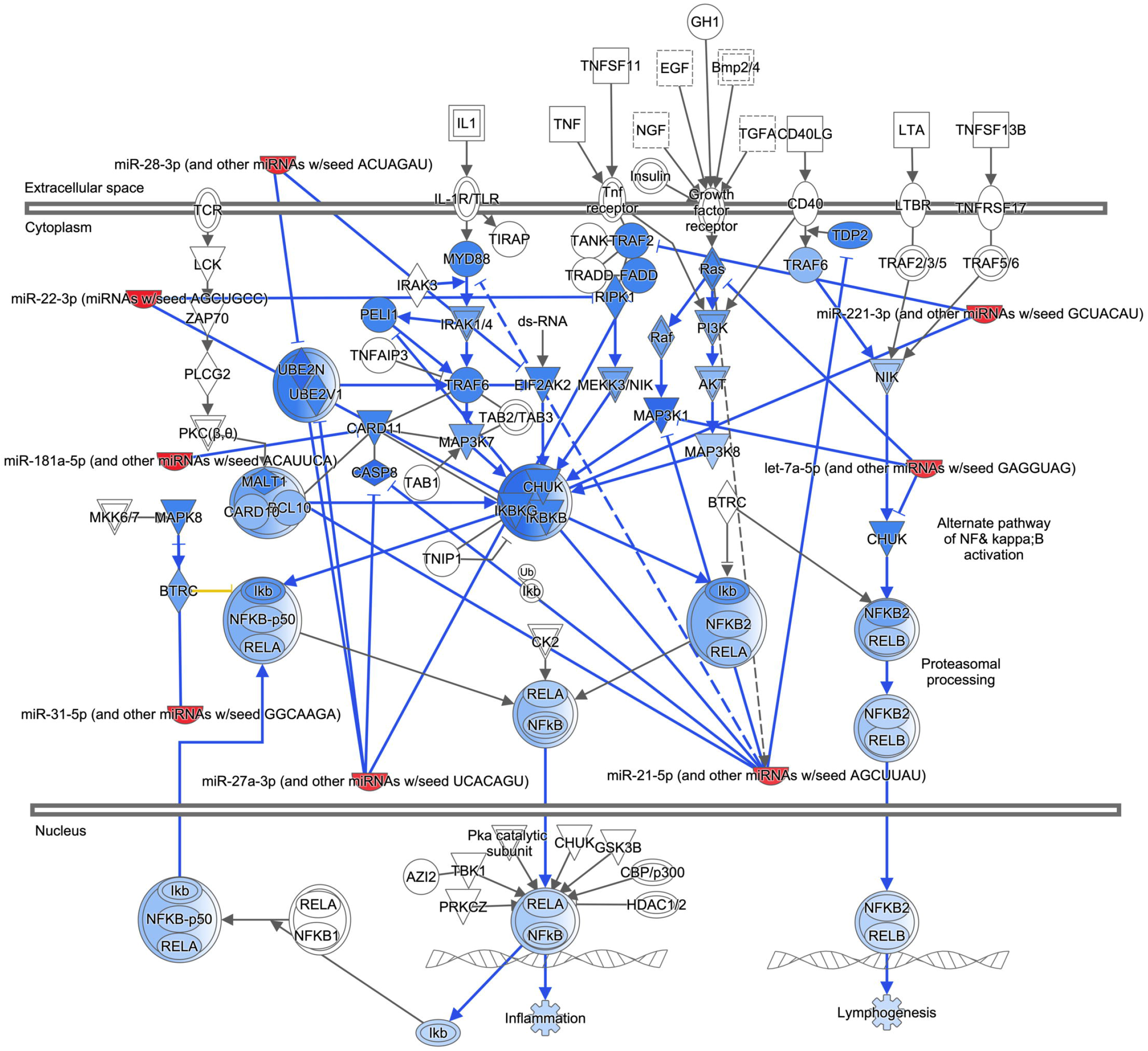
Effect of Ad14 miRNAs on NF-κB signaling pathway. IPA was used to predict the effects of Ad14 miRNAs on proteins involved in activating NF-κB dependent transcription. Lines ending with an arrowhead show the direction of activation, while lines ending with a dash show the direction of inhibition. Solid blue lines indicate validated repressive effects of miRNA-target interactions, while dashed blue lines are for predicted repressive effects of miRNA-target interactions. Red crescents are miRNAs, and the remaining shapes are proteins in the NF-κB pathway. Dark blue shapes are direct targets of the Ad14 miRNAs, while light blue shapes are predicted to have decreased activation based on repression of an upstream activator.

**Fig 7.**
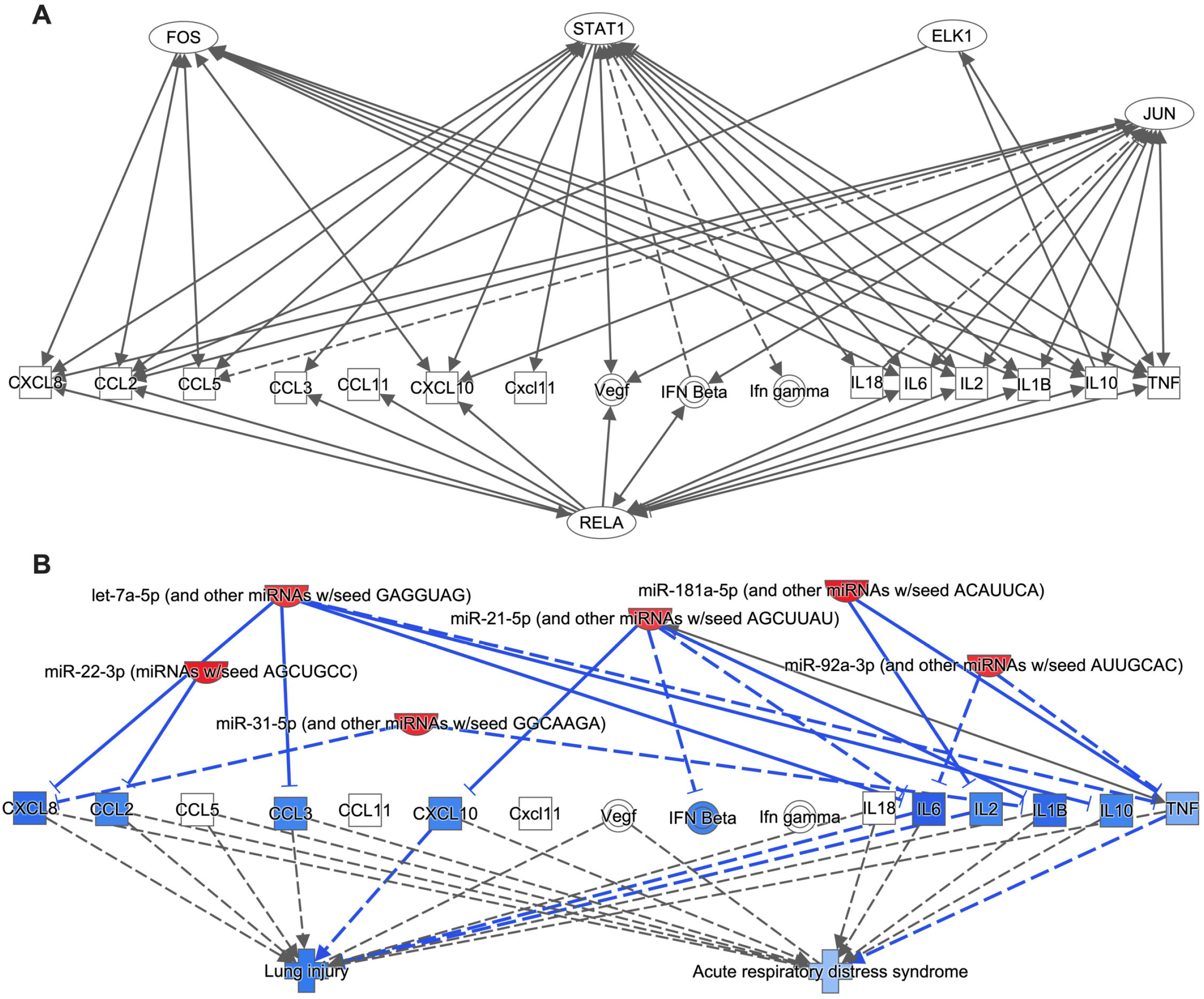
Bioinformatic analysis of regulation of chemokine and cytokine expression. **A & B**. IPA was used to probe the relationship of transcription factors (A) and Ad14 miRNA (B) on the expression of chemokines and cytokines involved in ALI and ARDS. Lines ending with an arrowhead show the direction of activation, while lines ending with a dash show the direction of inhibition. Solid blue lines indicate validated repressive effects of miRNA-target interactions, while dashed blue lines are for predicted repressive effects of miRNA-target interactions.

### Increased ALI/ARDS related cytokines and chemokines in the lungs of Ad14p1 infected Syrian hamsters compared to Ad14 infected hamsters

Our bioinformatic analysis would predict that there should be increased ALI/ARDS related cytokines and chemokines in the lungs of Ad14p1 infected hamsters compared to Ad14 infected hamsters. RNA-seq was to used to assess differential expression of cytokines and chemokines in the lungs of infected hamsters at day 5 post infection. Histopathology of lungs at 5 days post infection has shown the greatest difference in the inflammatory responses in between Ad14p1 and Ad14[47]. The lungs of Ad14p1 infected hamsters showed a significant increase in the expression of 17 cytokines/chemokines compared to Ad14 infected hamsters (Table 4). Eight of these chemokines/cytokines are predicted to be down regulated by the Ad14 miRNAs.

**Table 4.** Increased expression of cytokines and chemokines in the lungs of Ad14p1-infected vs. Ad14-infected Syrian hamsters^**a**^

| Gene | Log2 Fold Increase<sup>b</sup> | FDR p-value<sup>c</sup> |
| --- | --- | --- |
| CXCL11 | 5.146008 | 3.26E-12 |
| CXCL9 | 4.477917 | 4.35E-12 |
| CXCL10 | 4.101228 | 4.59E-11 |
| CCL7 | 4.250819 | 1.24E-10 |
| CCL5 | 2.892984 | 3.32E-09 |
| CCL2 | 3.429453 | 3.88E-09 |
| CCL8 | 2.801141 | 1.41E-07 |
| CCL4 | 2.784724 | 1.11E-05 |
| IFNG | 3.160325 | 1.31E-05 |
| IL10 | 3.692095 | 0.000147 |
| CXCL2 | 2.36665 | 0.000153 |
| CCL8 | 2.488043 | 0.000261 |
| CCL11 | 1.751661 | 0.000619 |
| CXCL6 | 2.46484 | 0.001136 |
| IL12B | 2.828453 | 0.024716 |
| CCL3 | 1.603861 | 0.039708 |
| IL21 | 7.092791 | 0.048245 |

<sup>a</sup> n=4 infected Syrian hamsters per virus
<sup>b</sup> Log<sub>2</sub> fold increase in mRNA expression induced by infection with Ad14p1 vs Ad14
<sup>c</sup> Benjamini-Hochberg test for false discovery rate

## Discussion

The majority of adenovirus infections in healthy individuals are mild and self-limiting. However, with the increased surveillance for respiratory virus infections by PCR over the last 20 years, there has been increased detection of outbreaks of severe respiratory Ad infections in healthy patients, some even resulting in ALI/ARDS. Ad14p1 is a strain of Ad that can induce ALI/ARDS in otherwise healthy patients. We have shown that, although Ad14 and Ad14p1 are 99.9% genetically identical, they have different effects on the host inflammatory response. Ad14p1 infection of Syrian hamsters results in an ALI-like response, whereas Ad14 infected hamsters show minimal lung inflammation[47]. Ad14 CPE corpses repress macrophage pro-inflammatory cytokine expression, while Ad14p1 CPE corpses fail to repress these pro-inflammatory responses because of reduced expression of the Ad protein, E1B 20K[49]. Similar results were also observed with a deletion mutant of Ad5 that lacks E1B 19/20K expression[48]. The objective of these studies was to determine whether differential expression of cellular miRNA between Ad14 and Ad14p1 CPE corpses occurs and whether those differentially expressed miRNA control Ad14 CPE corpse immunomodulation of macrophage inflammatory responses.

Infection of lung cell lines with human Ad (Ad2 or Ad3) alters cellular miRNA expression[25,26]. Here we infected human lung A549 cells with Ad14 and examined cellular miRNA expression using small RNA-seq. Our results (Table 1) show that infection with Ad14 or Ad14p1 results in a decrease in the total cellular miRNA reads over the course of infection, which is consistent with what has previously been observed[25]. The overall decrease in cellular miRNA reads at and beyond 12hpi is associated with the processing of Ad14 VA RNA I into its mivaRNAs (Table 1). At 48hpi, the total cellular miRNAs are further reduced in Ad14p1 infected cells compared to Ad14 infected cells. This is consistent with our previous findings that, at 48hpi, there are more Ad14 mivaRNAs present in Ad14p1 infected cells than in Ad14 infected cells[55]. Zhao and colleagues reported that, during Ad2 infection, the majority of the significantly deregulated miRNA are repressed[25]. In contrast, our data (Table 2) show that, throughout Ad14 and Ad14p1 infection through complete cytopathic effect, roughly the same number of miRNAs are up-regulated or down-regulated. This difference is most likely explained by the fact that our analysis included more miRNAs, since we allowed a lower fold change and did not restrict miRNAs based on overall expression levels. Both Ad14 and Ad14p1 infection resulted in the same degree of miRNA deregulation at all time points, except for 48hpi when Ad14p1 deregulated nearly twice as many (315 vs 173) cellular miRNAs (Table 2). Comparison of the miRNAs deregulated showed that Ad14 and Ad14p1 deregulated the same miRNAs at each time point (Fig2) except at 48hpi. The reason for this difference at 48hpi is unclear. The only molecular difference we have identified so far, between Ad14 and Ad14p1 infected cells is a marked reduction of Ad E1B 20K mRNA and protein in infected cells. Whether this difference in viral gene expression is the cause of the differential expression of cellular miRNA requires further study[49]. Expression differences in other viral genes between Ad14 and Ad14p1 would also have to be considered as possible factors in alterations in cellular miRNA expression. For example, E1A expression has been reported to down-regulate the expression of miR-27a, miR-520h, miR-7b and miR-197 in breast cancer cell lines[56].

Macrophages play a key role in removing cell corpses from sites of inflammation through efferocytosis. This process allows for the transfer of proteins, lipids and nucleic acids from the dying cellular corpses to macrophages[20,57,58]. It has been demonstrated that miRNA delivered from either exosomes or apoptotic corpses to macrophages can alter macrophage-mediated inflammatory responses, through repression of NF-κB dependent cytokine expression and the transition of M1 (pro-inflammatory) to M2 (anti-inflammatory) macrophages[17,20,59–66]. We have reported that Ad5 or Ad14 CPE corpses repress both NF-κB dependent transcription induced by PMA and pro-inflammatory cytokine expression induced by either LPS/IFNγ or Ad viral particles in macrophages. In contrast, Ad CPE corpses dying as the result of infection with adenoviruses that lack sufficient expression of E1B 20K (such as Ad14p1) fail to repress those same cellular functions[48,49]. However, the mechanism through which Ad CPE corpses convey that immunomodulatory activity is unknown. GO, KEGG and functional enrichment analysis (Fig 3 and 4) revealed that the 10 miRNAs enriched in Ad14 CPE corpses target genes that are involved in many signal transduction pathways that induce pro-inflammatory cytokine expression. Restricting IPA analysis to known and strongly predicted miRNA-protein interactions in macrophage/monocytes revealed that 8 of the 10 Ad14 miRNAs targeting 12 different proteins with 4 of those targeted by more than 1 miRNA in the NF-κB signaling pathway (Fig6). The Ad14 miRNAs also target proteins that result in the inhibition of ERK, JNK and p38 (S4-6 Fig) signal transduction pathways that activate FOS, JUN, ELK1, STAT1 and RELA transcription factors (Fig7A), which drive pro-inflammatory cytokine expression[64,67–76].

Prolonged acute inflammatory response during ALI can result in ARDS. Alveolar macrophages are long-lived resident macrophages that constitute >85% of the leukocytes in airspaces during ALI and, as such, are the first line of defense in the immune response to infection[77–80]. Many cytokines and chemokines produced by alveolar macrophages drive the progression of ALI/ARDS[81–87]. IPA analysis showed that 6 of the Ad14 miRNAs repress 10 of the cytokines/chemokines that are regulated by transcription factors regulated by the Ad14 miRNAs, and 9 of those cytokines/chemokines drive ALI/ARDS (Fig 7B). RNA-seq analysis showed that 8 cytokines/chemokines involved in ALI/ARDS are upregulated in the lungs at day 5 post infection of Syrian hamsters infected with Ad14p1 compared to Ad14 infected hamsters (Table 4). These results are consistent with our previous observations using qPCR on a limited number of cytokines/chemokines[47]. The possible relationship between reduced expression of the Ad14 miRNA during Ad14p1 infection and macrophage-mediated inflammatory responses is speculative. However, others have reported that many of the miRNA associated with Ad14 infection can regulate inflammatory responses, ALI/ARDS and other lung inflammatory diseases. For example, increased expression of miR-181a-5p decreases NK-κB activation and alleviates inflammatory responses in COPD, whereas miR-181a-5p inhibition contributes to macrophage M1 polarization[88,89]. Increased expression of miR-27a-3p decreases expression of TNFα and IL6, while increased expression of let-7a-5p decreases expression of TNFα, IL6 and IL1β[73,90]. Studies by Das and colleagues has shown that miR-21-5p in apoptotic bodies engulfed by macrophages induces an M2-like macrophage phenotype associated with repressed NK-κB activation and cytokine expression[19]. Other reports indicated that miR-21-5p can inhibit LPS induced inflammation in ulcerative colitis[70,71,91]. MiR-22-3p has been reported to be involved in asthma, attenuating airway destruction and tissue damage and attenuating ALI[92,93]. Overall, it is increasingly apparent that miRNAs from extracellular vesicles (including apoptotic bodies) can regulate macrophage functions and might be targets for the design of therapeutic agents [60,66,94–97].

In this report we examined the expression of cellular miRNA during infection with two pathogenically distinct strains of Ad14 – Ad14 deWit (prototype Ad14) and the derivative outbreak strain, Ad14p1. Our data showed that, despite being over 99.9% genetically identical, Ad14 and Ad14p1 infections have distinct effects on expression of cellular miRNAs. The bioinformatic analysis of the miRNAs enriched in Ad14 CPE corpses help explain the previously observed repression of inflammatory cytokine expression in macrophages that have engulfed Ad14 CPE corpses. In contrast, the decreased expression of Ad14 miRNAs in Ad14p1 CPE corpses might explain the increased inflammatory responses following efferocytosis of Ad14p1 CPE corpses by macrophages.

Based on our data and the literature, we propose the following model to explain the divergent, macrophage-mediated inflammatory lung responses during Ad14 and Ad14p1 infection (Fig 8). Engulfment by macrophages of Ad14 CPE corpses enriched in the Ad14 miRNAs results in decreased inflammatory cytokine/chemokine expression during Ad14 infection, limiting ALI/ARDS as a result of the collective actions of the Ad14 miRNAs to repress inflammatory gene expression. Conversely, efferocytosis of Ad14p1 CPE corpses, which express less Ad14 miRNAs, results in increased macrophage inflammatory responses to viral infection and the observed ALI/ARDS-like response, since there are insufficient levels of Ad14 miRNA to repress inflammatory gene expression. We propose that the Syrian hamster model of comparative lung immunopathogenesis between Ad14 and Ad14p1 can be used to understand the roles of miRNAs in resolving or promoting ALI/ARDS, the viral genetic basis of miRNA regulation and the possible basis for development of a strategy for therapeutic delivery of miRNAs to treat virally induced ALI/ARDS.

**Fig 8.**
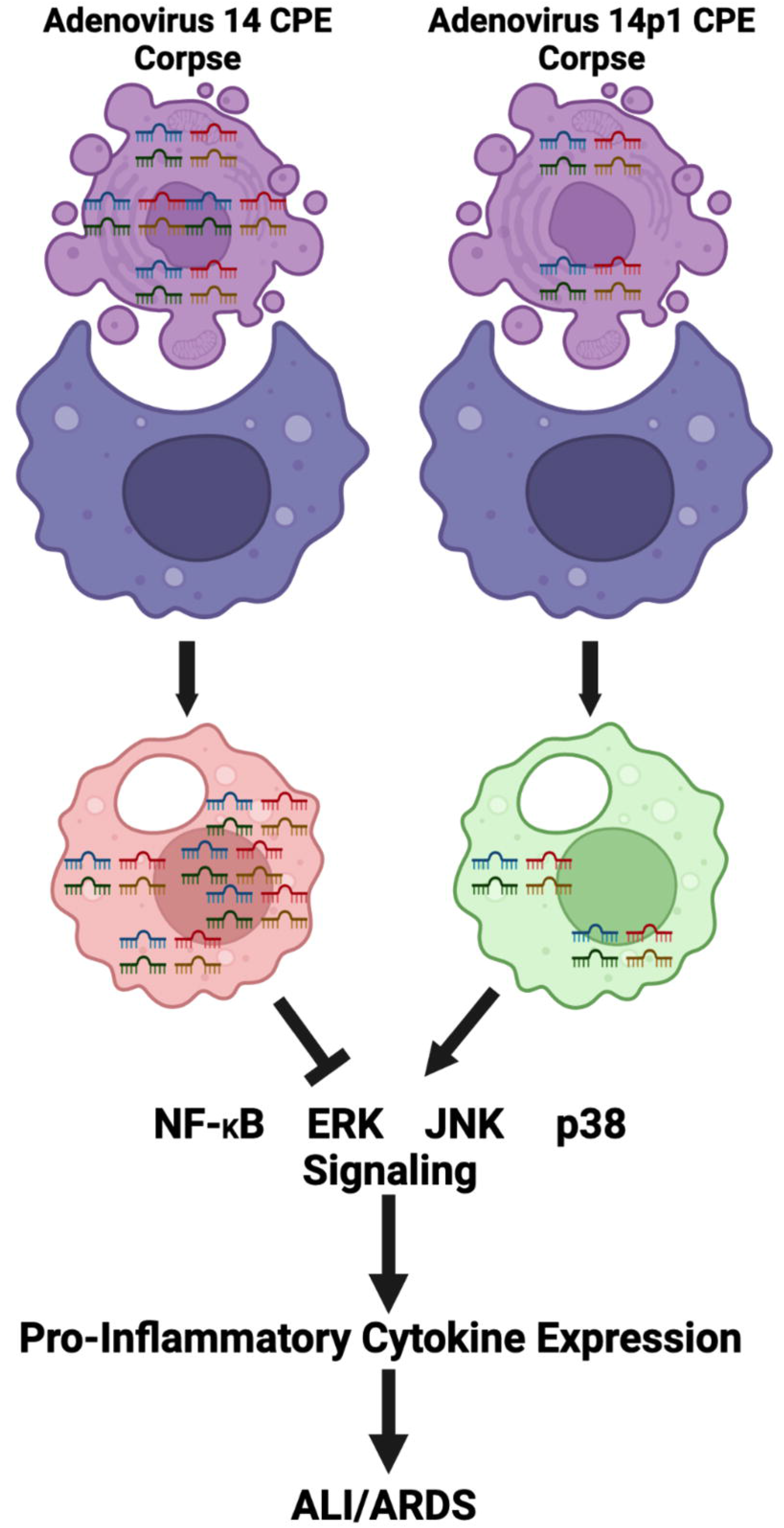
Model of Ad14/Ad14p1 CPE corpse modulation of ALI/ARDS. As Ad14 or Ad14p1 infected cells die, the resulting Ad CPE corpses are phagocytized by macrophages. This results in the transfer of Ad CPE corpse miRNAs to the macrophage, altering the macrophage inflammatory response according the array of miRNAs present in the corpses. MiRNAs from Ad14 CPE corpses decrease activation of macrophage signaling pathways that lead to the production of pro-inflammatory chemokines and cytokines, preventing induction of ALI/ARDS in Syrian hamsters infected with Ad14. The lower level of immunoregulatory miRNAs in Ad14p1 CPE corpses results in failure to repress virally triggered, pro-inflammatory pathways in macrophages, resulting in increased pro-inflammatory chemokine and cytokine expression and the associated ALI/ARDS responses seen in hamsters infected with Ad14p1.

## Methods and Materials

### Ethics statement

Studies were conducted in strict accordance with recommendations of the Guide for the Care and Use of Laboratory Animals of the National Institutes of Health and approved by the Boise, VA (RDIS#0001; 9/25/17 to present) IACUC committees. Hamsters were anesthetized with 2.5% isoflurane delivered in a stream of oxygen by a controlled precision vaporizer. Hamsters were euthanized using CO_2_ in accordance with American Veterinary Medical Association guidelines.

### Cells and viruses

A549 cells (CCL-185, ATCC, Manassas, VA) were maintained in DMEM and grown at 37°C and 5% CO_2_ as described previously[48,49]. A549 cells were validated by short tandem repeat markers (STR, ATCC) and monitored for mycoplasma contamination by PCR (ATCC). HAd14 deWit (VR-15) was obtained from ATCC and HAd14p1 (1986T) was obtained from Kevin Russel (United States Naval Health Research Center, San Diego, CA)[46]. Viruses were propagated in A549 cells and plaque titered in A549 cells.

### Infection of A549 cells and isolation of total cellular RNA

A549 cells were infected with either Ad14 or Ad14p1 at a MOI of 10 pfu/cell in suspension for 1 hr at 37°C, after which cells were plated and allowed to adhere until collected. Adherent and non-adherent cells were collected at 6, 12, 24, 36 and 48 hr post-infection. Total RNA was isolated using miRNeasy kit (Qiagen, Germantown, MD) with on-column DNase treatment. The total RNA in each sample was quantified using the Qubit 2.0 Fluorometer (Invitrogen, Carlsbad, CA), and quality was measured using the RNA6000 Nano chip on the Agilent 2100 Bioanalzyer (Agilent Technologies, Santa Clara, CA). Samples with an RNA integrity number (RIN) greater than 8 were used for sequencing.

### miRNA library preparation, sequencing and initial data processing

Sequencing libraries were generated using the TruSeq Small RNA library prep kit (Illumina, San Diego, CA). The libraries were size selected using a 6% polyacrylamide gel and concentrated using ethanol precipitation. Purified libraries were normalized and pooled to create a double stranded cDNA library ready for sequencing. The samples were sequenced on the Illumina MiSeq to render 50 base pair single end reads. Adapter sequences were removed, and low quality reads were trimmed from raw sequencing reads using Cutadapt (v. 1.11) and samples demultiplexed.

### Human miRNA data analysis

CLC Workbench 22 (Qiagen) was utilized for further data analysis. Quantify miRNA 1.3 was used to quantify miRNA counts allowing for 2 additional upstream or downstream bases, 2 missing upstream or downstream bases and with no mismatches. Reads were annotated with miRbase 22 that also included HAdV14 mivaRNA and a piRNA database 1.7.6 (add ref to VA RNA paper once submitted to BioRx). Results were grouped based on mature and seed sequences. Principle component analysis (PCA) was performed with PCA for RNA-seq 1.3 with data grouped on seed sequence. Heat map was conducted with Create Heat Map for RNA-seq 1.5 with data grouped on seed sequence. Samples were clustered based on Euclidean distance and average linkage with 30 features.

### Differential expression analysis

Differential Expression for RNA-Seq 2.7 was used, using small RNA option on samples grouped on seed sequence. Normalization was performed with trimmed mean of M values. Comparisons were made between all groups using the Wald test. False Discovery Rate (FDR) p-values were determined by the Benjamini Hochberg method. Maximum average of group RPKM values were determined between groups testing for differential expression based on viral strain. Differential expression of miRNA was considered significant if the FDR was ≤ 0.05. Venn diagram for RNA-seq 0.2 was used to create the Venn diagrams of significant differentially expressed miRNAs.

### Bioinformatics analysis

KEGG and GO analysis was performed with MirPath V.3 (05/07/22) using Fishers Exact Test with a p-value threshold of 0.05 with FDR correction on the 10 enriched miRNA in Ad14 CPE corpses[53]. Dotplots were made using ggplot with the MirPath data. Heatmaps were downloaded from MiRPath. Mienturnet (http://userver.bio.uniroma1.it/apps/mienturnet/) was used on 05/11/22 for target enrichment analysis[54]. TargetScan and miRTarBase were used for miRNA-target enrichment with a threshold of a minimum of 2 miRNA-target interactions and a threshold adjusted p-value of 1. Network analysis of the miRNA-targets was done using miRTarBase and allowing for both strong and weak interactions. Functional enrichment was done using miRTarBase with KEGG, REACTOME and WikiPathways databases. The miRNA Target Filter and Network/My Pathways were generated through Qiagen Ingenuity Pathway Analysis[98]. Briefly, Ad14 miRNAs were uploaded with expression data to IPA. Results were filtered by (1) cells/immune cells/macrophages, (2) Pathways to include Cellular Stress and Injury, Cytokine Signaling, Disease Specific Pathways and Pathogen Influenced Signaling and (3) Confidence to include experimentally observed and highly predicted. Interaction of Ad14 miRNAs on signaling pathways were generated by overlaying the miRNA express data on IPA Canonical Pathways.

### Viral infection of Syrian hamsters and RNA-seq analysis

Four to six-week-old female Syrian hamsters (*Mesocricetus auratus*) were obtained from Charles River (Wilmington, MA, USA) and had access to food and water ad libitum. Hamsters were allowed to acclimate for 72 h after shipping, before experimental use. Hamsters were infected with 5×10^9^ genomes of Ad14 or Ad14p1 as previously described[47]. Briefly, hamsters were sedated with isoflurane to induce slower, deeper respiration and held in an upright but slightly recumbent position to allow infection via intratracheal aspiration during panting respiration. Hamsters were euthanized at 5 days post infection and the left lobes were removed. Lobes were homogenized with TissueRupor II (Qiagen) according to recommendations for total RNA extraction using RNeasy (Qiagen). All RNA samples had RIN > 8. Total RNA was used as the input for KAPA mRNA HyperPrep and indexes were added for Illumina Sequencing. Libraries were pooled and sequenced on an Illumina NextSeq550. CLC Genomics Workbench (Qiagen) was used to map reads to the Syrian hamster genome (BCM_Maur_2.0 (GCF_017639785.1) and for differential gene expression between Ad14 and Ad14p1 infected hamsters. False Discovery Rate (FDR) p-values were determined by the Benjamini Hochberg method.

## Supporting Information

**S1 Table. Differentially expressed miRNA during Ad14 and Ad14p1 infection.**

**S2 Table. MirPath predicted targets for Ad14 miRNAs.**

**S3 Table. IPA predicted targets for Ad14 miRNAs.**

**S1 Fig. Heatmaps of KEGG and GO analysis of Ad14 miRNAs**. The Ad14 miRNAs were uploaded to mirPath V3. **A-C**. Heatmaps of the KEGG Pathways Union (A), KEGG Genes Union (B) and GO Categories Union (C) analysis.

**S2 Fig. Dotplots of KEGG and Reactome functional enrichment of Ad14 miRNAs. A&B**. The 10 enriched miRNAs in Ad14 CPE corpses were uploaded to MIENTURNET. Targent enrichment was performed with miRTarBase with a minimum of 2 miRNA-RNA interactions and an FDR threshold of 1. Dotplots of KEGG (A) and Reactome (B) functional enrichment. Number of genes targeted by each miRNA are shown in parentheses.

**S3 Fig. IPA network analysis of Ad14 miRNA and predicted targets**. IPA microRNA filter analysis was performed on the Ad14 miRNA. Network analysis of miRNA and targets filtered for cell type (macrophages), pathways (cellular stress & injury, cytokine signaling, disease specific pathways and pathogen-influenced signaling) and confidence level (experimental observed and high predicted).

**S4 Fig. Effect of Ad14 miRNAs on ERK signaling pathway**. IPA was used to predict the effects of Ad14 miRNAs on proteins involved in the ERK signaling pathway. Lines ending with an arrowhead show the direction of activation, while lines ending with a dash show the direction of inhibition. Solid blue lines indicate validated repressive effects of miRNA-target interactions, while dashed blue lines are for predicted repressive effects of miRNA-target interactions. Red cresents are miRNAs, and the remaining shapes are proteins in the ERK pathway. Dark blue shapes are direct targets of the Ad14 miRNAs, while light blue shapes are predicted to have decreased activation based on repression of an upstream activator.

**S5 Fig. Effect of Ad14 miRNAs on p38 MAPK signaling pathway**. IPA was used to predict the effects of Ad14 miRNAs on proteins involved in p38 MAPK signaling. Lines ending with an arrowhead show the direction of activation, while lines ending with a dash show the direction of inhibition. Solid blue lines indicate validated repressive effects of miRNA-target interactions, while dashed blue lines are for predicted repressive effects of miRNA-target interactions. Red cresents are miRNAs, and the remaining shapes are proteins in the p38 MAPK pathway. Dark blue shapes are direct targets of the Ad14 miRNAs, while light blue shapes are predicted to have decreased activation based on repression of an upstream activator.

**S6 Fig. Effect of Ad14 miRNAs on JNK signaling pathway**. IPA was used to predict the effects of Ad14 miRNAs on proteins involved in JNK signaling. Lines ending with an arrowhead show the direction of activation, while lines ending with a dash show the direction of inhibition. Solid blue lines indicate validated repressive effects of miRNA-target interactions, while dashed blue lines are for predicted repressive effects of miRNA-target interactions. Red cresents are miRNAs, and the remaining shapes are proteins in the JNK pathway. Dark blue shapes are direct targets of the Ad14 miRNAs, while light blue shapes are predicted to have decreased activation based on repression of an upstream activator.

## Acknowledgements

We thank Drs. David Metzgar and Adriana Kajon for sharing emergent HAdV14p1 isolates. The authors are indebted to Dr. James L. Cook for his continued support of our research through thoughtful discussions and critical reading of this manuscript. We appreciate the efforts and hard work of Drs. Dennis Stevens, Amy Bryant, Carolyn Bohach and Sam Minnich for providing research opportunities for undergraduate students in underfunded states such as Idaho and Nevada through their respective COBRE (D.S. and A.B) and INBRE (C.B. and S.M.) grants as mentioned above.

## Notes

### Competing Interest Statement

The authors have declared no competing interest.

