## Supplementary figures and images for "Adenovirus 14p1 induced changes in miRNA expression increases lung immunopathogenesis"

### Supplemental Figure 1a

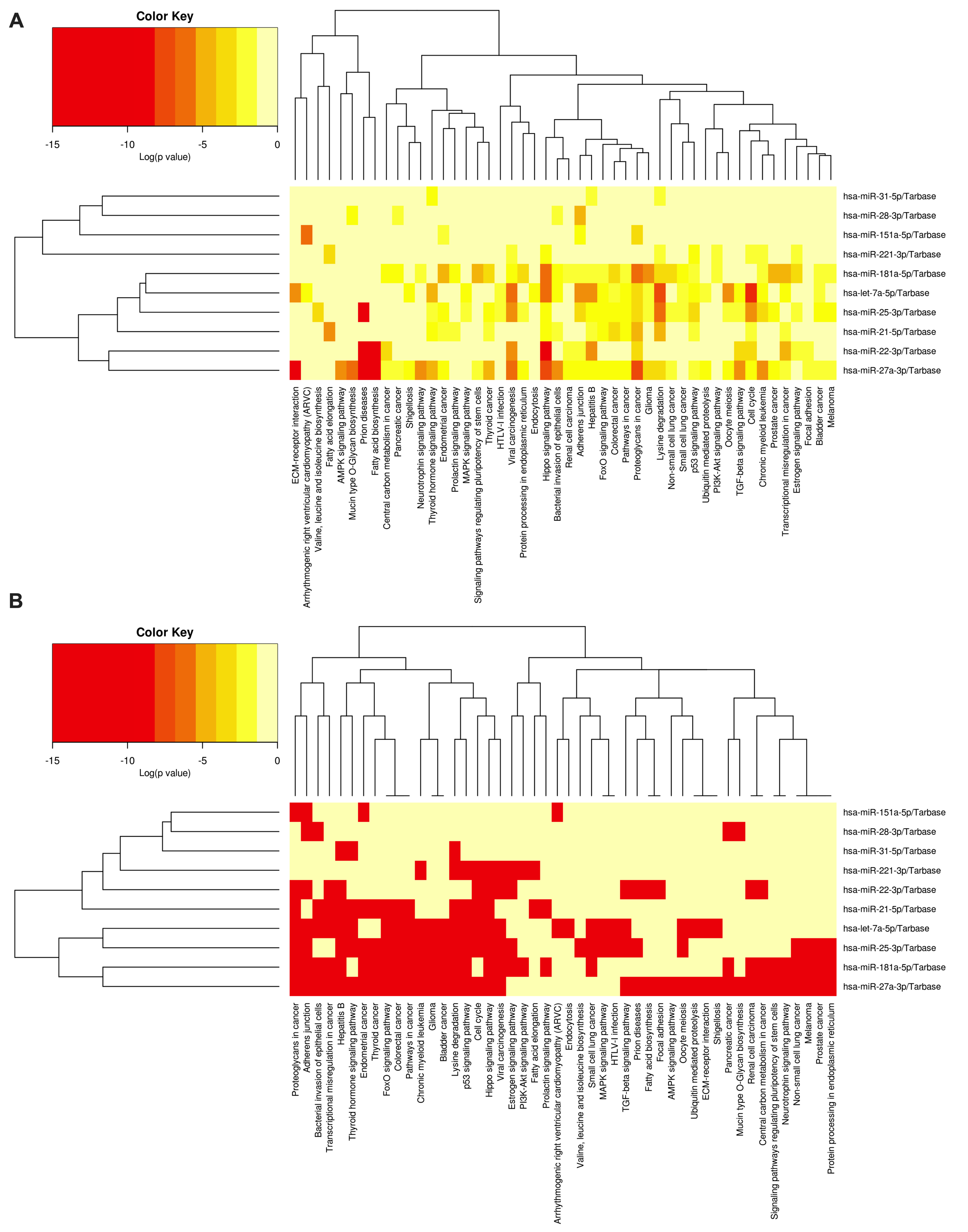

### Supplemental Figure 1b

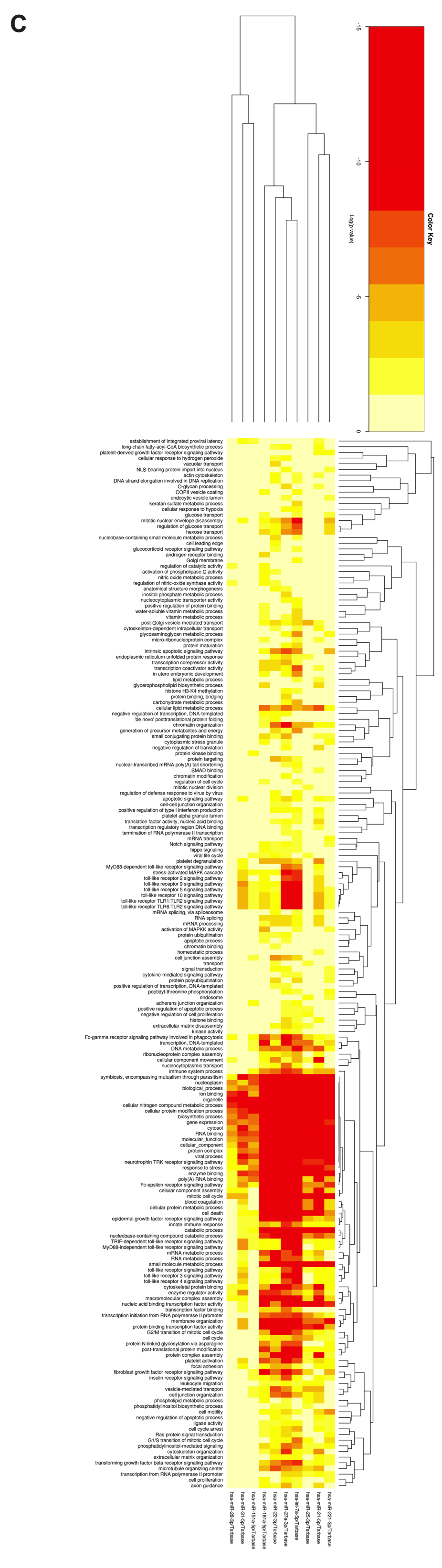

### Supplemental Figure 2

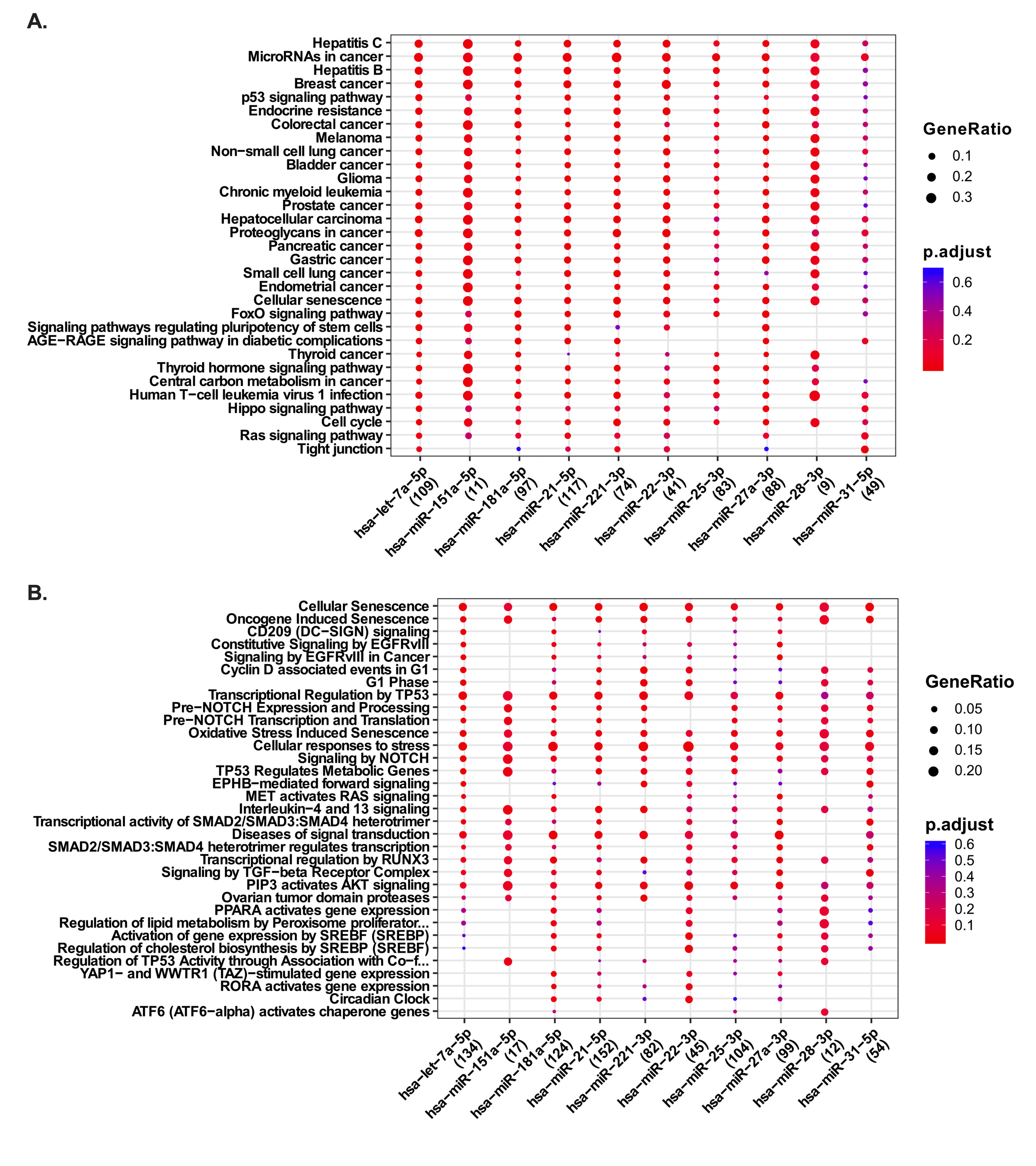

### Supplemental Figure 3

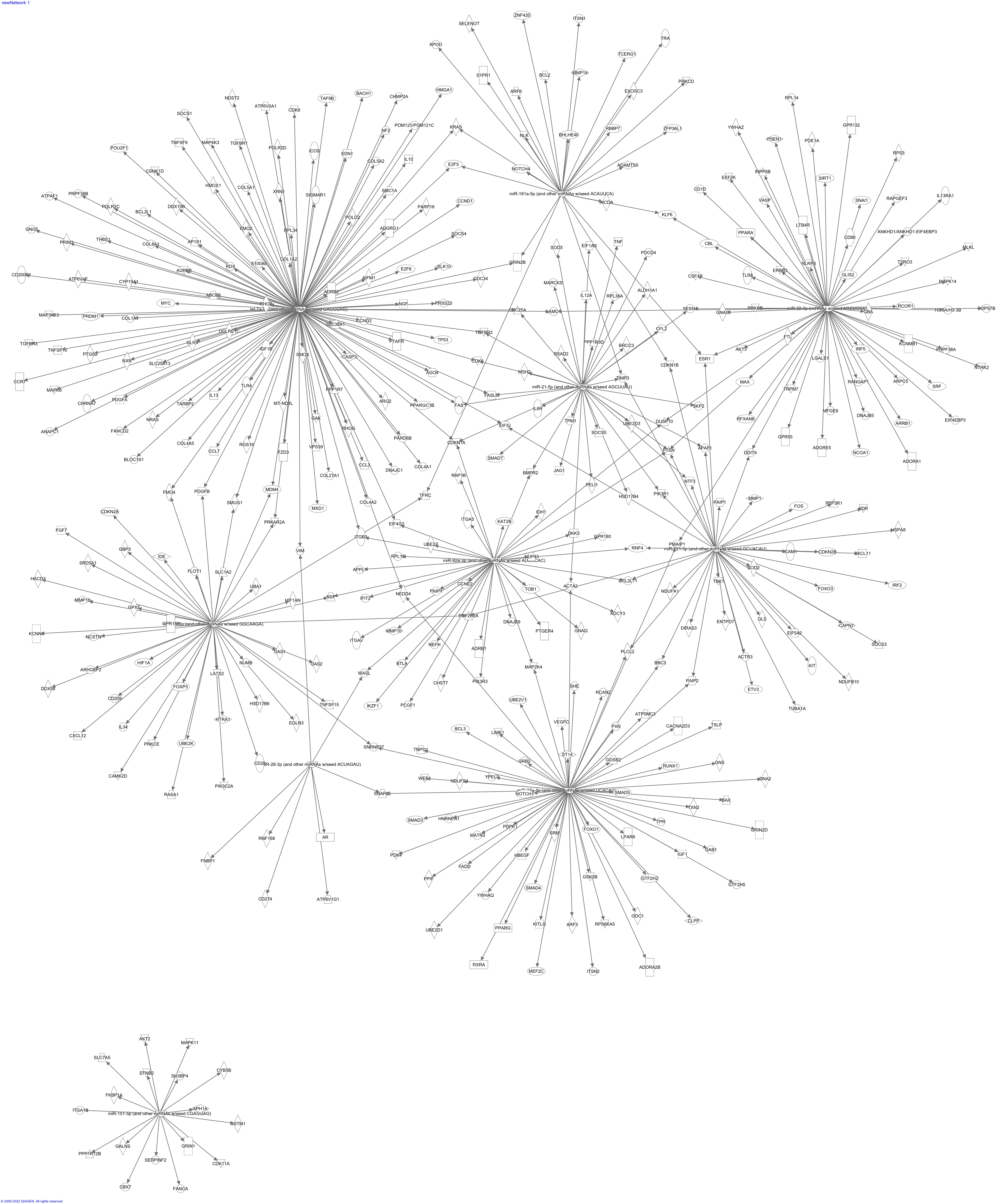

### Supplemental Figure 4

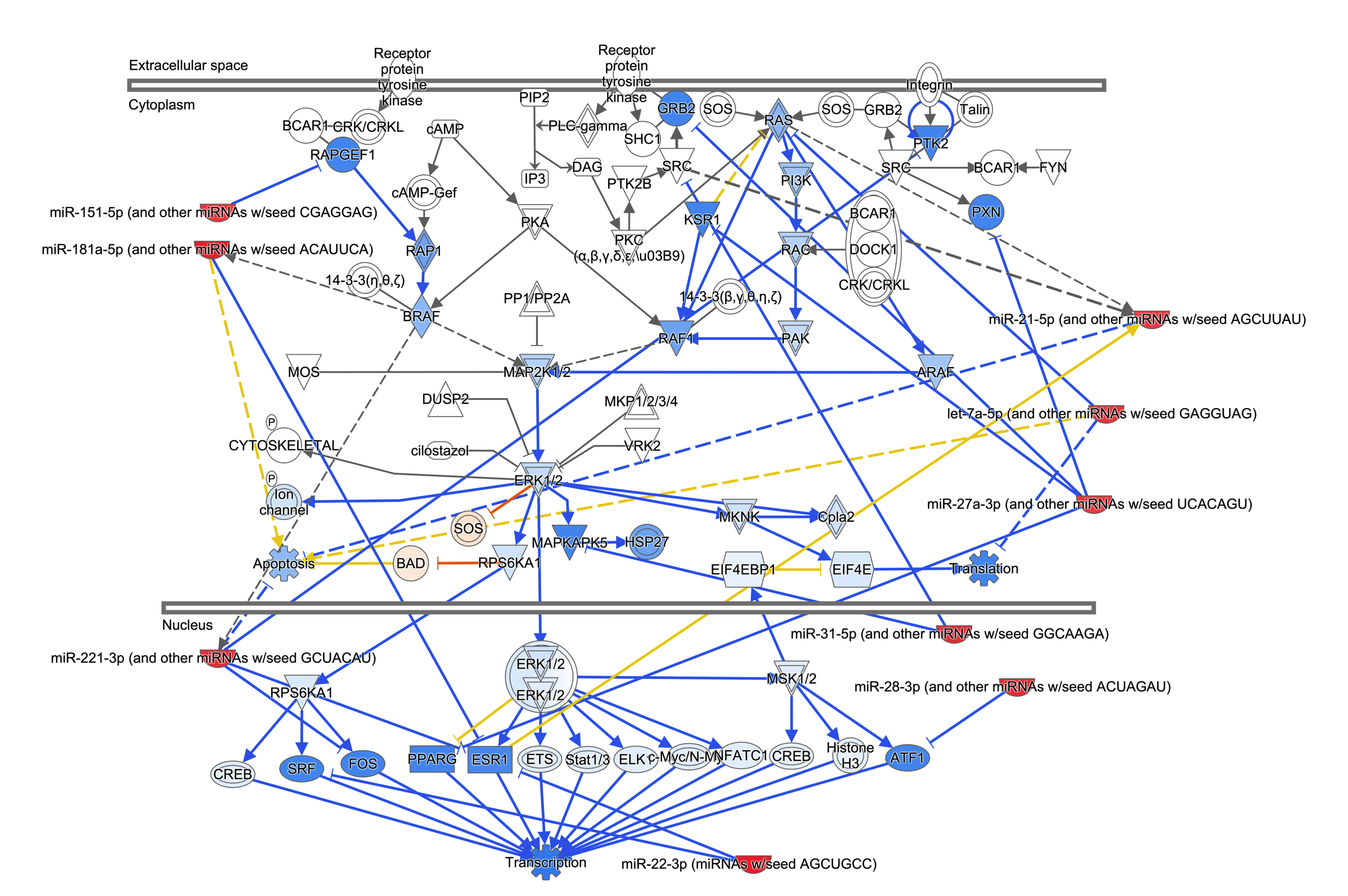

### Supplemental Figure 5

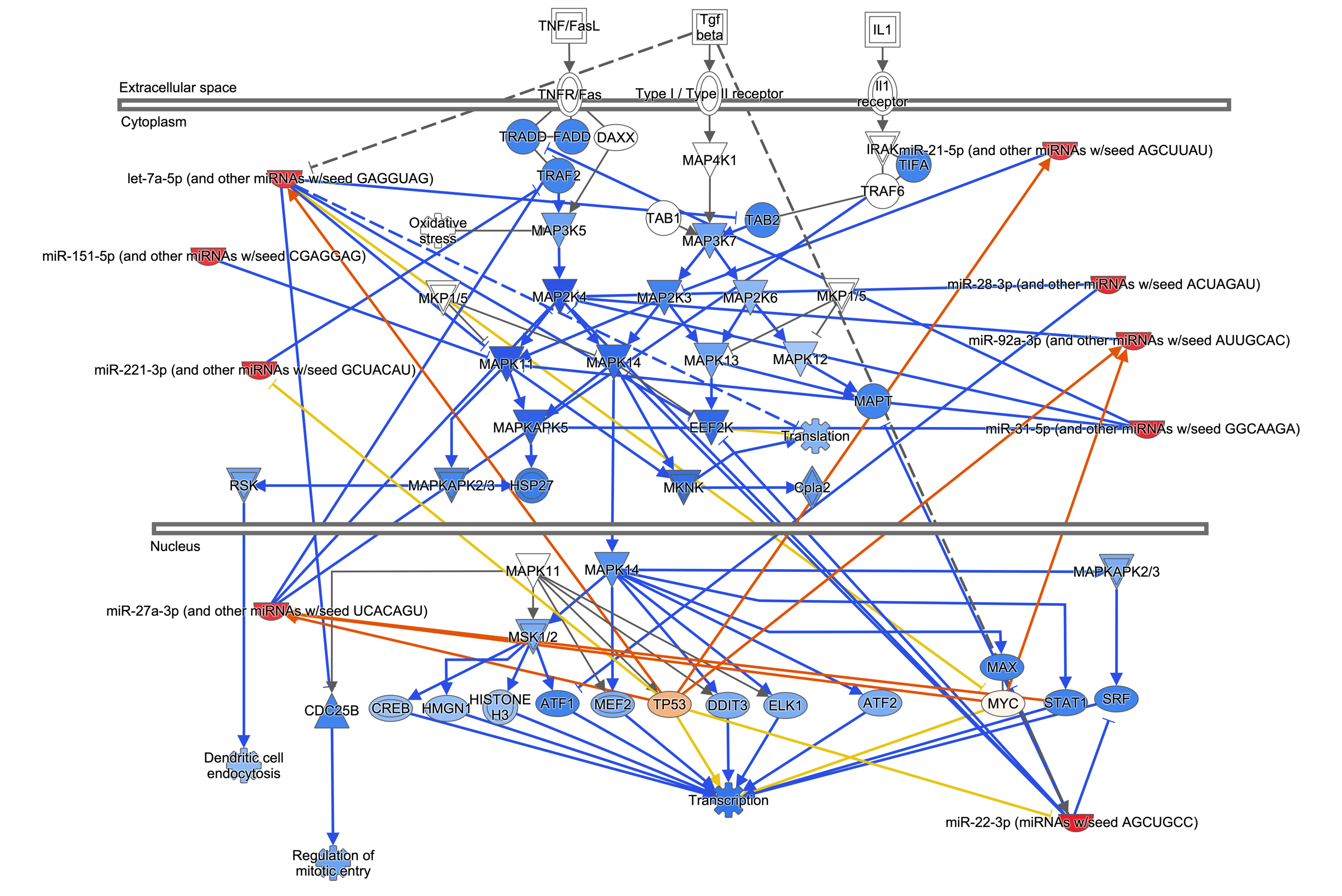

### Supplemental Figure 6

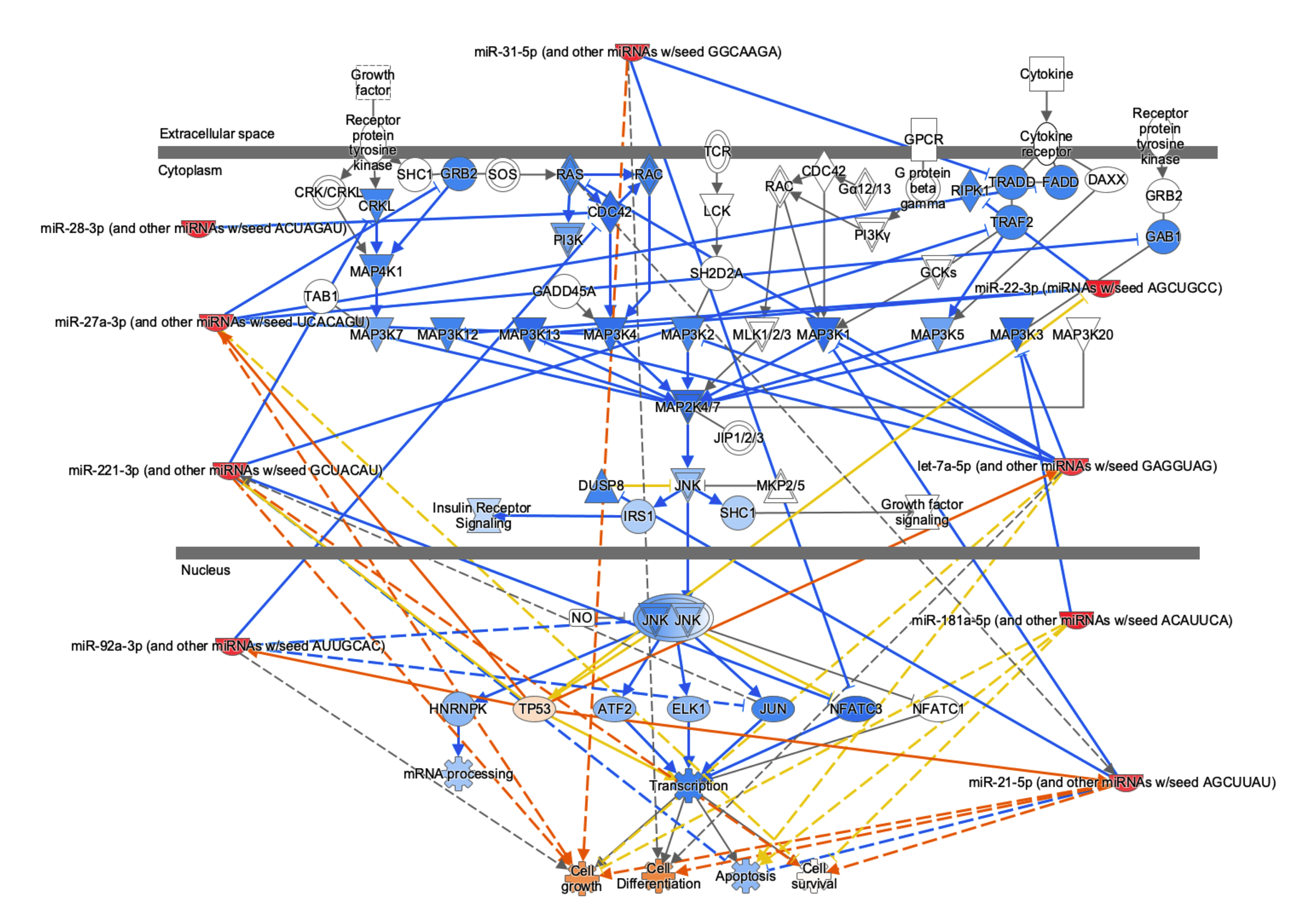
