## Supplemental Table 1 for "Adenovirus 14p1 induced changes in miRNA expression increases lung immunopathogenesis"

# Ad14 v C 6HPI     Ad14 v C 12H

| Identifier | Max group mean | Fold change | FDR p-value | Identifier | Max group mean | Fold change |
|---|---|---|---|---|---|---|
| hsa-miR-4490 | 7.75 | 21.29360261 | 0.002705967 | hsa-miR-684 | 7.5 | 61.3765228 |
| hsa-miR-3148 | 5.75 | 15.89349692 | 0.008746508 | hsa-miR-105 | 6.5 | 53.3198831 |
| hsa-miR-6799-5p | 6.5 | 10.61360467 | 0.003258432 | hsa-miR-375 | 7.25 | 20.5183667 |
| hsa-miR-3125 | 7 | 8.107721216 | 0.002094532 | hsa-miR-466 | 6.75 | 19.126241 |
| hsa-miR-155-5p | 3.75 | 6.222963807 | 0.039794513 | hsa-miR-155 | 9.75 | 16.6090772 |
| hsa-miR-493-3p | 9.5 | 5.851009 | 0.000368338 | hsa-miR-314 | 5 | 14.2587652 |
| hsa-miR-5187-3p | 23.5 | 4.31476471 | 7.48531E-08 | hsa-miR-557 | 3.75 | 10.7832213 |
| hsa-miR-369-3p | 8.25 | 3.890615923 | 0.005193682 | hsa-miR-427 | 10 | 9.58821453 |
| hsa-miR-191-3p | 6.25 | 3.879746152 | 0.013662149 | hsa-miR-679 | 5 | 8.64474076 |
| hsa-miR-493-5p | 18 | 3.4719067 | 1.83169E-05 | hsa-miR-520 | 12 | 7.90049093 |
| hsa-miR-409-3p | 96.25 | 3.179489594 | 1.6878E-19 | hsa-miR-119 | 11.5 | 7.5711708 |
| hsa-miR-152-5p | 8.75 | 3.039654065 | 0.012599283 | hsa-miR-888 | 15 | 7.54256363 |
| hsa-miR-411-5p | 141.5 | 3.03281733 | 6.12141E-22 | hsa-miR-472 | 4 | 6.93765675 |
| hsa-miR-1321 | 8.75 | 3.028666524 | 0.013774002 | hsa-miR-312 | 5 | 6.1763477 |
| hsa-miR-369-5p | 6.25 | 2.963453556 | 0.037776995 | hsa-miR-443 | 11.75 | 4.79623688 |
| hsa-miR-889-3p | 30.75 | 2.91809768 | 3.41365E-07 | hsa-miR-125 | 11.25 | 4.19775511 |
| hsa-miR-136-3p | 51.5 | 2.896757557 | 2.9294E-08 | hsa-miR-431 | 7.75 | 3.92820929 |
| hsa-miR-4286 | 61.25 | 2.776624931 | 8.88844E-12 | hsa-miR-493 | 5.75 | 3.83746357 |
| hsa-miR-6131 | 23.5 | 2.764115943 | 2.67008E-05 | hsa-miR-471 | 27.75 | 3.53560345 |
| hsa-miR-1468-5p | 8.5 | 2.718384147 | 0.017664112 | hsa-miR-299 | 6 | 3.45994983 |
| hsa-miR-375-3p | 200.5 | 2.714217555 | 1.68418E-21 | hsa-miR-681 | 5 | 3.33785533 |
| hsa-miR-432-5p | 7 | 2.677806016 | 0.035065231 | hsa-miR-889 | 31 | 3.17982768 |
| hsa-miR-92a-1-5p | 22.25 | 2.621802171 | 6.29183E-05 | hsa-miR-191 | 6.25 | 3.16891751 |
| hsa-miR-300 | 199.75 | 2.603313446 | 7.44937E-21 | hsa-miR-409 | 87.25 | 3.13616617 |
| hsa-miR-487b-3p | 9.25 | 2.545489719 | 0.018078389 | hsa-miR-493 | 13.75 | 2.87346553 |
| hsa-miR-410-3p | 92.5 | 2.507672452 | 9.35178E-14 | hsa-miR-410 | 96.25 | 2.83319402 |
| hsa-miR-4451 | 16.5 | 2.461544585 | 0.002535639 | hsa-miR-375 | 179.5 | 2.64757502 |
| hsa-miR-485-3p | 8.25 | 2.443592948 | 0.031811354 | hsa-miR-411 | 113 | 2.63232618 |
| hsa-miR-127-3p | 636 | 2.421938742 | 2.88248E-33 | hsa-miR-518 | 13 | 2.59192234 |
| hsa-miR-143-3p | 315.5 | 2.380045836 | 1.72869E-16 | hsa-miR-9-5 | 93.5 | 2.57577452 |
| hsa-miR-1269a | 53.5 | 2.197482974 | 3.02699E-06 | hsa-miR-711 | 13.5 | 2.57201761 |
| hsa-miR-132-3p | 96.5 | 2.186181066 | 3.18626E-09 | hsa-miR-126 | 7.25 | 2.50333582 |
| hsa-miR-3615 | 70.75 | 2.092940668 | 1.80019E-06 | hsa-miR-445 | 15.5 | 2.47623496 |
| hsa-miR-3184-5p | 1213.25 | 2.065687 | 2.61892E-17 | hsa-miR-432 | 10.5 | 2.45673931 |
| hsa-miR-1299 | 33 | 1.973929192 | 0.001195274 | hsa-miR-300 | 169.5 | 2.40199484 |
| hsa-miR-671-3p | 58.25 | 1.961748049 | 2.85195E-06 | hsa-miR-428 | 46.5 | 2.29099851 |
| hsa-miR-193a-5p | 86.75 | 1.950688586 | 3.53654E-07 | hsa-miR-127 | 525 | 2.21150546 |
| hsa-miR-7706 | 92 | 1.931055942 | 8.42023E-07 | hsa-miR-130 | 46.25 | 2.17975391 |
| hsa-miR-532-5p | 223.5 | 1.926116927 | 4.44536E-09 | hsa-miR-548 | 17.25 | 2.15957845 |
| hsa-miR-128-1-5p | 39.5 | 1.892038537 | 0.001730097 | hsa-miR-143 | 248 | 2.06501552 |
| hsa-miR-100-3p | 40.75 | 1.883549656 | 0.001105691 | hsa-miR-312 | 23.5 | 1.95246413 |

| | | | | | | |
|---|---|---|---|---|---|---|
| hsa-miR-106b-3p | 112.5 | 1.85173073 | 1.20901E-06 | hsa-miR-770 | 84 | 1.92989073 |
| hsa-miR-25-3p | 27555.25 | 1.787278253 | 5.28415E-20 | hsa-miR-132 | 76.5 | 1.90327849 |
| hsa-miR-654-3p | 95.75 | 1.733490638 | 2.50861E-06 | hsa-miR-121 | 719 | 1.88903601 |
| hsa-miR-192-5p | 61738.25 | 1.706559858 | 4.00527E-17 | hsa-miR-500 | 77.25 | 1.86303637 |
| hsa-miR-9-5p | 64.5 | 1.637345201 | 0.00080847 | hsa-miR-25- | 25360 | 1.83091898 |
| hsa-miR-548q | 24.25 | 1.632846092 | 0.048903728 | hsa-miR-106 | 100.5 | 1.80984009 |
| hsa-miR-1304-3p | 37.25 | 1.617382112 | 0.007244354 | hsa-miR-378 | 1447.75 | 1.79042718 |
| hsa-miR-5009-5p | 72.25 | 1.59727078 | 0.000368338 | hsa-miR-193 | 72.5 | 1.78400698 |
| hsa-miR-4499 | 24 | 1.589123898 | 0.048311271 | hsa-miR-671 | 47.75 | 1.75201068 |
| hsa-miR-181a-5p | 24514 | 1.580481845 | 1.24625E-12 | hsa-miR-318 | 927.5 | 1.75069192 |
| hsa-miR-3927-3p | 64.75 | 1.562260815 | 0.001236931 | hsa-miR-654 | 84.25 | 1.66974451 |
| hsa-miR-944 | 72.75 | 1.548236799 | 0.00265734 | hsa-miR-361 | 50 | 1.62070306 |
| hsa-miR-101-3p | 502.5 | 1.54727767 | 3.06091E-06 | hsa-miR-532 | 164 | 1.55134056 |
| hsa-miR-10396a-5p | 169 | 1.538163336 | 0.000989495 | hsa-miR-148 | 836.5 | 1.46622488 |
| hsa-miR-28-3p | 5317.75 | 1.533747571 | 6.66035E-06 | hsa-miR-797 | 133.25 | 1.42100513 |
| hsa-miR-378a-3p | 1358 | 1.518447654 | 2.03884E-09 | hsa-miR-192 | 44690.25 | 1.38272713 |
| hsa-miR-12135 | 625.25 | 1.492895713 | 9.21468E-08 | hsa-miR-103 | 136.75 | 1.37853362 |
| hsa-miR-151a-3p | 5954 | 1.488367646 | 2.03884E-09 | hsa-miR-151 | 4939.5 | 1.37669 |
| hsa-miR-421 | 184.75 | 1.479462281 | 0.002042086 | hsa-miR-183 | 440 | -1.3450415 |
| hsa-miR-182-5p | 19159 | 1.476702473 | 2.94109E-09 | hsa-miR-181 | 209.25 | -1.3591246 |
| hsa-miR-31-5p | 6341.5 | 1.460748601 | 1.17504E-08 | hsa-miR-210 | 270 | -1.3773038 |
| hsa-miR-221-3p | 3722.75 | 1.428738976 | 1.32586E-07 | hsa-miR-345 | 235.25 | -1.3892262 |
| hsa-miR-423-3p | 1086.25 | 1.426122021 | 3.35868E-05 | hsa-miR-130 | 509 | -1.4004877 |
| hsa-miR-30a-3p | 656 | 1.405641409 | 6.66035E-06 | hsa-miR-568 | 99.75 | -1.4290227 |
| hsa-miR-1180-3p | 104 | 1.368430725 | 0.027792309 | hsa-miR-27b | 343.75 | -1.4582561 |
| hsa-miR-148a-3p | 854.5 | 1.367934566 | 2.45375E-05 | hsa-miR-151 | 2481.25 | -1.559592 |
| hsa-miR-128-3p | 458 | 1.366291875 | 0.000105952 | hsa-miR-100 | 3983 | -1.5832476 |
| hsa-miR-339-3p | 76 | 1.352452022 | 0.046548104 | hsa-miR-574 | 61.25 | -1.6512258 |
| hsa-miR-769-5p | 515.75 | 1.351929526 | 0.000885321 | hsa-miR-22- | 31.75 | -1.7027277 |
| hsa-miR-126-5p | 254.5 | 1.333697734 | 0.01452405 | hsa-miR-125 | 43.75 | -1.7109354 |
| hsa-miR-27a-3p | 99179.25 | 1.327761257 | 2.87395E-05 | hsa-miR-29a | 1045.75 | -1.7114582 |
| hsa-miR-34a-5p | 451.5 | 1.30309192 | 0.001372993 | hsa-miR-185 | 23.5 | -1.7256726 |
| hsa-miR-5579-3p | 372 | 1.249778485 | 0.012583637 | hsa-miR-326 | 27.5 | -1.7541113 |
| hsa-miR-320a-3p | 767.25 | 1.242296311 | 0.047396446 | hsa-miR-342 | 55.5 | -1.7667971 |
| hsa-miR-21-5p | 140882 | 1.227083384 | 0.00481232 | hsa-miR-374 | 111.75 | -1.7884745 |
| hsa-miR-103a-3p | 2301.5 | 1.188810679 | 0.02897099 | hsa-miR-24- | 729.75 | -1.7894499 |
| hsa-miR-1297 | 8526.5 | 1.181463616 | 0.031108124 | hsa-miR-450 | 67.25 | -1.7955982 |
| hsa-miR-183-5p | 440 | -1.250792956 | 0.009012233 | hsa-miR-222 | 19.75 | -1.8850796 |
| hsa-miR-15a-5p | 3259.5 | -1.288993348 | 0.000368338 | hsa-miR-15b | 51.25 | -1.898002 |
| hsa-miR-29a-3p | 1045.75 | -1.293485218 | 0.000628184 | hsa-miR-365 | 238.75 | -1.9513444 |
| hsa-miR-138-5p | 657.5 | -1.298573064 | 0.000885321 | hsa-miR-103 | 16.25 | -1.9565968 |
| hsa-miR-210-3p | 270 | -1.309339562 | 0.026385908 | hsa-miR-125 | 5732 | -1.9699202 |
| hsa-miR-1307-5p | 509 | -1.370227826 | 5.98506E-05 | hsa-miR-23a | 1803.5 | -1.9812254 |
| hsa-miR-151a-5p | 2481.25 | -1.377516901 | 3.57627E-06 | hsa-miR-315 | 369 | -2.0112734 |
| hsa-miR-365a-3p | 238.75 | -1.390318785 | 0.008746508 | hsa-miR-513 | 19 | -2.0268608 |

| | | | | | | |
|---|---|---|---|---|---|---|
| hsa-miR-100-5p | 3983 | -1.39964092 | 7.5445E-07 | hsa-let-7a-3 | 84.75 | -2.0963686 |
| hsa-let-7a-3p | 84.75 | -1.410727125 | 0.012599283 | hsa-miR-33b | 11.75 | -2.181481 |
| hsa-miR-4282 | 54.25 | -1.467740483 | 0.011058595 | hsa-miR-10a | 13 | -2.1957384 |
| hsa-miR-652-3p | 48.5 | -1.467776135 | 0.030565873 | hsa-miR-542 | 23.5 | -2.2495162 |
| hsa-miR-23a-3p | 1803.5 | -1.481230527 | 1.01948E-08 | hsa-miR-365 | 12.25 | -2.2729575 |
| hsa-miR-450a-5p | 26.25 | -1.547024988 | 0.042193785 | hsa-miR-452 | 42.25 | -2.2931326 |
| hsa-miR-2392 | 67 | -1.547654599 | 0.037250532 | hsa-miR-193 | 1137.5 | -2.3159845 |
| hsa-miR-452-5p | 42.25 | -1.550674063 | 0.008777735 | hsa-let-7e-3 | 17.5 | -2.4019492 |
| hsa-miR-125a-5p | 5732 | -1.564632353 | 8.05385E-12 | hsa-miR-193 | 25.75 | -2.4030429 |
| hsa-miR-345-5p | 235.25 | -1.574087773 | 9.55507E-07 | hsa-miR-31- | 94.5 | -2.4462731 |
| hsa-miR-22-5p | 31.75 | -1.600383413 | 0.024123207 | hsa-miR-450 | 26.25 | -2.462525 |
| hsa-miR-15b-3p | 51.25 | -1.632061228 | 0.002042086 | hsa-miR-331 | 319.25 | -2.4841164 |
| hsa-miR-24-3p | 729.75 | -1.634641919 | 9.08398E-10 | hsa-miR-582 | 142.75 | -2.5044941 |
| hsa-miR-326 | 27.5 | -1.654716355 | 0.037776995 | hsa-miR-615 | 9.75 | -2.5639462 |
| hsa-miR-450b-5p | 67.25 | -1.735340524 | 0.000111875 | hsa-miR-121 | 47.5 | -2.6518458 |
| hsa-miR-3155a | 369 | -1.781436892 | 3.1056E-13 | hsa-miR-27a | 59.75 | -2.843397 |
| hsa-miR-23a-5p | 29.75 | -1.781993372 | 0.006755219 | hsa-miR-429 | 14 | -3.0397877 |
| hsa-miR-130b-5p | 13.25 | -1.847598379 | 0.046548104 | hsa-miR-508 | 7.25 | -3.2669327 |
| hsa-miR-513a-5p | 19 | -1.877960391 | 0.018817553 | hsa-miR-124 | 7.25 | -3.2689853 |
| hsa-miR-4650-3p | 22.25 | -1.880024287 | 0.005949352 | hsa-miR-23a | 29.75 | -3.3568341 |
| hsa-miR-193a-3p | 1137.5 | -1.959415616 | 1.06991E-14 | hsa-let-7i-3p | 17.5 | -3.4008877 |
| hsa-miR-141-3p | 18 | -1.961074044 | 0.023265012 | hsa-miR-141 | 5.75 | -3.4037493 |
| hsa-miR-3978 | 20.75 | -1.985347336 | 0.030939532 | hsa-miR-887 | 8.5 | -3.467956 |
| hsa-miR-12133 | 47.5 | -1.995823911 | 8.92639E-06 | hsa-miR-324 | 9.75 | -3.5403978 |
| hsa-miR-342-3p | 55.5 | -2.028239726 | 3.86737E-05 | hsa-miR-674 | 59 | -3.7187136 |
| hsa-miR-31-3p | 94.5 | -2.0315241 | 2.20116E-07 | hsa-miR-990 | 149 | -4.5840408 |
| hsa-miR-27a-5p | 59.75 | -2.10724928 | 4.7134E-06 | hsa-miR-578 | 10.5 | -4.7034336 |
| hsa-miR-331-3p | 319.25 | -2.143885835 | 7.07298E-11 | hsa-miR-442 | 35.75 | -4.7602787 |
| hsa-miR-7-5p | 20.75 | -2.212172667 | 0.001312309 | hsa-miR-342 | 7 | -5.9616441 |
| hsa-miR-193b-5p | 25.75 | -2.222033995 | 0.000441332 | hsa-miR-124 | 6.25 | -6.3608202 |
| hsa-miR-4425 | 35.75 | -2.243761671 | 0.006459381 | hsa-miR-673 | 115.5 | -10.116086 |
| hsa-miR-18a-5p | 24 | -2.33183721 | 0.00050715 | hsa-miR-445 | 4 | -10.349613 |
| hsa-miR-503-5p | 11.25 | -2.37390864 | 0.012599283 | hsa-miR-675 | 4.5 | -11.603304 |
| hsa-miR-5089-5p | 7.25 | -2.391673864 | 0.048172566 | hsa-miR-677 | 10.75 | -11.694156 |
| hsa-miR-3613-5p | 8.5 | -2.414682594 | 0.028536782 | hsa-miR-394 | 22 | -33.121885 |
| hsa-miR-505-3p | 8 | -2.441713801 | 0.034670523 | hsa-miR-607 | 7.5 | -57.203548 |
| hsa-miR-1296-5p | 18.5 | -2.489923644 | 0.000444053 | | | |
| hsa-miR-6734-5p | 11.5 | -2.553245405 | 0.006248326 | | | |
| hsa-miR-6808-5p | 6.5 | -2.55462373 | 0.048573728 | | | |
| hsa-miR-324-3p | 9.75 | -2.586659617 | 0.017664112 | | | |
| hsa-miR-33b-3p | 11.75 | -2.61139621 | 0.0047073 | | | |
| hsa-miR-582-5p | 142.75 | -2.766641133 | 1.45449E-15 | | | |
| hsa-miR-875-3p | 9.25 | -2.860994578 | 0.017903543 | | | |
| hsa-miR-6728-5p | 12.75 | -2.990014611 | 0.000989495 | | | |
| hsa-miR-6741-5p | 59 | -3.487797235 | 1.36775E-05 | | | |

| | | | |
|---|---|---|---|
| hsa-let-7i-3p | 17.5 | -3.497254545 | 1.65808E-05 |
| hsa-miR-6840-3p | 4.75 | -3.561358697 | 0.041186421 |
| hsa-miR-890 | 5 | -3.741905655 | 0.032218103 |
| hsa-miR-342-5p | 7 | -3.833847407 | 0.009012233 |
| hsa-miR-9900 | 149 | -3.843798339 | 0.000155283 |
| hsa-miR-6892-5p | 5.25 | -4.031068032 | 0.031108124 |
| hsa-miR-4718 | 6.75 | -4.260369825 | 0.006347337 |
| hsa-miR-578 | 10.5 | -4.523987514 | 0.000345771 |
| hsa-miR-1246 | 6.25 | -5.064507095 | 0.035591555 |
| hsa-miR-6752-5p | 4.5 | -5.262785062 | 0.027033868 |
| hsa-miR-34a-3p | 4.75 | -5.547756403 | 0.017664112 |
| hsa-miR-6070 | 7.5 | -6.790298146 | 0.001277869 |
| hsa-miR-6774-3p | 10.75 | -6.868021193 | 0.000160203 |
| hsa-miR-6737-5p | 115.5 | -9.070698429 | 3.24044E-35 |
| hsa-miR-584-3p | 3.5 | -9.325112396 | 0.038500193 |
| hsa-miR-3940-5p | 22 | -12.15629605 | 1.53367E-08 |

### PI Ad14 v C 24HPI Ad14 v

| FDR p-value | Identifier | Max group mean | Fold change | FDR p-value | Identifier | Max group mean |
|---|---|---|---|---|---|---|
| 0.02193832 | hsa-miR-127 | 471.5 | 2.68519268 | 8.3477E-33 | hsa-miR-518 | 34.75 |
| 0.0282903 | hsa-miR-990 | 149 | -4.7441554 | 4.9698E-31 | hsa-miR-105 | 8.75 |
| 0.00312531 | hsa-miR-25- | 20627.5 | 1.98453754 | 1.092E-27 | hsa-miR-684 | 5.75 |
| 0.00408364 | hsa-miR-673 | 115.5 | -16.537883 | 1.6771E-27 | hsa-miR-312 | 51.25 |
| 0.00024718 | hsa-miR-300 | 150.5 | 2.84743567 | 2.5809E-27 | hsa-miR-466 | 3.75 |
| 0.01332408 | hsa-miR-193 | 1137.5 | -2.0636122 | 2.1617E-25 | hsa-miR-155 | 6 |
| 0.03253087 | hsa-miR-27b | 343.75 | -2.46075 | 2.1617E-25 | hsa-miR-466 | 3.5 |
| 0.00036927 | hsa-miR-375 | 139 | 2.74120474 | 9.4741E-25 | hsa-miR-449 | 3 |
| 0.01226722 | hsa-miR-411 | 96.5 | 3.0081375 | 1.8804E-22 | hsa-miR-312 | 6 |
| 2.8359E-05 | hsa-miR-378 | 1138.5 | 1.89089378 | 2.0398E-21 | hsa-miR-548 | 12.75 |
| 2.8615E-05 | hsa-miR-143 | 249.25 | 2.81123466 | 2.4728E-21 | hsa-miR-428 | 144.25 |
| 1.9777E-06 | hsa-miR-23a | 1803.5 | -1.8808798 | 3.7085E-21 | hsa-miR-119 | 9.75 |
| 0.0282903 | hsa-miR-428 | 75.5 | 5.01768765 | 3.5617E-19 | hsa-miR-557 | 2.25 |
| 0.01855109 | hsa-miR-132 | 95 | 3.13342666 | 7.3968E-18 | hsa-miR-649 | 8.75 |
| 0.00010347 | hsa-miR-410 | 72.75 | 2.86576492 | 2.613E-17 | hsa-miR-472 | 3.25 |
| 0.00028925 | hsa-miR-315 | 369 | -2.0078678 | 4.5325E-17 | hsa-miR-427 | 5.5 |
| 0.00873903 | hsa-miR-9-5 | 74.75 | 2.75334893 | 9.8727E-17 | hsa-miR-888 | 10.5 |
| 0.02832155 | hsa-miR-331 | 319.25 | -2.305022 | 9.8727E-17 | hsa-miR-677 | 3 |
| 4.2753E-08 | hsa-miR-106 | 105.25 | 2.53822638 | 1.6255E-16 | hsa-miR-128 | 4 |
| 0.02267168 | hsa-miR-27a | 59.75 | -7.4686987 | 8.6459E-15 | hsa-miR-568 | 21.25 |
| 0.04511882 | hsa-miR-674 | 59 | -7.6231274 | 1.8044E-14 | hsa-miR-127 | 16.5 |
| 7.2353E-06 | hsa-miR-471 | 29.25 | 4.97320705 | 1.5504E-13 | hsa-miR-449 | 7.75 |
| 0.04248943 | hsa-miR-121 | 539.25 | 1.90016957 | 1.5504E-13 | hsa-miR-124 | 20.5 |
| 2.9929E-14 | hsa-miR-148 | 709.5 | 1.67279146 | 2.0712E-13 | hsa-miR-125 | 11.75 |
| 0.00091123 | hsa-miR-532 | 155 | 1.95680404 | 2.1819E-13 | hsa-miR-668 | 20 |
| 7.292E-12 | hsa-miR-318 | 728.75 | 1.84884124 | 4.4666E-13 | hsa-miR-493 | 6.25 |
| 1.3149E-14 | hsa-miR-125 | 5732 | -1.9206437 | 2.5222E-12 | hsa-miR-443 | 9.5 |
| 1.7555E-11 | hsa-miR-582 | 142.75 | -2.3360371 | 5.8167E-12 | hsa-miR-672 | 4 |
| 0.00434081 | hsa-miR-365 | 238.75 | -1.8311022 | 4.7384E-11 | hsa-miR-520 | 5.75 |
| 3.7739E-11 | hsa-miR-889 | 29 | 3.98232137 | 7.2047E-11 | hsa-miR-551 | 3.75 |
| 0.0028125 | hsa-miR-500 | 68.75 | 2.20542802 | 7.2047E-11 | hsa-miR-429 | 7 |
| 0.04729698 | hsa-miR-31- | 94.5 | -2.3523397 | 7.2047E-11 | hsa-miR-425 | 4.25 |
| 0.00878388 | hsa-let-7a-3 | 84.75 | -2.4799905 | 7.2047E-11 | hsa-miR-432 | 4.5 |
| 0.02645525 | hsa-miR-151 | 4017.25 | 1.48922224 | 1.3904E-09 | hsa-miR-889 | 27.75 |
| 6.9577E-09 | hsa-miR-101 | 348.25 | 1.58501891 | 2.5936E-09 | hsa-miR-120 | 8 |
| 1.6056E-05 | hsa-miR-409 | 49.75 | 2.38852319 | 8.496E-09 | hsa-miR-431 | 5.25 |
| 1.0218E-09 | hsa-miR-119 | 15.25 | 13.3534772 | 9.0093E-09 | hsa-miR-676 | 6.25 |
| 7.4213E-05 | hsa-miR-181 | 17184.75 | 1.63388193 | 1.0265E-08 | hsa-miR-106 | 130 |
| 0.0212453 | hsa-miR-24- | 729.75 | -1.5222461 | 1.5302E-08 | hsa-miR-410 | 74.75 |
| 2.3243E-05 | hsa-miR-125 | 43.75 | -3.1326036 | 4.3014E-08 | hsa-miR-806 | 68.75 |
| 0.01113146 | hsa-miR-442 | 35.75 | -3.9751653 | 6.3813E-08 | hsa-miR-447 | 4.25 |

| | | | | | | |
|---|---|---|---|---|---|---|
| 2.4945E-05 | hsa-miR-312 | 14.5 | 8.68625261 | 7.1767E-08 | hsa-miR-369 | 4.25 |
| 0.00016257 | hsa-miR-193 | 65.75 | 2.1922762 | 7.1767E-08 | hsa-miR-9-5 | 76 |
| 6.3294E-06 | hsa-miR-770 | 63 | 1.94239561 | 9.6117E-08 | hsa-miR-126 | 5.75 |
| 0.00011091 | hsa-miR-29a | 1045.75 | -1.4630058 | 1.2261E-07 | hsa-miR-127 | 472.5 |
| 9.4469E-06 | hsa-miR-181 | 209.25 | -1.6543437 | 1.2495E-07 | hsa-miR-300 | 137.25 |
| 5.8189E-05 | hsa-miR-449 | 12.5 | 9.50833686 | 1.4852E-07 | hsa-miR-268 | 10.75 |
| 2.5323E-06 | hsa-miR-182 | 12365.5 | 1.41128583 | 1.6259E-07 | hsa-miR-33a | 5.75 |
| 0.00051715 | hsa-miR-125 | 15 | 7.4574853 | 2.0868E-07 | hsa-miR-143 | 224 |
| 0.01308371 | hsa-miR-23a | 29.75 | -4.3317987 | 5.4095E-07 | hsa-miR-315 | 16 |
| 2.1711E-05 | hsa-miR-192 | 38121.75 | 1.59488073 | 1.3586E-06 | hsa-miR-375 | 114.25 |
| 0.00118262 | hsa-miR-126 | 196.25 | 1.51952294 | 1.6762E-06 | hsa-miR-518 | 8.25 |
| 0.01642955 | hsa-miR-315 | 20 | 2.81773851 | 1.8861E-06 | hsa-miR-432 | 7.25 |
| 0.01225814 | hsa-miR-128 | 39 | 2.16717259 | 3.4465E-06 | hsa-miR-471 | 12.75 |
| 0.00738567 | hsa-miR-601 | 29.25 | -3.1304262 | 4.3454E-06 | hsa-miR-7-5 | 31 |
| 0.00960823 | hsa-miR-27a | 71444.75 | 1.35342478 | 5.7987E-06 | hsa-miR-689 | 15 |
| 0.03870302 | hsa-miR-443 | 10.25 | 5.58786915 | 9.2315E-06 | hsa-miR-569 | 6 |
| 0.01123233 | hsa-miR-568 | 13 | 4.01832651 | 9.387E-06 | hsa-miR-500 | 64.75 |
| 0.03870302 | hsa-miR-394 | 22 | -25.537202 | 9.9402E-06 | hsa-miR-130 | 33 |
| 0.01650313 | hsa-miR-28- | 3228.25 | 1.3501866 | 1.1361E-05 | hsa-miR-128 | 36.75 |
| 0.0423163 | hsa-miR-130 | 34 | 2.13926752 | 1.8408E-05 | hsa-miR-411 | 65 |
| 0.01428901 | hsa-miR-797 | 100.75 | -1.696463 | 1.8408E-05 | hsa-miR-146 | 7.25 |
| 0.01543801 | hsa-miR-427 | 8.5 | 10.7868522 | 2.3876E-05 | hsa-miR-493 | 7.25 |
| 0.00392512 | hsa-miR-222 | 19.75 | -5.3199835 | 2.7209E-05 | hsa-miR-770 | 65.75 |
| 0.02178237 | hsa-miR-361 | 43.75 | 1.92164325 | 7.7359E-05 | hsa-miR-409 | 39.75 |
| 0.00344382 | hsa-miR-574 | 61.25 | -1.9858122 | 8.0701E-05 | hsa-miR-808 | 28 |
| 0.00093223 | hsa-miR-548 | 16.25 | 2.69069953 | 0.00012589 | hsa-miR-378 | 1133.25 |
| 0.00107339 | hsa-miR-465 | 22.25 | -3.0837119 | 0.00013111 | hsa-miR-548 | 10.5 |
| 0.01440197 | hsa-miR-493 | 11.5 | 3.21070924 | 0.00014031 | hsa-miR-193 | 53.25 |
| 0.01788754 | hsa-miR-124 | 10.75 | 3.51709756 | 0.00014189 | hsa-miR-181 | 19021.5 |
| 0.01088839 | hsa-miR-126 | 31.75 | 1.89333131 | 0.00016701 | hsa-miR-25- | 18406.25 |
| 1.6827E-05 | hsa-miR-151 | 2481.25 | -1.4229041 | 0.00018193 | hsa-miR-126 | 27.75 |
| 0.02167141 | hsa-miR-181 | 418.5 | -1.3647202 | 0.00021333 | hsa-miR-132 | 48 |
| 0.02660609 | hsa-miR-7-5 | 29 | 1.97939437 | 0.00022083 | hsa-miR-192 | 38400.5 |
| 0.0028125 | hsa-miR-155 | 7 | 15.9460388 | 0.00025075 | hsa-miR-449 | 15.75 |
| 0.00055591 | hsa-miR-450 | 67.25 | -1.829962 | 0.00027629 | hsa-miR-361 | 34 |
| 5.1624E-06 | hsa-miR-888 | 8 | 5.46026095 | 0.0002908 | hsa-miR-532 | 116 |
| 0.00149706 | hsa-miR-268 | 12.25 | 2.861235 | 0.00032789 | hsa-miR-182 | 13029.25 |
| 0.03177486 | hsa-miR-689 | 16.75 | 2.36179315 | 0.00033963 | hsa-miR-148 | 612.25 |
| 0.001821 | hsa-let-7i-3p | 17.5 | -3.2174871 | 0.00043147 | hsa-miR-543 | 54.75 |
| 1.3935E-07 | hsa-miR-33b | 11.75 | -7.5537075 | 0.00043966 | hsa-miR-318 | 582 |
| 0.02233453 | hsa-miR-34a | 335.75 | 1.33246007 | 0.00055878 | hsa-miR-769 | 362.75 |
| 2.6501E-07 | hsa-miR-342 | 55.5 | -1.7708359 | 0.0005743 | hsa-miR-151 | 3849.5 |
| 3.6933E-08 | hsa-miR-520 | 6.5 | 5.74671095 | 0.00057789 | hsa-miR-500 | 33.75 |
| 2.7776E-09 | hsa-miR-466 | 7.25 | 27.1050011 | 0.00069666 | hsa-miR-126 | 183.25 |
| 0.01224399 | hsa-miR-711 | 11 | 2.80066973 | 0.00069666 | hsa-miR-27a | 71444.75 |

| | | | | | | |
|---|---|---|---|---|---|---|
| 2.7787E-06 | hsa-miR-128 | 327.25 | 1.32536521 | 0.00078473 | hsa-miR-101 | 311 |
| 0.02733844 | hsa-let-7e-3p | 17.5 | -2.8486322 | 0.00110197 | hsa-miR-340 | 235.75 |
| 0.01855109 | hsa-miR-472 | 5.5 | 12.5790243 | 0.00111118 | hsa-miR-186 | 792.25 |
| 0.00091123 | hsa-miR-340 | 235.75 | 1.32876859 | 0.00152671 | hsa-miR-103 | 1881 |
| 0.01638408 | hsa-miR-186 | 792.25 | 1.27527134 | 0.00155986 | hsa-miR-129 | 6967.25 |
| 4.3285E-05 | hsa-miR-769 | 362.75 | 1.29815728 | 0.00187153 | hsa-miR-15a | 3259.5 |
| 2.3573E-11 | hsa-miR-684 | 13.75 | 148.627998 | 0.00232929 | hsa-miR-28- | 241.25 |
| 0.00678019 | hsa-miR-26a | 9.25 | -5.9616666 | 0.00321161 | hsa-miR-130 | 509 |
| 0.0001227 | hsa-miR-429 | 14 | -2.964567 | 0.00339392 | hsa-miR-22- | 28627.5 |
| 4.3143E-08 | hsa-miR-654 | 56.25 | 1.48094706 | 0.00368014 | hsa-miR-181 | 418.5 |
| 0.00093223 | hsa-miR-679 | 4.5 | 10.3359724 | 0.00373521 | hsa-miR-345 | 235.25 |
| 2.0329E-12 | hsa-miR-672 | 12.75 | -3.1733819 | 0.00377322 | hsa-miR-138 | 657.5 |
| 4.0256E-11 | hsa-miR-578 | 10.5 | -4.0285838 | 0.00385864 | hsa-miR-568 | 99.75 |
| 0.01852622 | hsa-miR-312 | 4.5 | 7.41036282 | 0.00418129 | hsa-miR-15b | 51.25 |
| 6.1316E-06 | hsa-miR-100 | 3983 | -1.3900736 | 0.00418129 | hsa-miR-374 | 111.75 |
| 1.4959E-06 | hsa-miR-105 | 10 | 107.793847 | 0.00489891 | hsa-miR-151 | 2481.25 |
| 0.00118402 | hsa-miR-365 | 12.25 | -3.0513148 | 0.00515769 | hsa-miR-125 | 80.75 |
| 0.01888326 | hsa-miR-518 | 9.5 | 2.53494172 | 0.00516848 | hsa-miR-335 | 56.75 |
| 0.01985943 | hsa-miR-887 | 8.5 | -5.4871954 | 0.00524647 | hsa-miR-224 | 111 |
| 6.2044E-07 | hsa-miR-685 | 8.75 | -5.685826 | 0.00524647 | hsa-miR-424 | 32 |
| 5.9824E-05 | hsa-miR-15a | 3259.5 | -1.2294541 | 0.00543767 | hsa-miR-24- | 729.75 |
| 0.03739501 | hsa-miR-450 | 26.25 | -2.1971072 | 0.00543767 | hsa-miR-181 | 209.25 |
| 0.03083442 | hsa-miR-449 | 19.25 | 1.84753864 | 0.00612982 | hsa-miR-318 | 42.25 |
| 0.00325623 | hsa-miR-374 | 111.75 | -1.3934326 | 0.00696708 | hsa-miR-30c | 47.25 |
| 0.00065725 | hsa-miR-607 | 7.5 | -15.017167 | 0.01022453 | hsa-miR-29a | 1045.75 |
| 0.00038088 | hsa-miR-493 | 4.75 | 4.23717654 | 0.01263867 | hsa-miR-797 | 100.75 |
| 0.00065724 | hsa-miR-471 | 6.75 | -7.9717754 | 0.01273355 | hsa-miR-315 | 369 |
| 6.9577E-09 | hsa-miR-185 | 23.5 | -1.8777877 | 0.01405266 | hsa-miR-365 | 238.75 |
| 0.00481579 | hsa-miR-193 | 25.75 | -1.934458 | 0.01416865 | hsa-miR-190 | 28.25 |
| 0.03778346 | hsa-miR-557 | 3.5 | 13.3735047 | 0.01467412 | hsa-miR-23a | 1803.5 |
| 1.9433E-20 | hsa-miR-649 | 4.5 | 4.00742701 | 0.01470728 | hsa-miR-450 | 29.75 |
| 0.03838287 | hsa-miR-221 | 20 | -2.3547426 | 0.01507472 | hsa-miR-22- | 31.75 |
| 0.02719952 | hsa-miR-548 | 4.75 | 3.66308638 | 0.01570749 | hsa-miR-125 | 5732 |
| 0.00025742 | hsa-miR-423 | 740.5 | 1.31551843 | 0.01606412 | hsa-miR-140 | 33.75 |
| 2.6174E-06 | hsa-miR-513 | 19 | -2.1100138 | 0.01617447 | hsa-miR-360 | 63.75 |
| 0.02953091 | hsa-miR-429 | 5 | 3.40001843 | 0.01656725 | hsa-miR-574 | 61.25 |
| | hsa-miR-106 | 1451.5 | 1.20527002 | 0.01784404 | hsa-miR-193 | 25.75 |
| | hsa-miR-335 | 176.25 | -1.2928748 | 0.01877473 | hsa-let-7a-3p | 84.75 |
| | hsa-miR-568 | 99.75 | -1.371823 | 0.01877473 | hsa-miR-193 | 1137.5 |
| | hsa-miR-125 | 20.75 | -1.8713685 | 0.01877473 | hsa-miR-450 | 26.25 |
| | hsa-miR-360 | 63.75 | -1.4777521 | 0.02078328 | hsa-miR-31- | 94.5 |
| | hsa-miR-130 | 94.25 | 1.31789964 | 0.02171496 | hsa-miR-182 | 52.5 |
| | hsa-miR-673 | 6.25 | -5.2389297 | 0.02266008 | hsa-miR-428 | 54.25 |
| | hsa-miR-369 | 4.75 | 3.23481319 | 0.02459502 | hsa-miR-331 | 319.25 |
| | hsa-miR-454 | 15 | 1.80550277 | 0.02467714 | hsa-miR-312 | 12.75 |

| | | | | | |
|---|---|---|---|---|---|
| hsa-miR-316 | 25.75 | -1.7183278 | 0.02624287 | hsa-miR-465 | 44 |
| hsa-miR-677 | 3 | 6.97138642 | 0.0271769 | hsa-miR-487 | 12.25 |
| hsa-miR-677 | 10.75 | -71.017019 | 0.0287718 | hsa-miR-452 | 42.25 |
| hsa-miR-447 | 4.75 | 3.23427718 | 0.02902086 | hsa-miR-582 | 142.75 |
| hsa-miR-224 | 111 | -1.3716802 | 0.0291866 | hsa-let-7i-3p | 17.5 |
| hsa-miR-561 | 78 | 1.32421329 | 0.02920105 | hsa-miR-715 | 22 |
| hsa-miR-210 | 13.75 | -2.1213476 | 0.02950606 | hsa-miR-664 | 12 |
| hsa-miR-15b | 51.25 | -1.5846487 | 0.03002462 | hsa-miR-615 | 9.75 |
| hsa-miR-100 | 21 | -1.9574526 | 0.03002462 | hsa-miR-542 | 23.5 |
| hsa-miR-314 | 2.75 | 10.5919415 | 0.03131551 | hsa-miR-10a | 13 |
| hsa-miR-29b | 9 | -2.7278176 | 0.03198622 | hsa-miR-27b | 343.75 |
| hsa-miR-132 | 5.5 | 2.76203781 | 0.03332851 | hsa-miR-26a | 9.25 |
| hsa-miR-465 | 44 | -1.507867 | 0.03336528 | hsa-miR-214 | 88.75 |
| hsa-miR-673 | 11.5 | -2.4365773 | 0.03456163 | hsa-miR-125 | 43.75 |
| hsa-miR-124 | 7.25 | -3.2348344 | 0.03558611 | hsa-miR-210 | 13.75 |
| hsa-miR-183 | 440 | -1.2829992 | 0.03684848 | hsa-let-7e-3p | 17.5 |
| hsa-miR-134 | 5 | -10.100308 | 0.0370233 | hsa-miR-222 | 19.75 |
| hsa-miR-471 | 8.25 | -2.8006826 | 0.03993485 | hsa-miR-685 | 8.75 |
| hsa-miR-889 | 5 | -5.9448202 | 0.04173024 | hsa-miR-315 | 8 |
| hsa-miR-678 | 6.5 | -3.4390856 | 0.04272732 | hsa-miR-397 | 20.75 |
| hsa-miR-34a | 4.75 | -9.6075645 | 0.04284866 | hsa-miR-316 | 25.75 |
| | | | | hsa-miR-429 | 14 |
| | | | | hsa-miR-990 | 149 |
| | | | | hsa-miR-192 | 10.5 |
| | | | | hsa-miR-33b | 11.75 |
| | | | | hsa-miR-442 | 35.75 |
| | | | | hsa-miR-601 | 29.25 |
| | | | | hsa-miR-465 | 22.25 |
| | | | | hsa-miR-27a | 59.75 |
| | | | | hsa-miR-715 | 5.25 |
| | | | | hsa-miR-673 | 11.5 |
| | | | | hsa-miR-29b | 9 |
| | | | | hsa-miR-342 | 7 |
| | | | | hsa-miR-607 | 7.5 |
| | | | | hsa-miR-677 | 10.75 |
| | | | | hsa-miR-674 | 59 |
| | | | | hsa-miR-34a | 4.75 |
| | | | | hsa-miR-471 | 8.25 |
| | | | | hsa-miR-394 | 22 |
| | | | | hsa-miR-430 | 5.25 |
| | | | | hsa-miR-23a | 29.75 |
| | | | | hsa-miR-471 | 6.75 |
| | | | | hsa-miR-672 | 12.75 |
| | | | | hsa-miR-673 | 115.5 |

C 36HPI Ad14 v C 48HPI Ad1

| Fold change | FDR p-value | Identifier | Max group mean | Fold change | FDR p-value | Identifier |
|---|---|---|---|---|---|---|
| 144.955825 | 1.467E-08 | hsa-miR-518 | 93 | 251.482132 | 3.8804E-10 | hsa-miR-127-3p |
| 100.547201 | 0.00310806 | hsa-miR-312 | 97 | 44.6022435 | 3.1356E-23 | hsa-miR-192-5p |
| 66.4768095 | 0.00834291 | hsa-miR-684 | 5.25 | 39.8543634 | 0.03117988 | hsa-miR-6737-5p |
| 34.7878185 | 7.2106E-24 | hsa-miR-807 | 5.25 | 39.1873894 | 0.03032373 | hsa-miR-300 |
| 15.9226355 | 0.00506871 | hsa-miR-681 | 21.5 | 21.9737687 | 0.00055992 | hsa-miR-181a-5p |
| 15.4362985 | 0.00025681 | hsa-miR-520 | 32.75 | 21.698749 | 3.5467E-11 | hsa-miR-3155a |
| 15.0631664 | 0.00870042 | hsa-miR-472 | 12 | 20.7112883 | 6.7885E-05 | hsa-miR-143-3p |
| 12.8338933 | 0.01173054 | hsa-miR-428 | 396.75 | 20.0218463 | 5.0603E-79 | hsa-miR-410-3p |
| 11.1113501 | 0.00020104 | hsa-miR-155 | 10 | 17.1972412 | 0.0002385 | hsa-miR-182-5p |
| 11.0299734 | 2.0152E-08 | hsa-miR-124 | 5.75 | 16.0118773 | 0.00626485 | hsa-miR-411-5p |
| 10.6290386 | 3.1634E-55 | hsa-miR-646 | 24 | 15.9688698 | 1.0041E-08 | hsa-miR-5187-3p |
| 9.7396477 | 1.1466E-06 | hsa-miR-785 | 5 | 13.9561887 | 0.00932211 | hsa-miR-409-3p |
| 9.7335371 | 0.03406263 | hsa-miR-312 | 11 | 13.4619261 | 6.7885E-05 | hsa-miR-582-5p |
| 8.75228409 | 4.2842E-06 | hsa-miR-557 | 4.75 | 13.2777963 | 0.01118698 | hsa-miR-4286 |
| 8.47655736 | 0.00847554 | hsa-miR-126 | 22.75 | 13.0829622 | 2.0317E-08 | hsa-miR-125a-5p |
| 7.96368971 | 0.0004546 | hsa-miR-429 | 23.5 | 11.9622904 | 1.4305E-09 | hsa-miR-1307-5p |
| 7.9244676 | 9.0981E-06 | hsa-miR-127 | 41 | 11.4194897 | 8.0103E-15 | hsa-miR-7706 |
| 7.84606117 | 0.01202846 | hsa-miR-596 | 11.75 | 11.2380055 | 2.4853E-05 | hsa-miR-27a-3p |
| 7.4703974 | 0.00411339 | hsa-miR-125 | 29.5 | 11.1207072 | 2.9912E-10 | hsa-miR-25-3p |
| 7.45476076 | 3.2755E-13 | hsa-miR-432 | 16.25 | 10.7993579 | 1.6024E-06 | hsa-miR-23a-3p |
| 6.93684108 | 6.3406E-09 | hsa-miR-888 | 19.75 | 10.0973519 | 4.8974E-08 | hsa-miR-24-3p |
| 6.73758493 | 3.1381E-05 | hsa-miR-607 | 6.5 | 7.95342707 | 0.03117988 | hsa-miR-889-3p |
| 6.66090956 | 1.2827E-08 | hsa-miR-121 | 24.5 | 7.40641653 | 2.0559E-08 | hsa-miR-375-3p |
| 6.63421445 | 8.2812E-07 | hsa-miR-200 | 7.5 | 7.21584573 | 0.00117432 | hsa-miR-193a-3p |
| 6.34146669 | 1.2942E-11 | hsa-miR-121 | 7.5 | 7.21485354 | 0.00164585 | hsa-miR-345-5p |
| 6.27975172 | 0.00025681 | hsa-miR-124 | 29 | 7.18423189 | 5.3279E-10 | hsa-miR-151a-5p |
| 5.90170515 | 5.5823E-06 | hsa-miR-472 | 16 | 7.02183105 | 0.0104206 | hsa-miR-138-5p |
| 5.82858566 | 0.0049853 | hsa-miR-551 | 7.25 | 6.97927054 | 0.00199598 | hsa-miR-151a-3p |
| 5.78770387 | 0.00068868 | hsa-miR-425 | 8.25 | 6.51131036 | 0.00094163 | hsa-miR-532-5p |
| 5.47995948 | 0.00718581 | hsa-miR-368 | 3.75 | 6.48619648 | 0.03163791 | hsa-miR-136-3p |
| 5.38171264 | 0.00017279 | hsa-miR-33a | 23.25 | 6.42123528 | 2.5311E-07 | hsa-miR-15a-5p |
| 4.83269465 | 0.01525482 | hsa-miR-427 | 6.5 | 6.2707947 | 0.00268433 | hsa-miR-29a-3p |
| 4.54952655 | 0.00605645 | hsa-miR-302 | 5 | 6.195241 | 0.01043621 | hsa-miR-365a-3p |
| 4.34818403 | 1.2923E-13 | hsa-miR-446 | 3.5 | 6.08096762 | 0.04697129 | hsa-miR-100-3p |
| 4.19205097 | 0.00025356 | hsa-miR-492 | 14.5 | 5.51168277 | 1.5672E-05 | hsa-miR-378a-3p |
| 4.05765543 | 0.00517889 | hsa-miR-426 | 6.75 | 5.33984333 | 0.00505255 | hsa-miR-106b-3p |
| 3.90228393 | 0.00153903 | hsa-miR-483 | 18.5 | 5.22283559 | 4.8155E-06 | hsa-miR-193a-5p |
| 3.54114155 | 6.413E-25 | hsa-miR-520 | 10 | 5.16969651 | 0.00050961 | hsa-miR-769-5p |
| 3.3495601 | 4.2437E-18 | hsa-miR-568 | 21.5 | 5.06108964 | 2.495E-07 | hsa-miR-671-3p |
| 3.33938819 | 1.8112E-14 | hsa-miR-366 | 8.5 | 5.02412015 | 0.01537905 | hsa-miR-3615 |
| 3.30020474 | 0.01841553 | hsa-miR-364 | 12.75 | 4.83090015 | 0.00015544 | hsa-miR-7-5p |

| | | | | | | |
|---|---|---|---|---|---|---|
| 3.27602226 | 0.0418568 | hsa-miR-548 | 8 | 4.65534784 | 0.0024787 | hsa-miR-30a-3p |
| 3.15139587 | 1.6103E-15 | hsa-miR-569 | 4.75 | 4.6488103 | 0.04802981 | hsa-miR-3940-5p |
| 3.02584624 | 0.00997081 | hsa-miR-478 | 13 | 4.5326925 | 0.00019029 | hsa-miR-1269a |
| 2.96789236 | 2.2207E-32 | hsa-miR-450 | 14.75 | 4.43785691 | 2.642E-05 | hsa-miR-148a-3p |
| 2.96764623 | 3.6121E-19 | hsa-miR-121 | 5.5 | 4.36873232 | 0.02032988 | hsa-miR-9-5p |
| 2.86923745 | 0.00095783 | hsa-miR-7-5 | 79.25 | 4.12252341 | 2.756E-05 | hsa-miR-5009-5p |
| 2.67447199 | 0.03312876 | hsa-miR-315 | 13.5 | 4.05983497 | 0.00022607 | hsa-miR-101-3p |
| 2.59697165 | 3.6725E-13 | hsa-miR-569 | 15.25 | 4.03762414 | 0.00015544 | hsa-miR-944 |
| 2.57799238 | 3.4675E-05 | hsa-miR-268 | 22.25 | 3.95876921 | 2.1335E-06 | hsa-miR-31-3p |
| 2.57652276 | 1.0436E-18 | hsa-miR-674 | 9.5 | 3.95346085 | 0.00326769 | hsa-miR-331-3p |
| 2.51640643 | 0.00578003 | hsa-miR-119 | 5.75 | 3.87500964 | 0.01954272 | hsa-miR-649 |
| 2.50456749 | 0.02053733 | hsa-miR-130 | 7.5 | 3.82247447 | 0.01472019 | hsa-let-7a-3p |
| 2.48812577 | 0.00059966 | hsa-miR-362 | 14 | 3.81196168 | 0.0008284 | hsa-miR-6741-5p |
| 2.42267517 | 1.743E-07 | hsa-miR-315 | 34 | 3.67835196 | 1.1103E-06 | hsa-miR-450b-5p |
| 2.39549512 | 0.00086639 | hsa-miR-443 | 8.75 | 3.64659806 | 0.0056062 | hsa-miR-12135 |
| 2.39298209 | 0.02824865 | hsa-miR-203 | 6 | 3.50774154 | 0.02569168 | hsa-miR-128-3p |
| 2.36997911 | 1.4708E-11 | hsa-miR-518 | 11.5 | 3.46654504 | 0.00093578 | hsa-miR-3181 |
| 2.3689883 | 1.5705E-07 | hsa-miR-668 | 16 | 3.4080598 | 0.00015964 | hsa-miR-676-3p |
| 2.33775388 | 5.4163E-08 | hsa-miR-106 | 178 | 3.36803097 | 8.0103E-15 | hsa-miR-2392 |
| 2.32477115 | 7.2334E-09 | hsa-miR-196 | 7.25 | 3.34242613 | 0.01648809 | hsa-miR-21-5p |
| 2.3217821 | 0.01789156 | hsa-miR-126 | 9.5 | 3.34154468 | 0.00760677 | hsa-miR-3927-3p |
| 2.32129724 | 0.01944682 | hsa-miR-451 | 13.75 | 3.27283056 | 0.00287964 | hsa-miR-574-3p |
| 2.30804408 | 1.0053E-09 | hsa-miR-448 | 7 | 3.23014884 | 0.02721769 | hsa-miR-22-5p |
| 2.16698379 | 6.0981E-07 | hsa-miR-361 | 25.25 | 3.19256012 | 4.8857E-06 | hsa-miR-342-3p |
| 2.16014257 | 9.9027E-05 | hsa-miR-716 | 7.5 | 3.13036605 | 0.01400245 | hsa-miR-132-3p |
| 2.076147 | 1.3017E-19 | hsa-miR-477 | 6 | 3.0983316 | 0.03077097 | hsa-miR-6774-3p |
| 2.00538865 | 0.03089478 | hsa-miR-132 | 8 | 3.04304715 | 0.02457075 | hsa-miR-15b-3p |
| 1.98558104 | 7.314E-07 | hsa-miR-548 | 8.25 | 2.89259165 | 0.01497872 | hsa-miR-543 |
| 1.96465528 | 5.4803E-19 | hsa-miR-673 | 15.75 | 2.72803098 | 0.00541852 | hsa-miR-214-5p |
| 1.9021763 | 2.8274E-16 | hsa-miR-548 | 21.5 | 2.71787008 | 0.00135777 | hsa-miR-654-3p |
| 1.88819217 | 0.00027328 | hsa-miR-711 | 13.5 | 2.62538752 | 0.00287964 | hsa-miR-128-1-5p |
| 1.80724998 | 2.5422E-05 | hsa-miR-121 | 14.25 | 2.51499099 | 0.00718222 | hsa-miR-424-3p |
| 1.72940232 | 1.2961E-08 | hsa-miR-808 | 12.25 | 2.49425849 | 0.01043621 | hsa-miR-27a-5p |
| 1.72536229 | 0.01847763 | hsa-miR-132 | 96.75 | 2.46929432 | 7.251E-08 | hsa-miR-10396a-5p |
| 1.6830711 | 0.00109455 | hsa-miR-808 | 42 | 2.2393916 | 0.00219648 | hsa-miR-335-5p |
| 1.66233207 | 2.8705E-05 | hsa-miR-17- | 23 | 2.10112833 | 0.0126492 | hsa-miR-23a-5p |
| 1.603921 | 5.4163E-08 | hsa-miR-806 | 60.5 | 2.00403012 | 0.00360206 | hsa-miR-1296-5p |
| 1.58682887 | 2.641E-06 | hsa-miR-9-5 | 65.75 | 1.85361519 | 0.00122212 | hsa-miR-146b-3p |
| 1.58092808 | 0.00062076 | hsa-miR-450 | 106.25 | 1.72627988 | 0.00360206 | hsa-miR-3978 |
| 1.4790424 | 0.00181171 | hsa-miR-239 | 90.5 | 1.47014396 | 0.02286603 | hsa-let-7e-3p |
| 1.44961714 | 0.00102812 | hsa-miR-186 | 1040.25 | 1.42495503 | 0.00292377 | hsa-miR-452-5p |
| 1.44604152 | 0.00010218 | hsa-miR-148 | 766.5 | 1.37991254 | 0.00688031 | hsa-miR-126-5p |
| 1.39123832 | 0.04746683 | hsa-miR-181 | 18607.75 | 1.35104133 | 0.01662154 | hsa-miR-3148 |
| 1.38569489 | 0.01304828 | hsa-miR-25- | 17419.5 | 1.29082859 | 0.04939511 | hsa-miR-4435 |
| 1.36318464 | 0.00086639 | hsa-miR-23a | 1803.5 | -1.3343118 | 0.01539949 | hsa-miR-6070 |

| | | | | | | |
|---|---|---|---|---|---|---|
| 1.35288293 | 0.02112289 | hsa-miR-22- | 28627.5 | -1.352864 | 0.0110381 | hsa-miR-1910-5p |
| 1.33865172 | 0.01482144 | hsa-miR-31- | 4277.25 | -1.4061964 | 0.00505255 | hsa-miR-5579-3p |
| 1.29956953 | 0.00517731 | hsa-miR-191 | 4219.5 | -1.4281851 | 0.00287964 | hsa-miR-4282 |
| 1.21262907 | 0.01048572 | hsa-miR-941 | 184.25 | -1.4314798 | 0.01574579 | hsa-miR-146a-5p |
| -1.2227397 | 0.00517731 | hsa-miR-21- | 109325 | -1.4922972 | 0.00015964 | hsa-miR-450a-5p |
| -1.2361751 | 0.04746683 | hsa-miR-360 | 63.75 | -1.5142683 | 0.04939662 | hsa-miR-30a-5p |
| -1.2800863 | 0.04687873 | hsa-miR-561 | 78 | -1.5583473 | 0.02231887 | hsa-miR-7158-3p |
| -1.3044366 | 0.02132098 | hsa-miR-122 | 9213 | -1.5786022 | 6.8122E-05 | hsa-miR-3613-5p |
| -1.3111023 | 0.00128283 | hsa-miR-193 | 1137.5 | -1.5788346 | 2.4853E-05 | hsa-miR-28-5p |
| -1.3217516 | 0.01304828 | hsa-miR-500 | 33.75 | -1.5869535 | 0.03678329 | hsa-miR-3125 |
| -1.3669272 | 0.01138405 | hsa-miR-143 | 125.5 | -1.6059013 | 0.01925312 | hsa-miR-6131 |
| -1.3830731 | 0.00106192 | hsa-miR-125 | 5732 | -1.6085768 | 8.4893E-06 | hsa-miR-129-5p |
| -1.4451281 | 0.02699224 | hsa-miR-103 | 105.25 | -1.6287843 | 0.00482583 | hsa-miR-3691-3p |
| -1.4797463 | 0.02569176 | hsa-miR-428 | 54.25 | -1.6318674 | 0.03938519 | hsa-miR-493-3p |
| -1.4804027 | 0.01961769 | hsa-miR-99a | 96 | -1.6684781 | 0.00631401 | hsa-miR-10a-5p |
| -1.4853525 | 0.00023216 | hsa-miR-30a | 25622.5 | -1.7892492 | 1.0041E-08 | hsa-miR-1307-3p |
| -1.4982633 | 0.00476292 | hsa-miR-121 | 409 | -1.8314564 | 6.1711E-06 | hsa-miR-493-5p |
| -1.569815 | 0.00870042 | hsa-miR-374 | 214 | -1.8422983 | 6.8122E-05 | hsa-miR-193b-5p |
| -1.6273369 | 0.00091951 | hsa-miR-224 | 111 | -1.8583637 | 0.00057394 | hsa-miR-875-3p |
| -1.6364319 | 0.0190902 | hsa-miR-557 | 288 | -1.8664083 | 2.4731E-06 | hsa-miR-130b-5p |
| -1.660586 | 1.1466E-06 | hsa-miR-331 | 319.25 | -1.8966138 | 1.9318E-05 | hsa-miR-6728-5p |
| -1.7012723 | 0.00023096 | hsa-miR-30a | 450 | -1.9867436 | 7.2032E-08 | hsa-miR-4490 |
| -1.7245086 | 0.01304828 | hsa-miR-210 | 13.75 | -2.0043828 | 0.0490135 | hsa-let-7i-3p |
| -1.7305935 | 0.00517731 | hsa-miR-134 | 33.75 | -2.0262118 | 0.00135777 | hsa-miR-660-5p |
| -1.7445449 | 1.153E-09 | hsa-let-7a-3 | 84.75 | -2.0496708 | 0.00089325 | hsa-miR-578 |
| -1.7566834 | 1.4993E-05 | hsa-miR-181 | 12.75 | -2.1088626 | 0.04437787 | hsa-miR-342-5p |
| -1.8010571 | 0.00012518 | hsa-miR-452 | 42.25 | -2.1480577 | 0.00326769 | hsa-miR-486-5p |
| -1.8626245 | 1.5161E-05 | hsa-miR-409 | 29.5 | -2.1606378 | 0.00157386 | hsa-miR-103a-3p |
| -1.8707468 | 0.00815986 | hsa-miR-424 | 32 | -2.2105802 | 0.00025738 | hsa-miR-191-5p |
| -1.9083491 | 1.309E-12 | hsa-miR-374 | 10 | -2.2640415 | 0.03046791 | hsa-miR-1304-3p |
| -1.9175481 | 0.0049853 | hsa-miR-181 | 418.5 | -2.3883464 | 4.5594E-09 | hsa-miR-6799-5p |
| -1.9195194 | 0.00412943 | hsa-miR-453 | 21 | -2.3928826 | 0.00505255 | hsa-miR-1257 |
| -1.9355077 | 1.0934E-10 | hsa-miR-443 | 10 | -2.4093485 | 0.02404454 | hsa-miR-9900 |
| -2.0144054 | 0.00134816 | hsa-miR-673 | 11.5 | -2.4480242 | 0.0350585 | hsa-miR-1827 |
| -2.0308761 | 1.06E-05 | hsa-miR-671 | 29 | -2.4619128 | 0.00027177 | hsa-miR-197-3p |
| -2.0312456 | 0.00039749 | hsa-miR-194 | 12.75 | -2.5652198 | 0.00528058 | hsa-miR-421 |
| -2.0959051 | 0.00351835 | hsa-miR-30c | 47.25 | -2.6388605 | 0.00022607 | hsa-miR-3184-5p |
| -2.1085603 | 2.4051E-07 | hsa-miR-582 | 573.25 | -2.6695341 | 8.5366E-16 | hsa-miR-210-5p |
| -2.1313407 | 1.944E-13 | hsa-miR-366 | 43.25 | -2.8474188 | 1.5672E-05 | hsa-miR-369-3p |
| -2.133774 | 0.00184656 | hsa-miR-397 | 20.75 | -2.8608991 | 0.00446305 | hsa-miR-7846-3p |
| -2.1471463 | 3.1632E-07 | hsa-miR-444 | 55 | -3.053342 | 2.3858E-06 | hsa-miR-155-5p |
| -2.1792092 | 4.9337E-05 | hsa-let-7e-3 | 17.5 | -3.166803 | 0.00060424 | hsa-miR-4425 |
| -2.1813587 | 2.6452E-06 | hsa-miR-193 | 25.75 | -3.2280112 | 1.5672E-05 | hsa-miR-5089-5p |
| -2.1851025 | 2.3711E-10 | hsa-miR-125 | 80.75 | -3.2519268 | 1.5928E-07 | hsa-miR-4326 |
| -2.2015646 | 0.03680409 | hsa-miR-181 | 209.25 | -3.2557729 | 5.8495E-17 | hsa-miR-374a-5p |

| | | | | | | |
|---|---|---|---|---|---|---|
| -2.2477213 | 3.268E-05 | hsa-miR-680 | 6.5 | -3.2709134 | 0.02992893 | hsa-miR-106a-5p |
| -2.273509 | 0.03797896 | hsa-miR-342 | 7 | -3.5155756 | 0.02461962 | hsa-miR-335-3p |
| -2.2988821 | 3.1906E-05 | hsa-miR-125 | 20.75 | -3.7515807 | 2.642E-05 | hsa-miR-1299 |
| -2.3162157 | 1.1415E-08 | hsa-miR-125 | 43.75 | -3.8887033 | 3.5055E-09 | hsa-miR-589-3p |
| -2.3591578 | 0.00718581 | hsa-miR-127 | 46.75 | -4.0224347 | 5.592E-10 | hsa-miR-4654 |
| -2.5468153 | 0.00117075 | hsa-miR-316 | 25.75 | -4.0547672 | 4.6185E-06 | hsa-miR-4718 |
| -2.5771145 | 0.04198701 | hsa-miR-128 | 20.25 | -4.0563477 | 2.1525E-05 | hsa-miR-5682 |
| -2.5865731 | 0.03738485 | hsa-miR-887 | 8.5 | -4.2533738 | 0.00718222 | hsa-miR-2355-3p |
| -2.5918396 | 0.00152572 | hsa-miR-607 | 7.5 | -4.3382924 | 0.01492882 | hsa-miR-126-3p |
| -2.6043033 | 0.01304828 | hsa-miR-365 | 12.25 | -4.3506032 | 0.00023754 | hsa-miR-6734-5p |
| -2.7068196 | 1.9917E-16 | hsa-miR-430 | 5.25 | -4.436326 | 0.02437539 | hsa-miR-5197-3p |
| -2.7458661 | 0.04188479 | hsa-miR-25- | 13.75 | -4.4476729 | 0.00015964 | hsa-miR-136-5p |
| -2.7896585 | 1.5901E-09 | hsa-miR-394 | 22 | -4.6302375 | 7.9338E-05 | hsa-miR-1249-3p |
| -2.8909038 | 1.474E-07 | hsa-miR-469 | 4.25 | -4.6546132 | 0.04891084 | hsa-miR-542-3p |
| -2.9947433 | 0.0049853 | hsa-miR-100 | 21 | -4.7200132 | 5.3106E-06 | hsa-miR-548u |
| -3.0136169 | 0.00153892 | hsa-miR-674 | 59 | -4.7260548 | 2.392E-10 | hsa-miR-625-5p |
| -3.0293606 | 0.0016771 | hsa-miR-465 | 22.25 | -4.7781274 | 0.00013695 | |
| -3.1643624 | 0.04594969 | hsa-miR-612 | 9.25 | -4.895971 | 0.0111364 | |
| -3.1649071 | 0.0418568 | hsa-miR-688 | 7.5 | -5.1384022 | 0.00717162 | |
| -3.2681179 | 0.00186285 | hsa-miR-158 | 4.75 | -5.1854137 | 0.02739551 | |
| -3.6707281 | 3.1381E-05 | hsa-miR-605 | 7.5 | -5.1931986 | 0.01364133 | |
| -3.7079009 | 0.00175211 | hsa-miR-312 | 12.75 | -5.5938283 | 0.00011774 | |
| -3.7819707 | 5.6364E-16 | hsa-miR-471 | 6.75 | -5.6685894 | 0.00717162 | |
| -4.0535315 | 0.00870042 | hsa-miR-450 | 29.75 | -5.9485678 | 1.9343E-08 | |
| -4.5599465 | 0.00220602 | hsa-miR-797 | 100.75 | -6.2204826 | 1.9873E-14 | |
| -4.9748645 | 3.8877E-08 | hsa-miR-990 | 149 | -6.4068338 | 1.5377E-25 | |
| -5.0173402 | 1.5705E-07 | hsa-miR-27b | 343.75 | -6.5116381 | 4.2925E-31 | |
| -5.2888588 | 5.0661E-06 | hsa-miR-214 | 88.75 | -7.0670065 | 5.4654E-12 | |
| -5.3824861 | 1.0042E-12 | hsa-miR-23a | 29.75 | -9.5784605 | 1.319E-09 | |
| -5.4761501 | 0.04434172 | hsa-miR-29b | 9 | -9.6839981 | 0.00094727 | |
| -6.4803468 | 0.00127007 | hsa-miR-684 | 9.5 | -10.211462 | 0.00064189 | |
| -6.5759334 | 0.0049853 | hsa-miR-601 | 29.25 | -11.405132 | 1.0923E-09 | |
| -7.2555598 | 0.01375883 | hsa-miR-677 | 10.75 | -11.53691 | 0.00025202 | |
| -7.7633529 | 0.01087258 | hsa-miR-33b | 11.75 | -12.595556 | 0.0001205 | |
| -7.8812243 | 0.00233986 | hsa-miR-442 | 35.75 | -12.603224 | 3.5467E-11 | |
| -8.3518806 | 5.4586E-15 | hsa-miR-222 | 19.75 | -13.151717 | 1.4569E-06 | |
| -8.4562195 | 0.04928831 | hsa-miR-715 | 5.25 | -13.472712 | 0.0148829 | |
| -8.5472576 | 0.00760846 | hsa-miR-465 | 44 | -14.138813 | 5.9209E-14 | |
| -8.6306948 | 1.0464E-05 | hsa-miR-672 | 12.75 | -19.171276 | 0.00017983 | |
| -9.3171749 | 0.0374062 | hsa-miR-471 | 8.25 | -21.282368 | 0.00403715 | |
| -11.46161 | 4.4301E-08 | hsa-miR-673 | 115.5 | -21.598701 | 1.0754E-23 | |
| -11.911404 | 0.01677273 | hsa-miR-27a | 59.75 | -34.001932 | 8.0103E-15 | |
| -13.092161 | 0.00080726 | | | | | |
| -17.323387 | 7.7745E-26 | | | | | |

### .4p1 v C 6HPI Ad14p1 v C 12HP

| Max group mean | Fold change | FDR p-value | Identifier | Max group mean | Fold change |
|---|---|---|---|---|---|
| 588.25 | 3.054929409 | 1.18459E-52 | hsa-miR-10524-5p | 5.25 | 48.18088227 |
| 56169.75 | 2.099939664 | 1.44066E-32 | hsa-miR-6841-5p | 4.75 | 43.69201414 |
| 115.5 | -8.75822958 | 1.5653E-28 | hsa-miR-375-5p | 7.5 | 23.6377344 |
| 157.75 | 2.771765589 | 1.1418E-22 | hsa-miR-4724-5p | 5.5 | 10.53291573 |
| 21568.25 | 1.836453145 | 7.36195E-22 | hsa-miR-155-5p | 5.25 | 10.06436869 |
| 369 | -2.104854288 | 5.14021E-19 | hsa-miR-6799-5p | 4 | 7.750974502 |
| 249.5 | 2.550402213 | 1.29897E-18 | hsa-miR-7846-3p | 3.5 | 6.786497169 |
| 79.75 | 2.928905485 | 5.67447E-18 | hsa-miR-520f-3p | 8.5 | 6.267494634 |
| 16352.75 | 1.701601349 | 8.2441E-17 | hsa-miR-888-5p | 10.5 | 5.901395702 |
| 93.75 | 2.715419171 | 9.09511E-17 | hsa-miR-493-3p | 7.75 | 5.714225717 |
| 28.25 | 6.987413203 | 8.72755E-14 | hsa-miR-1197 | 7.5 | 5.535452904 |
| 61.25 | 2.736039686 | 1.87535E-13 | hsa-miR-649 | 6.5 | 5.201515674 |
| 142.75 | -2.690101075 | 2.70941E-13 | hsa-miR-4431 | 10.5 | 4.778946472 |
| 48.5 | 2.968638337 | 1.52202E-12 | hsa-miR-200a-5p | 4 | 4.311523854 |
| 5732 | -1.568961286 | 7.60003E-12 | hsa-miR-1292-5p | 5.75 | 4.261202841 |
| 509 | -1.695906949 | 1.26997E-11 | hsa-miR-889-3p | 32.5 | 3.809385563 |
| 82.5 | 2.336591878 | 1.36153E-10 | hsa-miR-5187-3p | 16.75 | 3.714663167 |
| 84268.25 | 1.517333352 | 1.36153E-10 | hsa-miR-152-5p | 8.5 | 3.538366921 |
| 17346.75 | 1.509703989 | 2.78364E-10 | hsa-miR-323b-3p | 5.25 | 3.515156882 |
| 1803.5 | -1.543134529 | 2.80411E-10 | hsa-miR-676-3p | 7.25 | 3.340330724 |
| 729.75 | -1.643676083 | 1.25547E-09 | hsa-miR-409-3p | 74.25 | 2.976843356 |
| 27.25 | 3.489861598 | 2.71634E-09 | hsa-miR-668-5p | 13.25 | 2.964177489 |
| 107.75 | 1.972132727 | 3.0299E-09 | hsa-miR-1253 | 7 | 2.930499354 |
| 1137.5 | -1.687505979 | 5.84466E-09 | hsa-miR-411-5p | 99.75 | 2.645894723 |
| 235.25 | -1.746235954 | 8.14767E-09 | hsa-miR-127-3p | 545.25 | 2.61641719 |
| 2481.25 | -1.480353686 | 9.10858E-09 | hsa-miR-9-5p | 82.5 | 2.541351517 |
| 657.5 | -1.531830525 | 1.89475E-08 | hsa-miR-410-3p | 77.5 | 2.526715314 |
| 4310.5 | 1.452584931 | 2.27473E-08 | hsa-miR-5683 | 9.75 | 2.525168382 |
| 166 | 1.892402554 | 2.70057E-08 | hsa-miR-493-5p | 10.25 | 2.394865607 |
| 37.75 | 2.866776825 | 1.14031E-07 | hsa-miR-4286 | 43 | 2.36248534 |
| 3259.5 | -1.43474329 | 1.14031E-07 | hsa-miR-375-3p | 134 | 2.206118119 |
| 1045.75 | -1.469103663 | 1.14031E-07 | hsa-miR-5009-5p | 80.25 | 2.195025458 |
| 238.75 | -1.870641557 | 1.47379E-07 | hsa-miR-128-1-5p | 37.5 | 2.183078867 |
| 40.5 | 2.568408109 | 2.19412E-07 | hsa-miR-7706 | 80.5 | 2.083535991 |
| 964 | 1.446297937 | 2.19412E-07 | hsa-miR-106b-3p | 97.5 | 2.000450434 |
| 86.75 | 1.943049065 | 2.83496E-07 | hsa-miR-136-3p | 26.75 | 1.859073197 |
| 65.75 | 2.000726613 | 2.98548E-07 | hsa-miR-4714-5p | 13 | 1.855238323 |
| 435.75 | 1.535186429 | 6.95813E-07 | hsa-miR-671-3p | 45 | 1.836003645 |
| 45.75 | 2.080184698 | 7.44258E-07 | hsa-miR-300 | 121 | 1.833451672 |
| 54 | 2.180987488 | 8.44694E-07 | hsa-miR-192-5p | 50585.75 | 1.794831144 |
| 20.75 | -6.03582443 | 1.64355E-06 | hsa-miR-1285-3p | 37.75 | 1.749551554 |

| | | | | | |
|---|---|---|---|---|---|
| 496.75 | 1.440238129 | 1.7107E-06 | hsa-miR-378a-3p | 1236.5 | 1.711691901 |
| 22 | -19.23528517 | 2.14588E-06 | hsa-miR-1304-3p | 31.5 | 1.655151706 |
| 41.5 | 2.270425632 | 2.33955E-06 | hsa-miR-193a-5p | 58.5 | 1.606996015 |
| 655.75 | 1.411544156 | 3.39786E-06 | hsa-miR-12135 | 531 | 1.586779549 |
| 56.25 | 1.928686862 | 4.27254E-06 | hsa-miR-654-3p | 69.5 | 1.529412904 |
| 60.75 | 1.813070181 | 4.62705E-06 | hsa-miR-143-3p | 171 | 1.503249496 |
| 371 | 1.540245881 | 5.50223E-06 | hsa-miR-181a-5p | 17741 | 1.421533842 |
| 64.25 | 1.854318407 | 9.15249E-06 | hsa-miR-182-5p | 14467 | 1.411548043 |
| 94.5 | -1.902606136 | 9.69868E-06 | hsa-miR-151a-3p | 4446 | 1.396606483 |
| 319.25 | -1.70611757 | 1.32569E-05 | hsa-miR-3184-5p | 671 | 1.382368804 |
| 9.5 | 7.881583668 | 2.25164E-05 | hsa-miR-7974 | 116 | 1.368007819 |
| 84.75 | -1.773864984 | 4.74478E-05 | hsa-miR-138-5p | 657.5 | -1.375758231 |
| 59 | -3.336407084 | 6.7033E-05 | hsa-miR-196a-5p | 128 | -1.418778586 |
| 67.25 | -1.827758958 | 0.000104586 | hsa-miR-151a-5p | 2481.25 | -1.442020793 |
| 422 | 1.363129771 | 0.000108543 | hsa-miR-28-5p | 241.25 | -1.484718457 |
| 327.25 | -1.390200542 | 0.000198605 | hsa-miR-5682 | 99.75 | -1.500378318 |
| 42.25 | -2.198980991 | 0.000198605 | hsa-miR-125a-5p | 5732 | -1.524561848 |
| 9 | 4.609942673 | 0.000208007 | hsa-miR-22-3p | 28627.5 | -1.578717343 |
| 67 | -2.143179493 | 0.000212321 | hsa-miR-210-3p | 270 | -1.586107041 |
| 110775 | 1.290856687 | 0.000230607 | hsa-miR-574-3p | 61.25 | -1.688464273 |
| 50 | 1.629450175 | 0.000720767 | hsa-miR-3155a | 369 | -1.702468175 |
| 61.25 | -1.922283235 | 0.000829117 | hsa-miR-374a-5p | 111.75 | -1.75047078 |
| 31.75 | -2.126611047 | 0.000914985 | hsa-miR-224-5p | 111 | -1.799233523 |
| 55.5 | -1.865404774 | 0.000951298 | hsa-miR-365a-3p | 238.75 | -1.803143383 |
| 54 | 1.656957664 | 0.00102281 | hsa-miR-29a-3p | 1045.75 | -1.822781522 |
| 10.75 | -13.46127055 | 0.001046694 | hsa-miR-23a-3p | 1803.5 | -1.833418116 |
| 51.25 | -1.758124373 | 0.001222624 | hsa-let-7a-3p | 84.75 | -1.835025369 |
| 64.5 | 1.551148756 | 0.001449729 | hsa-miR-140-5p | 33.75 | -1.841263031 |
| 88.75 | -1.668474411 | 0.001552424 | hsa-miR-1307-5p | 509 | -1.844188772 |
| 62.25 | 1.522131806 | 0.00158729 | hsa-miR-345-5p | 235.25 | -1.873886957 |
| 30.5 | 1.952646206 | 0.001626843 | hsa-miR-450a-5p | 26.25 | -1.962383749 |
| 32 | -1.969239262 | 0.001663863 | hsa-miR-24-3p | 729.75 | -1.962492228 |
| 59.75 | -1.749809869 | 0.001726218 | hsa-miR-193a-3p | 1137.5 | -1.971025347 |
| 127.5 | 1.525733982 | 0.001904849 | hsa-let-7e-3p | 17.5 | -2.017029441 |
| 56.75 | -1.633596495 | 0.002030203 | hsa-let-7d-3p | 26.75 | -2.088460447 |
| 29.75 | -2.036334229 | 0.002675513 | hsa-miR-331-3p | 319.25 | -2.089539324 |
| 18.5 | -2.402434335 | 0.002993996 | hsa-miR-22-5p | 31.75 | -2.092955454 |
| 10.25 | 2.679966721 | 0.00326778 | hsa-miR-200b-3p | 13 | -2.16420509 |
| 20.75 | -2.759403283 | 0.003423449 | hsa-miR-33b-3p | 11.75 | -2.171651994 |
| 17.5 | -2.372672046 | 0.005088075 | hsa-miR-31-3p | 94.5 | -2.197238955 |
| 42.25 | -1.676418361 | 0.005208055 | hsa-miR-513a-5p | 19 | -2.265469261 |
| 195.75 | 1.389378667 | 0.005415414 | hsa-miR-450b-5p | 67.25 | -2.273445239 |
| 5 | 18.63472459 | 0.005670558 | hsa-miR-452-5p | 42.25 | -2.288597715 |
| 10 | -3.616244916 | 0.005804252 | hsa-miR-324-3p | 9.75 | -2.303248031 |
| 7.5 | -9.350584387 | 0.006177602 | hsa-miR-326 | 27.5 | -2.314767863 |

| | | | | | |
|---|---|---|---|---|---|
| 6 | 3.825983329 | 0.006417431 | hsa-miR-578 | 10.5 | -2.31852434 |
| 288 | 1.279280034 | 0.006417431 | hsa-miR-2355-5p | 13.5 | -2.362922714 |
| 54.25 | -1.561852653 | 0.006601283 | hsa-miR-125a-3p | 43.75 | -2.371308598 |
| 58.5 | 1.451582923 | 0.007310918 | hsa-miR-487a-5p | 12.25 | -2.393468921 |
| 26.25 | -1.878005963 | 0.007774057 | hsa-miR-27a-5p | 59.75 | -2.39610817 |
| 25622.5 | 1.216963154 | 0.008017374 | hsa-miR-23a-5p | 29.75 | -2.432377448 |
| 22 | -1.999970386 | 0.008017374 | hsa-miR-582-5p | 142.75 | -2.472884561 |
| 8.5 | -4.033674136 | 0.008726532 | hsa-miR-4446-3p | 55 | -2.533770474 |
| 241.25 | -1.313972414 | 0.008951781 | hsa-miR-193b-5p | 25.75 | -2.862037676 |
| 4.25 | 6.689729818 | 0.00948781 | hsa-miR-4425 | 35.75 | -2.951290464 |
| 13.25 | 2.108418065 | 0.009513116 | hsa-let-7i-3p | 17.5 | -3.053714967 |
| 4.75 | 4.717306197 | 0.009905534 | hsa-miR-5089-5p | 7.25 | -3.33457132 |
| 12 | -2.725383898 | 0.010109989 | hsa-miR-6741-5p | 59 | -3.525322758 |
| 5 | 4.192200384 | 0.010737139 | hsa-miR-192-3p | 10.5 | -4.234113822 |
| 20541.75 | 1.208786472 | 0.010924701 | hsa-miR-4718 | 6.75 | -4.243825593 |
| 96.5 | 1.346902424 | 0.01131455 | hsa-miR-342-5p | 7 | -4.400613668 |
| 9.25 | 2.419874001 | 0.011377144 | hsa-miR-9900 | 149 | -4.816584447 |
| 25.75 | -1.88876832 | 0.011573122 | hsa-miR-134-3p | 5 | -4.99197571 |
| 9.25 | -3.864934748 | 0.011573122 | hsa-miR-2110 | 6.5 | -5.008201314 |
| 13.25 | -2.459087576 | 0.011779686 | hsa-miR-141-5p | 5.75 | -5.718248061 |
| 12.75 | -2.517820106 | 0.011822604 | hsa-miR-6070 | 7.5 | -7.423616315 |
| 4 | 14.98888442 | 0.012076181 | hsa-miR-6784-5p | 6.5 | -9.059387782 |
| 17.5 | -2.094163757 | 0.012687572 | hsa-miR-6737-5p | 115.5 | -9.494417894 |
| 56.5 | 1.448802954 | 0.013651338 | hsa-miR-6774-3p | 10.75 | -10.5871185 |
| 10.5 | -2.800582168 | 0.014503957 | hsa-miR-3940-5p | 22 | -16.81738267 |
| 7 | -4.821712438 | 0.01453124 | | | |
| 175 | 1.353028042 | 0.016322977 | | | |
| 1881 | 1.204763125 | 0.016815549 | | | |
| 4219.5 | 1.19927746 | 0.017181979 | | | |
| 27 | 1.585287751 | 0.017676901 | | | |
| 3.5 | 7.800047988 | 0.017823692 | | | |
| 80.75 | -1.389868119 | 0.018921852 | | | |
| 149 | -2.479379695 | 0.018921852 | | | |
| 52.5 | -1.524702603 | 0.019187432 | | | |
| 84.5 | -1.418611493 | 0.019272586 | | | |
| 128 | 1.380167899 | 0.019337886 | | | |
| 566.5 | 1.268632115 | 0.021939358 | | | |
| 13.75 | -2.154982023 | 0.021939358 | | | |
| 5.25 | 3.358382989 | 0.022186742 | | | |
| 3.25 | 7.260437806 | 0.023680936 | | | |
| 3.25 | 7.257327375 | 0.02404583 | | | |
| 35.75 | -2.043021753 | 0.024051002 | | | |
| 7.25 | -3.446670857 | 0.024162883 | | | |
| 17 | -1.95775673 | 0.024247597 | | | |
| 111.75 | -1.397277477 | 0.024262906 | | | |

| | | |
|---|---|---|
| 1451.5 | -1.195647349 | 0.02593104 |
| 176.25 | -1.373216113 | 0.02593104 |
| 21 | 1.693107526 | 0.028997656 |
| 12.25 | -2.31641731 | 0.030767705 |
| 5.5 | -4.899300342 | 0.031106567 |
| 6.75 | -3.212284992 | 0.039203905 |
| 99.75 | -1.353381289 | 0.042820926 |
| 4.5 | -9.535056514 | 0.045907802 |
| 32 | -1.533134065 | 0.047570521 |
| 11.5 | -2.134403945 | 0.047570521 |
| 2.75 | 6.18102734 | 0.047854391 |
| 4.25 | 3.09775598 | 0.047854391 |
| 7.25 | -3.019291789 | 0.047854391 |
| 23.5 | -1.642125027 | 0.048366114 |
| 12.25 | -2.027163044 | 0.048366114 |
| 7.5 | -2.721101328 | 0.048366114 |

PI Ad14p1 v C 24HPI Ad14

| FDR p-value | Identifier | Max group mean | Fold change | FDR p-value | Identifier |
|---|---|---|---|---|---|
| 0.041380785 | hsa-miR-6841-5p | 17.5 | 146.6999358 | 0.002256142 | hsa-miR-6841-5p |
| 0.04904175 | hsa-miR-10524-5p | 14.75 | 124.4069907 | 0.003214475 | hsa-miR-10524-5p |
| 0.002087178 | hsa-miR-4661-3p | 12 | 35.31082274 | 0.000173434 | hsa-miR-518a-5p |
| 0.005232353 | hsa-miR-518a-5p | 10.25 | 30.16048955 | 0.000312876 | hsa-miR-4661-3p |
| 0.006542093 | hsa-miR-4668-3p | 6.5 | 19.26972423 | 0.002749831 | hsa-miR-3128 |
| 0.023551115 | hsa-miR-155-5p | 10.75 | 19.13164636 | 6.07305E-05 | hsa-miR-1197 |
| 0.041121933 | hsa-miR-1197 | 25.5 | 17.46608391 | 6.18522E-11 | hsa-miR-4668-3p |
| 0.000495066 | hsa-miR-5571-5p | 4.75 | 14.18721837 | 0.009591098 | hsa-miR-4490 |
| 8.15796E-05 | hsa-miR-375-5p | 4.75 | 14.18648751 | 0.009364267 | hsa-miR-4279 |
| 0.002178744 | hsa-miR-668-5p | 58 | 12.67561642 | 1.45871E-21 | hsa-miR-155-5p |
| 0.001295484 | hsa-miR-4279 | 12.5 | 12.40892696 | 4.3678E-06 | hsa-miR-4724-5p |
| 0.011244053 | hsa-miR-4490 | 4 | 12.00907028 | 0.017495062 | hsa-miR-668-5p |
| 0.000168304 | hsa-miR-4714-5p | 85.75 | 11.40432052 | 9.93041E-35 | hsa-miR-5571-5p |
| 0.044404514 | hsa-miR-4724-5p | 5 | 9.023828617 | 0.006140873 | hsa-miR-888-5p |
| 0.012564097 | hsa-miR-6799-5p | 4.5 | 8.145576632 | 0.010081031 | hsa-miR-520f-3p |
| 1.71614E-07 | hsa-miR-4431 | 18 | 7.665048309 | 1.77381E-08 | hsa-miR-4714-5p |
| 1.73267E-05 | hsa-miR-888-5p | 14.25 | 7.532660828 | 3.45652E-06 | hsa-miR-6799-5p |
| 0.006505105 | hsa-miR-3128 | 15.5 | 7.269181339 | 9.31326E-07 | hsa-miR-4462 |
| 0.041121933 | hsa-miR-520f-3p | 9.25 | 6.396144743 | 0.000135823 | hsa-miR-3125 |
| 0.017805593 | hsa-miR-6870-3p | 3.25 | 5.959980403 | 0.044996778 | hsa-miR-4759 |
| 2.01288E-12 | hsa-miR-369-3p | 11.25 | 5.941089308 | 2.40958E-05 | hsa-miR-4286 |
| 0.003987982 | hsa-miR-1253 | 15 | 5.895000333 | 7.28844E-06 | hsa-miR-1253 |
| 0.021478913 | hsa-miR-4496 | 9.25 | 5.545261848 | 0.00018734 | hsa-miR-548c-3p |
| 3.81278E-11 | hsa-miR-885-3p | 3.75 | 4.89416878 | 0.033499608 | hsa-miR-4299 |
| 1.0487E-13 | hsa-miR-493-3p | 7 | 4.863988171 | 0.002999772 | hsa-miR-215-3p |
| 2.83841E-10 | hsa-miR-4722-5p | 4.25 | 4.306561423 | 0.027730732 | hsa-miR-1245b-3p |
| 5.03204E-09 | hsa-miR-6792-5p | 4.25 | 4.305742337 | 0.027730732 | hsa-miR-369-3p |
| 0.018455906 | hsa-miR-431-5p | 8 | 4.244035403 | 0.001580028 | hsa-miR-33a-3p |
| 0.01603807 | hsa-miR-4286 | 81.25 | 4.149316235 | 3.61666E-15 | hsa-miR-1261 |
| 1.17977E-05 | hsa-miR-1321 | 10.25 | 4.007745297 | 0.000512282 | hsa-miR-518a-3p |
| 2.26937E-09 | hsa-miR-889-3p | 36.75 | 3.953129941 | 3.38194E-11 | hsa-miR-5683 |
| 6.07653E-07 | hsa-miR-1245b-3p | 14.75 | 3.77844637 | 2.18854E-05 | hsa-miR-646 |
| 0.000103444 | hsa-miR-129-5p | 4.5 | 3.728211717 | 0.046302955 | hsa-miR-3156-3p |
| 2.74147E-06 | hsa-miR-649 | 5.25 | 3.669213489 | 0.019238167 | hsa-miR-889-3p |
| 2.318E-06 | hsa-miR-5187-3p | 16.75 | 3.49006048 | 1.20257E-05 | hsa-miR-4431 |
| 0.012329875 | hsa-miR-5683 | 13.75 | 3.334620395 | 0.000164261 | hsa-miR-431-5p |
| 0.049905117 | hsa-miR-556-5p | 10.5 | 3.257557529 | 0.000954223 | hsa-miR-3649 |
| 0.00776634 | hsa-miR-548n | 8.5 | 3.066738886 | 0.005319228 | hsa-miR-520g-5p |
| 0.00025867 | hsa-miR-410-3p | 98.75 | 3.047260195 | 1.32413E-20 | hsa-miR-1301-5p |
| 2.2772E-05 | hsa-miR-136-5p | 5 | 3.0340709 | 0.048927873 | hsa-miR-12120 |
| 0.007468368 | hsa-miR-9-5p | 104 | 3.001222388 | 6.24795E-21 | hsa-miR-9-5p |

| | | | | | |
|---|---|---|---|---|---|
| 2.14773E-05 | hsa-miR-300 | 202 | 2.999296831 | 1.20287E-31 | hsa-miR-492 |
| 0.035604584 | hsa-miR-3156-3p | 26.25 | 2.897202745 | 3.05926E-07 | hsa-miR-410-3p |
| 0.010941786 | hsa-miR-127-3p | 654.25 | 2.896955364 | 3.77608E-39 | hsa-miR-8087 |
| 0.002681342 | hsa-miR-152-5p | 7.25 | 2.850131987 | 0.017495062 | hsa-miR-4474-3p |
| 0.01603807 | hsa-miR-485-3p | 8.25 | 2.758440267 | 0.010081031 | hsa-miR-1266-5p |
| 0.04904175 | hsa-miR-143-3p | 295 | 2.594451698 | 2.10397E-18 | hsa-miR-493-5p |
| 0.017127771 | hsa-miR-1266-5p | 7 | 2.534670195 | 0.032510208 | hsa-miR-4511 |
| 0.014151984 | hsa-miR-3620-3p | 9.25 | 2.526504153 | 0.014057157 | hsa-miR-106b-3p |
| 0.034868963 | hsa-miR-146b-3p | 11.25 | 2.468587536 | 0.007004278 | hsa-miR-300 |
| 0.038093335 | hsa-miR-106b-3p | 128.75 | 2.4319943 | 1.30004E-15 | hsa-miR-8066 |
| 0.032110946 | hsa-miR-487b-3p | 7.75 | 2.415119424 | 0.027850742 | hsa-miR-1276 |
| 0.032110946 | hsa-miR-411-5p | 97.75 | 2.387299286 | 2.61085E-14 | hsa-miR-127-3p |
| 0.02207209 | hsa-miR-375-3p | 150.5 | 2.323101417 | 9.98662E-18 | hsa-miR-1321 |
| 0.012018506 | hsa-miR-3654 | 8.75 | 2.256215316 | 0.032024731 | hsa-miR-375-3p |
| 0.006429775 | hsa-miR-6131 | 17 | 2.251105128 | 0.005530072 | hsa-miR-7-5p |
| 0.01143671 | hsa-miR-5009-5p | 88 | 2.211063472 | 1.62861E-11 | hsa-miR-1285-3p |
| 0.004134013 | hsa-miR-493-5p | 10 | 2.196505398 | 0.020522929 | hsa-miR-3620-3p |
| 0.000513279 | hsa-miR-132-3p | 84.5 | 2.195212032 | 1.41582E-08 | hsa-miR-5187-3p |
| 0.000202706 | hsa-miR-409-3p | 56.25 | 2.117282355 | 7.14421E-07 | hsa-miR-7706 |
| 0.014523315 | hsa-miR-193a-5p | 80 | 2.069680708 | 4.82126E-07 | hsa-miR-1304-3p |
| 1.58946E-05 | hsa-miR-136-3p | 31.25 | 2.037495789 | 0.000164261 | hsa-miR-5009-5p |
| 0.001485711 | hsa-miR-7706 | 82.25 | 1.98769674 | 8.73246E-09 | hsa-miR-411-5p |
| 0.000324401 | hsa-miR-1285-3p | 45.5 | 1.981716469 | 3.80249E-05 | hsa-miR-2682-5p |
| 7.82898E-06 | hsa-miR-378a-3p | 1522.25 | 1.980477048 | 8.7415E-25 | hsa-miR-589-3p |
| 1.65824E-06 | hsa-miR-7-5p | 37 | 1.979920368 | 0.000114968 | hsa-miR-4677-3p |
| 2.0997E-06 | hsa-miR-2682-5p | 10.75 | 1.974940757 | 0.040240253 | hsa-miR-548a-5p |
| 0.000244086 | hsa-miR-6890-3p | 17.5 | 1.936167521 | 0.008379084 | hsa-miR-143-3p |
| 0.018455906 | hsa-miR-7158-3p | 36.75 | 1.855449828 | 0.000250412 | hsa-miR-7113-3p |
| 2.1584E-08 | hsa-miR-3184-5p | 932.75 | 1.840476363 | 5.78808E-13 | hsa-miR-378a-3p |
| 6.07653E-07 | hsa-miR-548a-5p | 14 | 1.822228924 | 0.042296983 | hsa-miR-409-3p |
| 0.021478913 | hsa-miR-148a-3p | 987.25 | 1.821215893 | 2.50666E-18 | hsa-miR-148a-3p |
| 1.39599E-07 | hsa-miR-101-3p | 506.75 | 1.810346156 | 9.39113E-16 | hsa-miR-3613-5p |
| 1.91602E-07 | hsa-miR-8087 | 34.25 | 1.803678914 | 0.013659231 | hsa-miR-25-3p |
| 0.043056107 | hsa-miR-12135 | 650 | 1.790881685 | 2.44199E-11 | hsa-miR-4499 |
| 0.014831213 | hsa-miR-532-5p | 178.75 | 1.770488641 | 4.51471E-10 | hsa-miR-7158-3p |
| 3.00264E-08 | hsa-miR-671-3p | 45.25 | 1.733891468 | 0.00057178 | hsa-miR-532-5p |
| 0.001485711 | hsa-miR-181a-5p | 23304.25 | 1.727513375 | 1.19346E-10 | hsa-miR-193a-5p |
| 0.032110946 | hsa-miR-25-3p | 22837.75 | 1.724822252 | 6.85668E-18 | hsa-miR-3176 |
| 0.043056107 | hsa-miR-8066 | 50.5 | 1.711005143 | 0.002104529 | hsa-miR-4492 |
| 3.26167E-06 | hsa-miR-3615 | 49.25 | 1.680177694 | 0.002158321 | hsa-miR-1269a |
| 0.006559594 | hsa-miR-192-5p | 49640.75 | 1.614386073 | 6.95434E-07 | hsa-miR-181a-5p |
| 1.51158E-05 | hsa-miR-1269a | 33 | 1.543605337 | 0.016829866 | hsa-miR-3184-5p |
| 0.000104005 | hsa-miR-1304-3p | 31 | 1.531627195 | 0.036517335 | hsa-miR-192-5p |
| 0.049905117 | hsa-miR-340-5p | 321.5 | 1.518095902 | 2.67957E-07 | hsa-miR-132-3p |
| 0.001485711 | hsa-miR-769-5p | 471 | 1.442357508 | 1.95135E-06 | hsa-miR-101-3p |

| | | | | | |
|---|---|---|---|---|---|
| 0.042117758 | hsa-miR-34a-5p | 428.5 | 1.426575862 | 5.91449E-06 | hsa-miR-186-5p |
| 0.017805593 | hsa-miR-543 | 67.5 | 1.371700476 | 0.019238167 | hsa-miR-340-5p |
| 5.6427E-05 | hsa-miR-1827 | 63.25 | 1.340754088 | 0.044996778 | hsa-miR-671-3p |
| 0.031425168 | hsa-miR-182-5p | 14762.5 | 1.33320864 | 2.18854E-05 | hsa-miR-182-5p |
| 9.58343E-05 | hsa-miR-1307-3p | 111.75 | 1.318928431 | 0.014740977 | hsa-miR-769-5p |
| 0.000206753 | hsa-miR-27a-3p | 84171.25 | 1.318431754 | 4.61125E-05 | hsa-miR-12135 |
| 3.75739E-10 | hsa-miR-186-5p | 937 | 1.317888533 | 0.00018734 | hsa-miR-151a-3p |
| 0.001485711 | hsa-miR-561-5p | 92.5 | 1.317142395 | 0.02375061 | hsa-miR-126-5p |
| 2.78572E-05 | hsa-miR-423-3p | 861 | 1.311988298 | 0.015366841 | hsa-miR-194-5p |
| 2.30311E-05 | hsa-miR-151a-3p | 4498 | 1.307721479 | 0.000114927 | hsa-miR-28-5p |
| 0.000406054 | hsa-miR-126-5p | 215.25 | 1.307377427 | 0.004177148 | hsa-miR-7974 |
| 0.029126038 | hsa-miR-28-3p | 3670 | 1.263935424 | 0.001033302 | hsa-miR-151a-5p |
| 0.001680462 | hsa-miR-22-3p | 28627.5 | -1.27779398 | 0.000396665 | hsa-miR-1297 |
| 0.002792285 | hsa-miR-1297 | 6967.25 | -1.288975787 | 0.000259209 | hsa-miR-181a-2-3p |
| 0.019130172 | hsa-miR-28-5p | 241.25 | -1.289974109 | 0.007084033 | hsa-miR-345-5p |
| 0.016301813 | hsa-miR-210-3p | 270 | -1.295996532 | 0.004648724 | hsa-miR-22-3p |
| 0.00031005 | hsa-miR-5682 | 99.75 | -1.298478382 | 0.044076664 | hsa-miR-4446-3p |
| 0.045411897 | hsa-miR-345-5p | 235.25 | -1.325765094 | 0.009907573 | hsa-miR-4659a-5p |
| 0.019130172 | hsa-miR-151a-5p | 2481.25 | -1.361898482 | 0.001306185 | hsa-miR-3155a |
| 0.021805961 | hsa-miR-181a-2-3p | 418.5 | -1.384808561 | 6.00575E-05 | hsa-miR-4502 |
| 0.005671851 | hsa-miR-196a-5p | 128 | -1.392669187 | 0.019957831 | hsa-miR-30c-1-3p |
| 0.010941786 | hsa-miR-335-3p | 176.25 | -1.424016983 | 0.000304488 | hsa-miR-134-5p |
| 1.44039E-18 | hsa-miR-214-5p | 88.75 | -1.445275543 | 0.042296983 | hsa-miR-210-3p |
| 0.000661193 | hsa-miR-342-3p | 55.5 | -1.464159302 | 0.016351721 | hsa-miR-3606-3p |
| 1.04069E-06 | hsa-miR-224-5p | 111 | -1.469042548 | 0.003518159 | hsa-miR-374a-5p |
| | hsa-miR-197-3p | 84.5 | -1.472807419 | 0.002158321 | hsa-miR-29a-3p |
| | hsa-miR-29a-3p | 1045.75 | -1.529585236 | 1.63457E-09 | hsa-miR-224-5p |
| | hsa-miR-3155a | 369 | -1.572403617 | 1.8587E-08 | hsa-miR-24-3p |
| | hsa-miR-181a-3p | 209.25 | -1.600251101 | 2.76551E-07 | hsa-miR-181a-3p |
| | hsa-miR-374a-5p | 111.75 | -1.612960293 | 1.84329E-05 | hsa-miR-15b-3p |
| | hsa-miR-3165 | 25.75 | -1.617165385 | 0.028931576 | hsa-miR-125b-1-3p |
| | hsa-miR-125a-5p | 5732 | -1.618661399 | 5.72803E-07 | hsa-miR-574-3p |
| | hsa-miR-24-3p | 729.75 | -1.621130966 | 2.40094E-11 | hsa-let-7i-3p |
| | hsa-miR-193b-5p | 25.75 | -1.635440065 | 0.048519654 | hsa-miR-125a-5p |
| | hsa-miR-320e | 38.75 | -1.652276373 | 0.006464854 | hsa-miR-23a-3p |
| | hsa-miR-450b-5p | 67.25 | -1.66129945 | 0.001279053 | hsa-miR-365a-3p |
| | hsa-miR-3606-3p | 63.75 | -1.673975099 | 0.000682994 | hsa-miR-450a-5p |
| | hsa-miR-365a-3p | 238.75 | -1.682838958 | 3.517E-09 | hsa-let-7d-3p |
| | hsa-miR-574-3p | 61.25 | -1.683641297 | 0.001522066 | hsa-miR-31-3p |
| | hsa-miR-15b-3p | 51.25 | -1.733374782 | 0.004573892 | hsa-miR-214-5p |
| | hsa-let-7a-3p | 84.75 | -1.747452056 | 6.57125E-06 | hsa-miR-3165 |
| | hsa-miR-542-3p | 23.5 | -1.812777531 | 0.032929171 | hsa-miR-331-3p |
| | hsa-miR-23a-3p | 1803.5 | -1.867717201 | 5.56887E-21 | hsa-miR-193a-3p |
| | hsa-miR-424-3p | 32 | -1.876472569 | 0.005192507 | hsa-let-7a-3p |
| | hsa-let-7i-3p | 17.5 | -1.889235025 | 0.018521346 | hsa-miR-4282 |

| | | | | |
|---|---|---|---|---|
| hsa-miR-31-3p | 94.5 | -1.91747758 | 4.5737E-08 | hsa-miR-4650-3p |
| hsa-miR-4282 | 54.25 | -1.924402721 | 2.32159E-05 | hsa-miR-601 |
| hsa-miR-4650-3p | 22.25 | -1.935446277 | 0.007172993 | hsa-miR-222-5p |
| hsa-miR-193a-3p | 1137.5 | -2.011570739 | 1.86171E-24 | hsa-miR-424-3p |
| hsa-miR-452-5p | 42.25 | -2.013413179 | 0.000122497 | hsa-miR-582-5p |
| hsa-let-7d-3p | 26.75 | -2.054013503 | 0.0042833 | hsa-miR-452-5p |
| hsa-miR-125b-1-3p | 20.75 | -2.054604677 | 0.002696089 | hsa-miR-200b-3p |
| hsa-miR-331-3p | 319.25 | -2.152959373 | 2.70724E-15 | hsa-miR-9900 |
| hsa-miR-3691-3p | 12 | -2.232261316 | 0.016351721 | hsa-miR-542-3p |
| hsa-miR-601 | 29.25 | -2.269060734 | 8.62899E-05 | hsa-miR-513a-5p |
| hsa-miR-9900 | 149 | -2.301348367 | 7.65011E-16 | hsa-miR-26a-1-3p |
| hsa-miR-450a-5p | 26.25 | -2.306350225 | 0.001445636 | hsa-let-7e-3p |
| hsa-miR-582-5p | 142.75 | -2.30780123 | 8.42442E-13 | hsa-miR-6741-5p |
| hsa-miR-4435 | 10 | -2.343861186 | 0.023461004 | hsa-miR-125a-3p |
| hsa-miR-6850-3p | 8.75 | -2.375124892 | 0.04715106 | hsa-miR-4425 |
| hsa-miR-27b-5p | 343.75 | -2.41153794 | 1.75154E-26 | hsa-miR-10a-3p |
| hsa-miR-4425 | 35.75 | -2.503415087 | 1.13757E-05 | hsa-miR-365a-5p |
| hsa-miR-6728-5p | 12.75 | -2.639897628 | 0.004132465 | hsa-miR-27b-5p |
| hsa-miR-125a-3p | 43.75 | -2.648108543 | 1.07066E-07 | hsa-miR-193b-5p |
| hsa-miR-513a-5p | 19 | -2.68040177 | 0.000614751 | hsa-miR-4292 |
| hsa-miR-6734-5p | 11.5 | -2.685348892 | 0.009670706 | hsa-miR-5089-5p |
| hsa-miR-6741-5p | 59 | -2.737279465 | 7.51588E-08 | hsa-miR-6847-3p |
| hsa-miR-6784-5p | 6.5 | -2.79279604 | 0.042296983 | hsa-miR-3978 |
| hsa-miR-6125 | 9.25 | -2.806902202 | 0.042296983 | hsa-miR-141-5p |
| hsa-let-7e-3p | 17.5 | -2.942091611 | 0.000194996 | hsa-miR-134-3p |
| hsa-miR-6808-5p | 6.5 | -3.163659155 | 0.026876936 | hsa-miR-3691-3p |
| hsa-miR-192-3p | 10.5 | -3.314311545 | 0.004177148 | hsa-miR-6850-3p |
| hsa-miR-5089-5p | 7.25 | -3.520816927 | 0.012080857 | hsa-miR-6125 |
| hsa-miR-365a-5p | 12.25 | -3.550434332 | 0.000593525 | hsa-miR-647 |
| hsa-miR-29b-1-5p | 9 | -3.845865458 | 0.002999772 | hsa-miR-27a-5p |
| hsa-miR-4666b | 4.75 | -3.868622858 | 0.043115816 | hsa-miR-6774-3p |
| hsa-miR-875-3p | 9.25 | -3.964296648 | 0.005319228 | hsa-miR-6070 |
| hsa-miR-222-5p | 19.75 | -4.254883042 | 1.68835E-05 | hsa-miR-4718 |
| hsa-miR-4718 | 6.75 | -4.468495741 | 0.00853684 | hsa-miR-6850-5p |
| hsa-miR-33b-3p | 11.75 | -4.47949024 | 0.000249122 | hsa-miR-4303 |
| hsa-miR-342-5p | 7 | -4.631447527 | 0.007004278 | hsa-miR-33b-3p |
| hsa-miR-23a-5p | 29.75 | -4.987753929 | 8.62253E-09 | hsa-miR-23a-5p |
| hsa-miR-4303 | 5.25 | -5.471866968 | 0.016752189 | hsa-miR-6728-5p |
| hsa-miR-27a-5p | 59.75 | -6.209496648 | 2.47828E-16 | hsa-miR-342-5p |
| hsa-miR-5587-3p | 4.5 | -6.567815593 | 0.02490893 | hsa-miR-6737-5p |
| hsa-miR-6774-3p | 10.75 | -7.066744292 | 0.000173398 | hsa-miR-3940-5p |
| hsa-miR-6737-5p | 115.5 | -9.230006921 | 1.34179E-32 | |
| hsa-miR-6070 | 7.5 | -10.8179573 | 0.002154838 | |
| hsa-miR-6752-5p | 4.5 | -10.84212361 | 0.02606927 | |
| hsa-miR-3940-5p | 22 | -17.47132824 | 1.02368E-07 | |

**4p1 v C 36HPI** **Ad14p1 v C 48HP**

| Max group mean | Fold change | FDR p-value | Identifier | Max group mean | Fold change |
|---|---|---|---|---|---|
| 13 | 111.8774827 | 0.002585727 | hsa-miR-8077 | 18 | 111.6841957 |
| 10.25 | 88.43632392 | 0.004598096 | hsa-miR-518a-5p | 46.5 | 107.7627164 |
| 15.75 | 49.34835163 | 2.00696E-05 | hsa-miR-499b-5p | 15.5 | 96.24142476 |
| 6.5 | 20.55117935 | 0.001654959 | hsa-miR-4724-5p | 37.5 | 53.47115708 |
| 37.25 | 19.03923794 | 1.95098E-16 | hsa-miR-5008-5p | 7 | 42.15384151 |
| 21.75 | 16.22846911 | 2.55803E-10 | hsa-miR-4440 | 6.5 | 40.74714374 |
| 5 | 15.53063934 | 0.007538347 | hsa-miR-646 | 65.5 | 36.65519694 |
| 4.5 | 14.35031197 | 0.007474469 | hsa-miR-6802-3p | 5.75 | 36.12139902 |
| 12.75 | 13.73329374 | 1.33323E-06 | hsa-miR-6841-5p | 5.75 | 35.97531384 |
| 6.75 | 13.04103779 | 0.000754786 | hsa-miR-4286 | 845.25 | 35.53372348 |
| 6.5 | 12.56587769 | 0.000953076 | hsa-miR-10524-5p | 5 | 31.49837834 |
| 45.75 | 10.86842901 | 3.41648E-21 | hsa-miR-7850-5p | 13.25 | 30.76733686 |
| 2.75 | 8.922585641 | 0.046167916 | hsa-miR-520f-3p | 53.25 | 29.76077057 |
| 14.5 | 8.889362469 | 1.11504E-06 | hsa-miR-4462 | 20.75 | 29.51806441 |
| 11.75 | 8.821435048 | 3.64052E-06 | hsa-miR-1253 | 81.25 | 25.79937814 |
| 55.75 | 8.116550084 | 1.19419E-24 | hsa-miR-551a | 31.5 | 25.29185058 |
| 4 | 7.824024343 | 0.011328433 | hsa-miR-596 | 28.75 | 23.08476657 |
| 3.75 | 7.349689134 | 0.017783988 | hsa-miR-3168 | 86 | 22.96925854 |
| 5 | 6.999890517 | 0.00450588 | hsa-miR-3125 | 21.25 | 21.84425674 |
| 3.5 | 6.877112045 | 0.021292681 | hsa-miR-1261 | 45.25 | 21.81313517 |
| 119.5 | 6.725080448 | 1.03219E-35 | hsa-miR-124-3p | 9.25 | 21.55649267 |
| 15 | 6.451742865 | 7.24191E-07 | hsa-miR-6888-5p | 8.5 | 19.82472699 |
| 9.75 | 6.362316554 | 4.3912E-05 | hsa-miR-4668-3p | 8.5 | 19.66115233 |
| 10.75 | 6.19045729 | 1.74108E-05 | hsa-miR-155-5p | 13.5 | 19.33518306 |
| 3 | 5.928406949 | 0.040959118 | hsa-miR-3128 | 50 | 19.19268147 |
| 19.25 | 5.159997085 | 9.28819E-07 | hsa-miR-12120 | 74.75 | 18.96236181 |
| 8.5 | 5.120142801 | 0.000987569 | hsa-miR-412-3p | 7.5 | 17.52604534 |
| 15.5 | 5.071048763 | 1.63843E-05 | hsa-miR-6877-5p | 12 | 17.14935488 |
| 7.5 | 4.913894705 | 0.000892917 | hsa-miR-4759 | 11.5 | 16.44310312 |
| 4.25 | 4.668086234 | 0.017300492 | hsa-miR-4661-3p | 7 | 16.37260148 |
| 17.5 | 4.625861787 | 1.13909E-07 | hsa-miR-4696 | 20.25 | 16.28819986 |
| 6 | 4.608094317 | 0.008263164 | hsa-miR-4260 | 24 | 15.83488102 |
| 36.25 | 4.372678936 | 2.39138E-14 | hsa-miR-1296-3p | 11 | 15.72905154 |
| 36 | 4.240496437 | 5.27349E-14 | hsa-miR-12113 | 19.5 | 15.68886951 |
| 9 | 4.212135935 | 0.000474499 | hsa-miR-6762-5p | 10.5 | 15.025418 |
| 7.25 | 4.197856617 | 0.002888642 | hsa-miR-4754 | 10.5 | 15.02241803 |
| 9.25 | 3.953663663 | 0.00052624 | hsa-miR-4481 | 6.25 | 14.64497721 |
| 6.75 | 3.912413164 | 0.003324203 | hsa-miR-520g-5p | 32.75 | 14.01667933 |
| 6 | 3.485330154 | 0.009233216 | hsa-miR-200a-5p | 17 | 13.68833619 |
| 10.25 | 3.475270004 | 0.000696289 | hsa-miR-296-3p | 9.25 | 13.25346041 |
| 106.75 | 3.348619593 | 1.30459E-18 | hsa-miR-4299 | 27.5 | 11.80978755 |

| | | | | | |
|---|---|---|---|---|---|
| 7.5 | 3.250816517 | 0.032287576 | hsa-miR-3154 | 11.25 | 11.61222061 |
| 92.5 | 3.121417855 | 3.00681E-17 | hsa-miR-888-3p | 4.75 | 11.18952287 |
| 53.5 | 3.105158936 | 4.56337E-11 | hsa-miR-6763-3p | 4.75 | 11.16904398 |
| 5.25 | 3.059063776 | 0.027117276 | hsa-miR-492 | 35 | 11.14047159 |
| 7.75 | 3.056923156 | 0.006470966 | hsa-miR-4323 | 7.75 | 11.12905382 |
| 12.25 | 2.933614531 | 0.000696289 | hsa-miR-302f | 10.25 | 10.59014501 |
| 11 | 2.923704274 | 0.001132918 | hsa-miR-888-5p | 23.25 | 10.00230251 |
| 140.5 | 2.89971719 | 1.30459E-18 | hsa-miR-3136-3p | 6.75 | 9.930891972 |
| 170 | 2.733036183 | 1.53365E-17 | hsa-miR-6079 | 9.5 | 9.862232306 |
| 74.25 | 2.71596924 | 1.52716E-10 | hsa-miR-4431 | 28.25 | 9.843559433 |
| 8.5 | 2.700603482 | 0.01231764 | hsa-miR-33a-3p | 40.75 | 9.632645146 |
| 560 | 2.676352299 | 1.43227E-27 | hsa-miR-3928-3p | 22 | 9.441600955 |
| 6.25 | 2.673836554 | 0.047371504 | hsa-miR-6866-5p | 4 | 9.31408261 |
| 155.25 | 2.623030176 | 3.41648E-21 | hsa-miR-665 | 33.5 | 9.102687613 |
| 44.25 | 2.589374335 | 1.60124E-09 | hsa-miR-3911 | 8.75 | 9.058037092 |
| 54 | 2.573607234 | 6.63455E-11 | hsa-miR-5100 | 21 | 8.990252931 |
| 8.5 | 2.539725899 | 0.011552442 | hsa-miR-196a-1-3p | 23.25 | 8.952265622 |
| 11 | 2.514237904 | 0.003919977 | hsa-miR-759 | 6.25 | 8.908041039 |
| 88.25 | 2.329189585 | 7.34693E-11 | hsa-miR-299-5p | 6 | 8.650395565 |
| 42.75 | 2.306995148 | 1.04068E-07 | hsa-miR-3193 | 8.25 | 8.545706127 |
| 83 | 2.283668835 | 1.74114E-11 | hsa-miR-3610 | 8.25 | 8.530025701 |
| 86.25 | 2.267253555 | 4.10505E-09 | hsa-miR-4508 | 33 | 8.34209565 |
| 10.75 | 2.1568401 | 0.02403568 | hsa-miR-218-1-3p | 24 | 8.319683728 |
| 21.25 | 2.100358585 | 0.00121319 | hsa-miR-3649 | 25.75 | 8.200978161 |
| 13.5 | 2.091959841 | 0.025762042 | hsa-miR-518d-5p | 32 | 8.088832102 |
| 14.5 | 2.053409172 | 0.017466813 | hsa-miR-4762-3p | 25.25 | 8.048971522 |
| 217.5 | 2.037214073 | 9.07395E-08 | hsa-miR-1301-5p | 18.25 | 7.828110149 |
| 9.25 | 2.026407273 | 0.038662288 | hsa-miR-6085 | 7.5 | 7.779046166 |
| 1423 | 1.989703849 | 5.56636E-18 | hsa-miR-134-3p | 43 | 7.686738826 |
| 46.5 | 1.911034126 | 2.30556E-05 | hsa-miR-219b-3p | 7 | 7.370702961 |
| 952.75 | 1.875910315 | 1.90688E-11 | hsa-miR-4781-3p | 25 | 7.331129563 |
| 13 | 1.852018453 | 0.030840823 | hsa-miR-6753-5p | 7 | 7.267491471 |
| 22915.25 | 1.848823336 | 6.50375E-15 | hsa-miR-3678-3p | 5 | 7.234041669 |
| 22.25 | 1.832476379 | 0.004192479 | hsa-miR-7-5p | 169.75 | 7.228518455 |
| 32 | 1.769101123 | 0.001189616 | hsa-miR-6819-3p | 9 | 7.122236935 |
| 162 | 1.744275281 | 1.09013E-06 | hsa-miR-4279 | 8.75 | 7.087394898 |
| 60.75 | 1.704478868 | 0.000124549 | hsa-miR-483-5p | 29.5 | 7.084072218 |
| 19 | 1.677410648 | 0.020550286 | hsa-miR-6749-3p | 19.75 | 6.89241036 |
| 18.75 | 1.655530831 | 0.032011401 | hsa-miR-5692b | 8.5 | 6.887671275 |
| 32.25 | 1.650411921 | 0.004430824 | hsa-miR-2114-3p | 4.75 | 6.879378137 |
| 20364.25 | 1.63080805 | 2.81508E-10 | hsa-miR-4489 | 17.75 | 6.841994157 |
| 732.25 | 1.563205122 | 0.000268192 | hsa-miR-1305 | 6.5 | 6.756823956 |
| 44262.75 | 1.515466999 | 3.2449E-05 | hsa-miR-6081 | 6.5 | 6.756731953 |
| 52.75 | 1.494083398 | 0.00580898 | hsa-miR-3150b-3p | 26.5 | 6.708683688 |
| 389.75 | 1.469318677 | 0.001794921 | hsa-miR-330-3p | 27.25 | 6.560148379 |

| | | | | | |
|---|---|---|---|---|---|
| 967.5 | 1.461252282 | 1.33088E-05 | hsa-miR-203b-3p | 15.25 | 6.558139075 |
| 282.5 | 1.435465907 | 0.001197074 | hsa-miR-6873-5p | 4.5 | 6.536839707 |
| 34 | 1.427626504 | 0.047371504 | hsa-miR-3685 | 4.5 | 6.525929307 |
| 14831 | 1.425219674 | 9.55755E-05 | hsa-miR-626 | 4.5 | 6.525538319 |
| 434 | 1.423329372 | 0.00161221 | hsa-miR-8080 | 38 | 6.487622685 |
| 463 | 1.373109186 | 0.001654959 | hsa-miR-5089-5p | 52 | 6.474332156 |
| 4347.75 | 1.338434489 | 0.003266249 | hsa-miR-4654 | 38.75 | 6.337934271 |
| 205.25 | 1.327546878 | 0.034129631 | hsa-miR-556-3p | 6 | 6.245423284 |
| 1295.5 | -1.232797558 | 0.049433699 | hsa-miR-7851-3p | 4.25 | 6.171725793 |
| 241.25 | -1.322400202 | 0.01589475 | hsa-miR-6510-5p | 4.25 | 6.171109856 |
| 100.75 | -1.404118682 | 0.00523035 | hsa-miR-1288-3p | 4.25 | 6.165947535 |
| 2481.25 | -1.411287757 | 0.001654959 | hsa-miR-6165 | 12.5 | 6.102088863 |
| 6967.25 | -1.412679468 | 2.16846E-07 | hsa-miR-580-3p | 7.5 | 6.086242801 |
| 418.5 | -1.414548784 | 0.000989502 | hsa-miR-12136 | 40.75 | 6.080542085 |
| 235.25 | -1.423452571 | 0.002218114 | hsa-miR-1469 | 15.5 | 6.057119697 |
| 28627.5 | -1.4773713 | 1.11539E-06 | hsa-miR-1236-5p | 5.75 | 6.048206265 |
| 55 | -1.480778492 | 0.025232118 | hsa-miR-5692a | 7.25 | 5.828430614 |
| 44 | -1.48696331 | 0.022167143 | hsa-miR-4686 | 4 | 5.818825913 |
| 369 | -1.509661161 | 0.008456732 | hsa-miR-12114 | 8.75 | 5.818695496 |
| 29.75 | -1.546263768 | 0.039254867 | hsa-miR-6799-5p | 4 | 5.817906861 |
| 47.25 | -1.630756096 | 0.006467387 | hsa-miR-1825 | 4 | 5.817260595 |
| 33.75 | -1.646680112 | 0.006826131 | hsa-miR-4433b-3p | 5.5 | 5.733672176 |
| 270 | -1.648278412 | 3.47872E-05 | hsa-miR-4773 | 7 | 5.686904028 |
| 63.75 | -1.656579006 | 0.000481085 | hsa-miR-3620-3p | 25 | 5.579889932 |
| 111.75 | -1.668375227 | 0.000956423 | hsa-miR-3156-3p | 61.5 | 5.577729188 |
| 1045.75 | -1.676340662 | 6.48067E-09 | hsa-miR-8083 | 6.75 | 5.566932272 |
| 111 | -1.681754271 | 0.000127021 | hsa-miR-6757-5p | 12.75 | 5.500150368 |
| 729.75 | -1.693741237 | 1.87066E-07 | hsa-miR-4443 | 5.25 | 5.478520849 |
| 209.25 | -1.713454017 | 8.41641E-05 | hsa-miR-6768-5p | 3.75 | 5.46391814 |
| 51.25 | -1.729441929 | 0.000587988 | hsa-miR-4424 | 9.5 | 5.356015159 |
| 20.75 | -1.736445073 | 0.036329211 | hsa-miR-6833-5p | 6.5 | 5.287142231 |
| 61.25 | -1.777388095 | 0.001862657 | hsa-miR-5699-5p | 9.25 | 5.216160889 |
| 17.5 | -1.780666903 | 0.039155471 | hsa-miR-7113-3p | 31.75 | 5.185189305 |
| 5732 | -1.876664336 | 7.11474E-10 | hsa-miR-6783-5p | 20.25 | 5.1434762 |
| 1803.5 | -1.921424806 | 3.09029E-13 | hsa-miR-1197 | 9 | 5.080017862 |
| 238.75 | -1.935295017 | 1.22068E-06 | hsa-miR-642a-3p | 11.75 | 5.077332027 |
| 26.25 | -1.943912808 | 0.00196597 | hsa-miR-4691-3p | 4.75 | 4.966981744 |
| 26.75 | -1.958671096 | 0.004598096 | hsa-miR-12128 | 4.75 | 4.966556285 |
| 94.5 | -1.961253108 | 7.24191E-07 | hsa-miR-6776-3p | 6 | 4.88662012 |
| 88.75 | -1.961387124 | 5.83768E-06 | hsa-miR-1276 | 20.5 | 4.864994153 |
| 25.75 | -1.987923471 | 0.005420812 | hsa-miR-4433a-3p | 9.75 | 4.841176597 |
| 319.25 | -1.990733614 | 4.02046E-09 | hsa-miR-5584-5p | 9.75 | 4.771392048 |
| 1137.5 | -2.060214476 | 6.70961E-13 | hsa-miR-4776-5p | 10.5 | 4.538339046 |
| 84.75 | -2.060539568 | 3.09382E-08 | hsa-miR-1268a | 8 | 4.492164562 |
| 54.25 | -2.063869836 | 8.32824E-07 | hsa-miR-4699-5p | 4.25 | 4.455456163 |

| | | | | | |
|---|---|---|---|---|---|
| 22.25 | -2.067320622 | 0.002218114 | hsa-miR-432-3p | 7.75 | 4.378961991 |
| 29.25 | -2.119862784 | 0.000268192 | hsa-miR-1245b-3p | 20.75 | 4.358583489 |
| 19.75 | -2.141971779 | 0.011297456 | hsa-miR-4254 | 6.5 | 4.341466521 |
| 32 | -2.175742911 | 7.78463E-05 | hsa-miR-6736-3p | 5.25 | 4.331319732 |
| 142.75 | -2.313162638 | 6.86679E-10 | hsa-miR-5691 | 19.25 | 4.293469263 |
| 42.25 | -2.459747621 | 7.95975E-07 | hsa-miR-6784-5p | 30.5 | 4.231984609 |
| 13 | -2.464931891 | 0.007659698 | hsa-miR-1199-5p | 6.25 | 4.177267222 |
| 149 | -2.525590898 | 1.3192E-10 | hsa-miR-3917 | 8.5 | 4.167112685 |
| 23.5 | -2.54142776 | 0.000558906 | hsa-miR-6793-5p | 5 | 4.086362703 |
| 19 | -2.670717631 | 0.000979198 | hsa-miR-4475 | 7 | 3.959077665 |
| 9.25 | -2.677589015 | 0.025762042 | hsa-miR-10392-3p | 8 | 3.936759061 |
| 17.5 | -2.692448042 | 0.000989502 | hsa-miR-668-5p | 21.5 | 3.858639412 |
| 59 | -2.70881196 | 4.13034E-10 | hsa-miR-3169 | 5.75 | 3.856253618 |
| 43.75 | -2.742556425 | 1.01451E-08 | hsa-miR-6823-5p | 5.5 | 3.684477248 |
| 35.75 | -2.770577934 | 3.16572E-06 | hsa-miR-6816-3p | 8 | 3.468054236 |
| 13 | -2.972937493 | 0.002181299 | hsa-miR-4648 | 8.25 | 3.460008201 |
| 12.25 | -3.011544874 | 0.002797744 | hsa-miR-5683 | 16.75 | 3.333143834 |
| 343.75 | -3.047230814 | 3.59797E-22 | hsa-miR-1266-5p | 11.25 | 3.332232724 |
| 25.75 | -3.08734618 | 5.49591E-06 | hsa-miR-495-3p | 9.25 | 3.247402086 |
| 14 | -3.211639857 | 0.000989502 | hsa-miR-4511 | 16 | 3.210796615 |
| 7.25 | -3.231848552 | 0.020729573 | hsa-miR-8087 | 70.5 | 3.09193126 |
| 9.5 | -3.329544023 | 0.006412804 | hsa-miR-4318 | 13 | 3.090419521 |
| 20.75 | -3.347415714 | 0.000525248 | hsa-miR-1238-5p | 8.75 | 3.073287751 |
| 5.75 | -3.513796156 | 0.036329211 | hsa-miR-324-5p | 43 | 3.064253038 |
| 5 | -3.747668127 | 0.047742975 | hsa-miR-3613-5p | 28.5 | 3.041487669 |
| 12 | -3.79361216 | 0.000953076 | hsa-miR-889-5p | 16.75 | 3.010460361 |
| 8.75 | -4.052866557 | 0.010071719 | hsa-miR-4522 | 7.75 | 2.986774444 |
| 9.25 | -4.561950425 | 0.006467387 | hsa-miR-4436a | 10 | 2.950908053 |
| 5 | -4.751581821 | 0.047429664 | hsa-miR-548a-5p | 27.25 | 2.888637295 |
| 59.75 | -4.995871987 | 4.1175E-15 | hsa-miR-1908-3p | 15.25 | 2.881862448 |
| 10.75 | -5.530125422 | 0.000989502 | hsa-miR-548u | 38.75 | 2.880817446 |
| 7.5 | -5.568879922 | 0.004898444 | hsa-miR-4714-5p | 25.75 | 2.851220032 |
| 6.75 | -6.455274597 | 0.005988459 | hsa-miR-517-5p | 12.5 | 2.793677255 |
| 5.25 | -7.068128445 | 0.017466813 | hsa-miR-3960 | 9 | 2.658646097 |
| 5.25 | -7.068969836 | 0.017300492 | hsa-miR-548n | 9 | 2.658317412 |
| 11.75 | -7.092654744 | 0.000169609 | hsa-miR-132-3p | 122 | 2.604698753 |
| 29.75 | -7.264672779 | 3.52892E-10 | hsa-miR-2392 | 177 | 2.436560488 |
| 12.75 | -7.685491185 | 9.55755E-05 | hsa-miR-2682-5p | 15.25 | 2.296177555 |
| 7 | -9.363727766 | 0.004482853 | hsa-miR-5587-3p | 10.5 | 2.09701497 |
| 115.5 | -10.98380767 | 3.65399E-35 | hsa-miR-345-3p | 10.25 | 2.048232263 |
| 22 | -13.27430672 | 4.08947E-07 | hsa-miR-10400-5p | 20 | 1.966032711 |
| | | | hsa-miR-629-5p | 22.75 | 1.965147176 |
| | | | hsa-miR-410-3p | 74.75 | 1.904860223 |
| | | | hsa-miR-106b-3p | 120.25 | 1.899130205 |
| | | | hsa-miR-877-5p | 62.75 | 1.876190509 |

| | | |
|---|---|---|
| hsa-miR-9-5p | 69 | 1.649482716 |
| hsa-miR-8066 | 59.25 | 1.64002017 |
| hsa-miR-148a-3p | 862.5 | 1.310381842 |
| hsa-miR-182-5p | 12365.5 | -1.327007068 |
| hsa-miR-130a-3p | 726.25 | -1.328818165 |
| hsa-miR-29a-3p | 1045.75 | -1.346100479 |
| hsa-miR-192-5p | 34179.25 | -1.353066897 |
| hsa-miR-3155a | 369 | -1.373899941 |
| hsa-miR-769-5p | 362.75 | -1.379516883 |
| hsa-miR-589-5p | 140.75 | -1.389220535 |
| hsa-miR-34a-5p | 335.75 | -1.395513524 |
| hsa-miR-103a-3p | 1881 | -1.440104023 |
| hsa-miR-300 | 75 | -1.454629284 |
| hsa-miR-128-3p | 327.25 | -1.470034005 |
| hsa-miR-196a-5p | 128 | -1.517905524 |
| hsa-miR-151a-5p | 2481.25 | -1.544890875 |
| hsa-miR-331-3p | 319.25 | -1.548807819 |
| hsa-let-7a-5p | 34460.75 | -1.574642609 |
| hsa-miR-191-5p | 4219.5 | -1.576126093 |
| hsa-miR-3184-5p | 566.5 | -1.591665565 |
| hsa-miR-561-5p | 78 | -1.616011874 |
| hsa-miR-1297 | 6967.25 | -1.629281297 |
| hsa-miR-31-3p | 94.5 | -1.630793527 |
| hsa-miR-193a-3p | 1137.5 | -1.633705436 |
| hsa-miR-190a-5p | 28.25 | -1.660561423 |
| hsa-miR-99a-3p | 96 | -1.685729808 |
| hsa-miR-221-3p | 2559.5 | -1.689672214 |
| hsa-miR-365a-3p | 238.75 | -1.696477917 |
| hsa-miR-941 | 184.25 | -1.707178068 |
| hsa-miR-28-3p | 3228.25 | -1.724482471 |
| hsa-miR-452-5p | 42.25 | -1.725566946 |
| hsa-miR-320a-3p | 597.25 | -1.747892258 |
| hsa-miR-12135 | 409 | -1.767612403 |
| hsa-miR-28-5p | 241.25 | -1.779384408 |
| hsa-miR-944 | 46 | -1.791958058 |
| hsa-miR-143-3p | 125.5 | -1.805323236 |
| hsa-miR-122b-3p | 9213 | -1.806804551 |
| hsa-miR-148a-5p | 26.5 | -1.826875814 |
| hsa-miR-409-3p | 29.5 | -1.858406481 |
| hsa-miR-455-3p | 22.5 | -1.877809806 |
| hsa-miR-125a-5p | 5732 | -1.879402414 |
| hsa-miR-574-5p | 50.25 | -1.881567781 |
| hsa-miR-1296-5p | 18.5 | -1.912584921 |
| hsa-miR-210-3p | 270 | -1.933925824 |
| hsa-miR-3927-3p | 40.5 | -1.954695029 |

| | | |
|---|---|---|
| hsa-miR-21-5p | 109325 | -1.960311034 |
| hsa-miR-513a-5p | 19 | -1.962171689 |
| hsa-miR-128-1-5p | 20.25 | -1.996662345 |
| hsa-miR-328-3p | 9.25 | -2.001596785 |
| hsa-miR-3667-3p | 77.75 | -2.025869749 |
| hsa-miR-320e | 38.75 | -2.027622307 |
| hsa-miR-193b-5p | 25.75 | -2.031607287 |
| hsa-miR-1180-3p | 73.5 | -2.031894006 |
| hsa-miR-31-5p | 4277.25 | -2.039888477 |
| hsa-miR-22-3p | 28627.5 | -2.107871049 |
| hsa-miR-30a-5p | 25622.5 | -2.122476471 |
| hsa-miR-3145-3p | 14.75 | -2.12585818 |
| hsa-miR-30a-3p | 450 | -2.179418022 |
| hsa-miR-10396a-5p | 105.25 | -2.180056037 |
| hsa-miR-423-3p | 740.5 | -2.200908026 |
| hsa-miR-4536-5p | 21 | -2.220098319 |
| hsa-miR-374a-3p | 214 | -2.225178794 |
| hsa-miR-224-5p | 111 | -2.251200314 |
| hsa-miR-500a-3p | 33.75 | -2.253021345 |
| hsa-miR-5579-3p | 288 | -2.330844707 |
| hsa-miR-671-5p | 9.75 | -2.336559084 |
| hsa-miR-134-5p | 33.75 | -2.360783525 |
| hsa-let-7f-2-3p | 12.25 | -2.405040757 |
| hsa-miR-10a-3p | 13 | -2.441494106 |
| hsa-miR-6734-5p | 11.5 | -2.480284689 |
| hsa-miR-221-5p | 20 | -2.504449184 |
| hsa-miR-10396b-5p | 16.25 | -2.600996806 |
| hsa-miR-26a-1-3p | 9.25 | -2.646617195 |
| hsa-let-7a-3p | 84.75 | -2.649634701 |
| hsa-miR-6499-3p | 6.25 | -2.67009498 |
| hsa-miR-6847-3p | 9.5 | -2.72085572 |
| hsa-miR-6774-3p | 10.75 | -2.720975419 |
| hsa-miR-141-5p | 5.75 | -2.721440732 |
| hsa-miR-744-3p | 5.75 | -2.721442084 |
| hsa-miR-6850-3p | 8.75 | -2.724818531 |
| hsa-miR-3122 | 12.75 | -2.745708652 |
| hsa-miR-887-5p | 8.5 | -2.802121404 |
| hsa-miR-374b-3p | 10 | -2.861438907 |
| hsa-miR-3909 | 6.25 | -2.952035382 |
| hsa-miR-6741-5p | 59 | -2.985050735 |
| hsa-miR-582-3p | 573.25 | -2.985479123 |
| hsa-miR-605-5p | 7.5 | -2.994762595 |
| hsa-miR-3978 | 20.75 | -3.000026599 |
| hsa-miR-3165 | 25.75 | -3.011498759 |
| hsa-miR-424-3p | 32 | -3.013263805 |

| | | |
|---|---|---|
| hsa-miR-1257 | 80.75 | -3.076535996 |
| hsa-miR-6752-5p | 4.5 | -3.145294913 |
| hsa-miR-181a-2-3p | 418.5 | -3.226645718 |
| hsa-miR-4446-3p | 55 | -3.242970085 |
| hsa-miR-1587 | 4.75 | -3.315167681 |
| hsa-miR-6844 | 5.75 | -3.453557472 |
| hsa-miR-4280 | 5.75 | -3.454916646 |
| hsa-miR-4502 | 29.75 | -3.47655896 |
| hsa-miR-3662 | 43.25 | -3.492804486 |
| hsa-miR-1273h-5p | 8.25 | -3.501938522 |
| hsa-miR-1272 | 46.75 | -3.593042878 |
| hsa-miR-210-5p | 13.75 | -3.679272052 |
| hsa-miR-125b-1-3p | 20.75 | -3.722756519 |
| hsa-let-7e-3p | 17.5 | -3.758369957 |
| hsa-miR-4716-3p | 8.25 | -3.890023959 |
| hsa-miR-30c-1-3p | 47.25 | -4.067681857 |
| hsa-miR-181a-3p | 209.25 | -4.270415582 |
| hsa-miR-6850-5p | 5.25 | -4.324649297 |
| hsa-miR-4303 | 5.25 | -4.32515007 |
| hsa-miR-671-3p | 29 | -4.45933078 |
| hsa-miR-194-3p | 12.75 | -4.513186171 |
| hsa-miR-584-3p | 3.5 | -4.626785585 |
| hsa-miR-4697-3p | 3.5 | -4.627120172 |
| hsa-miR-34a-3p | 4.75 | -4.810853417 |
| hsa-miR-6747-5p | 6 | -4.925532593 |
| hsa-miR-6881-5p | 7.5 | -5.175835324 |
| hsa-miR-3940-5p | 22 | -5.5118804 |
| hsa-miR-365a-5p | 12.25 | -5.721322478 |
| hsa-miR-25-5p | 13.75 | -5.794629661 |
| hsa-miR-23a-5p | 29.75 | -5.800309829 |
| hsa-miR-9900 | 149 | -5.867455772 |
| hsa-miR-125a-3p | 43.75 | -6.07404552 |
| hsa-miR-7974 | 100.75 | -6.836231045 |
| hsa-miR-6758-3p | 3.75 | -6.979265175 |
| hsa-miR-100-3p | 21 | -8.052426202 |
| hsa-miR-6125 | 9.25 | -8.760242985 |
| hsa-miR-29b-1-5p | 9 | -8.984354109 |
| hsa-miR-27b-5p | 343.75 | -9.17095664 |
| hsa-miR-214-5p | 88.75 | -9.201974603 |
| hsa-miR-4659a-5p | 44 | -9.400441614 |
| hsa-miR-601 | 29.25 | -9.522141634 |
| hsa-miR-7158-5p | 5.25 | -9.661980925 |
| hsa-miR-4425 | 35.75 | -10.83146896 |
| hsa-miR-4718 | 6.75 | -12.3458122 |
| hsa-miR-6737-5p | 115.5 | -12.61499195 |

| | | |
|---|---|---|
| hsa-miR-33b-3p | 11.75 | -15.09187341 |
| hsa-miR-6728-5p | 12.75 | -23.08080419 |
| hsa-miR-222-5p | 19.75 | -24.92786296 |
| hsa-miR-27a-5p | 59.75 | -30.74480434 |

#### PI Ad14 v Ad14p1 6HPI Ad14

| FDR p-value | Identifier | Max group mean | Fold change | FDR p-value | Identifier |
|---|---|---|---|---|---|
| 0.002140831 | hsa-miR-3664-5p | 5 | -5.964865829 | 0.033493107 | hsa-miR-12133 |
| 1.27972E-07 | hsa-miR-649 | 9.5 | -2.653126764 | 0.01990159 | hsa-miR-100-5p |
| 0.00281383 | hsa-miR-33b-3p | 9 | -2.651364662 | 0.020683305 | hsa-miR-183-5p |
| 2.15497E-08 | hsa-miR-8066 | 34.75 | -1.630505591 | 0.020220309 | hsa-miR-128-3p |
| 0.027731349 | hsa-miR-5187-3p | 28.25 | -1.619419298 | 0.033493107 | hsa-miR-214-5p |
| 0.020512503 | hsa-miR-183-5p | 394.75 | -1.48873377 | 1.88204E-06 | hsa-miR-134-5p |
| 6.30103E-15 | hsa-miR-100-5p | 3152.5 | -1.430918301 | 8.26962E-07 | hsa-miR-4714-5p |
| 0.026631636 | hsa-miR-127-3p | 636 | -1.261357008 | 0.018712706 | hsa-miR-2355-5p |
| 0.029711624 | hsa-miR-192-5p | 61738.25 | -1.230510406 | 0.018712706 | |
| 3.574E-114 | hsa-miR-10a-5p | 21353.25 | -1.224502342 | 0.020220309 | |
| 0.034386439 | hsa-miR-31-5p | 6341.5 | 1.257203009 | 0.009012972 | |
| 0.000174149 | hsa-miR-28-5p | 248.75 | 1.324011722 | 0.020683305 | |
| 7.37042E-14 | hsa-miR-221-3p | 3722.75 | 1.336694513 | 0.000194933 | |
| 2.25581E-06 | hsa-miR-28-3p | 5317.75 | 1.354301694 | 0.015353497 | |
| 3.04849E-19 | hsa-miR-375-3p | 200.5 | 1.376285439 | 0.01990159 | |
| 2.66143E-09 | hsa-miR-423-3p | 1086.25 | 1.482479239 | 2.62102E-05 | |
| 3.82103E-09 | hsa-miR-3184-5p | 1213.25 | 1.628278972 | 4.37638E-07 | |
| 1.4968E-05 | hsa-miR-1827 | 57.75 | 1.636968991 | 0.018712706 | |
| 4.72312E-07 | hsa-miR-214-5p | 102.25 | 1.864449176 | 0.00022777 | |
| 2.60905E-12 | hsa-miR-128-3p | 458 | 1.899419705 | 5.47723E-15 | |
| 0.001361002 | hsa-miR-424-3p | 32.25 | 1.940666901 | 0.011407212 | |
| 0.002271266 | hsa-miR-3181 | 38.5 | 2.003010296 | 0.009012972 | |
| 0.002613402 | hsa-miR-4435 | 9.25 | 3.287985095 | 0.039013719 | |
| 7.43855E-05 | | | | | |
| 5.74043E-14 | | | | | |
| 9.54856E-19 | | | | | |
| 0.002889 | | | | | |
| 0.000147254 | | | | | |
| 0.000182953 | | | | | |
| 0.004913994 | | | | | |
| 2.12932E-07 | | | | | |
| 5.00138E-08 | | | | | |
| 0.000262478 | | | | | |
| 8.56487E-07 | | | | | |
| 0.000415272 | | | | | |
| 0.000397101 | | | | | |
| 0.007149017 | | | | | |
| 1.44669E-10 | | | | | |
| 1.7655E-06 | | | | | |
| 0.000861594 | | | | | |
| 9.45367E-10 | | | | | |

| |
|---|
| 0.000198995 |
| 0.016695345 |
| 0.022072437 |
| 1.81884E-11 |
| 0.001712197 |
| 0.000175463 |
| 3.10808E-08 |
| 0.006229992 |
| 0.001078427 |
| 3.74333E-09 |
| 3.49326E-11 |
| 4.37408E-07 |
| 0.042060548 |
| 1.27711E-09 |
| 0.000928848 |
| 3.92702E-07 |
| 6.63196E-08 |
| 0.008250439 |
| 0.006669719 |
| 0.001712197 |
| 0.002506338 |
| 2.96489E-11 |
| 1.54826E-07 |
| 2.37982E-08 |
| 5.29974E-11 |
| 1.96307E-07 |
| 8.66107E-06 |
| 0.002525334 |
| 6.41377E-13 |
| 0.006419167 |
| 5.79891E-08 |
| 0.003733281 |
| 0.017789269 |
| 4.87853E-10 |
| 0.042260725 |
| 0.000693757 |
| 7.48556E-09 |
| 2.18294E-06 |
| 0.002366702 |
| 0.016496021 |
| 8.6897E-06 |
| 0.005248282 |
| 0.005371948 |
| 2.6017E-08 |
| 2.53441E-07 |

| |
|---|
| 2.59105E-05 |
| 0.030274785 |
| 0.022270148 |
| 0.022599648 |
| 4.39865E-11 |
| 6.30103E-15 |
| 4.15744E-11 |
| 0.006700858 |
| 0.030425483 |
| 0.035782302 |
| 0.042940205 |
| 9.30943E-05 |
| 0.003929794 |
| 1.01291E-10 |
| 0.000163772 |
| 0.021446904 |
| 0.010949734 |
| 0.035789896 |
| 0.002045368 |
| 0.035789896 |
| 0.033989199 |
| 0.011735418 |
| 0.004495674 |
| 2.03873E-06 |
| 4.82962E-12 |
| 0.012326588 |
| 5.89543E-05 |
| 0.019516736 |
| 0.042720252 |
| 0.001643786 |
| 0.006586379 |
| 0.000973636 |
| 9.45367E-10 |
| 2.94962E-05 |
| 0.001719905 |
| 0.000805586 |
| 0.022599648 |
| 0.03200933 |
| 0.016117814 |
| 2.10967E-06 |
| 0.00455405 |
| 0.002366702 |
| 0.000722317 |
| 0.007463098 |
| 0.047300498 |

| |
|---|
| 0.006972933 |
| 7.43163E-06 |
| 0.012510678 |
| 0.036354522 |
| 3.27811E-05 |
| 5.95651E-08 |
| 0.012357715 |
| 0.004075768 |
| 0.042720252 |
| 0.033950956 |
| 0.034386439 |
| 8.75001E-06 |
| 0.047300498 |
| 0.029931395 |
| 0.018791776 |
| 0.036127699 |
| 0.000244383 |
| 0.005239892 |
| 0.006239066 |
| 0.002358064 |
| 1.8017E-06 |
| 0.001296398 |
| 0.022072437 |
| 0.001629931 |
| 6.37672E-06 |
| 0.00281383 |
| 0.047211757 |
| 0.004980501 |
| 0.000321012 |
| 0.002140831 |
| 5.60532E-05 |
| 0.000229079 |
| 0.008697675 |
| 0.016668714 |
| 0.02057595 |
| 4.02013E-09 |
| 1.11483E-10 |
| 0.009388309 |
| 0.035789896 |
| 0.044286186 |
| 0.013677756 |
| 0.012357715 |
| 0.001842895 |
| 0.000127624 |
| 0.007463098 |

| |
|---|
| 0.009012233 |
| 0.045549159 |
| 0.022755879 |
| 0.027267948 |
| 0.012354374 |
| 0.00765963 |
| 0.013499172 |
| 0.007536917 |
| 0.01385976 |
| 0.030425483 |
| 0.009736445 |
| 0.001376012 |
| 0.047879888 |
| 0.004254398 |
| 0.013058147 |
| 0.00011828 |
| 0.004320653 |
| 3.09731E-05 |
| 4.58109E-05 |
| 0.000776606 |
| 0.007463098 |
| 6.05261E-06 |
| 0.014905808 |
| 2.62627E-06 |
| 0.029949595 |
| 0.003302219 |
| 1.19203E-06 |
| 4.83346E-05 |
| 6.13024E-05 |
| 1.99767E-06 |
| 0.030477454 |
| 2.25581E-06 |
| 1.31308E-05 |
| 7.93463E-05 |
| 0.015333471 |
| 0.001572763 |
| 5.73348E-08 |
| 0.011949783 |
| 0.006306097 |
| 0.022123613 |
| 6.12555E-10 |
| 0.004913994 |
| 0.025623894 |
| 1.09349E-06 |
| 0.004948749 |

| |
|---|
| 1.00001E-11 |
| 0.016496021 |
| 0.011575326 |
| 0.045549159 |
| 0.00267913 |
| 0.002610845 |
| 0.00281383 |
| 0.001039747 |
| 4.15744E-11 |
| 4.74654E-13 |
| 2.7665E-14 |
| 0.04594627 |
| 3.31484E-10 |
| 8.60579E-07 |
| 5.00138E-08 |
| 0.005464114 |
| 4.12746E-08 |
| 1.63871E-06 |
| 4.78042E-05 |
| 2.96489E-11 |
| 0.044246754 |
| 3.2836E-05 |
| 0.007526684 |
| 0.009115041 |
| 0.017968027 |
| 0.00455405 |
| 0.005239892 |
| 0.02057595 |
| 1.63871E-06 |
| 0.04704758 |
| 0.013710034 |
| 0.008706378 |
| 0.04594627 |
| 0.038788936 |
| 0.035789896 |
| 0.002409609 |
| 0.021398053 |
| 0.00302575 |
| 0.031699447 |
| 5.75088E-07 |
| 9.74034E-20 |
| 0.046073395 |
| 0.001532472 |
| 3.68715E-05 |
| 2.19972E-07 |

| |
|---|
| 1.79034E-07 |
| 0.045764551 |
| 7.42508E-16 |
| 1.79034E-07 |
| 0.034791215 |
| 0.023088116 |
| 0.02534038 |
| 1.56016E-06 |
| 8.45399E-08 |
| 0.00624954 |
| 5.09602E-10 |
| 0.000174149 |
| 5.60043E-06 |
| 3.68569E-05 |
| 0.010085611 |
| 5.39108E-08 |
| 3.70828E-25 |
| 0.016378425 |
| 0.011575326 |
| 1.03031E-08 |
| 2.86936E-05 |
| 0.04078839 |
| 0.04078839 |
| 0.013570908 |
| 0.006402449 |
| 0.00281383 |
| 4.86478E-06 |
| 1.37658E-05 |
| 8.0377E-06 |
| 1.94127E-09 |
| 1.11882E-25 |
| 1.06721E-13 |
| 2.13561E-16 |
| 0.024980803 |
| 2.46572E-08 |
| 0.001186036 |
| 0.000246112 |
| 2.69738E-42 |
| 4.64993E-15 |
| 1.91652E-15 |
| 5.29974E-11 |
| 0.004948749 |
| 1.04816E-12 |
| 0.001629931 |
| 1.71166E-23 |

| |
|---|
| 2.29882E-05 |
| 4.58109E-05 |
| 4.22271E-07 |
| 1.49202E-17 |

### v Ad14p1 12HPI

| Max group mean | Fold change | FDR p-value |
|---|---|---|
| 41.75 | -2.757765809 | 2.2951E-05 |
| 3794.75 | -1.720162935 | 0.000309963 |
| 429.75 | -1.557962488 | 0.000309963 |
| 389.25 | 1.479834716 | 0.009429037 |
| 114.25 | 1.497393678 | 0.045811723 |
| 42.75 | 1.865288605 | 0.00995941 |
| 27.75 | 1.905740845 | 0.010901471 |
| 13.5 | 2.510544794 | 0.026976219 |

### Ad14 v Ad14p1 24HPI

| Identifier | Max group mean | Fold change | FDR p-value |
|---|---|---|---|
| hsa-miR-668-5p | 58 | -7.801974142 | 3.17337E-15 |
| hsa-miR-6741-5p | 19 | -2.784928446 | 0.003802427 |
| hsa-miR-7158-3p | 36.75 | -2.387925819 | 3.45142E-05 |
| hsa-miR-4714-5p | 85.75 | -2.29315216 | 1.15776E-09 |
| hsa-miR-9900 | 58 | -2.061467729 | 1.06965E-05 |
| hsa-miR-12133 | 50.75 | -1.705700608 | 0.009700209 |
| hsa-miR-7974 | 85 | -1.602602726 | 0.000904 |
| hsa-miR-183-5p | 406.25 | -1.33086895 | 0.035649421 |
| hsa-miR-3155a | 210.25 | -1.276941705 | 0.042450099 |
| hsa-miR-1297 | 4804 | 1.22072751 | 0.022948938 |
| hsa-miR-128-3p | 308.25 | 1.250976584 | 0.034935898 |
| hsa-miR-132-3p | 95 | 1.427391348 | 0.034935898 |

### Ad14 v Ad14p1 36HPI

| Identifier | Max group mean | Fold change | FDR p-value |
|---|---|---|---|
| hsa-miR-7158-3p | 32 | -4.505574034 | 8.93838E-08 |
| hsa-miR-589-3p | 21.25 | -3.482885422 | 0.00037624 |
| hsa-miR-487a-5p | 15 | -3.385319589 | 0.005580633 |
| hsa-miR-4714-5p | 55.75 | -3.262114064 | 8.9574E-10 |
| hsa-miR-6741-5p | 17.75 | -3.083226431 | 0.002747215 |
| hsa-miR-1827 | 56.75 | -2.883077547 | 8.93838E-08 |
| hsa-miR-548u | 14.75 | -2.424916241 | 0.043280537 |
| hsa-miR-140-5p | 28 | -2.041263118 | 0.010262679 |
| hsa-miR-668-5p | 45.75 | -1.713866766 | 0.012409825 |
| hsa-miR-3156-3p | 36.25 | -1.696156657 | 0.025539276 |
| hsa-miR-4286 | 144.25 | 1.580506861 | 0.002351945 |
| hsa-miR-3128 | 51.25 | 1.827164438 | 0.000151355 |
| hsa-miR-1276 | 16.5 | 2.568626294 | 0.005580633 |
| hsa-miR-649 | 8.75 | 2.857521473 | 0.019607293 |
| hsa-miR-518a-5p | 34.75 | 2.937399531 | 4.81607E-06 |

# Ad14 v Ad1

| Identifier | Max group mean |
|---|---|
| hsa-miR-499b-5p | 15.5 |
| hsa-miR-4762-3p | 25.25 |
| hsa-miR-3168 | 86 |
| hsa-miR-134-3p | 43 |
| hsa-miR-4696 | 20.25 |
| hsa-miR-4754 | 10.5 |
| hsa-miR-5089-5p | 52 |
| hsa-miR-6877-5p | 12 |
| hsa-miR-3154 | 11.25 |
| hsa-miR-1268a | 8 |
| hsa-miR-665 | 33.5 |
| hsa-miR-412-3p | 7.5 |
| hsa-miR-6736-3p | 5.25 |
| hsa-miR-6079 | 9.5 |
| hsa-miR-4654 | 38.75 |
| hsa-miR-5100 | 21 |
| hsa-miR-4424 | 9.5 |
| hsa-miR-6085 | 7.5 |
| hsa-miR-6762-5p | 10.5 |
| hsa-miR-219b-3p | 7 |
| hsa-miR-5584-5p | 9.75 |
| hsa-miR-6833-5p | 6.5 |
| hsa-miR-4462 | 20.75 |
| hsa-miR-330-3p | 27.25 |
| hsa-miR-1238-5p | 8.75 |
| hsa-miR-218-1-3p | 24 |
| hsa-miR-6753-5p | 7 |
| hsa-miR-4433b-3p | 5.5 |
| hsa-miR-6892-5p | 9 |
| hsa-miR-6165 | 12.5 |
| hsa-miR-6783-5p | 20.25 |
| hsa-miR-4318 | 13 |
| hsa-miR-1908-3p | 15.25 |
| hsa-miR-4759 | 11.5 |
| hsa-miR-551a | 31.5 |
| hsa-miR-1296-3p | 11 |
| hsa-miR-3928-3p | 22 |
| hsa-miR-4440 | 6.5 |
| hsa-miR-3917 | 8.5 |
| hsa-miR-642a-3p | 11.75 |
| hsa-miR-4650-3p | 16.75 |

| | |
|---|---|
| hsa-miR-3911 | 8.75 |
| hsa-miR-4260 | 24 |
| hsa-miR-5587-3p | 10.5 |
| hsa-miR-8077 | 18 |
| hsa-miR-6757-5p | 12.75 |
| hsa-miR-345-3p | 10.25 |
| hsa-miR-520g-5p | 32.75 |
| hsa-miR-4431 | 28.25 |
| hsa-miR-196a-1-3p | 23.25 |
| hsa-miR-8080 | 38 |
| hsa-miR-4724-5p | 37.5 |
| hsa-miR-12120 | 74.75 |
| hsa-miR-12136 | 40.75 |
| hsa-miR-4436a | 10 |
| hsa-miR-518d-5p | 32 |
| hsa-miR-1253 | 81.25 |
| hsa-miR-646 | 65.5 |
| hsa-miR-7850-5p | 13.25 |
| hsa-miR-12113 | 19.5 |
| hsa-miR-6784-5p | 30.5 |
| hsa-miR-596 | 28.75 |
| hsa-miR-492 | 35 |
| hsa-miR-7113-3p | 31.75 |
| hsa-miR-4508 | 33 |
| hsa-miR-3606-3p | 81.25 |
| hsa-miR-4286 | 845.25 |
| hsa-miR-2392 | 177 |
| hsa-miR-21-5p | 67880.5 |
| hsa-miR-378a-3p | 867.5 |
| hsa-miR-151a-5p | 1980.75 |
| hsa-miR-27a-3p | 71095.25 |
| hsa-miR-320a-3p | 421.75 |
| hsa-let-7a-5p | 27083.75 |
| hsa-miR-34a-5p | 300.5 |
| hsa-miR-25-3p | 17419.5 |
| hsa-miR-221-3p | 1947.5 |
| hsa-miR-1307-5p | 529 |
| hsa-miR-345-5p | 260.25 |
| hsa-miR-28-3p | 2501 |
| hsa-miR-31-5p | 2736.25 |
| hsa-miR-181a-5p | 18607.75 |
| hsa-miR-22-3p | 19373.75 |
| hsa-miR-450b-5p | 106.25 |
| hsa-miR-28-5p | 197.5 |
| hsa-miR-128-3p | 325.5 |

| | |
|---|---|
| hsa-miR-210-3p | 214.25 |
| hsa-miR-421 | 131.5 |
| hsa-miR-339-3p | 61.5 |
| hsa-miR-3184-5p | 557.75 |
| hsa-miR-106b-3p | 178 |
| hsa-miR-3667-3p | 62.25 |
| hsa-miR-423-3p | 561 |
| hsa-miR-3128 | 97 |
| hsa-miR-518a-5p | 93 |
| hsa-miR-1276 | 41 |
| hsa-miR-432-3p | 16.25 |
| hsa-miR-1303 | 5 |

### 4p1 48HPI

| Fold change | FDR p-value |
|---|---|
| -21.32369586 | 0.000116902 |
| -19.22662547 | 2.22568E-06 |
| -14.44730214 | 0.000667838 |
| -13.8857482 | 5.34888E-12 |
| -10.62499985 | 2.744E-06 |
| -10.31778036 | 0.001456994 |
| -9.526474287 | 1.82511E-14 |
| -9.147474116 | 0.000669297 |
| -8.578034286 | 0.001092371 |
| -7.900058802 | 0.007124941 |
| -7.848442803 | 7.7885E-08 |
| -7.40012597 | 0.006631442 |
| -7.229678621 | 0.040615588 |
| -7.177316788 | 0.005044215 |
| -7.101235101 | 4.50371E-10 |
| -6.779978959 | 2.03047E-05 |
| -5.930504428 | 0.004096229 |
| -5.746126869 | 0.010636312 |
| -5.544228664 | 0.003977253 |
| -5.335696009 | 0.026234875 |
| -5.143208926 | 0.005475076 |
| -4.990490964 | 0.01907494 |
| -4.854172275 | 8.01663E-05 |
| -4.70262946 | 4.97165E-05 |
| -4.622906492 | 0.012130687 |
| -4.393451515 | 0.00010477 |
| -4.387781628 | 0.022821092 |
| -4.235831012 | 0.046092102 |
| -4.117410098 | 0.010381572 |
| -4.066006593 | 0.002040164 |
| -3.927738862 | 0.000959114 |
| -3.860509174 | 0.001201102 |
| -3.85033058 | 0.001064569 |
| -3.744753777 | 0.006856418 |
| -3.623853015 | 4.97165E-05 |
| -3.58257512 | 0.010707379 |
| -3.46330796 | 0.001092371 |
| -3.449046635 | 0.040615588 |
| -3.43276113 | 0.020600071 |
| -3.405231484 | 0.012941455 |
| -3.313746103 | 0.010039264 |

| | |
|---|---|
| -3.159163681 | 0.030214328 |
| -2.965420525 | 0.001171517 |
| -2.872841916 | 0.01303362 |
| -2.850003511 | 0.00470403 |
| -2.807808966 | 0.007581805 |
| -2.804967705 | 0.01633974 |
| -2.71131562 | 0.000569968 |
| -2.699381526 | 0.001214578 |
| -2.678373554 | 0.002739051 |
| -2.60102259 | 0.000158282 |
| -2.5817398 | 0.002096908 |
| -2.560261333 | 0.000116902 |
| -2.417719227 | 0.000669297 |
| -2.359339631 | 0.041702997 |
| -2.33339882 | 0.000669297 |
| -2.319940433 | 0.000569968 |
| -2.295415863 | 0.001862114 |
| -2.204565838 | 0.023390315 |
| -2.17452363 | 0.033891128 |
| -2.132303541 | 0.003970428 |
| -2.054169362 | 0.00803314 |
| -2.021246876 | 0.006996308 |
| -1.975018647 | 0.004785212 |
| -1.879757629 | 0.010039264 |
| -1.775938046 | 0.004096229 |
| -1.774747562 | 4.84052E-05 |
| -1.657361829 | 0.001064569 |
| 1.313619695 | 0.02575761 |
| 1.341250887 | 0.022648599 |
| 1.345792782 | 0.02575761 |
| 1.348751499 | 0.012899168 |
| 1.352687866 | 0.040615588 |
| 1.353496054 | 0.017770006 |
| 1.369579788 | 0.030184929 |
| 1.386439525 | 0.010636312 |
| 1.414808675 | 0.005475076 |
| 1.418195801 | 0.013595954 |
| 1.439457092 | 0.041430516 |
| 1.445522529 | 0.005461007 |
| 1.45064262 | 0.003481963 |
| 1.547176433 | 0.000477427 |
| 1.558080552 | 0.000146972 |
| 1.600776649 | 0.020164164 |
| 1.609576925 | 0.004096229 |
| 1.626684968 | 0.000569968 |

| | |
|---|---|
| 1.662748695 | 0.000959114 |
| 1.663520118 | 0.014157646 |
| 1.692773276 | 0.014157646 |
| 1.735558987 | 0.000146972 |
| 1.773459747 | 0.000790428 |
| 1.820630858 | 0.025888489 |
| 1.90020314 | 8.43759E-05 |
| 2.323919321 | 0.000146972 |
| 2.333665516 | 0.000116902 |
| 2.347277167 | 0.000136251 |
| 2.466191275 | 0.010707379 |
| 4.464356826 | 0.030360483 |
