## Supplemental Table 2 for "Adenovirus 14p1 induced changes in miRNA expression increases lung immunopathogenesis"

| miR27a-3p | miR-21-5p | let-7a-5p | miR22-3p |
|---|---|---|---|
| TERF2 | FAM102B | TERF2 | AHI1 |
| CALU | RTN4 | CALU | FAM102B |
| BMI1 | SLK | ARHGAP1 | RTN4 |
| TMSB10 | RPRD2 | RNF41 | C17orf85 |
| ZKSCAN5 | WWC2 | VPS4A | ARPC5 |
| RNF41 | TMEM16C(hsa) | ERLIN1 | RPRD2 |
| CCNT2 | YTHDC2 | OSBP2 | COPS7A |
| POLR2B | CELF2 | SLK | B2M |
| GOLPH3L | ESR1 | RPRD2 | ESR1 |
| RTN4 | PELI1 | ESPL1 | PLXNA2 |
| VAPB | DDI2 | NEK3 | ATF6 |
| SLK | PLXNA2 | YTHDC2 | SRRT |
| RPRD2 | VPS13C | SEMA6A | SCLT1 |
| ESPL1 | XKR3 | NDUFS2 | RBM14 |
| YTHDC2 | PTPDC1 | PPP1CA | DENND3 |
| SEMA6A | KLHL28 | KIAA0754 | HSPA13 |
| NDUFS2 | RNF11 | TRIB3 | HGSNAT |
| CELF2 | HGSNAT | RBM11 | POU2F1 |
| PPP1CA | TNFRSF11B | SLC44A2 | FASN |
| E2F7 | ADAM9 | PLXNA2 | CDC6 |
| DDI2 | ACTB | C5orf28 | HIST1H1A |
| ABCB10 | CCNG1 | VPS13C | EIF3L |
| PLXNA2 | TANC1 | RGS17 | SLC38A1 |
| BRK1 | EIF3L | MFSD12 | TTC7B |
| PTPDC1 | PRLR | PFN1 | NACA |
| TMPRSS2 | BLOC1S1 | AK2 | DUSP4 |
| C12orf73 | NEK7 | IRS2 | PRLR |
| KLHL28 | PAQR3 | TMPRSS2 | IQCJ-SCHIP1 |
| CAMK2D | MSI2 | CAMK2D | XXYLT1 |
| BRAF | IGFBP5 | BRAF | TENM3 |
| FHL2 | PPP1R37 | POU2F1 | LIG1 |
| HGSNAT | EGR1 | ANP32E | DCP1A |
| ANP32E | ZMAT3 | SLC41A2 | IGFBP5 |
| LAMB2 | NETO2 | UBASH3B | EGR1 |
| FASN | HIPK2 | CDH26 | LRP10 |
| ADAM9 | CNOT6 | ADAM9 | ZEB2 |
| CCNG1 | SCYL3 | ACTB | MLK4 |
| TANC1 | TNS1 | CCNG1 | PICALM |
| PHLPP2 | FBXO11 | HDX | DPP3 |
| PRLR | KYNU | SNAP23 | CNOT6 |
| TMEM50A | URGCP | DUSP4 | BICD2 |
| NRCAM | MARCHF1 | RAB4A | GSK3B |
| NEK7 | ALS2 | PRLR | RNF44 |

| | | | |
|---|---|---|---|
| BSCL2 | SLMAP | MAGI2 | BRWD3 |
| MSI2 | TSC1 | NEK7 | MOV10 |
| SERTAD3 | PTPN3 | PAQR3 | ASB7 |
| SEC16A | RBM27 | SEC16A | FAM120A |
| DCP1A | TXNL1 | PCNXL4 | RBM25 |
| IGFBP5 | VWA9 | EGR1 | SOX4 |
| EGR1 | BRMS1L | FOS | PDPR |
| LRP10 | ENC1 | SFSWAP | ARAP2 |
| PDLIM5 | UGCG | PICALM | TRA2B |
| PICALM | SAR1A | CASP8AP2 | SIPA1 |
| ZMAT3 | STAT3 | FAM195A | WAPAL |
| PDK3 | B3GNT5 | NFE2L3 | TMED9 |
| HIPK2 | CCAR2 | HIPK2 | CHST12 |
| CNOT6 | EPRS | CNOT6 | ESYT1 |
| TMED8 | TNPO1 | RPL31 | RIC8A |
| EDRF1 | GALNT7 | BICD2 | ZBED5 |
| WIPF2 | GLG1 | MEN1 | FARP1 |
| RPL31 | FERMT2 | RNF44 | UBXN7 |
| CRTC3 | ARFGEF1 | BRWD3 | NDUFB10 |
| GSK3B | G3BP1 | FAM120A | SCAMP3 |
| BRWD3 | RAB3GAP2 | C5orf30 | SDF2 |
| CXADR | PDCD4 | GPR63 | FAM96B |
| ASB7 | CDS1 | SOX4 | TRIM11 |
| FAM120A | TGFBR1 | PDPR | VWA9 |
| RBM25 | MTERF | CDIP1 | POLR3D |
| HID1 | PHRF1 | SLC15A4 | TBCD |
| PDPR | CAPN2 | PAGR1 | FKBP3 |
| SLC15A4 | VASH2 | CCNB2 | SPARC |
| FBXL3 | DSC2 | FBXO11 | UGCG |
| FBXO11 | FBXO46 | RNF43 | DNAJC22 |
| HBEGF | CEP152 | RBL2 | GUCD1 |
| UBFD1 | SAFB2 | C9orf40 | STAT3 |
| KYNU | TIPIN | LRRC47 | MYO9B |
| IL1RAP | IRF4 | DUSP22 | RREB1 |
| OAS3 | S100A16 | SCAMP3 | THY1 |
| PYGL | IPO11 | TSC1 | SIGMAR1 |
| URGCP | SMC1A | PHF19 | TNPO1 |
| ZBED5 | RPS7 | ARG2 | PTPRK |
| UBXN7 | C8orf57(hsa) | RBM27 | DIP2A |
| USMG5 | UBXN4 | ANKRD13B | FERMT2 |
| DAZAP2 | PTGFR | VWA9 | ARFGEF1 |
| SMG5 | SPRYD4 | SIN3B | SWAP70 |
| SCAMP3 | MEGF9 | BRMS1L | EEF2K |
| FST | VDAC1 | POLR3D | TOB2 |
| TSC1 | PHF20L1 | ENC1 | NBAS |

| | | | |
|---|---|---|---|
| CLU | TM4SF18 | TUBGCP3 | TM4SF1 |
| APMAP | TTL | FKBP3 | PRR22 |
| RPA3OS | SOCS4 | PCNA | EMP1 |
| PEAK1 | HIF-1Ã□Å½Ã□Â±(hsa) | UGCG | APEX1 |
| ARL6IP5 | WNK3 | STAT3 | EPHB2 |
| PSD4 | FBXL5 | B3GNT5 | SAFB2 |
| SPARC | RPN1 | MYO9B | FEM1B |
| UGCG | CEP97 | RREB1 | TWF2 |
| EPRS | POLR1C | SIGMAR1 | PTPRU |
| MAN2A2 | TACC1 | NAGA | S100A16 |
| RREB1 | EZR | UTP15 | MET |
| SFXN4 | YAP1 | PIEZO2 | MAP2 |
| NAGA | HMGB3 | ARFGEF1 | SENP3 |
| CLPP | PBK | ELL2 | PHF8 |
| TNPO1 | ZBTB44 | TOB2 | C14orf1 |
| GALNT7 | ANKRD26 | G3BP1 | BLCAP |
| UBE2Z | KAT6A | MCAT | MEGF9 |
| ARFGEF1 | CDS2 | RAB3GAP2 | TTYH3 |
| EEF2K | RNF38 | PDCD4 | ANO6 |
| TOB2 | ZBTB38 | TGFBR1 | RPL17 |
| E2F1 | GCLC | TM4SF1 | OPTN |
| TMEM204 | SOS2 | MACC1 | ATP2C1 |
| RCC1 | LTA4H | RTTN | ZBTB39 |
| MASTL | MAP4K2 | ADAM10 | PTMA |
| ASB1 | LSM3 | ADAMTS5 | EPCAM |
| TGFBR1 | WDR47 | PDE4DIP | RNF2 |
| MSI1 | FUS | NAT6 | GPN1 |
| PDE4DIP | BDH2 | EMP1 | SOCS4 |
| CXCL16 | PLEC | COX8A | BCL9 |
| WDR1 | ITGB8 | CHST3 | USP39 |
| TCEB3 | PCSK6 | ANKRD13A | SNX19 |
| APEX1 | FAM126B | TSSC4 | CIC |
| GGCT | RHOBTB3 | LEPROTL1 | UBE3B |
| DSC2 | WDR7 | IRF4 | FAM109A |
| EPHB2 | UBE2T | PDHX | CDR2L |
| FBXO46 | CCDC39 | ARID3A | TACC1 |
| NFKB1 | LAMB1 | PTPRU | YAP1 |
| FAM189B | PTPLB | FAM129B | LSM7 |
| IRF4 | ATF2 | SMC1A | TOP3B |
| UNKL | DDX50 | MET | SSR1 |
| PDHX | RAPGEF1 | UTP14A | ELF2 |
| EXT1 | GPR137B | TROVE2 | SPIRE1 |
| FEM1B | TPM1 | GP5 | DDT |
| RAP2B | ZNF12 | LRBA | C1orf43 |

| | | | |
|---|---|---|---|
| EIF3F | OSR1 | PHF8 | GATM |
| PTPRU | ZNF791 | YWHAH | KAT6A |
| SNX11 | ZNF70 | SRSF9 | RPL32 |
| SMC1A | EIF2AK3 | SPRYD4 | FPGS |
| TOP3A | ALMS1 | SSH2 | ANGEL2 |
| MET | ZGRF1 | UBP1 | RNF38 |
| UTP14A | C11orf95 | TOM1L2 | FUS |
| MFSD1 | RNASEL | C20orf24 | PLEC |
| TROVE2 | TDRD1 | CSK | NELFCD |
| UBXN4 | SCRN1 | GPR85 | PCSK6 |
| SRSF9 | DICER1 | CDT1 | PGR |
| SIPA1L3 | DOCK3 | TTYH3 | RHOBTB3 |
| CBFA2T3 | DNM2 | PPP4C | TM9SF1 |
| BLCAP | RCN1 | TOR1A | TOR2A |
| CNOT11 | AMFR | ATP2C1 | DDX50 |
| MEGF9 | NRAS | ZBTB39 | FLNC |
| SSH2 | UGGT1 | ZNF691 | RAPGEF1 |
| RRAGD | LTN1 | ARL8A | NUCKS1 |
| ID2 | PRKAA2 | NUP62 | WWP2 |
| CSK | ALG11 | GPN1 | FGD5 |
| VDAC1 | IP6K1 | UBR1 | RAPGEFL1 |
| TTYH3 | CRKL | MED13L | HIST1H1D |
| ITGB1 | ATXN2L | MAGEB1 | RFX5 |
| PPP4C | MAP1S | PLCXD1 | PREX1 |
| RPL17 | WHSC1L1 | BCL9 | ATN1 |
| TOR1A | CHCHD3 | SNX19 | FAM83G |
| TTL | TET1 | RPF1 | HDGFRP3 |
| ZBTB39 | LPIN1 | FGD6 | RAD23B |
| GPNMB | SLC7A6 | UBE3B | GNG11 |
| WARS2 | PRRC1 | STOX2 | SH3PXD2A |
| NUP62 | TSC22D3 | FBXL5 | PKP1 |
| UBR1 | CD38 | TACC2 | KRT8 |
| MED13L | PIGX | EZR | CAMSAP1 |
| RAB35 | BID | YAP1 | ABCA2 |
| SRR | FILIP1L | ZBED4 | GEMIN4 |
| SNX19 | BTRC | ADRBK1 | CBL |
| RPF1 | AP3S1 | WASL | USP36 |
| FGD6 | DOCK2 | DDHD1 | C11orf95 |
| CIC | HIST1H2AE | EIF2AK1 | INTS10 |
| STOX2 | SKIDA1 | ZBTB44 | SCRN1 |
| RPN1 | TGFBRAP1 | AREL1 | DICER1 |
| GNA12 | SPPL2A | C1orf43 | ARHGAP11A |
| PET112 | CYB561D1 | PACS1 | DNM2 |
| TACC2 | NT5C2 | ZDHHC23 | RCN1 |
| EFS | ZNF532 | GATM | FBXO21 |

| | | | |
|---|---|---|---|
| ZFAND5 | EXOC5 | KAT6A | PLEKHM2 |
| TTC39C | DCAF8 | MRS2 | EMC10 |
| YAP1 | RFFL | LTA4H | AMFR |
| ADRBK1 | TCF4 | RB1CC1 | PTBP2 |
| NDFIP2 | HNRNPA3 | DNAJC6 | BCAT2 |
| FOXN4 | NBPF1 | RIPK2 | CENPT |
| ZNF621 | KLHL8 | PKN3 | NRAS |
| ZBTB44 | ZNFX1 | PEX19 | IGF2BP2 |
| SSR1 | HIP1 | THEM6 | LPHN2 |
| ELF2 | YOD1 | PLEC | SIK3 |
| AREL1 | WARS | PCSK6 | ADORA2B |
| C1orf43 | ZNF10 | TCERG1 | NDRG3 |
| GATM | APC | PWWP2A | PANK4 |
| ZIC2 | RRN3 | RHOBTB3 | PTPN23 |
| KAT6A | BRD8 | CDK4 | ATXN2L |
| ANGEL2 | KIAA1598 | JMJD4 | ARHGAP19 |
| RNF38 | GATA3 | GPHN | CD59 |
| ZBTB38 | CFL1 | ZCCHC9 | CNOT4 |
| LTA4H | COL12A1 | PTPLB | DGKE |
| ZNF827 | UBR5 | ATF2 | RASA2 |
| RB1CC1 | UHRF1BP1L | LDB2 | PPP2R3A |
| DNAJC6 | TRIAP1 | INHBC | SLC7A6 |
| IL17RA | AFTPH | BLM | YWHAE |
| ACSL3 | ARMC8 | IARS | TRIM52 |
| ALPK3 | THBS1 | KCTD12 | CRIM1 |
| KSR1 | HSPA4 | PREX1 | INPPL1 |
| LSM3 | IL1B | ZNF12 | ELK4 |
| MCM9 | TOR1AIP2 | WIPI1 | COTL1 |
| TRAPPC11 | GTF3C4 | KDM5C | CLIC4 |
| PLEC | ZNF780B | POU3F2 | PEX26 |
| FAM126B | PHACTR2 | ARHGAP27 | FILIP1L |
| LAMB1 | ZMYND11 | PLEKHO1 | BTRC |
| SAR1B | IL12A | ZNF585B | CYR61 |
| TOR2A | CMBL | KCNN4 | SEPT8 |
| ATF2 | LGALS14 | CAMSAP1 | CCNL2 |
| LDB2 | SEPT10 | ABCA2 | CCNA2 |
| NUCKS1 | PRRG4 | GEMIN4 | SEL1L3 |
| BLM | TTLL12 | GARS | KIAA0195 |
| HIST1H1D | CUL2 | CBL | CCT7 |
| KCTD12 | PAK2 | SCRN1 | RMI1 |
| PREX1 | ATMIN | DICER1 | AGAP3 |
| ZNF12 | RPSA | FRMD4A | BTF3 |
| CCDC92 | C11orf30 | ACOX1 | SCARA3 |
| CTNND1 | EDEM2 | C19orf55 | GPATCH8 |
| CHCHD1 | FAT3 | CBX2 | DCAF8 |

| | | | |
|---|---|---|---|
| SH3PXD2A | MAP2K7 | PHB2 | FBXO30 |
| BNIP3 | SAMD8 | LZTS3 | ZNF394 |
| MRPS35 | REST | TRIM27 | CALM3 |
| PKP1 | PGLS | PLEKHM2 | CALM1 |
| KRT8 | PITHD1 | SMUG1 | ZMYM4 |
| ZNF70 | CEBPZ | ZBTB10 | RGL3 |
| ALMS1 | LRRC8B | MAST4 | TCF4 |
| USP25 | B4GALT5 | C15orf41 | ESRRG |
| GARS | GAPDH | SLC37A3 | HNRNPA3 |
| BRD2 | MPDZ | HCN2 | NUP153 |
| CBL | MCM6 | NRAS | TMEM30B |
| C11orf95 | MED21 | IGF2BP2 | LGALS3BP |
| SH2B3 | FKTN | H2AFX | MAP3K3 |
| BCL3 | PLAGL2 | DIEXF | ZNFX1 |
| HNRNPA1 | NF2 | SCAF4 | INHBB |
| DICER1 | ZYG11B | DUSP2 | NME4 |
| CTR9 | CA5B | CRKL | GNAS |
| ARHGAP11A | USP1 | TMEM194A | WARS |
| CBX2 | GSTO2 | NDRG3 | ZNF10 |
| CAMK2G | ANTXR1 | PTPN23 | PPAT |
| DRG1 | WNT5A | ATXN2L | ALDH16A1 |
| DNM2 | RFXDC2(hsa) | C6orf136 | VPS13D |
| CSRP2BP | DLG1 | HN1 | SMTN |
| EN2 | RSF1 | MTHFD2 | PSMA6 |
| TRIM27 | PURA | WHSC1L1 | MAP1LC3A |
| RBM12 | HSH2D | ATXN1L | CC2D1A |
| ZBTB10 | ZNF275 | CNOT4 | SIRT1 |
| PTBP2 | SEPT11 | STK4 | PTPN2 |
| CBFA2T2 | XRCC6 | LPIN1 | MED14 |
| LAMB3 | PAFAH1B1 | ZNF124 | THBS1 |
| MMGT1 | DENND1B | SLC7A6 | IPO9 |
| NRAS | HIST1H1E | YWHAE | HMGA2 |
| PDK4 | ARHGAP24 | HDAC3 | TOMM7 |
| SMIM21 | NAP1L1 | TSC22D3 | CENPM |
| UGGT1 | KDM4B | CD80 | TMEM101 |
| LTN1 | SOWAHC | INPPL1 | GTF3C4 |
| CTB-50L17.10 | ARL2BP | REXO1 | PHACTR2 |
| ADORA2B | NUBPL | TM7SF2 | POLD2 |
| HIST1H1C | PWP1 | ELK4 | CDC25B |
| ARMCX1 | ARHGEF12 | COTL1 | HDAC7 |
| ATXN2L | FLI1 | BID | CYC1 |
| HN1 | ZNF286A | CLIC4 | SKAP1 |
| MAP1S | FAM46C | BTRC | AGO3 |
| WHSC1L1 | SERPINB2 | MMS22L | DBN1 |
| ATXN1L | CD83 | PRR3 | P4HA2 |

| | | | |
|---|---|---|---|
| STK4 | CPS1 | CHD1 | KIAA1522 |
| DGKE | CCND2 | ABT1 | DHCR24 |
| MAP2K2 | PSMD2 | 8-Sep | NINJ1 |
| VPS18 | ANXA1 | CCNA2 | IMPDH1 |
| TPCN1 | NUDT21 | GNS | NPLOC4 |
| PPP2R3A | CORO2A | AGAP3 | LENG8 |
| ZNF124 | FOXM1 | NT5C2 | RPSA |
| YWHAE | SMARCA2 | ZNF532 | C11orf30 |
| TSC22D3 | DOCK6 | MBTPS1 | ANKRD52 |
| CRIM1 | RABGAP1 | SCARA3 | PRRC2B |
| FEN1 | RAB6A | GPATCH8 | SPATA2 |
| ELK4 | HOXA9 | SIMC1 | MTUS1 |
| TMEM59 | LSG1 | ALDH7A1 | REST |
| MORC2 | TTC33 | ATP11C | GBA2 |
| BID | RC3H2 | ZNF710 | LMNB2 |
| CLIC4 | AUTS2 | HOXA1 | LRRC8B |
| HOXB4 | SNX30 | ERO1L | B4GALT5 |
| TIMELESS | SEMA5A | ECE2 | SEL1L |
| PRR3 | LMBR1 | CALM3 | SLC35B1 |
| KBTBD2 | U2SURP | CALM1 | ZNF431 |
| CHD1 | EIF5A | NBPF1 | RBMXL1 |
| CCR3 | ROCK2 | NUP153 | NPAT |
| FAM195B | TESK2 | MAP3K3 | USP31 |
| KLHDC2 | ZNF148 | KDM2B | COPS7B |
| SPPL2A | AKAP9 | NME4 | MCM6 |
| TMED7-TICAM2 | NKTR | YOD1 | PLAGL2 |
| TMEM39A | PITX2 | SLC25A3 | TUBA1B |
| RCAN1 | SKA2 | SOX11 | TMED3 |
| CCNL2 | OTUD6B | CRK | PIK3R4 |
| CCNA2 | MYH10 | SMTN | SSSCA1 |
| GNS | KLF9 | ORC2 | FUCA2 |
| SEL1L3 | DPP8 | NSRP1 | AP2B1 |
| FAM78A | PTPN14 | SIK1 | CHD8 |
| BTBD1 | SYPL1 | MED14 | GPAA1 |
| ZNHIT2 | ZFC3H1 | THBS1 | COL4A5 |
| PPM1K | BACH1 | MEF2BNB | GCA |
| EXOC5 | SAMSN1 | TMEM167A | ODF2 |
| LPCAT1 | GNG12 | JAM3 | ZNF275 |
| ANKRD30B | HUWE1 | MTMR6 | MARVELD2 |
| MAGEF1 | FRS2 | HMGA2 | SEPT11 |
| ZBTB17 | CBX4 | CPSF7 | NRG2 |
| FHL3 | TRIM7 | TOR1AIP2 | PAFAH1B1 |
| ATP11C | SNRK | TMEM101 | ACSF3 |
| ERO1L | STXBP5 | ATF6B | OAF |
| CALM3 | TPCN2 | SUPT4H1 | PITPNA |

| | | | |
|---|---|---|---|
| **DCTN5** | **BUB1** | **MED12** | **RRP9** |
| **LRRC61** | **FICD** | **LGR4** | **STXBP4** |
| **ZMYM4** | **ANKRD50** | **ZNF267** | **MAZ** |
| **RBM48** | **LCOR** | **CYC1** | **BLOC1S1** |
| **NUP153** | **RALA** | **SOX18** | **KLF2** |
| **MIER2** | **MCM4** | **AGO3** | **PVR** |
| **HIP1** | **HRK** | **P4HA2** | **HTT** |
| **KDM2B** | **ACLY** | **KIAA1522** | **ARHGEF12** |
| **GNAS** | **REV1** | **ETV3** | **FLI1** |
| **PCLO** | **KIAA1549** | **TBC1D9** | **SGPL1** |
| **IP6K2** | **TMEM167B** | **EIF3D** | **TP53I13** |
| **FAM115A** | **IQGAP1** | **NPLOC4** | **TRAF7** |
| **GOSR1** | **GAN** | **CUL2** | **YTHDF1** |
| **MKL1** | **ARHGAP32** | **FBXW5** | **YWHAG** |
| **WARS** | **WNK1** | **RPSA** | **SEC23A** |
| **TMED7** | **RUFY3** | **EDEM2** | **MCL1** |
| **IWS1** | **GPATCH2** | **TPR** | **DUSP6** |
| **SMAD9** | **PHF20** | **FAT3** | **VWDE** |
| **LAMA5** | **PBX1** | **ANKRD52** | **TBL3** |
| **BRPF3** | **NBR1** | **PRRC2B** | **RAB7A** |
| **PPME1** | **SETD1B** | **MOB3A** | **PSMD2** |
| **BRD8** | **PREPL** | **PIK3IP1** | **MRPL34** |
| **HMGCR** | **HSPA5** | **THBS2** | **RNF14** |
| **SOX11** | **SLC17A5** | **WDR41** | **ACTG1** |
| **CRK** | **MGA** | **SPATA2** | **PPIL1** |
| **BEND5** | **SNRNP48** | **MTUS1** | **TSPAN3** |
| **CUZD1** | **SOD1** | **REST** | **FTSJ3** |
| **VPS13D** | **SMEK1** | **ELK3** | **NHP2** |
| **FBXW7** | **TPRG1L** | **UAP1** | **SATB2** |
| **DHX29** | **LSS** | **PPP2CA** | **PXN** |
| **SSH1** | **CCDC93** | **TPP2** | **FKBP14** |
| **GATA3** | **SMC5** | **XK** | **ASCL1** |
| **TRIM37** | **TMEM64** | **CAV1** | **RC3H2** |
| **COL12A1** | **LIMCH1** | **INTS1** | **MERTK** |
| **UBR5** | **VPS36** | **IL18** | **STK38** |
| **TRIM24** | **KIFAP3** | **SMARCA4** | **EHD4** |
| **ARF3** | **NUP93** | **B4GALT5** | **MTMR4** |
| **WSCD1** | **CCT5** | **SEL1L** | **INPP5E** |
| **SHC1** | **ZNF664** | **RBBP9** | **HNRNPF** |
| **NDUFAF6** | **ARL14EP** | **CPEB4** | **TACC3** |
| **AFTPH** | **BCL11A** | **NPAT** | **ARPP19** |
| **ARMC8** | **PTBP1** | **UBIAD1** | **ZNF148** |
| **ARRDC4** | **NT5E** | **EXTL3** | **AKAP9** |
| **NR2F6** | **APPBP2** | **TIMP4** | **NKTR** |
| **SIRT1** | **OSER1** | **SPTLC2** | **OTUD6B** |

| | | | |
|---|---|---|---|
| SIK1 | SRGAP1 | MCM6 | PTTG1IP |
| HOXB8 | MAP2K3 | MED21 | HM13 |
| MED14 | FOXK1 | FKTN | DYNC1H1 |
| THBS1 | UQCC2 | TCF7L2 | RNF31 |
| STOM | SPG11 | PLAGL2 | RPS6KC1 |
| NLN | MYH9 | ZBTB7B | ARHGAP17 |
| RUNX1 | HBP1 | DIDO1 | BACH1 |
| TMEM167A | MED13 | MTX3 | GUK1 |
| IPO9 | BIRC5 | NF2 | PKM |
| JAM3 | TUBB | NIRF(hsa) | HUWE1 |
| ZNF521 | C12orf66 | NDOR1 | NTRK3 |
| RBM18 | SLC31A1 | USP1 | MAP4 |
| USP6NL | MACF1 | ACVR1B | FRS2 |
| VANGL1 | PTPN21 | RBM28 | SNRK |
| VPS8 | NADK2 | TSC22D1 | ITCH |
| CENPM | SNTB2 | LRRC8A | BCL9L |
| NUP62CL | PAPPA | UBQLN1 | STX3 |
| GTF3C4 | ATP11B | NR6A1 | SERINC5 |
| ZNF780B | SLC16A10 | XRCC1 | RBFOX2 |
| PHACTR2 | ERG | CSNK1G3 | MKRN2 |
| PIK3CB | FAM83F | PURA | MEX3A |
| CDC25B | PER3 | MYO5B | LTBP2 |
| TRABD | HIPK3 | IST1 | ATXN7L1 |
| HDAC7 | CCR6 | 11-Sep | ANKRD50 |
| ZMYND11 | APH1A | PAFAH1B1 | ANGPT2 |
| SRSF5 | PAPOLA | OGT | CDCA8 |
| PYCR2 | PPP6C | RPS15A | MCM4 |
| AGO3 | CCDC14 | IFITM10 | SETD3 |
| DBN1 | PLD1 | LANCL1 | RNF150 |
| PHF14 | ATP6V1D | MAZ | ACLY |
| SASH3 | CAND1 | NAP1L1 | C7orf50 |
| P4HA2 | RAVER2 | BLOC1S1 | SLC6A8 |
| NCOA7 | AGO2 | PVR | ACAN |
| TBC1D9 | TMX1 | HTT | TOB1 |
| PAXIP1 | AFF4 | ARL2BP | KIAA1549 |
| IMPDH1 | ANKRD17 | ZNF200 | STARD8 |
| GLRB | STAG2 | SZRD1 | UBTD1 |
| AMPD3 | MBTPS2 | YTHDF1 | TRIP11 |
| SMIM8 | CSDE1 | LMBR1L | ASXL1 |
| ATAD2 | DHX30 | NEDD4L | WNK1 |
| LYAR | AXIN1 | YWHAG | RUFY3 |
| TMEM248 | RBM22 | PDZD2 | PARK7 |
| PRKCI | FBXL2 | P2RX5 | PBX1 |
| C11orf30 | USP13 | PTPN4 | C16orf46 |
| EDEM2 | MIB1 | DUSP6 | NBR1 |

| | | | |
|---|---|---|---|
| HLTF | PRDM11 | CD83 | KDM3A |
| C10orf54 | LMNB1 | CPS1 | ARID1B |
| KIAA1211L | MTMR9 | LRRC20 | ZNF562 |
| AKIRIN2 | HMGB2 | POMT2 | SRCAP |
| TPR | SP3 | CDK2 | ITGA5 |
| ANKRD52 | TOMM70A | CCND2 | RAF1 |
| PRRC2B | ATP1B1 | ADAT1 | FNBP1L |
| NSUN5 | TECPR2 | TYK2 | DCAF5 |
| SAMD8 | ARID4A | FAM35A | SMAD3 |
| PVRL2 | NA | BMP5 | PALD1 |
| TNXB | ZNF292 | DNAJB11 | SNX10 |
| THBS2 | TNFAIP3 | COL27A1 | BPTF |
| SPATA2 | CCSER2 | ACTG1 | TMEM43 |
| MTUS1 | ZNF587 | TSPAN3 | CIT |
| REST | ERF | CD74 | SMARCAD1 |
| ELK3 | WEE1 | FOXM1 | SF3B1 |
| UAP1 | AIM1 | FBXL14 | CCKBR |
| TRAPPC2P1 | ZSCAN30 | ETS2 | BCL11A |
| SETD7 | SFT2D2 | RALGAPA2 | PTDSS1 |
| PITHD1 | RCOR1 | ZNF260 | NT5E |
| LMNB2 | MSL2 | ATP6V1F | ERLIN2 |
| TPP2 | IGF1R | EEA1 | GFOD1 |
| CAV1 | CD164 | PXN | AFAP1 |
| CEP128 | OSBPL3 | MARS | LDB1 |
| LRRC8B | CENPQ | HOXA9 | SULF1 |
| GTF3C3 | RPS27A | TOMM20 | ZNHIT6 |
| B4GALT5 | ACBD5 | RC3H2 | TMCO4 |
| SEL1L | FMR1 | ORC1 | TMED5 |
| ZNF431 | DDX17 | GLI2 | CENPW |
| KCNN3 | DOCK11 | SLC25A32 | SRGAP1 |
| CPEB4 | DSE | SNX30 | FOXK1 |
| GAPDH | BTBD3 | PSMB7 | CD70 |
| ADAR | SRPK2 | IFNLR1 | NAP1L4 |
| SUGP2 | TSHZ3 | EHD4 | MYH9 |
| USP31 | CSTB | MAPKAP1 | GOLGA2 |
| DPF1 | KPNA2 | HNRNPF | STYX |
| SPTLC2 | AGO4 | TACC3 | ZC3H7A |
| AFF1 | KLF7 | TRIM35 | BIRC5 |
| MCM6 | ZFHX3 | HIF3A | TUBB |
| PLAGL2 | TPX2 | GMFB | CUL1 |
| ZDBF2 | TMOD3 | U2SURP | MACF1 |
| DIDO1 | LRRIQ2(hsa) | MYOF | CCNF |
| ZNF346 | CLTC | ARPP19 | ADNP2 |
| WFS1 | KPNA4 | BDKRB2 | PHACTR4 |
| MTX3 | SPCS3 | ZNF551 | SQLE |

| | | | |
|---|---|---|---|
| PIK3R4 | SERAC1 | GDF11 | ANKIB1 |
| TSC22D1 | GLS | TALDO1 | KIF13A |
| NR6A1 | VCL | ANPEP | PAPOLA |
| C10orf10 | USP47 | NKTR | AAGAB |
| HHIP | IMPDH2 | CMAS | ELTD1 |
| DLG1 | CTSC | STIM2 | CAND1 |
| PHGDH | VAPA | PTPN14 | PXYLP1 |
| RSF1 | ZNF75D | EIF2AK4 | ALK7(hsa) |
| TAF2 | SNN | DYNC1H1 | MCM5 |
| E2F8 | MAP3K4 | HS2ST1 | CORO7 |
| CHD8 | MAP3K1 | LSM8 | AFF4 |
| PURA | TLR4 | SYPL1 | ANKRD17 |
| GCA | EIF3B | BACH1 | STAG2 |
| ODF2 | HSBP1 | TMEM63A | UBE4A |
| ZNF275 | NA | EFTUD2 | TIMM8A |
| TMEM110 | HN1L | HUWE1 | MBTPS2 |
| IST1 | HADH | MAP4 | ZHX1-C8orf76 |
| ELN | NGFRAP1 | SF3B4 | MRPS16 |
| SEPT11 | VPS26A | FRS2 | HNRNPA0 |
| MBOAT7 | PGM3 | GLMN | OLFM4 |
| MRPL51 | APPL1 | NRBF2 | CSDE1 |
| PAFAH1B1 | SPG20 | PCTP | DHX30 |
| OGT | TMEM164 | NOP10 | AXIN1 |
| IDS | CD2AP | BCL9L | MARCHF9 |
| RPS15A | RAB1A | PHF1 | MCFD2 |
| IFITM10 | PDCD6 | STX3 | MYO1C |
| PRPF40A | HIC2 | HAND1 | QSOX1 |
| ZNF597 | EIF1AX | SERINC5 | HMGB2 |
| NDUFS4 | QSER1 | ALG8 | CLN8 |
| NAP1L1 | HNRNPU | RAD54B | TNKS2 |
| SOWAHC | CXorf21 | RBFOX2 | SP3 |
| HTT | ALDH18A1 | KIF2A | TOMM70A |
| CGN | TSPAN31 | GTF3C1 | PRPF19 |
| SGPL1 | VLDLR | ZNF566 | MOB1B |
| SZRD1 | OLR1 | MESDC1 | ATP1B1 |
| TRAF7 | GID8 | NBEAL2 | STAG1 |
| FAM46C | DCAF11 | ZNF121 | LRP5 |
| APOF | ATP6V1G1 | DIP2B | CDCA5 |
| HYI | DCAF10 | LCOR | TM7SF3 |
| NEDD4L | SPTAN1 | CDCA8 | TECPR2 |
| YWHAG | FOXN2 | GLRX3 | YWHAB |
| PPP1CC | IPO7 | SLC25A25 | SCAND1 |
| MCL1 | DDX3X | ACLY | ZNF292 |
| DUSP6 | PARP9 | SLC6A8 | DAD1 |
| ZDHHC20 | IGF2BP1 | ZNF24 | SLC25A5 |

| | | | |
|---|---|---|---|
| MAPK14 | FLAD1 | PGPEP1 | DTNB |
| DUSP15 | SNRPD1 | RDX | TUBB6 |
| KIAA1191 | ECI2 | IFT122 | FAM83H |
| NUP133 | FSTL1 | ZFP36 | WEE1 |
| GNA13 | MARK3 | KIAA1549 | CDK1 |
| CELSR2 | PRKCE | INPP5F | USF2 |
| TNIP1 | USP15 | WAC | MLLT3 |
| TYK2 | SMARCD1 | GAN | SETD5 |
| BAX | ARHGDIB | NME6 | SFT2D2 |
| TBX3 | FNBP1 | RUFY3 | PHF3 |
| ZNF639 | PPIA | TBC1D12 | NUP160 |
| HSP90AA1 | FASLG | PBX1 | AGFG1 |
| TRMU | MTMR12 | B4GALT6 | NFYA |
| SMARCA2 | SNIP1 | RCSD1 | RCOR1 |
| ADSS | SRSF2 | ZFYVE26 | CLUH |
| RALGAPA2 | DEK | KIAA1671 | TMEM109 |
| MKL2 | CDK2AP1 | SETD1B | BLMH |
| SATB2 | GPR64 | TAP1 | CTDNEP1 |
| PXN | ADD3 | ARID1B | MSL2 |
| MARS | CELF1 | IRGQ | IGF1R |
| TM9SF3 | KLF5 | MARK2 | TRIOBP |
| PNPLA6 | SLC9A6 | ZNF562 | MINK1 |
| RC3H2 | CKAP5 | SRCAP | DDX17 |
| FBXL4 | TAOK1 | MED11 | RAB12 |
| ORC1 | CAPZA1 | SERPINB9 | RFNG |
| CFDP1 | ABHD2 | MGA | PABPC4 |
| TASP1 | SMG8 | CDH11 | DCTPP1 |
| ZZZ3 | KBTBD6 | CDCA4 | TSNAXIP1 |
| EHD4 | TIPARP | PALD1 | C6orf211 |
| MTMR4 | CAPRIN1 | LONP2 | EEF1A2 |
| HNRNPF | CNTRL | C14orf169 | KPNA2 |
| GMFB | RFX7 | BPTF | ZYX |
| ROCK2 | ZBTB20 | FBXO45 | ZFHX3 |
| FAM198B | HK2 | TMEM138 | ARF1 |
| ARPP19 | VPS13A | LIPH | PTGR2 |
| NABP1 | ABI2 | PKD1 | PRR5L |
| VGF | NCOA3 | CHUK | RPUSD1 |
| KATNBL1 | PKNOX1 | NHLRC3 | CLTC |
| ZNF148 | HAUS3 | HOXD11 | ZNF580 |
| SKA2 | CHM | MAP4K3 | MAD2L1BP |
| PTTG1IP | AGPAT5 | VPS36 | NOTCH2 |
| NRBP1 | LPP | SMARCAD1 | PTMS |
| CEP78 | DUS2 | KIFAP3 | SAPCD2 |
| LMBRD2 | ATP2B1 | GMNN | DAZAP1 |
| SH3BP2 | ANLN | ATPAF1 | VCL |

| | | | |
|---|---|---|---|
| PTPN14 | PCDH17 | PIGM | PSKH1 |
| DYNC1H1 | DDX39A | SF3B1 | PTPRF |
| ADCY3 | PLEKHA8 | ZSCAN22 | C6orf48 |
| VARS2 | NCL | ARID3B | CTSC |
| HS2ST1 | PHTF1 | LDB1 | KLF11 |
| ARHGAP17 | RPLP1 | DBF4 | CNIH4 |
| PKM | QSOX2 | TNFRSF11A | WWTR1 |
| GNG12 | MYCBP2 | SELT | IKZF4 |
| EFTUD2 | ZGLP1 | ECHDC1 | ABHD14B |
| MAP4 | MPP5 | TMED5 | ARFGEF2 |
| SF3B4 | TC2N | RNF111 | TIMP2 |
| AGRN | MRFAP1 | FOXK1 | NSMF |
| FRS2 | NHP2L1 | NDST2 | ZNF768 |
| SNRK | ATXN10 | OSTF1 | SPTBN2 |
| CTC1 | SOCS6 | CEP120 | HN1L |
| NPAS3 | CBFB | MYH9 | SMIM14 |
| ITCH | BAG4 | HBP1 | RTKN2 |
| GOLGA8B | TNRC6B | ZNF516 | GTF2B |
| ALG8 | SPTY2D1 | MED13 | CHD7 |
| STXBP5 | C6orf117(hsa) | IMMT | NDRG2 |
| RBFOX2 | EXOC4 | TUBB | APITD1-CORT |
| KIF2A | B3GALNT1 | SLC31A1 | KDM2A |
| LOXL2 | DDX21 | CUL1 | UBE2J1 |
| MESDC1 | PHF6 | MACF1 | DTD1 |
| LTBP2 | PRRC2C | CCNF | RAB1A |
| ZNF121 | LIMS1 | XPOT | KIAA0196 |
| ANKRD50 | CEP104 | TMTC3 | ERBB3 |
| EVA1C | DCTN4 | PAPPA | C14orf80 |
| MDFI | ASH1L | DDX6 | PSAP |
| LCOR | ZSWIM6 | NUP98 | TAB2 |
| RBM23 | PANK3 | PHACTR4 | FBLN1 |
| SLC25A25 | HS3ST3B1 | SLC16A10 | ALDH18A1 |
| ACLY | VARS | HIPK3 | HSPH1 |
| ARHGAP31 | RERE | APH1A | RPL24 |
| SLC6A8 | PURB | PAPOLA | SLC7A5 |
| KIAA0319L | VPS13B | NR4A2 | MAGED1 |
| PAPOLG | SREBF2 | AGO2 | VEPH1 |
| ZNF24 | FAM53C | LIN28B | WDR6 |
| SOAT1 | RECK | IMP4 | CDK6 |
| SAMD9 | CTTN | AFF4 | ATP6V1G1 |
| TOB1 | ICAM1 | SH3RF3 | DCAF10 |
| ZFP36 | RNF149 | FBXO6 | SPTAN1 |
| KIAA1549 | ZNF207 | HMOX2 | SLC25A29 |
| PLEKHJ1 | ITSN2 | EPG5 | IPO7 |
| ARHGAP32 | RFC1 | MBTPS2 | STAT5B |

| | | | |
|---|---|---|---|
| WNK1 | TRIP12 | FOXRED1 | WBP2 |
| MNT | BCAT1 | ZHX1-C8orf76 | PTOV1 |
| RUFY3 | AZIN1 | PRKAR1B | DDX3X |
| ZWINT | RALGPS2 | AGAP1 | PDIA6 |
| GSE1 | PPFIA4 | CRNKL1 | SETD2 |
| GPATCH2 | NOP14 | CHST9 | PARS2 |
| PHF20 | DCAF7 | HNRNPA0 | KIAA1217 |
| PBX1 | TMEM66 | CSDE1 | COPB1 |
| CSNK2B-LY6G5B--991 | RBMS1 | DHX30 | OXSR1 |
| KDM3A | SEC63 | TTC17 | TGFB1 |
| SPATA13 | YME1L1 | AXIN1 | FSTL1 |
| TPST1 | TRIM41 | DGCR2 | CASP2 |
| SNAI2 | SLFN11 | SLC38A7 | NID1 |
| METTL4 | PRR14L | VCAM1 | MVB12B |
| IRGQ | HECTD1 | MIB1 | BDNF |
| PAIP1 | APAF1 | LMNB1 | EYA3 |
| ZNF562 | FAM217B | MEF2D | CORO1C |
| SRCAP | EFHD2 | MTMR9 | MCM7 |
| TRPS1 | FAM46A | ALDH1L2 | FKBP1A |
| MGA | ABCD3 | CHRND | CHD4 |
| CDH11 | MLLT4 | CLN8 | SMARCD1 |
| FNBP1L | LAMTOR5 | TNKS2 | TMEM106A |
| DCAF5 | UBA6 | SP3 | FNBP1 |
| B3GNT7 | FIS1 | SRM | PPIA |
| LONP2 | KRIT1 | MOB1B | MTMR12 |
| SOD1 | EIF5 | ZNF384 | GCC1 |
| BPTF | ARHGAP35 | GOLT1B | HIRIP3 |
| FBXO45 | PTK2 | ANKRA2 | MSN |
| NET1 | PTPN13 | SERINC3 | ASF1B |
| C5orf34 | IDI1 | ADRB2 | ZCCHC2 |
| NHLRC3 | CNN2 | PPP2R5D | ITGB5 |
| TMEM64 | XPO7 | RNF217 | WDR26 |
| CIT | DUSP10 | THRA | POLR3E |
| EYA4 | MAVS | ZNF587 | SOWAHB |
| C7orf43 | PTGES2 | ERF | QTRTD1 |
| CCT5 | ZNF667 | PI4KB | NAA15 |
| MAP4K4 | STK38L | NOD1 | PRPF8 |
| SYT4 | HADHA | ABCC10 | ACAD9 |
| SLC25A51 | ETNK1 | WEE1 | CELF1 |
| C18orf32 | LTV1 | PCNXL3 | PDS5B |
| PTBP1 | LCORL | RFK | SRSF7 |
| ERLIN2 | COPB2 | NUP160 | FBXL7 |
| GFOD1 | CYP4V2 | SLC37A4 | ALKBH5 |
| PEX5 | TFDP2 | CLUH | EPAS1 |
| LIN9 | NCEH1 | ZBTB26 | RABEP1 |

| | | | |
|---|---|---|---|
| CAT | CCDC127 | MSL2 | LBR |
| AFAP1 | BRCA1 | EIF4EBP1 | TUBGCP2 |
| APPBP2 | ZNF217 | PLXNA1 | CAPZA1 |
| PDXK | ARL1 | IGF1R | AMMECR1L |
| SULF1 | SALL3 | IFT57 | ZBTB45 |
| LRP11 | NUP214 | CD164 | CANX |
| U2AF2 | SH3BP4 | ASB13 | SV2A |
| NNMT | LARP1 | PIK3C2B | DPP9 |
| WNT2B | KLHL42 | TNFSF10 | PDS5A |
| ECHDC1 | TAF1 | ACBD5 | TMEM38B |
| TMED5 | PIAS3 | DDX17 | ADIPOR2 |
| MAP2K3 | TRIM59 | DCUN1D2 | CAPRIN1 |
| RNF111 | ZFYVE16 | KARS | SLC3A2 |
| SYNJ1 | CCAR1 | BTBD3 | B4GALT2 |
| RNF144B | LRRC59 | TLE1 | C19orf43 |
| PHB | DOCK10 | ACE | MICB |
| CEP120 | DMD | C6orf211 | KIAA1109 |
| STYX | CSE1L | PSME4 | ACVR1 |
| TPBG | NID2 | KPNA2 | SART1 |
| AP4E1 | POP7 | AGO4 | IER3 |
| MED13 | CERS2 | KLF7 | RFX7 |
| MAP7D2 | ANP32A | ZFHX3 | OXLD1 |
| BIRC5 | OSTC | ZNF195 | FAT4 |
| CUL1 | RAB11FIP2 | SERPINH1 | FJX1 |
| PPP2R4 | KIAA1432 | SPCS3 | LMTK3 |
| MACF1 | BRD7 | LEFTY1 | AGPAT5 |
| XPOT | SMC4 | SLC25A10 | LPP |
| SNTB2 | OSBPL11 | GLS | DPM2 |
| TMTC3 | SFXN1 | VCL | ASGR1 |
| PAPPA | ZNF185 | USP47 | GIGYF1 |
| DDX6 | TXNDC13(hsa) | CTSC | SKP1 |
| ANKIB1 | NFAT5 | GNB1 | YWHAQ |
| PTPN1 | EDC4 | KLF11 | FAM96A |
| PPTC7 | APOLD1 | AMOT | NSD1 |
| HIPK3 | SLC5A3 | VAPA | C12orf10 |
| KIF13A | ATP5G3 | PA2G4 | NCL |
| RPL38 | SF3B3 | ERCC6 | ADORA2A |
| PAPOLA | AGO1 | TXNDC11 | RPLP1 |
| PPP6C | SLIT2 | BMP6 | FAM222B |
| UBL3 | RBM7 | MAP3K1 | UNG |
| CCDC14 | HAPLN1 | CTD-2368P22.1 | SLC16A3 |
| BOD1L1 | CHORDC1 | TLR4 | CPSF1 |
| DCUN1D4 | POLR2A | CHTF18 | ACP1 |
| DNAJB6 | HuD(hsa) | KCTD5 | MPP5 |
| AGO2 | BTG2 | ETF1 | CCDC85B |

| | | | |
|---|---|---|---|
| **WDR46** | **PHC3** | **ASPH** | **RHOV** |
| **GLUD1** | **FBN1** | **HSBP1** | **INSIG1** |
| **AFF4** | **ANKRD49** | **PARG** | **ARFIP2** |
| **IBTK** | **GPD1L** | **PRDM5** | **DFFA** |
| **UBE4A** | **PABPC1L** | **ZNF768** | **RALGAPB** |
| **ADH1B** | **SYBU** | **SPTBN2** | **PAQR4** |
| **UTP20** | **RNF4** | **GTF2B** | **CNP** |
| **AGAP1** | **FAM208B** | **MIEF1** | **PACSIN3** |
| **ITPKB** | **TOP2A** | **CHD7** | **TP53** |
| **MRPS16** | **CCND1** | **H6PD** | **ALYREF** |
| **PRKCD** | **GRPEL2** | **TJP1** | **TBC1D16** |
| **RRP7A** | **C11orf1** | **KDM2A** | **SAMD1** |
| **N4BP2L2** | **PPARA** | **PMP22** | **EMC7** |
| **CSDE1** | **ST6GAL1** | **UBE2J1** | **FGFRL1** |
| **IRF2BP2** | **FNDC3B** | **ZNF510** | **KRT18** |
| **RASIP1** | **C10orf12** | **SLC30A7** | **NTMT1** |
| **PRUNE2** | **CTNNB1** | **FAM13A** | **LITAF** |
| **MYO1C** | **AP3B1** | **NARS** | **LEMD3** |
| **SLC7A11** | **EFR3A** | **HIC2** | **FAM178A** |
| **LMNB1** | **SPEN** | **ATG16L1** | **PRRC2C** |
| **RHOBTB1** | **SNX27** | **QSER1** | **UBB** |
| **HMGB2** | **PTBP3** | **HNRNPU** | **CXorf36** |
| **ALDH1L2** | **VRK3** | **TAB2** | **C14orf119** |
| **TNKS2** | **GPD2** | **PPP1R18** | **FOXO6** |
| **TOMM70A** | **ASNS** | **COQ9** | **CEP104** |
| **PRPF19** | **IRF2BPL** | **ABCC8** | **GHR** |
| **MOB1B** | **RAB27B** | **HSPH1** | **RPL23** |
| **GOLT1B** | **FAM89A** | **UBE2I** | **HDAC4** |
| **MAN2A1** | **BOD1** | **EP400** | **C9orf69** |
| **YWHAB** | **TMEM56** | **GXYLT1** | **ASH1L** |
| **TMEM68** | **PABPC3** | **CDK6** | **MBD3** |
| **RNF217** | **CCT6AP1(hsa)** | **PLOD2** | **SSRP1** |
| **CCDC117** | **SSFA2** | **IGDCC3** | **VARS** |
| **ZNF292** | **RASGRP1** | **NOTCH4** | **EDEM1** |
| **TNFAIP3** | **LYRM2** | **ATP6V1G1** | **SPECC1L** |
| **IPPK** | **STRBP** | **TFAP2A** | **SH3PXD2B** |
| **THRA** | **ZADH2** | **COL3A1** | **HSPG2** |
| **ACAP2** | **SBNO1** | **VPS37B** | **CCDC97** |
| **WEE1** | **GIGYF2** | **HOMER2** | **SDHC** |
| **SETD5** | **DOCK1** | **C1orf21** | **LRPPRC** |
| **SFT2D2** | **INPP5K** | **FOXN2** | **SNX8** |
| **PHF3** | **SKP2** | **FYN** | **ZXDB** |
| **NUP160** | **TSPAN2** | **KAZN** | **ZMIZ2** |
| **RCOR1** | **SOX9** | **DDX3X** | **ARIH1** |
| **PPP2R2B** | **GCN1L1** | **IGF2BP1** | **SDHAF2** |

| | | | |
|---|---|---|---|
| EIF4EBP1 | PI4KA | PARS2 | ADAM17 |
| PLXNA1 | AXL | SLC35A4 | EEF1D |
| LONRF1 | SAMD4A | OXSR1 | DPF2 |
| IL24 | NUFIP2 | FSTL1 | RNF149 |
| MINK1 | POGK | FBXL16 | ZNF207 |
| ASB13 | ZNF650(hsa) | MARK3 | RFC1 |
| PIK3C2B | COPS5 | INIP | TRIP12 |
| MFN2 | NEK1 | HERC1 | MIS18BP1 |
| DDX17 | BLOC1S6 | SHMT2 | AZIN1 |
| RAB12 | TMEM243 | GBF1 | ATP9A |
| DCUN1D2 | DYNC1LI2 | CORO1C | RALGPS2 |
| KARS | CLDN7 | SLC16A9 | VAV2 |
| DNAJC5 | HFE | PHF10 | NPR1 |
| AFG3L2 | PM20D2 | FBL | DYRK2 |
| C6orf211 | NDUFS1 | MID1 | ZNF317 |
| DOC2B | E2F3 | AP1M1 | DCAF7 |
| UGP2 | PAIP2 | GSTO1 | PLCB3 |
| KPNA2 | SNAPIN | FKBP1A | CYTH3 |
| CCDC149 | SMCHD1 | PREP | FRAT2 |
| TMEM41B | DPYSL2 | SMARCD1 | YME1L1 |
| ZFHX3 | ANP32B | TMEM106A | RYK |
| HSPD1 | DMXL1 | FZD3 | FAM43A |
| TPX2 | YEATS4 | CPD | SZT2 |
| WWC1 | TLK2 | ZBTB41 | CHPF2 |
| NACC1 | TARBP1 | SEMA6B | HECTD1 |
| TARDBP | PPAP2B | MTMR12 | NPTX2 |
| SERPINH1 | KLHL6 | MSN | NUMA1 |
| SERPINE2 | ZNF493 | ASF1B | APAF1 |
| CLTC | PAG1 | FARSA | ASAP1 |
| KPNA4 | CSNK1A1 | ZCCHC2 | LAPTM4A |
| SPCS3 | CREBRF | TRAM2 | MTR |
| MAD2L1BP | SPTLC3 | ZNF107 | FAM69B |
| NOTCH2 | RNF24 | ACTA2 | TRIM14 |
| SQSTM1 | MKRN1 | EPC1 | LAMTOR5 |
| GABARAP | F2R | API5 | PCGF5 |
| ZNF800 | KCTD10 | CCDC71L | UBQLN2 |
| GLS | ACTR2 | NAA15 | SCAMP1 |
| ID4 | FOXN3 | PRPF8 | SLC7A1 |
| VCL | MYC | ACAD9 | KRIT1 |
| USP47 | RMND5B | HPS3 | MYO1D |
| WDFY2 | SRPK1 | SPECC1 | SLC6A16 |
| GNB1 | MRPL16 | FBXL7 | ARHGAP35 |
| RASGEF1B | HAX1 | RABEP1 | SLAMF7 |
| ZNF75D | PBRM1 | GALC | CLCN4 |
| PA2G4 | NR2C2 | SLC9A6 | MRPS7 |

| | | | |
|---|---|---|---|
| GNL1 | RNF185 | HS6ST3 | LCMT1 |
| SNN | EPM2A | 7-Mar | FOXP1 |
| LUC7L3 | RAB6C | XPNPEP1 | SOGA1 |
| MINPP1 | SF3B2 | LBR | SHKBP1 |
| MAP3K4 | RAPGEF6 | TUBG1 | XPO7 |
| WWTR1 | FBXL17 | TAOK1 | TIMM22 |
| GM2A | FBXO3 | ALAS1 | RFXANK |
| CXXC1 | DOCK5 | TFPI | TRAFD1 |
| SPAG17 | SACM1L | ERGIC2 | MAVS |
| ARFGEF2 | MYO9A | MLH1 | GNA11 |
| XBP1 | ASB6 | TAB3 | GTF3C2 |
| PPP2R5C | RAD23A | DST | PEG10 |
| COA6 | ELMSAN1 | KIAA2026 | MLEC |
| EIF3B | GPAM | AMMECR1L | ARHGAP18 |
| ASPH | DOCK8 | PLAGL1 | FAH |
| FIZ1 | MMP9 | FAM160A1 | AP000350.10 |
| RHOA | SERPINI1 | SYNM | AMPD2 |
| B3GALNT2 | GPRC5A | RALY | MED16 |
| PCDH10 | LIFR | EPHA7 | ZNF217 |
| TMEM199 | ACY1L2(hsa) | FOXK2 | LARP4B |
| SEC61A1 | SUB1 | SIKE1 | TRIM22 |
| MGLL | DIAPH1 | OSGEPL1 | FAF2 |
| CACUL1 | PTP4A2 | PDS5A | GGA3 |
| SF1 | COG5 | ADIPOR2 | NUP214 |
| RTKN2 | CCT8 | CAPRIN1 | NAA20 |
| CHD7 | CLIP4 | MICB | SH3BP4 |
| HADH | ITPRIPL2 | CNTRL | FAM213B |
| NGFRAP1 | ZFP36L1 | RFX7 | LARP1 |
| VPS26A | GOLGA4 | PQLC2 | KLHL42 |
| KDM2A | KLF3 | NCOA3 | CPT1A |
| PGM3 | UBA1 | ZNF622 | HNRNPUL2 |
| ZNF330 | DEGS1 | LIMD1 | KIAA1147 |
| UBE2J1 | TBCEL | MTUS2 | CSRP1 |
| SNRNP27 | PCBP1 | EIF3K | LRRC59 |
| NAAA | ACOT7 | GIGYF1 | DERL2 |
| APPL1 | MSH2 | LRRC42 | DMD |
| C18orf25 | CPNE3 | ATP2B1 | CSE1L |
| TMEM164 | SLC16A1 | ANLN | BCS1L |
| CD2AP | MTPN | FAM96A | CNIH1 |
| GALK1 | PLAT | PLEKHA8 | NPC2 |
| PRSS23 | SOCS5 | CCR7 | KANSL1 |
| PSAP | SKIL | PREB | BEX1 |
| PDP1 | TFRC | RPS6KB2 | POP7 |
| GOLM1 | BLVRB | ATP5B | MB21D1 |
| HOXC11 | CCL20 | RPLP1 | KBTBD4 |

| | | | |
|---|---|---|---|
| HIC2 | VIM | FAM222B | ANP32A |
| QSER1 | FGB | PNMA2 | SP4 |
| HNRNPU | DYRK3 | ZNF852 | CDH1 |
| MRPS14 | TMED10 | MYCBP2 | KEAP1 |
| CYLD | ATAD2B | ZDHHC3 | FZD4 |
| HSDL1 | REL | UNG | NUDT16L1 |
| PNPLA8 | DMTF1 | ALCAM | DDIT4 |
| HSPH1 | ATRX | SLC16A3 | SFXN1 |
| RCAN2 | FKBP5 | KLF10 | MYH14 |
| VLDLR | ME1 | CPSF1 | NFAT5 |
| SLC7A5 | ECT2 | ZNF57 | UBL7 |
| OLR1 | SET | CSRNP2 | PIKFYVE |
| GXYLT1 | TTC1 | SMARCB1 | PPP1R12A |
| WDR6 | ZMYM2 | NHP2L1 | TET3 |
| WIZ | ENPP2 | G6PD | CMIP |
| SNX15 | FAM20B | C15orf39 | MRPL40 |
| ICAM2 | WASF2 | ALOX5 | DDX54 |
| EZH2 | HSP90AB1 | USP12 | EDC4 |
| CDK6 | ARHGDIA | PPP2R5A | CSNK1D |
| EPDR1 | MYO5A | FAM104A | ZBED6 |
| HEG1 | SLC38A2 | RPS6KA5 | RIF1 |
| CNTNAP2 | NCOR1 | CERCAM | ATF5 |
| MAP3K9 | DDX46 | MFHAS1 | TEX10 |
| RHEB | EIF4G1 | MAML3 | AGO1 |
| DCAF10 | RAB11A | TP53 | NCLN |
| IPO7 | MIA3 | SPTY2D1 | EIF2A |
| GATA2 | ANO5 | EXOC4 | NAV2 |
| DDX3X | SAV1 | LRIG2 | CLIP1 |
| MDH2 | HDDC3 | ADAMTS12 | NOTCH1 |
| SETD2 | CERK | B3GALNT1 | RSU1 |
| RER1 | JMJD1C | ZNF543 | GANAB |
| TGFB1 | NOL4L | LEMD3 | BTG2 |
| FSTL1 | VEGFC | NUAK1 | PHC3 |
| CNNM2 | TIMP3 | PRRC2C | PDLIM1 |
| MARK3 | NCAPG | CDC34 | ORMDL1 |
| INIP | TRIM33 | RICTOR | ARFGAP2 |
| RPIA | HSF2 | ASH1L | FBN1 |
| MVB12B | TGOLN2 | ZSWIM6 | ENO1 |
| TCEB1 | CYB5R4 | RBBP6 | COMMD4 |
| MCM7 | AGGF1 | SRSF1 | NCOR2 |
| SLC16A9 | MUC1 | MFSD4 | SLC16A14 |
| PHF10 | ARIH2 | HSPG2 | EIF4H |
| MID1 | ASNA1 | FADD | PDCD10 |
| SLC35A5 | NCOA2 | HIVEP2 | JUN |
| NOL9 | ZNF354C | ZDHHC12 | CRTAP |

| | | | |
|---|---|---|---|
| GSTO1 | NIPAL1 | HDHD1 | PPP1R15B |
| CHD4 | PIK3R1 | HIF1AN | RBMS2 |
| ARHGDIB | ARMCX3 | PRDM1 | WDR33 |
| MBD1 | SREK1 | PNKD | DCP2 |
| FZD3 | SATB1 | DNAH1 | ENDOG |
| CPD | SKI | GABPB1 | XPO6 |
| FNBP1 | ANKRD27 | ARIH1 | TXLNA |
| ZBTB41 | CSNK2A1 | LEPRE1 | FAM208B |
| BUB3 | MARCKSL1 | DPF2 | PBXIP1 |
| MSN | POMT1 | ITSN2 | RSAD1 |
| SLC35C2 | DHX38 | TMEM132B | TNFAIP2 |
| TPM3 | LRRC57 | BCAT1 | WDR82 |
| RPS20 | CD97 | SOCS3 | USP32 |
| WDR26 | DOCK4 | AMD1 | FMNL3 |
| TRAM2 | TMEM246 | VAV2 | RBM3 |
| ZNF107 | HSP90B1 | ZNF343 | JUNB |
| EPC1 | COPS4 | NPR1 | GIT1 |
| API5 | GLCCI1 | DYRK2 | ZNF805 |
| SREBF1 | HDAC2 | H2AFV | UNK |
| NDUFC2 | NAA30 | ZNF317 | RHNO1 |
| ADD3 | NUSAP1 | TEX2 | MMADHC |
| TANK | PEX5L | DCAF7 | ALG14 |
| PRPF8 | RGS1 | SBF1 | ELOVL1 |
| SLCO3A1 | TMPO | PLCB3 | SPEN |
| CELF1 | SRRM2 | ZNF589 | SNX27 |
| PDS5B | TEX261 | RBMS1 | PPT1 |
| DBF4B | ADAM22 | SEC63 | TUBA1C |
| EPAS1 | KLHL15 | FLII | MSH6 |
| MED12L | FXR1 | UTY | CTNNA1 |
| LZTR1 | DCUN1D3 | YME1L1 | CRLS1 |
| LBR | SENP1 | RIOK2 | SNRPA |
| CKAP5 | EEF1A1 | HEATR3 | MAPRE2 |
| TAOK1 | SLC26A2 | FAM43A | TRIM65 |
| SIK2 | TBK1 | SZT2 | GFPT1 |
| ALAS1 | EPHA2 | TRIM41 | SLC25A13 |
| TFPI | PRS6(hsa) | PSMA2 | SMARCC2 |
| ERGIC2 | DNM1L | SLCO5A1 | NR3C1 |
| ALG9 | EIF5B | TRANK1 | RFTN1 |
| DST | LARP4 | C7orf73 | SHROOM4 |
| CAPZA1 | CUX1 | CHPF2 | ST3GAL2 |
| ABHD2 | ZBTB43 | HECTD1 | MB21D2 |
| HES1 | ZC3H14 | PRTG | PIGC |
| PLAGL1 | SYNE1 | NUMA1 | SLC6A2 |
| BMP8B | LDHA | APAF1 | NRD1 |
| CANX | FAM3C | PEX14 | POLR2H |

| | | | |
|---|---|---|---|
| CDR2 | EIF3J | ASAP1 | KCTD2 |
| F3 | SPRY4 | CRAMP1L | NCOA6 |
| SMG8 | MAPRE1 | FAM217B | MYLIP |
| ZNF746 | MYH11 | JPH3 | NONO |
| SIKE1 | DUSP8 | GREB1L | QARS |
| ADIPOR2 | HMGN3 | ACVR2B | MAOA |
| KIAA1109 | PEBP1 | EDEM3 | PNP |
| ACVR1 | ATP6V1A | PCDH18 | INPP5K |
| IER3 | KMT2D | ELF4 | NPEPPS |
| CAMKK2 | MRPS10 | CD81 | N4BP3 |
| ZBTB20 | SLC25A22 | OXA1L | SH2D3C |
| ABI2 | MAP7 | MTR | PRDM8 |
| NCOA3 | TUBGCP5 | FAM69B | LILRB4 |
| HDAC9 | PEA15 | ATP6V1B2 | FAM179B |
| NFX1 | SESN1 | SLC7A1 | WWC3 |
| WDR5 | CLNS1A | HMGXB3 | ARHGEF37 |
| LIMD1 | ANKRD46 | HCFC2 | NUFIP2 |
| LPP | RHOQ | CKAP2L | POGK |
| IFNAR2 | CDK11A | SERP1 | RING1 |
| TMEM123 | RNMTL1 | PTK2 | TPD52L2 |
| LRRC42 | SCO1 | SLC9A3R1 | IGFBP4 |
| ATP2B1 | TMED4 | C8orf58 | LRRC58 |
| SKP1 | SLC36A1 | FOXP1 | PPP1R10 |
| YWHAQ | TNFRSF10B | NLK | MCC |
| LSM5 | TSNAX | SOGA1 | POGZ |
| SUPT3H | ZNF595 | LAMC3 | NUP107 |
| RCN2 | PIK3C2A | SHKBP1 | MXD4 |
| PCDH17 | MXD1 | XPO7 | SACS |
| PLEKHA8 | RPL19 | DUSP10 | ANP32B |
| PRCC | PUS7 | MAVS | HIST1H2AG |
| TMEM11 | PPM1F | PTGES2 | SEC14L1 |
| FAM222B | CXXC6(hsa) | TES | AKT1 |
| SAFB | KIAA0247 | MYO1F | UBE3A |
| CMTR2 | TMEM2 | EIF2S3 | PGF |
| SNX14 | SNRNP40 | ZBTB37 | ZC3H7B |
| MYCBP2 | PSMC3 | LYPD3 | PARVB |
| ALCAM | CCDC88A | TTC26 | FAM160B2 |
| KLF10 | AP2A1 | GTF3C2 | CSNK1A1 |
| MPP5 | CALD1 | PEG10 | UQCR10 |
| TMEM181 | ADNP | MLEC | CEP68 |
| NHP2L1 | HSPA8 | ARHGAP18 | TRMT1L |
| INSIG1 | NF1 | LRCH3 | FAM134C |
| ATXN10 | FIGN | SRRD | DLC1 |
| ARFIP2 | JPH1 | COPB2 | ACTR2 |
| FAM104A | LYRM7 | SF3B5 | FOXN3 |

| | | | |
|---|---|---|---|
| RPS6KA5 | STRN | POLR2D | IRS4 |
| RALGAPB | IPP | YARS2 | MYC |
| SOCS6 | MED28 | ZNF217 | QKI |
| CBFB | SUV420H1 | ITGAV | TMEM209 |
| MFHAS1 | CLIC2 | GGA3 | CYHR1 |
| SIDT2 | CASC5 | NXF1 | JKAMP |
| MAML3 | YY1 | SALL3 | ACAP3 |
| CUL4A | SRP72 | NUP214 | PCDH1 |
| TP53 | SP1 | NAA20 | SON |
| BAG4 | THRB | ZNF385A | HCFC1 |
| TNRC6B | ARID5B | LARP1 | BMF |
| SPTY2D1 | LURAP1L | KLHL42 | MAGED2 |
| ALDH5A1 | ZPR1 | ABCF2 | ASB8 |
| CPPED1 | JAG1 | MAPKBP1 | FAM83D |
| MOCS3 | RBPJ | PGM2L1 | DOCK5 |
| MITF | ARHGAP21 | CENPK | MARCHF8 |
| ADAMTSL3 | MDM4 | LRRC59 | PLK2 |
| LITAF | LARS | C4orf29 | NIPSNAP3A |
| GLTSCR1L | SLC2A4RG | PTCHD1 | PPM1A |
| DDX21 | BCL6 | GSN | TUFM |
| PRRC2C | BTF3L4 | POLR3A | ACO1 |
| SEMA3F | SMIM13 | TRAK2 | CCKAR |
| RPL23 | CDC73 | ANKRD9 | TP53INP1 |
| ANXA5 | NPTN | CNIH1 | GBP2 |
| C9orf69 | FAXDC2 | THG1L | HIST1H3D |
| RICTOR | BCAS2 | PMAIP1 | SLC4A2 |
| DCTN4 | TNKS1BP1 | RNF145 | MRPS2 |
| ADAMTS2 | TSEN2 | CERS2 | ITFG3 |
| TOP2B | UBE2O | ANP32A | WDR11 |
| ZSWIM6 | ARRDC3 | URB1 | LZTFL1 |
| EGR3 | FAS | SLC9A1 | POLR1B |
| PANK3 | PGRMC1 | OSTC | PGBD2 |
| SSRP1 | ANAPC5 | SP4 | HOXA3 |
| PTER | NMT1 | RAB11FIP2 | DIAPH1 |
| IMPA2 | PRKAB2 | MICU1 | MLST8 |
| RBBP6 | ARID4B | PITPNM3 | PTP4A2 |
| SRSF1 | LAMC1 | DENND5A | SERPINB6 |
| SH3PXD2B | PRICKLE2 | KIAA1432 | TMEM9B |
| PURB | RAI14 | SMC4 | CHERP |
| FADD | RPS6KA3 | NFAT5 | PSMD11 |
| AMMECR1 | TRAPPC2 | PPP1R12A | NLRP1 |
| EVL | NFE2L1 | LIPT2 | ITPRIPL2 |
| HIF1AN | DXO | TET3 | ZFP36L1 |
| SLC35F5 | CDK19 | DROSHA | COL5A1 |
| PNKD | LAMP2 | SLC5A6 | DBNL |

| | | | |
|---|---|---|---|
| GABPB1 | TGFB2 | CMIP | PLEKHA3 |
| EEF1D | FAM160B1 | NAV1 | GPBP1L1 |
| DPF2 | PROSER1 | CCSER1 | BLOC1S5-TXNDC5 |
| RNF40 | MOXD1 | ZBED6 | KLF3 |
| TMEM170B | SUN1 | SLC41A3 | ATXN1 |
| ZNF207 | HDLBP | LFNG | UBA1 |
| ITSN2 | DDAH1 | RUNX1T1 | STRAP |
| UROD | CBLL1 | AGO1 | FAM49B |
| RFC1 | ITPA | FBXL6 | LRRC32 |
| HNRNPK | GAB1 | CRYBG3 | DEGS1 |
| COX19 | AHSA2 | PODXL2 | UMPS |
| TRIP12 | PGRMC2 | EIF2A | CLDND1 |
| DHRS7 | MGEA5 | GGA2 | NMD3 |
| GRN | SLC44A1 | VGLL3 | TMEM86B |
| SIPA1L1 | CKB | ARNT2 | HNRNPA1L2 |
| AKAP8L | TP53BP1 | NAV2 | COMT |
| AZIN1 | GNAQ | ROBO4 | TULP3 |
| GLTP | BCCIP | SMG9 | RILPL1 |
| VAV2 | PLAG1 | KHSRP | YARS |
| DYRK2 | ATXN7L3B | EEF2 | CYP2U1 |
| TEX2 | NFIB | PLCG1 | KANK1 |
| RELT | REV3L | PGS1 | MPLKIP |
| G6PC3 | RASEF | RAB3IP | GPRASP2 |
| DCAF7 | TXLNG2P | POLR2A | MTPN |
| ZNF253 | TTC22 | ARHGEF2 | INTS7 |
| TMEM66 | SNX29 | BTG2 | BIRC6 |
| CREB1 | KLHL3 | CASP3 | FLNB |
| TSG101 | PITPNB | FBN1 | VIM |
| POGLUT1 | MRE11A | ENO1 | DAPK1 |
| PPP4R2 | ATF7IP | NCOR2 | RGS5 |
| CCSAP | WDR3 | CCDC51 | BMP2K |
| HEATR3 | ZNF281 | RGS3 | RPS16 |
| DAP3 | WHSC1 | SLC16A14 | REL |
| CCNT1 | KAT7 | CLCF1 | CCDC47 |
| ARL4C | TOPORS | B4GALT1 | SLC25A24 |
| ANKRD40 | PCDHA9 | ANKRD49 | SRPR |
| ATP6V0B | SCAF11 | MCM3AP | PDGFB |
| EZH1 | CENTB2(hsa) | ARHGEF40 | FRMD6 |
| FAM122B | BCL7A | CRTAP | NDUFB7 |
| PRR14L | HNRPH1(hsa) | PPP1R15B | ATRX |
| ARL8B | VPS41 | PTPN6 | ACTN1 |
| HECTD1 | PARP1 | RBMS2 | POLR2E |
| KLF13 | GK5 | CCT3 | CBX6 |
| ZNF608 | CCR1 | DCP2 | TTC3 |
| ASAP1 | FOXP2 | CLINT1 | KCTD3 |

| | | | |
|---|---|---|---|
| GALNT6 | ZFYVE20 | LRCH2 | RPS6KL1 |
| FAM217B | SERPINB5 | XPO6 | SLC25A17 |
| LAPTM4A | BRD1 | FBXO17 | IRF3 |
| DHX32 | CERS6 | FANCM | HMOX1 |
| ACVR2B | C6orf151(hsa) | DHPS | SMURF1 |
| EFHD2 | GAPVD1 | TMEM173 | C19orf48 |
| SDPR | G3BP2 | MARS2 | ENPP2 |
| OXA1L | SOD2 | DSCR3 | WASF2 |
| MTR | PAPD4 | HMGXB4 | GDE1 |
| MLLT4 | CEP170 | RTKN | IL32 |
| DPY19L4 | TP53I11 | NCAPD2 | TM2D3 |
| SLC7A1 | TNS3 | TOP2A | PRNP |
| ICE1 | SUZ12 | CCND1 | HSP90AB1 |
| FOXP4 | ARCN1 | GRPEL2 | BEND3 |
| EIF5 | TRAPPC10 | ASTE1 | SMNDC1 |
| SERP1 | COL6A3 | URM1 | ARHGDIA |
| PTK2 | PDGFD | RBMX | MYO5A |
| SLC2A3 | SNX13 | USP32 | RPLP0 |
| C9orf41 | ZNF326 | ERGIC3 | ACVRL1 |
| VPS4B | ESYT2 | FMNL3 | CENPB |
| MRPS7 | GNB2L1 | FNDC3B | DDB1 |
| CNN2 | ELOVL7 | JUNB | DDX46 |
| NLK | GALNT2 | MEAF6 | EIF4G1 |
| SOGA1 | LONRF2 | ICMT | RAB11A |
| MAVS | PJA2 | ZNF805 | NIPA2 |
| TIMM10 | GPSM2 | RHNO1 | CEMIP |
| TMEM242 | ABAT | CNOT7 | FLT3LG |
| TES | PPP1R3B | MMADHC | PPP4R1 |
| TICRR | SCD | ZNF678 | VASH1 |
| ZBTB37 | ACSL4 | ACTR3 | RBL1 |
| LYPD3 | GTF2I | LPAR1 | PDK1 |
| SETDB1 | KLHL9 | MAPKAPK5 | NOL4L |
| ETNK1 | SDC2 | B3GNT1 | HNRNPM |
| PEG10 | MYEF2 | RAE1 | CHTOP |
| PPP1R16B | PAPD7 | CRY2 | ZNF382 |
| MOB3B | IL6ST | MSH6 | GBAS |
| PISD | ZNF462 | GPD2 | ZNF138 |
| LCORL | THOC2 | CTNNA1 | AAMP |
| AKT1S1 | PLEKHA1 | RNF25 | FLNA |
| CBLB | FUBP1 | SNRPA | C5orf51 |
| TMEM254 | KMT2A | COL15A1 | TRIB2 |
| GDA | CEP95 | S1PR1 | TRIM33 |
| GPR125 | AKR1A1 | CCNE2 | ATP11A |
| EML5 | GOLGB1 | TRIM65 | TGOLN2 |
| ZNF217 | PHF16 | SLC25A13 | HECA |

| | | | |
|---|---|---|---|
| LARP4B | BAMBI | INTU | SFPQ |
| JOSD1 | UBAP2L | TAPBP | CDC42BPA |
| FAF2 | RTF1 | RFTN1 | RMDN3 |
| NXF1 | MALT1 | E2F5 | ARL6IP6 |
| NUP214 | GRPEL1 | SSFA2 | PABPC1 |
| NAA20 | UBN1 | SEC24C | METTL15 |
| PSD3 | FOXO3 | PHLDB2 | DHX8 |
| LIF | ANKRD28 | SBNO1 | ATIC |
| LARP1 | DERL1 | NCOA6 | ZNF362 |
| CEP192 | GATAD2B | MPHOSPH6 | PNRC2 |
| KLHL42 | ANXA11 | QARS | SCYL2 |
| PRDX4 | RAB22A | USP5 | AEN |
| TAF1 | RAD54L2 | UBXN8 | SMARCC1 |
| HNRNPUL2 | USP7 | CDV3 | SKI |
| ABCF2 | EIF4G3 | PNP | TGIF1 |
| PIAS3 | TMEM30A | NPEPPS | ETV1 |
| NCAM2 | BAG6 | SKP2 | NR1H2 |
| KIAA1147 | RBM12B | PBX3 | SEPP1 |
| KIF3B | BRD2 | ZNF264 | TMCC1 |
| ZNF697 | STMN1 | SH2D3C | PANX2 |
| ARAP3 | UHMK1 | FAR1 | DOCK4 |
| TRIM59 | CCDC34 | TMEM19 | FGF9 |
| PGM2L1 | POFUT1 | COL18A1 | PRR12 |
| WDHD1 | PKD2 | LILRB4 | ARPC1A |
| MAL2 | SYNE2 | SLC35C1 | NPNT |
| DDX24 | SLC25A19 | NUFIP2 | PHC2 |
| FUBP3 | OSBPL1A | POGK | YWHAZ |
| EBF3 | CLSPN | WBP1L | YIF1A |
| C4orf29 | HIAT1 | RAPGEF2 | YKT6 |
| CRTC2 | DOCK9 | TSR1 | ACVR2A |
| DUS1L | TNRC6A | DUSP18 | MBD6 |
| KANSL1 | MDN1 | SLC19A2 | PSMB2 |
| DDX18 | MAP3K2 | ADPGK | RGS2 |
| PMAIP1 | RSPRY1 | TMEM243 | GABRA6 |
| CERS2 | FAM213A | DPYSL5 | TMPO |
| SLC9A1 | USP9X | COX14 | SRRM2 |
| SP4 | SAMD5 | COL4A2 | PLCXD3 |
| ELAVL4 | POLR3B | SLC4A7 | KLHL15 |
| MRPS18B | MBNL1 | TRAK1 | SLC30A9 |
| PDHA1 | PRC1 | DYNC1LI2 | EPHA2 |
| FZD4 | RP2 | SOCS1 | PIGV |
| KIAA1432 | MON2 | CCDC28B | DDX5 |
| BRD7 | SRSF11 | LRIG3 | LARP4 |
| SFXN1 | MYBBP1A | CNNM3 | CUX1 |
| CUL5 | COBLL1 | PM20D2 | MIF |

| | | | |
|---|---|---|---|
| TRMT5 | ATP2B4 | GIT2 | WSCD2 |
| NFAT5 | OGFOD1 | E2F3 | FAM219A |
| MFSD9 | ZRANB1 | MAST1 | ZC3H14 |
| PPP1R12A | PAN3 | RGMB | SLC35B2 |
| LIPT2 | AVL9 | SACS | RAC1 |
| SLC35F3 | MCAM | ZNF443 | S100A10 |
| MBTD1 | KIF21A | ITM2C | H3F3C |
| TET3 | EIF4EBP2 | GDAP2 | ANKRD13C |
| DROSHA | MLXIP | ACER2 | RANGAP1 |
| SLC5A6 | RASGRP3 | SEC14L1 | POF1B |
| NAV1 | PTX3 | MAPK8 | MAK16 |
| CCSER1 | HNRNPA2B1 | KPNA5 | CTNNAL1 |
| KLHDC10 | NAA25 | AKAP8 | ANKFY1 |
| ZNF655 | ASRGL1 | SMAGP | CCDC58 |
| DDX54 | PFKFB2 | RQCD1 | CTPS2 |
| CSNK1D | TPT1 | SLC35D2 | PAK4 |
| SLC5A3 | SPC24 | PARVB | PEBP1 |
| ZBED6 | ZBTB1 | CDK9 | ABR |
| LFNG | OSBPL6 | PAG1 | NOLC1 |
| ATP5G3 | TRIM2 | CSNK1A1 | ATP6V1A |
| SF3B3 | GPR155 | CCR5 | DNMT3A |
| RAB30 | CCRN4L | UQCR10 | KMT2D |
| KDELC1 | TCEAL3 | CEP68 | ADAMTS1 |
| KCTD21 | SUGT1 | DNAJC13 | FSCN1 |
| AGO1 | KANSL1L | TRMT1L | UBE2K |
| NECAP1 | DDX1 | ZFHX2 | THAP2 |
| S1PR3 | LIMA1 | SLC29A1 | TSPAN33 |
| TBC1D4 | NDUFS5 | F2R | KDM6B |
| MICU3 | CCNI | WDR48 | MANSC1 |
| AP1S2 | CUL3 | C1orf226 | SESN1 |
| DLX5 | BTBD7 | DLC1 | RARA |
| OGDH | GID4 | POLD3 | CTIF |
| KLRG2 | FAM63B | DNMT1 | ZFP64 |
| ATG101 | PCBP2 | HSBP1L1 | HES6 |
| YIF1B | RASSF8 | MYC | BCL2L2 |
| NAV2 | AHNAK | DEDD | ANKRD46 |
| SLIT2 | TRPM7 | ZDHHC6 | PPP1R9B |
| KLKB1 | NRIP1 | NDUFA3 | DCLK2 |
| ROBO4 | PRKDC | IFI44L | SESN2 |
| PCNX | SASH1 | p27(hsa) | PHKG2 |
| TTC37 | ZBTB34 | PBRM1 | HIST1H3H |
| PKP4 | OLA1 | CTC-534A2.2 | TRIO |
| C1orf52 | SMAD7 | NR2C2 | KIF11 |
| STAMBPL1 | UBE2W | PCDH1 | CELF6 |
| KHSRP | ZNF726 | SON | FRYL |

| | | | |
|---|---|---|---|
| **CLIP1** | **MKNK2** | **TRIM3** | **GMPPB** |
| **EEF2** | **PSRC1** | **SF3B2** | **PIK3C2A** |
| **EXOSC1** | **HERPUD2** | **CD46** | **OR10S1** |
| **RAB3IP** | **PLK1** | **RGL2** | **SURF4** |
| **BTG2** | **ZNF367** | **FBXO3** | **MPI** |
| **PDLIM1** | **JADE1** | **PRKAG1** | **C16orf72** |
| **ARFGAP2** | **TSN** | **MKLN1** | **PPM1F** |
| **FBN1** | **ARHGEF7** | **SMIM15** | **IGF2BP3** |
| **TMTC4** | **TGFBI** | **MRPS9** | **TMEM2** |
| **TOMM5** | **DLAT** | **PLK2** | **SSX2IP** |
| **GPD1L** | **PPARÃ□Å½Ã□Â±(hsa )** | **GALNT1** | **MGST3** |
| **PAQR7** | **HIST1H1B** | **HLA-DPA1** | **DHX15** |
| **ZNF711** | **CLCN5** | **MYO9A** | **EMP2** |
| **MFN1** | **DLX2** | **PSAT1** | **DYNLL2** |
| **PIK3R3** | **ABCC5** | **PIGP** | **CCDC50** |
| **ZDHHC2** | **CMTM6** | **HSPA14** | **PSMD10** |
| **CRTAP** | **RNF6** | **COPS3** | **ZC3H11A** |
| **PPP1R15B** | **MOAP1** | **ASB6** | **ADNP** |
| **WDR33** | **KLHL24** | **MATN2** | **HSPA8** |
| **ABHD17B** | **ALDOA** | **VEZT** | **CRHR2** |
| **CCT3** | **NR1D2** | **DOK3** | **PSEN1** |
| **FBXO28** | **PTAR1** | **VCAN** | **SMARCD2** |
| **DCP2** | **EXOC8** | **USP21** | **CDK13** |
| **FBN2** | **DOCK7** | **PCDHAC2** | **UBXN11** |
| **XPO6** | **ACAT1** | **WSB1** | **EIF2S1** |
| **RNF4** | **IVNS1ABP** | **TP53INP1** | **GPX8** |
| **TXLNA** | **CDC42SE1** | **CMKLR1** | **SUV420H1** |
| **CATSPERB** | **RP1-130H16.16** | **CCL7** | **GNG5P2** |
| **TMEM173** | **SPIN1** | **SLC4A2** | **CXXC5** |
| **MARS2** | **PDZD8** | **C12orf29** | **HIVEP1** |
| **HMGXB4** | **UTRN** | **MRPS2** | **SP1** |
| **FAM208B** | **MAP2K4** | **WDR11** | **DAB2IP** |
| **TOP2A** | **CNOT6L** | **GPRC5A** | **ARID5B** |
| **CCND1** | **EHD1** | **LIFR** | **SCARF1** |
| **IMPAD1** | **ZNF518A** | **ST3GAL6** | **LURAP1L** |
| **PBXIP1** | **CLSTN1** | **SUB1** | **CLDN1** |
| **ILF3** | **XRCC5** | **IGF2** | **MCRS1** |
| **TSSC1** | **RYBP** | **PTP4A2** | **MFSD5** |
| **LHFPL2** | **YAE1D1** | **CCT8** | **IDH2** |
| **WDR82** | **RND3** | **THOC5** | **DZIP1L** |
| **EPHB4** | **PAMCI(hsa)** | **CHERP** | **SLC39A6** |
| **ERGIC3** | **ALPP** | **RBM19** | **SYNJ2** |
| **FNDC3B** | **TXNL4A** | **TK1** | **FAM168B** |
| **JUNB** | **GTF2A1** | **PSMD11** | **C19orf10** |

| | | | |
|---|---|---|---|
| MEAF6 | EPHA4 | FLRT2 | DNTTIP1 |
| CTNNB1 | TRIM71 | ARPC2 | GTPBP2 |
| CNOT7 | UNC13B | SPRYD3 | NPTN |
| ZNF678 | PHIP | ORMDL2 | ZNF451 |
| POLDIP3 | ST5 | PRKCSH | MAX |
| PUM1 | VEGFA | MAPRE3 | CCDC61 |
| ELOVL1 | CPEB3 | ZFP36L1 | INO80E |
| SPEN | PTEN | SMAP1 | AHSA1 |
| MAPKAPK5 | SYT14 | GOLGA4 | TNKS1BP1 |
| THOC3 | ZC3H4 | CDCA7L | UBE2O |
| SNX27 | DUSP16 | ARID1A | ARRDC3 |
| PTBP3 | MARCHF6 | PANK2 | IKZF2 |
| CBX1 | DTX3L | FAM49B | ANAPC5 |
| PPT1 | HPS5 | MTRF1 | STX6 |
| CRY2 | PALLD | DEGS1 | ILF2 |
| NEBL | SPTBN1 | CLDND1 | SLC20A1 |
| GPD2 | PI4K2B | NMD3 | NFE2L1 |
| CTNNA1 | ATL3 | LLGL1 | NEDD4 |
| IRF2BPL | LATS1 | DPH3 | CNPPD1 |
| PMM2 | ACSL6 | FBXO8 | ATP5E |
| GFPT1 | IRAK4 | MTPN | MKI67 |
| INPP4A | DAAM1 | RRAGA | RRP1B |
| NR3C1 | SOX2 | CTSZ | UBE2V2 |
| TEX22 | CAP1 | SOCS5 | PRKAR1A |
| RFTN1 | ALDH1A1 | E2F6 | PEF1 |
| E2F5 | RNF213 | SKIL | VMA21 |
| SHROOM4 | TAF5 | TFRC | CSNK1E |
| SSFA2 | DYNC2H1 | CABLES1 | SGK494 |
| ST3GAL2 | PIGN | BLVRB | CYB561A3 |
| MB21D2 | MAPK1 | PCNXL2 | EP300 |
| NRD1 | LAMTOR1 | TMEM38A | POM121 |
| MMP1 | IMP3 | ADAM12 | PPWD1 |
| POLR2H | NIPBL | XKR7 | AMOTL2 |
| GIGYF2 | SLAIN2 | GPR45 | SLC10A7 |
| KCTD2 | NUPL1 | TMED10 | GHITM |
| MYLIP | RNF168 | KPNA1 | ZNF35 |
| RPL17-C18orf32 | TLN1 | MTMR14 | MGEA5 |
| TRAPPC9 | NUP205 | IFI16 | MPRIP |
| CTTNBP2 | RCC2 | ANO2 | ARFIP1 |
| BHLHE40 | RRAGC | SLC25A39 | HARS |
| ARL6IP1 | PPP2R1B | P2RX4 | ZFYVE19 |
| EPHA3 | CHD9 | SLC25A24 | CASZ1 |
| ASPM | SESTD1 | FCHO2 | GNAQ |
| CDV3 | PRDX3 | PDGFB | REV3L |
| DOCK1 | CXCL10 | COL1A1 | EXOC2 |

| | | | |
|---|---|---|---|
| **NPEPPS** | **IL10** | **CHD3** | **ECI1** |
| **C2orf44** | **MMP-2(hsa)** | **KCTD3** | **AKT3** |
| **FAR1** | **AP3M1** | **XPR1** | **PRPF4** |
| **AQP1** | **BOC** | **IRF3** | **ZNF484** |
| **SAMD4A** | **KMT2C** | **CDC42EP4** | **CELSR3** |
| **NUFIP2** | **CYBRD1** | **PSMD14** | **PITPNB** |
| **GCC2** | **MGAT4A** | **CNNM4** | **CLN6** |
| **NEK4** | **CCNG2** | **SLC11A2** | **MYOZ3** |
| **WBP1L** | **GOLGA3** | **NDUFV3** | **ATF7IP** |
| **RAPGEF2** | **ZSCAN9** | **WASF2** | **BRD3** |
| **TSR1** | **GTSF1** | **GDE1** | **GRK6** |
| **SEC23B** | **TLDC1** | **ARHGDIA** | **YIPF5** |
| **BLOC1S6** | **NUCB1** | **ZBTB5** | **POMC** |
| **MSL1** | **PNPLA3** | **MYO5A** | **PHF12** |
| **AOX1** | **ZNF567** | **LRP1** | **TOPORS** |
| **TMEM243** | **MRPL49** | **SLC38A2** | **HNRNPH2** |
| **MRPL30** | **C20orf194** | **MYCBP** | **MZT1** |
| **LRRC58** | **COL5A2** | **COQ10B** | **DMPK** |
| **SLC27A1** | **ARF4** | **BAZ1A** | **CREB3L2** |
| **DYNC1LI2** | **ENAH** | **NCOR1** | **TSHZ1** |
| **CISH** | **SGCB** | **ZNF598** | **DHRS13** |
| **PAICS** | **RNF17** | **HIC1** | **CDH5** |
| **CCDC28B** | **KIF5C** | **TBRG4** | **RIN2** |
| **HIF1A** | **FAM79A(hsa)** | **DDB1** | **GBE1** |
| **PPP1R10** | **RHOB** | **EIF4G1** | **PARP1** |
| **HELZ** | **BRWD1** | **RAB11A** | **CTSB** |
| **POGZ** | **CBX5** | **RBBP5** | **H3F3B** |
| **PAIP2** | **TEK** | **NIPA2** | **PPFIBP1** |
| **RGMB** | **TBL1XR1** | **CEMIP** | **MPHOSPH9** |
| **SACS** | **SETX** | **CEP44** | **CAPN1** |
| **C2CD2** | **BAZ2B** | **SHISA5** | **GRAMD1A** |
| **CTNNBL1** | **PIGO** | **DENND6A** | **CLIP2** |
| **C8orf4** | **IREB2** | **LYL1** | **DDX39B** |
| **SLIT1** | **NA** | **VASH1** | **SOD2** |
| **ANP32B** | **TGFBR2** | **CERK** | **PAPD4** |
| **SEC14L1** | **COLGALT1** | **HIST1H2BG** | **FLT4** |
| **MAPK8** | **MEF2A** | **XYLT2** | **CEP170** |
| **KPNA5** | **GRSF1** | **KDM3B** | **AHCY** |
| **ICE2** | **DDHD2** | **JMJD1C** | **CHGA** |
| **PPAP2B** | **COPS6** | **NOL4L** | **ARCN1** |
| **STAT2** | **ZBTB47** | **HNRNPM** | **HNRNPH1** |
| **PDE8A** | **SNRNP70** | **RNMT** | **COL6A3** |
| **DNAJB9** | **RAPH1** | **SNX16** | **DOK4** |
| **RIT1** | **GCNT2** | **SMAD5** | **CCDC88C** |
| **HERC2** | **NBEA** | **BTN2A1** | **BSG** |

| | | | |
|---|---|---|---|
| CYP24A1 | ITFG1 | PDE12 | RACGAP1 |
| CSNK1A1 | MDM2 | FLNA | FASTKD5 |
| CREBRF | ELOVL4 | CYP2R1 | CAPN15 |
| CGGBP1 | PSMB5 | C5orf51 | EIF3M |
| DNAJC13 | PGK1 | PSMC4 | SYNGAP1 |
| KIDINS220 | SLC39A14 | TGOLN2 | TPST2 |
| LNPEP | TOMM6 | PTGS2 | BAHD1 |
| MED26 | NIN | SLC39A9 | GPSM2 |
| AFF3 | LPGAT1 | RNF20 | C5orf24 |
| STAU1 | PPAP2A | DAG1 | ASS1 |
| OGG1 | FANCI | RNGTT | NTN4 |
| FOXN3 | DDR2 | SF3A2 | NCDN |
| RMND5B | TRIM38 | WAF1(hsa) | CYP1B1 |
| SRPK1 | KIAA1715 | UQCRH | SCD |
| HOXA5 | VPS54 | MRPL17 | PDIK1L |
| LIMK2 | PCMT1 | MSANTD2 | NR1D1 |
| MRPL16 | MEIS1 | TBC1D25 | LGALS1 |
| HAX1 | CNDP2 | CFL2 | CHAF1A |
| NR2C2 | HAT1 | HOOK1 | PLEKHG2 |
| KIF4A | BAZ1B | KLHDC3 | NGRN |
| CENPF | CDC123 | ID1 | EDC3 |
| SULT4A1 | SMG1 | GINS4 | KIF18B |
| IKZF5 | MCM3 | APBA2 | ZNF462 |
| SON | BMPR2 | FZD2 | NBEAL1 |
| HMCN1 | KIAA1551 | SEC24A | NUP43 |
| PIM3 | EIF4G2 | TCF20 | FAM102A |
| HCFC1 | CLOCK | TEAD1 | KMT2A |
| SF3B2 | KBTBD7 | RB1 | RARG |
| CD46 | KIF5B | WDR13 | RPN2 |
| AHCTF1 | COG1 | AEN | SCAF1 |
| RGL2 | 39701(hsa) | BCAP29 | EPS8 |
| FBXO3 | LMO4 | NEK9 | LGALSL |
| PRKAG1 | CCDC6 | SREK1 | BAMBI |
| AHDC1 | ZDHHC7 | BAZ2A | TDP1 |
| FAM83D | LIN7C | SMARCC1 | LRRC1 |
| MKLN1 | PRRX1 | SKI | HPS6 |
| PLK2 | PLS3 | NADK | UBAP2L |
| GALNT1 | USP34 | CSNK2A1 | ALDH9A1 |
| TMEM91 | GNE | FAM118A | MALT1 |
| MYO9A | CD44 | MARCKSL1 | DNHD1 |
| STRN4 | KIAA0020 | TMEM185B | TPP1 |
| EDNRB | CD47 | PRPF31 | FOXO3 |
| AGAP2 | KIAA1430 | ARSK | HDAC6 |
| SFXN5 | COL4A1 | ARL4D | ZNF585A |
| VCAN | ZKSCAN1 | PRPF38B | DDR1 |

USP21
MTCH1
USP46
WSB1
ACO1
TP53INP1
INSIG2
GPAM
MLH3
ITFG3
FXYD6
LIFR
LBH
POLR1B
BCL7B
SUB1
DIAPH1
PTP4A2
TMEM9B
CHERP
SPRYD3
PRKCSH
CLIP4
RELN
ZFP36L1
COL5A1
PLEKHA3
GOLGA4
GPBP1L1
BLOC1S5-TXNDC5
KLF3
ATXN1
UBA1
EPS15
ESM1
PRDX1
STK35
PPP2R1A
FAM126A
IGF2R
DYNLT1
KANK1
UBAP1
SLC16A1
CLPX
SNAP29
CDC25A
BNIP2
FNIP2
C2orf43
RXRA
ATP5O
ZBTB8A
EIF3I
DSP
EDIL3
GSPT1
ZDHHC13
RMND5A
C7orf55-LUC7L2
ZCCHC3
TXNIP
CCR6
NCOA4
TADA2A
RP1-130H16.18
GTF2I
SLC6A14
CTDSP2
CPNE1
ZNF28
USO1
CNKSR3
UBR2
DOCK4
IDH1
POM121C
HSP90B1
FARP2
H2AFY2
PHC2
YWHAZ
C11orf84
YLPM1
CUEDC2
EIF4ENIF1
GLCCI1
HDAC2
FBXL19
NRP1
KIAA0284
PRKAR2A
ZNF256
NAA30
NTPCR
MBD6
GDI1
PHLPP1
PSMB2
WDR81
SEMA4C
ZC3HAV1
DHTKD1
PPP2R3C
CHAC1
RAP2C
KAT2A
TMPO
SRRM2
RNFT1
TEX261
RPAP1
EOGT
PEX11B
SENP1
TBK1
MAPK8IP3
LMNA
SLC7A2
TULP4
TSC22D4
FAM136A
STARD7
ATF3
IPO8
MYO1E
BASP1
SGTA
CTCF
GFRA1
PXDN
FAM173A
RAD54L2
AKTIP
COBL
APOOL
MATR3
LRIF1
TNC
AKAP12
GSDMD
C21orf59
TMEM30A
PPAN-P2RY11
BAG6
CDKN1A
STMN1
UHMK1
PRMT2
BDP1
KIAA0232
SYNE2
OSBPL1A
NANS
EDARADD
GPT2
TNRC6A
TP53INP2
MBNL1
WDR43
ZFHX4

**GALNT3**
**SOCS5**
**E2F6**
**SKIL**
**TFRC**
**BIRC6**
**IKBKAP**
**SLC25A23**
**BLVRB**
**FLNB**
**DSN1**
**DAPK1**
**BMP2K**
**NEK6**
**JRKL**
**SPAG9**
**SUPT6H**
**MAPKAPK3**
**TMED10**
**ATAD2B**
**REL**
**IFI16**
**SRPR**
**FRMD6**
**HAUS6**
**DMTF1**
**WT1**
**GLO1**
**VCP**
**CBX6**
**CHD3**
**ZMYM1**
**ECT2**
**XPR1**
**MAFK**
**ANKLE2**
**ZBED6CL**
**ZMYM2**
**SMURF1**
**GPR56**
**APLP2**
**GIPC1**
**NDUFV3**
**TM2D3**
**GMPS**
**ICOSLG**
**EPHA2**
**ASCC3**
**SYT7**
**CUX1**
**SEC31A**
**XRN1**
**ZNF74**
**RAC1**
**INSR**
**SNX2**
**SRC**
**TCOF1**
**MAPRE1**
**ZSWIM4**
**STC2**
**PRKAA1**
**PRMT1**
**CAP2**
**ANKFY1**
**DDOST**
**CTPS2**
**FAM208A**
**PTP4A1**
**PEBP1**
**NOLC1**
**SHOC2**
**PMPCB**
**KMT2D**
**HOXC5**
**ADAMTS1**
**FHDC1**
**FSCN1**
**LDLRAD3**
**KDM6A**
**UBE2K**
**THAP2**
**KDM6B**
**IPO4**
**SESN1**
**DOLK**
**FAM122A**
**ANKRD46**
**TSEN34**
**SESN2**
**ANKHD1-EIF4EBP3**
**CNEP1R1**
**MCAM**
**KIF21A**
**ATG9A**
**ZBTB18**
**TDRP**
**EIF4EBP2**
**TUBA4A**
**SMG7**
**D2HGDH**
**FAT1**
**HNRNPA2B1**
**FOCAD**
**POLG**
**NOP9**
**MYL6**
**HK1**
**ZBTB1**
**CHST2**
**ADARB1**
**TTI1**
**PPM1B**
**TNPO2**
**ZBTB4**
**NOL8**
**IFI27L1**
**GORASP2**
**PTPN12**
**LMF2**
**CCNI**
**TMEM263**
**ERalpha(hsa)**
**HNRNPL**
**RUFY2**
**PHF13**
**CCNJL**
**HSD17B4**
**RASSF8**
**STAT1**
**HIPK1**
**AHNAK**
**LUZP1**
**YTHDF3**
**PRKDC**

| | | |
|---|---|---|
| **PRNP** | **NARF** | **SASH1** |
| **HSP90AB1** | **RAP1GDS1** | **ZBTB34** |
| **ZFYVE9** | **TRIO** | **AP1G1** |
| **NFKBIA** | **BMP2** | **SMAD7** |
| **MYO5A** | **KCNC4** | **PAFAH1B2** |
| **BAZ1A** | **TMED4** | **KIF1C** |
| **HIC1** | **CS** | **PRKAR2B** |
| **ZNF100** | **TNFRSF10B** | **EHD2** |
| **SNAPC4** | **NLGN2** | **NDUFV1** |
| **RBBP5** | **KDELR1** | **SETD6** |
| **NIPA2** | **TIA1** | **RHBDD1** |
| **CEMIP** | **SURF4** | **SPHK1** |
| **PPP4R1** | **DZIP1** | **PPIF** |
| **SHISA5** | **C16orf72** | **TRRAP** |
| **RAB18** | **MRGBP** | **ZFP36L2** |
| **TMEM87B** | **TSC22D2** | **C11orf31** |
| **C17orf58** | **PYGO2** | **PIEZO1** |
| **DENND6A** | **SLC12A9** | **PPM1G** |
| **HIST1H2BG** | **IGF2BP3** | **ZNF3** |
| **XYLT2** | **ELAVL1** | **HIST1H1B** |
| **KDM3B** | **COPG1** | **ZNF106** |
| **JMJD1C** | **CCER2** | **PSMD8** |
| **COPZ1** | **PHKA1** | **FAM134A** |
| **RNMT** | **TMEM2** | **ZC3HAV1L** |
| **GLUL** | **LYPLA2** | **KHNYN** |
| **RBM26** | **SSX2IP** | **MAFG** |
| **GBAS** | **EMC4** | **AGPAT3** |
| **GALK2** | **AP2A1** | **ULK1** |
| **KIAA2013** | **CCDC50** | **GATAD1** |
| **TLE4** | **CALD1** | **PRKD2** |
| **MID2** | **ADNP** | **UQCRC1** |
| **NCAPD3** | **HSPA8** | **CTDSPL2** |
| **CLK1** | **MAD2L1** | **HTR2C** |
| **ZNF138** | **LCLAT1** | **CMTR1** |
| **H2AFZ** | **C6orf120** | **NR1D2** |
| **C5orf51** | **FIGN** | **RPS4Y1** |
| **TRIM33** | **CCNY** | **FGFR2** |
| **ELOVL5** | **PNPO** | **METTL3** |
| **MMD** | **CDK13** | **UBXN2B** |
| **CDC42BPA** | **STK25** | **MOB1A** |
| **ARIH2** | **RNF152** | **HEY1** |
| **PATZ1** | **MED28** | **KIRREL** |
| **DAG1** | **UBAP2** | **CHST1** |
| **RMDN3** | **RANBP2** | **ITGA6** |
| **CACNG4** | **DSG2** | **SLC30A3** |

| | | |
|---|---|---|
| **RNGTT** | **SUV420H1** | **PRKRA** |
| **RP11-385D13.1** | **ULK3** | **PHLDB1** |
| **NCOA2** | **YY1** | **ATP6V0D1** |
| **POP5** | **CXXC5** | **UTRN** |
| **SAP130** | **SP1** | **PBX2** |
| **PABPC1** | **ARID5B** | **CNOT6L** |
| **CDYL** | **B4GALT3** | **EHD1** |
| **EI24** | **DUSP7** | **RUSC2** |
| **MEPCE** | **VSNL1** | **FLOT2** |
| **RBM47** | **KLHDC8B** | **SHH** |
| **THAP11** | **COL1A2** | **RPS6KA4** |
| **CDCA7** | **CCDC15** | **CLSTN1** |
| **HOOK1** | **FAM98A** | **DKK3** |
| **KLHDC3** | **DUSP3** | **SSBP3** |
| **SEC62** | **RBPJ** | **PHEX** |
| **NOL11** | **MFSD5** | **CHKA** |
| **SEC24A** | **MDM4** | **RELA** |
| **PIK3R1** | **IDH2** | **ITPR3** |
| **NEO1** | **ACVR1C** | **TMCO3** |
| **PCMTD2** | **ADRM1** | **CHMP1A** |
| **ARMCX3** | **SYNJ2** | **TANC2** |
| **TEAD1** | **FUT11** | **PSIP1** |
| **PNRC2** | **SMIM13** | **TMEM170A** |
| **SCYL2** | **PLXND1** | **FAM171A1** |
| **GEM** | **C19orf10** | **PTEN** |
| **ERP29** | **EGLN2** | **ZNF652** |
| **BAZ2A** | **ONECUT2** | **APBB2** |
| **SATB1** | **MGAT1** | **MTF2** |
| **SMARCC1** | **HIST1H2BL** | **TUBB4B** |
| **TGIF1** | **RBPMS** | **EWSR1** |
| **ASH2L** | **ANGPTL2** | **DUSP16** |
| **FKBP8** | **AKIRIN1** | **MARCHF6** |
| **MARCKSL1** | **CDC23** | **HPS5** |
| **TMEM185B** | **ZNF451** | **STX4** |
| **WDTC1** | **GNPTAB** | **SIX4** |
| **KDM5A** | **MAX** | **SPTBN1** |
| **SEPP1** | **NRARP** | **HNRNPUL1** |
| **CPEB2** | **TSPYL4** | **CCDC85C** |
| **TMCC1** | **MYH15** | **ATL3** |
| **RORA** | **ARRDC3** | **SLC35D3** |
| **SLC22A23** | **USP42** | **FAM160A2** |
| **CDK18** | **IKZF2** | **ACADM** |
| **MRPL43** | **FAS** | **BRAT1** |
| **PTPN11** | **PGRMC1** | **AP1B1** |
| **BUD31** | **ANAPC5** | **KIAA1841** |

HSP90B1
MAD2L2
ARPC1A
CDC42SE2
C2orf49
FARP2
TMEM39B
PHC2
DYNC2LI1
GTF2H2
ERAL1
NRP1
CRELD1
MPP6
PRKAR2A
PRKX
ZNF592
TUT1
BHLHE22
ZC3HAV1
TAF4B
SRRM2
SS18L1
DLGAP5
AGPAT6
KLHL15
FXR1
SLC30A9
PPIL4
PEX11B
SENP1
JAZF1
GNPNAT1
GALNT10
METTL9
EPHA2
MID1IP1
ASCC3
EIF5B
CERS5
LARP4
BACE2
PTPRJ
DIS3L
KITLG
MBNL2
NMT1
DOT1L
PRKAB2
MELK
PEX13
LAMC1
USP24
RAB40C
RPS6KA3
KCNJ2
NFE2L1
C14orf166
ERC1
ZNF414
MKI67
TSPAN14
RRP1B
TGFBR3
ASCC2
PCYT2
FAM160B1
MANEAL
TTLL11
PLIN2
SLC4A5
CD200R1
EP300
POM121
MIDN
PRDM2
HDLBP
SLC10A7
RNF165
CBLL1
BTG3
PAAF1
ARL5B
ZCCHC14
PIGT
GPCPD1
SAMD10
CTSA
TP53BP1
BCCIP
C9orf91
NFIC
SORD
RNF213
INF2
DCAF16
PPFIA1
RNF141
MAPK1
PNRC1
HSPA12B
AMER1
SLAIN2
TLN1
NUP205
RCC2
SPTSSB
AMOTL1
MICAL3
SRF
MAPK6
RRAGC
ADSL
CREBBP
C22orf29
CHD9
GREB1
ABL2
KDR
TRAF3
WRNIP1
RAD21
SERPINE1
NKIRAS2
ARHGEF4
SCARB2
AP3M1
CDK16
HEATR2
WDR77
PUS1
ADCY9
GOLGA3
BTG1
HOXA10

| | | |
|---|---|---|
| **TNFRSF21** | **TTLL4** | **NRN1** |
| **KBTBD11** | **RPUSD3** | **USP22** |
| **SYNE1** | **PLAG1** | **SERTAD2** |
| **SRPX2** | **S100PBP** | **GPR151** |
| **IRS1** | **PUM2** | **VAT1** |
| **WDFY3** | **ATXN7L3B** | **SEPHS2** |
| **LZTS2** | **PIAS4** | **CAPRIN2** |
| **GLCE** | **REV3L** | **UBE4B** |
| **RAC1** | **RSBN1L** | **NUCB1** |
| **NRP2** | **ZNF511** | **TMEM136** |
| **SNX2** | **OSBPL10** | **PCM1** |
| **GABPA** | **TTC22** | **ATP5G2** |
| **CD99L2** | **CELSR3** | **SERBP1** |
| **FAM120C** | **CLN6** | **TMUB1** |
| **TNFRSF25** | **RHBDF2** | **ENTPD4** |
| **SPRY4** | **FBXW2** | **SETD1A** |
| **CHRAC1** | **BRD3** | **KLF6** |
| **ANKRD13C** | **ARHGEF15** | **SLMO2** |
| **RANGAP1** | **TGM2** | **MYL12A** |
| **DUSP8** | **MYLK3** | **MRPL52** |
| **STC2** | **GRK6** | **PSMD6** |
| **MAPK3** | **GJC1** | **MAPKAPK2** |
| **PRKAA1** | **YIPF5** | **TMEM165** |
| **CAP2** | **CDK12** | **RBM4** |
| **UBXN2A** | **CNOT1** | **RHOB** |
| **CTNNAL1** | **ZNF281** | **NCKIPSD** |
| **ANKFY1** | **UBE2D3** | **CBX5** |
| **FAM208A** | **USP14** | **TNKS** |
| **SDAD1** | **CASP6** | **TEK** |
| **NOLC1** | **STARD13** | **NOP58** |
| **MESDC2** | **EIF4E2** | **JMY** |
| **ATP6V1A** | **COX10** | **HIST1H3J** |
| **CDC42** | **MPL** | **UQCR11** |
| **KMT2D** | **CREB3L2** | **HYOU1** |
| **SEPN1** | **DIABLO** | **GOLPH3** |
| **ADAMTS1** | **TSPYL1** | **TIE1** |
| **DNAJB14** | **CDH5** | **SLC1A5** |
| **FSCN1** | **CKAP4** | **EPHX2** |
| **PAK6** | **SHE** | **EIF1** |
| **KDM6A** | **POLQ** | **SLC39A1** |
| **UBE2K** | **GBE1** | **GLYR1** |
| **NUP50** | **ATP8B3** | **TONSL** |
| **KDM6B** | **PARP1** | **MAML1** |
| **BCORL1** | **CTSB** | **MGAT4B** |
| **PEA15** | **UBE2G2** | **RPL15** |

IPO5
IPO4
LDHB
SESN1
AIF1L
TSEN34
SESN2
VPS52
SDE2
HSPA4L
YRDC
POLR3C
KIF11
CS
FRYL
C2CD4A
COMMD2
DSC3
TIA1
PIK3C2A
CIPC
C16orf72
KCNJ6
MRGBP
RPL19
TSC22D2
TXLNG
KIAA0247
TMEM2
LYPLA2
ADCK3
DNAJC12
MIOS
SNRNP40
ZBTB21
DHX15
BAG2
DYNLL2
AP2A1
CALD1
ADNP
RPS27L
C6orf120
NF1
ZNF430
H3F3B
FOXP2
ZFYVE20
CLPTM1
GRAMD1A
RIIAD1
STAMBP
RGS16
RNF167
TRMT112
SOD2
MBD4
PAPD4
IDE
CYTH1
PCDH7
ZFR
VPS33A
TRAPPC10
PYGB
ACTL6A
SMIM3
NMNAT1
CDC27
KPNA6
SLC1A4
KAT5
BAHD1
TLCD1
GALNT2
KIAA0355
GPSM2
PHLDA3
PLXNC1
ABI1
SPINT1
RPL22L1
SLC6A9
ATP6V1C1
CYP1B1
SCD
SAMD4B
VPS53
LRGUK
TFIP11
SNRNP200
PTGES3
SGK1
PRRC2A
ATP5SL
RAPH1
SYNPO2
JAK1
RAB5B
DEF8
MAF1
CD63
AKAP11
MAP7D1
ELOVL4
ICT1
RNF126
DYNLRB1
SLC39A14
CMPK1
MTDH
TOMM6
LPGAT1
PRR11
FANCI
ABL1
ICA1L
RABL6
PCMT1
VKORC1L1
GNB5
BAZ1B
SMG1
BMPR2
RPS29
RTN3
TMBIM6
PGAM5
GINM1
EIF4G2
ECE1
KIF5B
MTF1
NCOA1
NOA1

STRN
HMG20A
CDK13
KIAA0100
UBAP2
NIFK
CACYBP
RANBP2
SUV420H1
DTNA
ZFP1
SPSB1
HIVEP1
SP1
ARID5B
B4GALT3
DUSP7
ABCB7
COL1A2
JAG1
FAM98A
MDM4
OGFRL1
CSNK1G1
ACVR1C
KRT80
SLC39A6
FUT11
SMIM13
PLXND1
C19orf10
ONECUT2
KLC1
TWSG1
AGAP5
AKIRIN1
NPTN
TNRC18
ZNF451
GNPTAB
MTERFD3
TLE3
AP3D1
CNOT2
RASSF3
C14orf28
DNA2
SUV420H2
NGRN
AURKA
ZNF644
ZNF512B
THOC2
GPN2
NOC3L
NUP43
PLEKHA1
SGOL1
KMT2A
ROGDI
RARG
RPN2
SCAF1
AKR1A1
VEZF1
MYO5C
TAF5L
MED19
PXT1
GRPEL1
TMEM8A
HOXD10
CHD5
MEX3D
TPP1
LASP1
RLF
FUT10
IGSF3
ZNF585A
FBXO22
LMNA
NDC1
DRG2
CANT1
FN1
TSC22D4
ACSL1
NWD1
MYCN
FBXW9
HLA-A
LIN7C
SIN3A
PLS3
MLLT10
ANKMY2
CDC42BPB
ZNF689
SIDT1
DDX49
C21orf33
SARS
PRKACB
XIAP
EGLN1
MAP1B
APH1B
CTGF
MCF2L
DCHS1
QRICH1
GSTCD
COL4A1
PFKFB3
UBR4
ZKSCAN1
SENP5
NACC2
TTC9
IDH3G
ANAPC16
TRIM28
WHAMM
ASAP2
ATP5O
TOR3A
DSP
RPL10
RIN3
KDM5B
GSPT1
CYB561D2
NDEL1
ADCY6

TAF7
USP42
NFE2L2
EPC2
ANAPC5
GFM1
MBNL2
STX6
DOT1L
SLC20A1
PPP6R1
NDUFA8
LAMC1
PRICKLE2
USP24
RAI14
RPS6KA3
NFE2L1
NEDD4
FOXJ3
ERC1
CDK19
ATP5E
MKI67
SCOC
IGF1
ELP6
TGFBR3
CNTLN
UBE2F
LAMP2
PRKAR1A
SUMO3
FAM160B1
UBL4A
KIF26A
MIDN
AMOTL2
BRE
PRDM2
ZNF714
DDAH1
SLC10A7
FCGRT
PTPRA
NECAB3
UBN2
MYO1E
SLC35E3
ATP7B
OR2AG1
PXDN
RAD54L2
PKN2
CALR
AP3S2
TEAD4
CDKN1A
PYGO1
PCIF1
UHMK1
LHFP
DCAF15
BIRC3
RHOF
ATG12
CCNJ
IRAK2
C3orf38
NANS
CHTF8
HIAT1
TNRC6A
TRAPPC1
SLC2A1
MAP3K2
RBM10
ARF6
RSPRY1
USP9X
SAMD5
TMEM201
WDR43
NFIA
GFM2
MON2
ZFHX4
SRSF11
PAN3
INPP5A
RMND5A
ZCCHC3
RBM15B
HIST1H3B
HIST1H3F
CYFIP1
OTUD7B
RP5-850E9.3
HIST1H3I
HIST1H3E
HIST1H4L
SNX12
GPR75-ASB3
HIST1H3G
SYNRG
LIX1L
ZFP91
CTDSP2
DDX52
HIST1H4A
BRD4

AHSA2
BTG3
PPARG
MYT1L
POU2F2
PAAF1
MGEA5
DENND2D
ARL5B
CASP8
NOTCH3
TTLL4
PLAG1
PUM2
NFIB
THUMPD1
C2orf47
TWISTNB
RASEF
C12orf57
CHML
OSBPL10
STON2
ATF7IP
IGFBP2
MEDAG
PLEKHB1
COL11A1
GJC1
LY6G5B
POMK
CNOT1
ZNF281
EFNB1
ATG3
CEP135
UBE2D3
ZZEF1
STARD13
PHF12
TSHZ1
PPARGC1A
TSPYL1
SCAF11
CBY1
ANKHD1-EIF4EBP3
AVL9
UBR3
PLD3
TNFAIP1
RFC5
ATG9A
POLL
MON1B
CARM1
SCYL1
EIF4EBP2
MLXIP
SUV39H2
RRM2
VGLL4
CAD
COPA
KLHL12
FAT1
C17orf51
EIF1AD
DONSON
ZNF555
GTPBP4
ZNF473
NUDT16
PFKFB2
PRDX6
PPM1H
NUP155
TPT1
WDR24
PARD3
TTI1
GORASP2
HMGCS1
BCOR
GAREM
HOXC6
NCKAP5L
MMRN1
BTBD7
HNRNPL
SLC25A27

CYB561
CDH5
CKAP4
PARP1
UBE2G2
H3F3B
GPR160
SEC24B
MPHOSPH9
CAPN1
SLC4A8
CLIP2
CNN3
TMEM150C
CD99
GAPVD1
G3BP2
RBBP7
B4GALNT1
TP53I11
ZFR
TTC39A
SUZ12
TRAPPC10
HNRNPH1
OAZ2
ABCA5
CCDC88C
FAM210A
RC3H1
SPX
CDC27
KPNA6
IGFBP3
FASTKD5
CAPN15
MYCT1
ZNF81
GALNT2
LONRF2
PJA2
ZBTB33
FAM193B
ABI1
VPS39
ERGIC1
PPIG
ATP2A2
CCNJL
AFAP1L1
MPZL3
RASSF8
EDN1
AHNAK
YTHDF3
PRKDC
CNBP
ZBTB34
SH2B1
SMAD7
PAFAH1B2
KIF1C
SRGN
E2F4
ENTHD2
BVES
TNFSF9
RPL9
GLRX5
UBE2W
C1GALT1
RBM4B
PDE7A
SPHK1
DUSP5
ZBTB32
PPIF
TRRAP
YIPF1
VPS25
ZFP36L2
ARHGEF7
PIEZO1
DLAT
AURKB
USP11
PSMA4
GPR157
ABCC5
ANAPC13

DDA1
ABAT
NCDN
ATP6V1C1
NGLY1
CYP1B1
LAPTM4B
KLHL9
MARCHF5
UBE2G1
SNRPB
TFIP11
RCBTB1
SREK1IP1
RLIM
IL6ST
SEMA7A
ZNF462
THOC2
NAPB
ENPP1
FAM102A
LEF1
OCRL
MRPL28
KMT2A
RPN2
EPS8
VEZF1
UPF2
BAMBI
CLK2
PRMT5
PACSIN2
ZER1
ALDH9A1
ZNF682
GPI
TPP1
LIMK1
LASP1
RLF
MECOM
FOXO3
DHX9
PRKD2
ZCCHC10
CTDP1
ARHGEF17
RNF6
FOSL1
SNX25
WIPI2
KLHL24
ALDOA
CTDSPL2
PTHLH
LEPROT
CLDN12
UHRF1BP1
SRSF10
KREMEN1
DPH2
ZNF449
MOB1A
DOCK7
PRDM4
VCPIP1
CDC42SE1
KIRREL
GFOD2
IGDCC4
SLC30A3
TADA1
JAGN1
PDZD8
LUC7L
PFKL
SUCO
UTRN
MAP2K4
PBX2
C9orf64
EHD1
KMT2E
PTPRB
USP3
CHKA
VAV3
ITPR3

ANKRD28
DERL1
IFNG
FBXO22
LMNA
SLC7A2
CANT1
TULP4
FN1
FAM136A
STARD7
IFFO1
SEPT2
IPO8
RNASET2
ERI3
ANXA11
CTCF
GFRA1
RRP12
PXDN
RAB22A
TMEM218
TXN2
RAD54L2
SEMA4D
LRIF1
TNC
ROBO1
AKAP12
CALR
ELOF1
EIF4G3
TMEM30A
RBM12B
CDKN1A
ERH
PCIF1
STMN1
UHMK1
RALB
POFUT1
NAPEPLD
SYNE2
CPT1B
DNAJA2
CEP290
SPHK2
RFWD3
XPO1
EPHA4
TRIM71
AP2S1
NFATC3
MTF2
FGFR1
FUCA1
HNRNPUL1
AGL
LIN54
TRIM25
PHF21A
LATS1
SLC35F6
FOXO1
AP1B1
SALL4
ZSWIM8
SOX2
BLOC1S4
RNF213
EMC1
CTPS1
BBX
INF2
CAB39
TAF5
PCYOX1
ATP13A3
TRIB1
MAPK1
MFI2
MVK
PNRC1
TTLL9
MAP3K7
FNDC3A
NUPL1
RAD54L
DDX26B

**C3orf38**
**GTPBP10**
**CDC42EP3**
**GPT2**
**DOCK9**
**TNRC6A**
**MDN1**
**ZNF662**
**AKR7A2**
**USP9X**
**IRX3**
**MBNL1**
**ISL1**
**WDR4**
**TMEM201**
**EIF6**
**TBP**
**MYBBP1A**
**COBLL1**
**ATP2B4**
**LTBP1**
**NOL10**
**PAN3**
**ANKHD1-EIF4EBP3**
**MKNK1**
**ZMAT2**
**MCAM**
**SEMA6D**
**KIF21A**
**PRKD3**
**ABCA12**
**ZNF579**
**CHAMP1**
**HSD17B12**
**KIAA0586**
**ZBTB18**
**REEP1**
**PRKCH**
**PPA1**
**HHLA3**
**MON1B**
**CARM1**
**TUBA4A**
**MLXIP**
**KIFC3**
**KANK3**
**LRRC41**
**TLN1**
**HIST1H2AC**
**ITPR2**
**SRF**
**MAPK6**
**MMP14**
**MED6**
**CREBBP**
**C22orf29**
**C9orf37**
**FBXW11**
**GATAD2A**
**RHOBTB2**
**CHD9**
**SESTD1**
**ZNF146**
**RNASEH2B**
**MAFB**
**ABL2**
**TMPPE**
**GNG5**
**TET2**
**SLC43A3**
**PCGF3**
**ZNF33A**
**ESRP1**
**AP3M1**
**DNAJC27**
**CDK16**
**APLN**
**TBPL1**
**PIP4K2C**
**MGAT4A**
**PUS1**
**ADCY9**
**PSMC6**
**HELLS**
**GOLGA3**
**KATNAL1**
**SERTAD2**
**ZSCAN9**
**RNF26**
**DLGAP4**

| | |
|---|---|
| PDZD11 | CAPRIN2 |
| RRM2 | VRK1 |
| SMG7 | TLDC1 |
| VGLL4 | UBE4B |
| CKAP2 | PCM1 |
| SESN3 | PATL1 |
| LDLR | ZNF567 |
| FAT1 | TRIM32 |
| TSEN54 | MTCL1 |
| C17orf51 | ALG3 |
| HNRNPA2B1 | SETD1A |
| TEX15 | FNBP4 |
| FOCAD | C20orf194 |
| HRSP12 | GLB1 |
| PDPK1 | ARF4 |
| NOP9 | ZNRF3 |
| PPM1H | ZNF623 |
| NUP155 | MYBL2 |
| TPT1 | OS9 |
| TRIM44 | TNPO3 |
| DKK1 | ZNF17 |
| LRP3 | KIF5C |
| ZBTB1 | CDC42EP2 |
| POLA1 | PLBD2 |
| FIP1L1 | GPR161 |
| HMGCS1 | STK40 |
| FAM47C | SLC12A7 |
| BNIP3L | RHOB |
| BCOR | SIM2 |
| LIMA1 | BRWD1 |
| PNMA1 | CBX5 |
| TUB | STARD4 |
| CCNI | TEK |
| HOXC6 | JUP |
| BTBD7 | JMY |
| OPA1 | PIAS1 |
| ERalpha(hsa) | DMKN |
| HNRNPL | 9-Sep |
| PHF13 | WDR37 |
| PPIG | DNASE1 |
| MAP1A | LEFTY2 |
| ATP2A2 | SNRPC |
| NAV3 | BAZ2B |
| RPL23A | DUSP1 |
| AFAP1L1 | GATA6 |

CCNK
ID3
RASSF8
HIPK1
AHNAK
TINAGL1
PRKDC
RINT1
SASH1
ZBTB34
SH2B1
AP1G1
SMAD7
COL14A1
KIF1C
EYA1
E2F4
KCNB2
TNFSF9
MEF2C
GLRX5
PLEKHH1
UBE2W
ZNF726
ABCA1
TMEM127
RAN
NDUFV1
MKNK2
PSRC1
FANCA
ZIC5
DCLRE1B
STAT6
STAB1
PDE7A
PALM2
PPIF
ADD1
GDI2
JADE1
MTOR
TRRAP
ARPC4
PRR13
FOXF2
ARHGAP28
BZW2
TIE1
SECISBP2L
TGFBR2
GPRIN3
EIF1
C3orf52
SLC52A3
BICC1
MAML1
PON2
USP10
TMEM60
SNRNP200
COPS6
PTGES3
PRRC2A
MBOAT2
FADS3
GCNT2
JAK1
EXO1
CABIN1
ZNF142
MDM2
SH3TC1
WNT9A
MTDH
GPR137C
LPGAT1
ZNF629
ABL1
NOP16
PLCB4
AP5M1
KIAA1715
NIPA1
VPS54
BMP4
SNX24
EHMT1
COIL
SVIL

VPS25
ZFP36L2
PPM1G
PLEKHA6
PHF5A
ZNF106
NCKAP1
MAGT1
GATAD1
DYRK1A
CTDP1
CMTM6
ABHD17C
FOSL1
MOAP1
WIPI2
C1orf115
ALDOA
BCAR3
NR1D2
SLC35B4
PTAR1
DENND2C
EXOC8
SLC30A2
METTL3
MOB1A
ZNF574
NFASC
VCPIP1
ITGA6
ZFY
SGPP1
PDZD8
ATP1A1
PFKL
SUCO
ST20
UBE2B
UTRN
MAP2K4
XPC
EHD1
RUSC2
AKAP6
BAZ1B
SMG1
MCM3
STK24
BMPR2
RPS29
TMBIM6
ATF4
H1FX
NNT
ZNF141
BCL2L11
RHOG
ECE1
MED8
SYT16
TBRG1
NR2F2
ZMYND8
HOXC8
SLC27A4
SOBP
DAXX
LIN7C
ABCF1
SIN3A
MLLT10
ANKMY2
ZNF689
WBP11
ARRB1
PAX3
ZNF695
MAP1B
FLCN
EPB41
SOX5
EIF3C
MICAL2
DHX33
COL4A1
PFKFB3
SENP5
CDC25A
FNIP2

| | |
|---|---|
| SHH | TRIM28 |
| ZNF518A | PEX1 |
| RYBP | SYNCRIP |
| RND3 | BIRC2 |
| PTRH2 | RIMKLA |
| TMCO3 | KCNH2 |
| COL5A3 | ATP5O |
| NME1 | DSP |
| ZNF618 | INO80D |
| GTF2A1 | IL6R |
| TANC2 | LRCH4 |
| XPO1 | UBOX5 |
| PHIP | C9orf142 |
| FAM171A1 | GSPT1 |
| FBXO33 | COA7 |
| SCNN1A | SMCR8 |
| ZNF408 | UTP11L |
| CPEB3 | DSEL |
| PCF11 | EFNA1 |
| ZC3H4 | TUSC2 |
| MTF2 | C7orf55-LUC7L2 |
| TUBB4B | ZCCHC3 |
| DUSP16 | LUC7L2 |
| MARCHF6 | KMT2B |
| DCTN6 | PIGW |
| HPS5 | GGNBP2 |
| PALLD | SOCS7 |
| SIX4 | MT-CO1 |
| FUCA1 | MT-ATP8 |
| SPTBN1 | MT-ND5 |
| CHD6 | OTUD7B |
| ATL3 | BOP1 |
| LIN54 | LHX1 |
| TRIM25 | RP11-201K10.3 |
| CD55 | NR4A1 |
| FOXO1 | ZNF25 |
| AP1B1 | MARCKS |
| IRAK1 | SERPINA3 |
| CAP1 | GPR75-ASB3 |
| RNF213 | ARHGAP23 |
| ADAMTS9 | ZNF792 |
| BBX | H2BFS |
| CAB39 | VAMP3 |
| PCYOX1 | RPS6KA2 |
| ATP13A3 | TMEM189-UBE2V1 |

**PSMF1**
**SPATS2**
**RNF141**
**DYNC2H1**
**TRIB1**
**NSFL1C**
**QPCTL**
**MAPK1**
**MAP3K7**
**FNDC3A**
**BRI3**
**SDCBP**
**SLAIN2**
**PAX9**
**NUPL1**
**MTMR3**
**RNF168**
**RCC2**
**METTL7A**
**AMOTL1**
**MAPK6**
**MED6**
**PTGFRN**
**CREBBP**
**FBXW11**
**GATAD2A**
**TMEM231**
**PPP2R1B**
**SESTD1**
**C9orf114**
**PRDX3**
**ABL2**
**CSF1**
**ZHX1**
**IL10**
**RAD21**
**HECTD4**
**HMMR**
**AP3M1**
**CIRBP**
**DNAJC27**
**KMT2C**
**HEATR2**
**APLN**
**TBPL1**
**ZFP92**
**HIST1H4D**
**SARS2**
**ZNF280B**
**PSMB3**
**MUC16**
**BRD4**

**GRB2**
**MGAT4A**
**FAM84B**
**CALCRL**
**BTG1**
**HOXA10**
**ITSN1**
**KIAA0895**
**USP22**
**EIF2B4**
**RNF26**
**C17orf70**
**DLL4**
**SLFN5**
**SMAD1**
**UBE4B**
**VBP1**
**RAB10**
**PCM1**
**PARD6B**
**SERBP1**
**TMUB1**
**CPNE8**
**TRIM32**
**DNMBP**
**MTCL1**
**C1GALT1C1**
**LYSMD3**
**TAF3**
**KLF6**
**SRP14**
**NGDN**
**KIAA1279**
**MYBL2**
**ENAH**
**ARHGEF26**
**ITPK1**
**RAB20**
**ZBTB25**
**GPR107**
**DNAJC2**
**MAPKAPK2**
**GPR176**
**ZNF675**
**RNF139**

**KNTC1**
**SLC12A7**
**KPNB1**
**RHOB**
**C19orf24**
**BRWD1**
**CBX5**
**TTPAL**
**CDK10**
**JMY**
**MXRA7**
**TBL1XR1**
**SEPT9**
**WDR37**
**SETX**
**GALNT5**
**C14orf132**
**LETM1**
**GATA6**
**RNASE1**
**ATRN**
**IREB2**
**SPIDR**
**CCP110**
**GPS1**
**SECISBP2L**
**TGFBR2**
**SLC1A5**
**MEF2A**
**EIF1**
**C3orf52**
**BICC1**
**RAB14**
**ORC5**
**PON2**
**MTSS1L**
**TMEM60**
**ATP2C2**
**PLA2G15**
**SGK1**
**PRRC2A**
**ZBTB47**
**ARHGAP5**
**ATP5SL**
**RAPH1**

**KHDRBS1**
**PLEKHG1**
**JAK1**
**RPA2**
**RAB5B**
**CABIN1**
**MAF1**
**CLDN3**
**AKAP11**
**GLYCTK**
**SLC46A3**
**MDM2**
**TTC5**
**SLC18B1**
**PGK1**
**ARMCX2**
**WNT9A**
**MTDH**
**TOMM6**
**SYTL5**
**NIN**
**ZNF669**
**TNRC6C**
**SLC35E2B**
**N4BP1**
**SCML1**
**ABL1**
**KIAA1429**
**EHMT1**
**SVIL**
**STK24**
**BMPR2**
**KIAA1551**
**PNISR**
**TMBIM6**
**ARHGAP12**
**TOR1AIP1**
**ZW10**
**EIF4G2**
**BCL2L11**
**ECE1**
**NR2F2**
**ZMYND8**
**MAP3K5**
**ZMYM3**

**KIF5B**
**MTF1**
**ILK**
**ZFP91-CNTF**
**LIN7C**
**SP6**
**SIN3A**
**CDC42BPB**
**PDGFRB**
**HOXA13**
**RPS6KB1**
**ARRB1**
**ERLEC1**
**CD44**
**ARHGEF28**
**KCTD20**
**EGLN1**
**MAP1B**
**FLCN**
**EPB41**
**MIER3**
**ADH5**
**DCHS1**
**DHX33**
**SIAH2**
**KIAA1430**
**DGKH**
**TRIM26**
**SLC30A1**
**UBR4**
**ZKSCAN1**
**SENP5**
**NACC2**
**CDC25A**
**C2orf69**
**BNIP2**
**FNIP2**
**VDAC3**
**TRIM28**
**YY1AP1**
**RXRA**
**RIMKLA**
**SPRY2**
**MEX3C**
**TRAPPC8**

**EIF3I**
**LRFN5**
**INO80D**
**PANK1**
**SEC11A**
**p53(hsa)**
**KDELR2**
**NEURL4**
**B4GALT7**
**KDM5B**
**GSPT1**
**NDEL1**
**COA7**
**ADCY6**
**SMCR8**
**UTP11L**
**TUSC2**
**ELF1**
**RMND5A**
**OR7C2**
**ZCCHC3**
**TXNIP**
**KMT2B**
**HIST1H2AH**
**HIST1H4E**
**PIGW**
**GGNBP2**
**PILRB**
**PIP4K2B**
**GTF2H5**
**ZNF8**
**NUDT3**
**C5orf22**
**HIST1H2BJ**
**SNX12**
**MARCKS**
**GPR75-ASB3**
**MYO19**
**VAMP3**
**RPS6KA2**
**ACACA**
**ZFP91**
**CTDSP2**
**CPNE1**
**PECAM1**

**BRD4**
**ZNF28**
**F11R**

| miR-181a-5p | miR25-3p | miR31-5p | miR28-3p |
|---|---|---|---|
| BMI1 | CALU | TBC1D20 | BMI1 |
| FAM47B | ZKSCAN5 | CAMTA1 | VAPB |
| CCNT2 | POLR2B | ATF6 | ESR1 |
| POLR2B | RTN4 | TMEM120A | BRK1 |
| RTN4 | ABCD4 | ANP32E | SLC25A44 |
| SLK | RPRD2 | ACTB | CECR1 |
| WWC2 | ZNF503 | CCNG1 | HOXB7 |
| E2F7 | PELI1 | PCNXL4 | ACTB |
| PELI1 | DDI2 | JARID2 | EAF1 |
| DDI2 | CAMTA1 | DFNA5 | SLC38A1 |
| RAB2B | RPA1 | CRTC3 | SNAP23 |
| TWF1 | IRS2 | CCNB2 | IQCJ-SCHIP1 |
| ZNF419 | RBM14 | IL1RAP | PPP1R2 |
| DTD2 | RNF11 | URGCP | FOS |
| IRS2 | BRAF | FARP1 | ZEB2 |
| FHL2 | FHL2 | ITPRIP | TCF3 |
| SFT2D1 | ANP32E | SCAMP2 | MTA3 |
| ANP32E | TNFRSF11B | SPARC | BICD2 |
| ZNF544 | ACTB | GUCD1 | ADIPOR1 |
| ACTB | CCNG1 | STAT3 | RNF44 |
| CCNG1 | SLC38A1 | TAGLN | CREBL2 |
| EAF1 | PHLPP2 | CCAR2 | WAPAL |
| EIF3L | MTO1 | C2CD3 | PKIA |
| SLC38A1 | TMEM50A | RAB3GAP2 | MFAP3 |
| RP6-24A23.6 | BRI3BP | ZSCAN25 | MYBL1 |
| DUSP4 | NRCAM | TBC1D13 | MGAT2 |
| PRLR | NEK7 | NPDC1 | SPARC |
| ZNF239 | PAQR3 | YIPF3 | UGCG |
| NEK7 | SAP18 | NFKB1 | SAR1A |
| MSI2 | SERTAD3 | FAM189B | B3GNT5 |
| SEC16A | PPP1R37 | OAZ1 | RREB1 |
| DCP1A | MLK4 | MET | FERMT2 |
| RAB3GAP1 | ZMAT3 | ATP6V0E1 | CDH2 |
| FOS | JARID2 | GRB10 | G3BP1 |
| MLK4 | EDRF1 | CLEC14A | ZNF638 |
| JARID2 | BICD2 | DCTD | DIRAS1 |
| NETO2 | GSK3B | IL1R1 | MORF4L1 |
| HIPK2 | RNF44 | RNF38 | HMG2L1(hsa) |
| TMED8 | ALKBH3 | HTRA1 | GGCT |
| GSKIP | ASB7 | WDR47 | CHST3 |
| C5orf30 | FAM120A | THEM6 | RFESD |
| SOX4 | C5orf30 | CDK4 | FAM53B |
| ARL4A | SOX4 | CCDC92 | LRP6 |

| | | | |
|---|---|---|---|
| FBXL3 | MCOLN2 | EIF2AK3 | IPO11 |
| FBXO11 | UIMC1 | BRD2 | SMC1A |
| PKIA | RBL2 | ASCC1 | MEGF9 |
| RBL2 | OTUD1 | DICER1 | ZNF616 |
| OTUD1 | DAZAP2 | TMEM205 | ADAMTS19 |
| TMPRSS11A | NDUFB10 | CBX2 | DDX3Y |
| MFAP3 | SCAMP2 | SDC1 | ZBTB44 |
| NUMB | RBM27 | MLF2 | CDS2 |
| KSR2 | PEAK1 | RP4-758J18.2 | ITGB8 |
| UBXN7 | ANKRD13B | EMC10 | FAM126B |
| POC1B-GALNT4 | BRMS1L | POU4F1 | ATF2 |
| SLMAP | FREM2 | TMEM194A | KCTD12 |
| SCAMP2 | TUBGCP3 | ATXN2L | POU3F2 |
| MYBL1 | PCNA | C12orf76 | USP36 |
| RBM27 | SPARC | CD59 | HNRNPA1 |
| PEAK1 | ZSCAN26 | THUMPD3 | DICER1 |
| SIN3B | ETAA1 | TET1 | SDC1 |
| BRMS1L | NAGA | YWHAE | ZBTB10 |
| C18orf21 | CLPP | DENND4B | LRSAM1 |
| MGAT2 | UTP15 | CALM1 | GAX(hsa) |
| UGCG | PTPRK | ZNFX1 | MTERFD2 |
| STAT3 | GALNT7 | RGS4 | MMGT1 |
| CEBPG | UBE2Z | VPS13D | PRKAA2 |
| SFXN4 | GLG1 | HIST1H2BK | CRKL |
| TNPO1 | TVP23B | SIK1 | DGKE |
| UBE2Z | HSPA1A | STOM | SLC7A6 |
| SWAP70 | EID2B | TMEM167A | YWHAE |
| WNT16 | TGFBR1 | DAB2 | TSC22D3 |
| G3BP1 | CHMP7 | AGO3 | ELK4 |
| ZNF638 | PDE4DIP | DHCR24 | TMEM59 |
| PDCD4 | TCEB3 | LENG8 | CYR61 |
| ZNF568 | COX8A | SMIM8 | TMED7-TICAM2 |
| TM4SF1 | PPP2R5E | MEIS2 | ATP5F1 |
| CDON | PLP2 | FAT3 | GNS |
| ADAMTS5 | FEM1B | ANKRD52 | CLDN11 |
| PDE4DIP | RAP2B | REST | DCTN5 |
| TCEB3 | TROVE2 | USP31 | TCF4 |
| GGCT | YWHAH | PLAGL2 | GTF2H1 |
| METTL2A | SPRYD4 | DIDO1 | GOSR1 |
| PPP2R5E | CNOT11 | NF2 | TMED7 |
| UNKL | ZNF616 | PIK3R4 | PPAT |
| FEM1B | ZDHHC21 | DDX56 | THBS1 |
| LRP6 | ADAMTS19 | UBQLN1 | UBTD2 |
| OAZ1 | GRAMD1B | PAFAH1B1 | ITGA8 |
| EIF3F | EPCAM | REEP3 | CMBL |

| | | | |
|---|---|---|---|
| SMC1A | UBR1 | PPP1CC | AGO3 |
| MET | MED13L | GNA13 | DBN1 |
| ATP6V0E1 | RPL6 | FAM35A | ETV3 |
| UTP14A | CIC | ZSCAN20 | ZNF681 |
| TROVE2 | FBXL5 | TOMM20 | CLCN3 |
| ZBTB11 | RPN1 | NUP210 | TMEM248 |
| WASF1 | ZFAND5 | SLC25A32 | MARVELD1 |
| SENP3 | DDX3Y | ZNF16 | SSU72 |
| YWHAH | PTGER4 | PDE3A | HOOK3 |
| CST5 | PLAA | SPRED1 | B4GALT5 |
| ZNF273 | NDFIP2 | PTPN14 | ZNF131 |
| BLCAP | GATM | MAP4 | RBM28 |
| HLCS | RNF38 | KIF2A | UTP6 |
| UBP1 | MRS2 | FZD6 | ZNF275 |
| USP53 | ZBTB38 | PPP3R1 | PITPNA |
| CXorf23 | ZNF827 | ASXL1 | SGPL1 |
| RRAGD | RB1CC1 | WNK1 | YWHAG |
| ITGB1 | FAM126B | NME6 | SEC23A |
| ANO6 | RHOBTB3 | PPRC1 | CCND2 |
| C5 | RPL7A | SETD1B | PRKCDBP |
| ZNF32 | CDK4 | SPATA13 | ANXA1 |
| CCNB1 | LAMB1 | SRCAP | DKK2 |
| HNRNPR | BSDC1 | TMEM138 | FAM35A |
| SOCS4 | FAM135A | TMEM64 | ETS2 |
| GNA12 | ATF2 | AKAP1 | RC3H2 |
| CEP97 | NUCKS1 | AFAP1 | LMBR1 |
| DDX3Y | C12orf49 | UBA52 | TRIM35 |
| YAP1 | RIOK3 | ECHDC1 | MYOF |
| ADRBK1 | CTNND1 | SRGAP1 | ABCG2 |
| HMGB3 | HDGFRP3 | NDST2 | NKTR |
| WASL | SMAD2 | RFX1 | ARNT |
| DDHD1 | SH3PXD2A | DCBLD2 | ETS1 |
| ZNF621 | ZNF791 | APH1A | DPP8 |
| ELF2 | MRPS35 | SVEP1 | TCF12 |
| SPIRE1 | BRD2 | AFF4 | KCNAB3 |
| AREL1 | USP36 | HNRNPA0 | FRS2 |
| SH3BGRL | USP38 | CSDE1 | STX3 |
| C1orf43 | TDRD1 | AXIN1 | MEX3A |
| TRIM23 | SCRN1 | WDR20 | LTBP2 |
| ZIC2 | DICER1 | MRPL48 | FICD |
| KAT6A | TMEM205 | MCU | DNAJC10 |
| ANGEL2 | PHB2 | ATP1B1 | ORMDL3 |
| NUDT12 | RBM12 | C19orf12 | GPX1 |
| DNAJC6 | MAST4 | SLC25A5 | GSE1 |

| | | | |
|---|---|---|---|
| PGAP1 | SLC37A3 | THRA | PARK7 |
| ZNF594 | HCN2 | SETD5 | RCSD1 |
| L1CAM | LRCH1 | SFT2D2 | SETD1B |
| FUS | BCAT2 | WBSCR22 | IRGQ |
| PLEC | NRAS | TMEM109 | TRPS1 |
| PGR | DNAJC3 | CD164 | RAF1 |
| TCERG1 | IP6K1 | DSE | FRK |
| FAM126B | CRKL | BTBD3 | ATP7A |
| SMS | MTHFD2 | P4HB | NHLRC3 |
| FAM135A | CD59 | ARF1 | TMEM64 |
| USP9Y | ATXN1L | CLTC | TIAM2 |
| NUCKS1 | DGKE | PTMS | NUP93 |
| GPR137B | REXO1 | ZNF225 | CCT5 |
| METAP1 | PIGX | GLS | GMNN |
| C12orf49 | ELK4 | VCL | C18orf32 |
| MFSD6 | CLIC4 | NOP56 | C20orf166 |
| CLASP2 | KBTBD2 | TIMP2 | CPSF6 |
| SHROOM3 | CHD1 | HN1L | MPZL1 |
| ZNF12 | KIF1B | SF1 | CDKN1B |
| RAD23B | HIST1H2AE | ENSA | TMED5 |
| CHCHD1 | ATP5F1 | MIEF1 | PTPRM |
| EIF2AK3 | CLDN11 | ANXA2 | MYH9 |
| H2AFJ | MBTPS1 | NGFRAP1 | DSTYK |
| GARS | BET1L | BANF1 | BCAR1 |
| STAM | EXOC5 | RAB1A | SNTB2 |
| USP38 | DENND4B | ISCU | PHACTR4 |
| ELAVL2 | ANKHD1 | FOXC1 | NUP88 |
| OFCC1 | ZMYM4 | PRSS23 | HIPK3 |
| SH2B3 | TCF4 | RCAN2 | PPP6C |
| DICER1 | LGALS3BP | GPX7 | CAND1 |
| ARHGAP11A | INHBB | WDR6 | DDX20 |
| IQCG | YOD1 | CNOT8 | UBE4A |
| PPP1R9A | WARS | HEG1 | HNRNPA0 |
| FBXO21 | PPAT | HOMER2 | SLC7A11 |
| PAM | ENTPD5 | SLC25A40 | CDKN2B |
| CAMK4 | RRN3 | IPO7 | TIAM1 |
| SLC37A3 | HMGCR | IGF2BP1 | HMGB2 |
| PTBP2 | FBXW7 | SLC35A4 | CLN8 |
| MGAT5 | GUCY1A3 | OXSR1 | TNKS2 |
| ETV6 | SSH1 | S100A11 | ZNF292 |
| NRAS | COL12A1 | HERC1 | WEE1 |
| IKBIP | WSCD1 | NOL9 | CDK1 |
| SIK3 | ARRDC4 | CALM2 | SETD5 |
| DIEXF | SIK1 | PPIA | SFT2D2 |
| TCTN3 | RUNX1 | FOXRED2 | CD164 |

| | | | |
|---|---|---|---|
| CRKL | SLC33A1 | APP | LSM14A |
| LMO2 | HSPA4 | MARCH7 | FMR1 |
| TMEM194A | HMGA2 | TAOK1 | DDX17 |
| NAT8L | VANGL1 | SIK2 | ZC3H15 |
| CD59 | TOR1AIP2 | SLC35A2 | KARS |
| TSPAN5 | GTF3C4 | AMMECR1L | ODC1 |
| KLF15 | UBTD2 | SIKE1 | KPNA2 |
| CNOT4 | ITGA8 | SPIN4 | ZFHX3 |
| VTA1 | PIK3CB | FAT4 | ARF1 |
| PPP2R3A | LGR4 | DPM2 | GEN1 |
| ZNF124 | CMBL | NSD1 | TMOD3 |
| MRPS23 | MMP15 | UNG | ZNF226 |
| CRIM1 | PIP5K1C | INSIG1 | SQSTM1 |
| BRIX1 | LENG8 | RPS6KA5 | GNB1 |
| DUSP11 | KIF20A | CBFB | MAP3K4 |
| PRR3 | ATMIN | CNP | WWTR1 |
| AP3S1 | CDKAL1 | TBC1D16 | ASPH |
| KBTBD2 | FAM47E-STBD1 | FYTTD1 | RHOA |
| CYR61 | RECQL | RICTOR | SPATS2L |
| CHD1 | PRKCI | ASH1L | ATAD5 |
| KIF1B | PRRC2B | USP19 | GTF2B |
| LYRM5 | TMEM184B | ZSWIM6 | CDK5R1 |
| TMEM39A | SETD7 | PANK3 | UBC |
| CCNL2 | EXTL2 | HIF1AN | ZNF510 |
| KIAA0195 | CEBPZ | FAM53C | MCEE |
| CCT7 | IFITM1 | PNKD | CAV2 |
| LPCAT1 | GNPDA1 | RNF149 | RAB1A |
| FBXO30 | SEL1L | ITSN2 | ERBB3 |
| CALM3 | ZNF431 | RYK | FOXC1 |
| CALM1 | CPEB4 | CCNT1 | QSER1 |
| RFFL | USP31 | LAPTM4A | ZKSCAN8 |
| C16orf87 | SPTLC2 | FAM46A | WDR6 |
| TCF4 | FTH1 | SLPI | GID8 |
| NUP153 | IL10RB | UBA6 | CDK6 |
| MAP3K3 | NF2 | UCKL1 | C10orf2 |
| IMMP2L | PIK3R4 | VPS4B | HEG1 |
| YOD1 | UBQLN1 | TTC28 | ATP6V1G1 |
| NKX3-2 | CTBP1 | SOGA1 | TFAP2A |
| IWS1 | DPF3 | ZNF134 | DDX3X |
| ZNF10 | RNF103 | WSB2 | SERINC1 |
| APC | PAFAH1B1 | EML5 | CORO1C |
| HMGCR | DENND1B | NXF1 | C1orf35 |
| VPS13D | PITPNA | RASA1 | PHF10 |
| OTX2 | KLF2 | LARP1 | CPD |
| CFL1 | SOWAHC | FH | PPIA |

| | | | |
|---|---|---|---|
| COL12A1 | DUSP12 | MED24 | MSN |
| UBR5 | PWP1 | PIAS3 | SFT2D3 |
| ARF3 | ZNF200 | CSRP1 | FARSA |
| AFTPH | SGPL1 | LRRC59 | SRSF2 |
| SIRT1 | LMBR1L | CERS2 | DEK |
| THBS1 | NEDD4L | MAP3K14 | CDK2AP1 |
| USP6NL | YWHAG | TET3 | LYSMD1 |
| HMGA2 | SEC23A | THRAP3 | APP |
| ZNF780B | MCL1 | EPT1 | TAOK1 |
| PIK3CB | DUSP6 | MAPK9 | DST |
| ATP6V0E2 | HLA-E | EIF2A | CANX |
| ZMYND11 | TMEM98 | SLIT2 | TIPARP |
| ZNF267 | GNA13 | ARHGEF2 | PDS5A |
| AGO3 | ZDHHC5 | IGFBP7 | VPS13A |
| ROPN1L | CCND2 | PDSS1 | NSD1 |
| RORB | RAB7A | ENO1 | RCN2 |
| FAM47E-STBD1 | NDNL2 | B4GALT1 | NCL |
| TMEM14A | PSMD2 | RPL13A | FAM222B |
| FKBP4 | ACTG1 | XPO6 | CSRNP2 |
| C11orf30 | SATB2 | TXLNA | C15orf39 |
| AKIRIN2 | MARS | MUL1 | INSIG1 |
| TPR | TM9SF3 | CCND1 | PAQR4 |
| ANKRD52 | DBT | C10orf12 | CBFB |
| PRRC2B | SLC25A32 | CNOT7 | CUL4A |
| SAMD8 | ZZZ3 | LPAR1 | SIX1 |
| SPATA2 | SNX30 | CTNNA1 | ZNF197 |
| TADA2B | STK38 | GIGYF2 | MBD3 |
| REST | LMBR1 | MYLIP | ZSWIM6 |
| GBA2 | MAPKAP1 | EPHA3 | PANK3 |
| PPP2CA | PERP | ASPM | PURB |
| SETD7 | ROCK2 | PRRG1 | SREBF2 |
| HOOK3 | ZNF551 | ZNF264 | ADAM17 |
| B4GALT5 | RECQL5 | GCN1L1 | EEF1D |
| SEL1L | ZNF148 | NUFIP2 | STK39 |
| ZNF431 | ZNF45 | SEC23B | PDE8B |
| IPO13 | SYPL1 | EAPP | TRIP12 |
| CACNB4 | ZFC3H1 | LRRC58 | BCAT1 |
| CPEB4 | BACH1 | HIF1A | STC1 |
| MIR1199 | CASP1 | SACS | DYRK2 |
| COPS7B | YPEL2 | MAPK8 | NFYB |
| TCF7L2 | EXOC6B | PPP2R2A | RELT |
| MTX3 | BCL9L | LNPEP | DCAF7 |
| ZYG11B | GOLGA8B | DESI2 | ETV5 |
| ZNF33B | RAD54B | FAM134C | FRAT2 |
| USP1 | MAP2K5 | IRS4 | PPP4R2 |

| | | | |
|---|---|---|---|
| TSC22D1 | PTPRD | MFSD10 | PRR14L |
| KIF2C | ZNF566 | SON | FAM46A |
| ATG2B | BUB1 | HNRNPC | PPP3CA |
| NR6A1 | LCOR | LPL | AGPAT1 |
| KCNQ5 | FAM69A | FAM83D | UBQLN2 |
| CDKN3 | FZD6 | HSPA14 | MME |
| AP2B1 | GPR180 | WDR36 | KIF16B |
| SORT1 | ZNF24 | GPRC5A | ARHGAP35 |
| RNF103 | TOB1 | PTGER2 | FOXP1 |
| RSF1 | MIEN1 | PTP4A2 | CNN2 |
| PQLC1 | GAN | COG5 | NLK |
| TAF2 | KIAA1671 | FLRT2 | HECTD2 |
| GPAA1 | SETD1B | ITPRIPL2 | ZBTB37 |
| CDADC1 | ARID1B | ZFP36L1 | HADHA |
| KIF3A | IRGQ | C4orf3 | PEG10 |
| SEPT11 | TRPS1 | ARID1A | MLEC |
| OGT | ITGA5 | UBA1 | WSB2 |
| IDS | FBXO45 | UBAP1 | LARP4B |
| ATAD1 | PLCL2 | ZNF460 | FAF2 |
| ZNF597 | EYA4 | RRAGA | SF3B6 |
| NAP1L1 | MAP4K3 | CABLES1 | KIF3B |
| ARL2BP | SLC4A4 | ATRX | ZFYVE16 |
| ZNF200 | CCT5 | COL1A1 | RAB11FIP2 |
| IL2RB | IER2 | CBX6 | HDGF |
| YWHAG | NDUFAF3 | HCN3 | DDIT4 |
| SEC23A | BCL11A | RNF19A | CCSER1 |
| PPP1CC | PTBP1 | ASL | RIF1 |
| MCL1 | ZNF507 | CHTOP | CDKN1C |
| COASY | DBF4 | REEP4 | CASP3 |
| PTPN4 | SELT | ITPR1 | ZNF227 |
| DUSP6 | CDKN1B | RPRD1B | ZNF711 |
| GNA13 | SYNJ1 | NEK9 | PPP1R15B |
| FRA10AC1 | MYH9 | SATB1 | PTPRE |
| CELSR2 | DSTYK | RPL5 | RNF4 |
| ZFP14 | RFX1 | DOCK4 | TXLNA |
| MRPL34 | TUBB | COMMD10 | BAG3 |
| NUDT21 | SLC31A1 | COPS4 | CCND1 |
| COL27A1 | MACF1 | GLCCI1 | LHFPL2 |
| HSP90AA1 | ADNP2 | NXT2 | WDR82 |
| ADSS | DDX6 | NRP1 | RBM3 |
| ZNF445 | ANKIB1 | SLC25A36 | FNDC3B |
| RABGAP1 | HIPK3 | CUTA | MSH6 |
| ZNF260 | DLG5 | VTI1A | SLAMF6 |
| SATB2 | CCDC14 | MBD6 | PABPC3 |
| HNRNPDL | CASD1 | LDOC1L | METTL17 |

| | | | |
|---|---|---|---|
| FKBP14 | CAND1 | SRRM2 | MYLIP |
| TM9SF3 | AFF4 | SYNE1 | RBM17 |
| TOMM20 | IBTK | WDFY3 | QARS |
| LSG1 | ANKRD17 | RAC1 | CDV3 |
| TTN | STAG2 | SPRY4 | PGBD4 |
| RC3H2 | EPG5 | MESDC2 | TMEM19 |
| MERTK | DIRC2 | KMT2D | TPD52L2 |
| SLC25A32 | CHST9 | VPS26B | C1orf198 |
| HNRNPF | CSDE1 | SLC25A11 | E2F3 |
| ACOX3 | TTC17 | C6orf89 | SMARCA5 |
| EPHA5 | NA | ATP6V0C | PAIP2 |
| SSB | WDR20 | CS | RGMB |
| PERP | MEF2D | TNFRSF10B | DPYSL2 |
| HERPUD1 | MTMR9 | HSPBP1 | ANP32B |
| FAM198B | TNKS2 | PLIN5 | RQCD1 |
| ARPP19 | STAG1 | UBXN11 | ZC3H7B |
| AKAP9 | MAN2A1 | YY1 | HERC2 |
| SPRED1 | C19orf12 | TFDP1 | PPP2R2A |
| SKA2 | DEPDC1B | SP1 | RNF24 |
| OTUD6B | TECPR2 | DUSP7 | YPEL3 |
| MYH10 | SETDB2 | JADE2 | LNPEP |
| ARNT | ZNF292 | PDCD11 | FAM134C |
| ETS1 | RAB21 | KRT80 | ACTR2 |
| ZNF850 | WEE1 | SYNJ2 | SUPT16H |
| HLA-DQA2 | SETD5 | RRNAD1 | PPP1R15A |
| DYNC1H1 | ZNF41 | EGLN2 | QKI |
| ZC3H12B | TMBIM1 | TNKS1BP1 | p27(hsa) |
| ARHGAP17 | CEP85 | SERF2 | ASXL3 |
| SYPL1 | AGFG1 | ARHGAP29 | EPM2A |
| BACH1 | PPCS | USP42 | CTDSPL |
| SAMSN1 | MSL2 | REPS2 | FBXO3 |
| PAK7 | SH3D19 | LAMC1 | FAM83D |
| LRRC8D | CD164 | MKI67 | PRELID1 |
| PRPF39 | LLPH | RRP1B | RPS24 |
| AGRN | RAB12 | SUMO3 | VCAN |
| FRS2 | SSR2 | MYADM | ELMSAN1 |
| CBX4 | C6orf211 | CSNK1E | LIFR |
| CTC1 | UGP2 | SLC39A13 | ATXN1 |
| ITCH | EVI5 | DESI1 | ARID1A |
| BCL9L | ZFHX3 | AMOTL2 | CLDND1 |
| TOM1L1 | GNAI3 | CBLN3 | AGPS |
| CAMSAP2 | HSPD1 | PPHLN1 | PPP2R1A |
| ZNF121 | ARF1 | GNAQ | GALNT3 |
| DIP2B | LY75 | TTLL4 | ZNF460 |
| ANKRD50 | RAB23 | FAM114A1 | TFRC |

| | | | |
|---|---|---|---|
| ANGPT2 | TARDBP | DPH5 | BIRC6 |
| RNF150 | CLTC | USP14 | KPNA1 |
| SLC25A25 | NOTCH2 | STARD13 | CEP350 |
| SOAT1 | SERAC1 | CREB3L2 | BRAP |
| PPP3R1 | USP47 | SCAF11 | ARNTL2 |
| SAMD9 | ARHGEF10 | CKAP4 | VCP |
| TOB1 | SNN | RGS16 | CHD3 |
| ENTPD6 | LUC7L3 | NAE1 | APLP2 |
| PLEKHJ1 | NOP56 | HNRNPH1 | CNNM4 |
| WAC | NHLRC2 | OAZ2 | NDUFV3 |
| GAN | ZNF224 | ABCA5 | HES4 |
| ASXL1 | ARFGEF2 | DDX19A | ZBTB5 |
| MTHFD1L | XBP1 | RC3H1 | MIA3 |
| WNK1 | PPP2R5C | GLP2R | ERC2 |
| NME6 | ITM2B | FBXO44 | KIT |
| GSE1 | ASPH | BAHD1 | PLIN3 |
| CDC5L | GRAMD3 | ZBTB33 | TRIM33 |
| MEST | SF1 | C5orf24 | CREBZF |
| SEMA3C | ANXA2 | CYP1B1 | ARIH2 |
| ZFYVE26 | ST6GAL2 | SCD | PABPC1 |
| TMEM41A | VPS26A | CSTF2 | MAP3K10 |
| CA12 | TJP1 | LETMD1 | DHX8 |
| ARID1B | CDK5R1 | ZNF644 | NOL11 |
| SNAI2 | FAM91A1 | ATXN2 | BAZ2A |
| MARK2 | ASCL2 | LASP1 | SATB1 |
| ZNF562 | APPL1 | IPO8 | CSNK2A1 |
| HSPA5 | TMEM164 | EPS15L1 | JAK3 |
| MGA | CD2AP | NHSL2 | TMCC1 |
| DCST1 | RAB1A | CALR | PTPN11 |
| ADO | KIAA0196 | BAG6 | HSP90B1 |
| CDCA4 | PRSS23 | RBM12B | PHC2 |
| C14orf169 | COPS2 | SGPP2 | HDAC2 |
| FRK | QSER1 | HIAT1 | NRP1 |
| CLCC1 | HNRNPU | TBP | SLC25A36 |
| BPTF | TCF19 | LTBP1 | UBE2N |
| FBXO45 | COQ9 | KIAA1919 | KLHL15 |
| BACH2 | SLC16A13 | EIF4EBP2 | PFDN6 |
| TMEM138 | KRAS | VGLL4 | RCHY1 |
| LSS | GXYLT1 | LDLR | TBK1 |
| PLCL2 | CNOT8 | DCK | ENTPD1 |
| CHUK | CDK6 | RIPK1 | LARP4 |
| NHLRC3 | VPS37B | DKK1 | PTPRJ |
| TMEM64 | KALRN | PPM1B | LDHA |
| NBN | FOXN2 | CREG1 | SPRY4 |
| EYA4 | FYN | TUB | IFRD1 |

| | | | |
|---|---|---|---|
| MAP4K3 | THOC1 | BTBD7 | CYBB |
| RRM2B | GATA2 | HIPK1 | CTPS2 |
| ZNF664 | DDX3X | AHNAK | MESDC2 |
| MAP4K4 | HACE1 | LEPR | FGF2 |
| CHL1 | SETD2 | UBE2W | DNMT3A |
| BCL2 | ELAC2 | HERPUD2 | RNF219 |
| AFAP1 | SLC35A4 | MFGE8 | BCL2L2 |
| CPSF6 | ECI2 | DYRK1A | SESN2 |
| CDKN1B | CNNM2 | UHRF1BP1 | C6orf89 |
| SLCO2A1 | HERC1 | PELP1 | CS |
| FOXK1 | WBP5 | PRDM4 | MXD1 |
| CEP120 | CORO1C | ATP1A1 | C16orf72 |
| SERTM1 | PHF10 | TTC9C | TXLNG |
| NDUFAB1 | PRKCE | RXRB | BUB1B |
| MED13 | CHD4 | GTF2A1 | PSMD12 |
| BIRC5 | ASF1B | CD55 | CLASP1 |
| TUBB | WDR26 | RNF213 | ZC3H11A |
| SLC31A1 | HRH1 | QPCTL | ERCC4 |
| SNX9 | TRAM2 | NUP188 | HMG20A |
| ADNP2 | ADD3 | PNRC1 | RANBP2 |
| DDX6 | CCDC71L | MAP3K7 | SUV420H1 |
| RALBP1 | CELF1 | WLS | ULK3 |
| ANKIB1 | FGFBP1 | ADSL | HIVEP1 |
| PPTC7 | NUDCD1 | PRDX3 | DAB2IP |
| ERG | RABEP1 | ABL2 | NDUFA1 |
| HIPK3 | LBR | RPL27A | CLP1 |
| CCR6 | CKAP5 | VPS35 | RBPJ |
| PPP6C | TAOK1 | CYB5R3 | SSR3 |
| NR4A2 | SIK2 | SERTAD2 | OSTM1 |
| CCDC14 | ERGIC2 | TIMM10B | SLC39A6 |
| CAND1 | UCHL5 | TMEM136 | FAM168B |
| TAAR6 | NPY1R | CPNE8 | PLXND1 |
| KLHL5 | PLAGL1 | LYSMD3 | AKIRIN1 |
| LIN28B | CANX | SLMO2 | NPTN |
| TMX1 | KBTBD6 | MAPKAPK2 | MARK1 |
| AFF4 | KIAA1109 | STK40 | TLE3 |
| IBTK | DCTN2 | JMY | TSPAN17 |
| STAG2 | NCOA3 | GATA6 | PEAR1 |
| UBE4A | RNF128 | SLC1A5 | UHRF1 |
| UTP20 | AGPAT5 | HIVEP3 | NMT1 |
| AGAP1 | ANK2 | RAB14 | SLC20A1 |
| STT3A | DUS2 | RFT1 | PRKAB2 |
| PRKCD | ZXDA | RAB5B | LAMC1 |
| N4BP2L2 | NSD1 | RABL6 | MKI67 |
| DGCR2 | CCT6A | RITA1 | METTL2B |

MCFD2
MIB1
WDR20
SLC7A11
C9orf16
SUDS3
TMEM257
MTMR9
HMGB2
SP3
TOMM70A
MOB1B
CHRM3
ATP1B1
URI1
C19orf12
CTSD
YWHAB
RNF217
PSMD3
WEE1
SETD5
SFT2D2
DGUOK
PHF3
ZNF736
RCOR1
IVD
EPN2
BLMH
IGF1R
SH3D19
LONRF1
CD164
OSBPL3
MINK1
DDX17
ZNF627
BTBD3
EGFR
C6orf211
HERC3
SRPK2
ANKRD1
PIBF1
PHTF1
NAB1
CHEK1
USP28
SLC35B3
C4BPB
MEGF8
INSIG1
FAM104A
CNP
PRRC2C
RPP14
RICTOR
ASH1L
PANK3
EDEM1
PURB
KLHL11
FADD
USP45
TAS2R30
ARIH1
IARS2
IFIH1
SLC25A38
STK39
BCAT1
DHRS7
FAM20C
CDC42BPG
AMD1
DYRK2
H2AFV
RAB29
SBF1
SNX3
HECTD1
NUMA1
PAPD5
ASAP1
KIAA1524
LAPTM4A
FAM46A
ACAA1
MTR
KRBA1
ZW10
MTF1
HLA-A
ZDHHC7
PLS3
USP34
COQ7
MAP1B
LIG4
YY1AP1
ZNF774
ASAP2
UBOX5
ZDHHC13
TXNIP
PALM2-AKAP2
SMEK2
MIDN
BBC3
HDLBP
ZCCHC14
SLC44A1
DCUN1D1
CHSY1
TAX1BP1
PPP2CB
ZBTB7A
REV3L
RASEF
SEP15
HOXC10
COX15
TOPORS
H3F3B
SEC24B
BRD1
STAMBP
UBE2D1
RBBP7
LYRM4
MFF
KIAA0040
PDGFD
RC3H1
CDC27
KPNA6
GNAI2
MYO10
ZBTB33
SLC6A9
PDE7B
CYP1B1
SCD
SEH1L
VPS53
SDC2
NUF2
NGRN
PCDHGA4
NUP43
KIAA0368
VEZF1

| | | |
|---|---|---|
| AGO4 | PKDCC | TRIM5 |
| CD96 | GET4 | p27kip1(hsa) |
| COX7A2L | ATP6V1B2 | UBN2 |
| TMOD3 | ZNF721 | GATAD2B |
| MSANTD4 | KRIT1 | SGTA |
| RAB23 | SERP1 | ABHD3 |
| PRR5L | ARID2 | CTCF |
| NACC1 | VPS4B | GJA1 |
| TARDBP | PRICKLE1 | STMN1 |
| CLTC | NPM1 | BIRC3 |
| KPNA4 | NLK | POFUT1 |
| DNAJC7 | DKC1 | NAPEPLD |
| NOTCH2 | DUSP10 | CCNJ |
| ZNF519 | TRAFD1 | HIAT1 |
| CARS | PTGES2 | MBNL1 |
| ZNF800 | TES | TMEM201 |
| GLS | CEBPB | OXR1 |
| C1QTNF9 | ZNF134 | SEMA6D |
| ID4 | GTF3C2 | ZBTB18 |
| VCL | SETDB1 | CSTF2T |
| PSKH1 | WSB2 | VGLL4 |
| C6orf48 | POLR2D | CAD |
| CTSC | EML5 | SESN3 |
| GNB1 | ZNF217 | COPA |
| AMOT | ITGAV | HNRNPA2B1 |
| ZNF625 | DCAF6 | PER2 |
| MTCH2 | MCM2 | ZIK1 |
| HAS2 | EHBP1 | BNIP3L |
| TIMP2 | FBN3 | NR2C2AP |
| ETF1 | ITGA11 | BCOR |
| PPP2R5C | SH3BP4 | EIF5A2 |
| POLR1D | FH | TUB |
| HNRNPH3 | TAF1 | CUL3 |
| MAP3K13 | ABCF2 | MMRN1 |
| HN1L | KIAA1147 | FAM214A |
| FAM175B | KIF3B | HIPK1 |
| CHD7 | NRF1 | AHNAK |
| SH3GLB1 | CCAR1 | YES1 |
| TJP1 | TMX4 | AP1G1 |
| UBE2J1 | MAL2 | PAFAH1B2 |
| FAM91A1 | CNIH1 | EHD2 |
| ZNF510 | PHTF2 | TMEM127 |
| TMEM164 | NDE1 | ZNF367 |
| CD2AP | CDH1 | MTOR |
| RAB1A | DENND5A | ZFP36L2 |

| | | |
|---|---|---|
| PDCD6 | KIAA1432 | OSBPL8 |
| CFI | PITPNM2 | ZNF106 |
| PSAP | SFXN1 | ABCC5 |
| COPS2 | CUL5 | NKRF |
| IRF8 | NFAT5 | DYRK1A |
| EIF1AX | PIKFYVE | FUT8 |
| STT3B | PPP1R12A | KLHL24 |
| HNRNPU | AEBP2 | CTDSPL2 |
| MRPS14 | KLHDC10 | LEPROT |
| HSDL1 | ZBED6 | DPH2 |
| ZKSCAN8 | PLEKHG3 | UBXN2B |
| UBE2I | NECAP1 | ACIN1 |
| MAGED1 | PKP4 | ATP1A1 |
| KRAS | CLIP1 | SUN2 |
| GXYLT1 | GANAB | NAPA |
| WIZ | BTG2 | TRIM71 |
| SPRYD7 | FBN1 | C9orf156 |
| RSL1D1 | PPP1R15B | PTEN |
| EPDR1 | HIST2H2AA3 | ZNF652 |
| HEG1 | DCP2 | NFATC3 |
| HEY2 | FBN2 | TRAM1 |
| IPO7 | RNF4 | FERMT3 |
| WBP2 | TXLNA | ISCA1 |
| DDX3X | TMEM173 | SLC35F6 |
| PDIA6 | GOLGA8A | ALDH1A1 |
| IGF2BP1 | CCND1 | PPFIA1 |
| MDH2 | N4BP2 | FNDC3A |
| COPB1 | LHFPL2 | SDCBP |
| GOLGA6L4 | FNDC3B | NUPL1 |
| FSTL1 | C2CD5 | SUCLG1 |
| INIP | NRXN3 | TET2 |
| EYA3 | CTNNB1 | USP18 |
| TCEB1 | ZNF805 | SLC40A1 |
| ZNF189 | AP3B1 | SCARB2 |
| MTX2 | EFR3A | AP3M1 |
| ARPC1B | MMADHC | KMT2C |
| PHF10 | LPAR1 | HELLS |
| FBL | ANXA7 | HOXA10 |
| SLC35A5 | PTBP3 | RNF26 |
| TCP1 | ASNS | SERBP1 |
| ACOT12 | PMM2 | DVL2 |
| FKBP1A | CCNE2 | TAF3 |
| CALM2 | AXIN2 | ARF4 |
| PRKCE | PLEKHB2 | ZCCHC24 |
| SMARCD1 | NR3C1 | MRPL19 |

| | | |
|---|---|---|
| CPD | SSFA2 | BRWD1 |
| ZBTB41 | SBNO1 | TBL1XR1 |
| BUB3 | GIGYF2 | ERO1LB |
| MTMR12 | KCTD2 | ZNF91 |
| ZCCHC2 | MYLIP | HYOU1 |
| SNIP1 | TRAPPC9 | BZW2 |
| SRSF2 | ASPM | SECISBP2L |
| SLC35C2 | CDV3 | EIF1 |
| MORF4L2 | ZNF264 | PTCD3 |
| WDR26 | FAR1 | TFG |
| ZNF107 | AXL | MAML1 |
| EPC1 | NUFIP2 | ZFAND3 |
| DEK | TMEM87A | TSPAN13 |
| ADD3 | MSL1 | ARHGAP5 |
| PRPF8 | LRRC58 | RAPH1 |
| CELF1 | ADM | RAB5B |
| PDS5B | DYNC1LI2 | MDM2 |
| SRSF7 | GIT2 | PSMB5 |
| FBXL7 | E2F3 | PGK1 |
| SLC9A6 | SMARCA5 | ZNF629 |
| CKAP5 | SACS | UBE2J2 |
| TAOK1 | ATP5A1 | TAF6L |
| TFPI | GDAP2 | BMPR2 |
| ERGIC2 | DMXL1 | TMBIM6 |
| DST | KPNA5 | TOR1AIP1 |
| KIAA2026 | COX5B | ATF4 |
| SUMF1 | RQCD1 | BCL2L11 |
| RBAK | DNAJB9 | DUSP19 |
| KIAA0101 | CGGBP1 | MTF1 |
| FOXK2 | DNAJC13 | LIN7C |
| ADIPOR2 | NEFH | PLS3 |
| ABI3BP | CHCHD10 | PIN4 |
| KIAA1109 | DPY30 | USP34 |
| HK2 | LNPEP | MIER3 |
| NCOA3 | AFF3 | SLC30A1 |
| C3orf14 | ZNF44 | UBR4 |
| PKNOX1 | IRS4 | NACC2 |
| ZNF622 | MYC | TCTEX1D2 |
| LPP | JKAMP | FNIP2 |
| TMEM123 | NR2C2 | ANAPC16 |
| GIGYF1 | CENPF | KDM5B |
| ATP2B1 | IKZF5 | EFNA1 |
| GPR183 | SON | ELF1 |
| NSD1 | HCFC1 | ACACA |
| TMEM184C | IQGAP2 | MLLT6 |

CCT6A
PLEKHA8
RNF220
PIP4K2A
ATP5B
PRCC
NAB1
RPLP1
RFC2
MYCBP2
ZDHHC3
USP28
RBM38
SLC16A3
POLR2K
MPP5
FOXB1
INSIG1
RALGAPB
MFHAS1
ZNF571
TP53
TNRC6B
RNF187
SPTY2D1
LRIG2
ZNF543
LITAF
LEMD3
GLTSCR1L
DDX21
ZNF197
RICTOR
ASH1L
MBD3
ADAMTS2
ZSWIM6
EGR3
PANK3
MYNN
PURB
KLHL11
ACSS2
AMMECR1
LCP1
RAPGEF6
MARCHF8
TXNRD1
PPM1A
BTBD2
VEZT
MTMR1
DEPDC1
WSB1
INSIG2
ELMSAN1
HIST1H3D
GPAM
MRPS2
OCIAD1
PGBD2
C21orf91
TMEM9B
ARPC2
GOLGA4
GPBP1L1
KLF3
ATXN1
FAM49B
STK35
HNRNPA1L2
PCBP1
YARS
ATP10B
CPNE3
MTPN
ZNF460
TFRC
JRKL
TMED10
CCDC47
SRPR
ARNTL2
ZC3H12C
SYT1
XPR1
MAFK
AARS
ZNF687
APLP2
ZNF749

| | |
|---|---|
| SREBF2 | CNNM4 |
| ZDHHC12 | CR1 |
| HDHD1 | PRNP |
| PRDM1 | FBXO10 |
| ARIH1 | MYO5A |
| ADAM17 | SLC38A2 |
| UBA2 | MYCBP |
| ZNF432 | MUC5B |
| PDE8B | ZNF100 |
| TRIP12 | MIA3 |
| BCAT1 | ABHD5 |
| DHRS7 | ZC3H13 |
| SIPA1L1 | MED31 |
| AZIN1 | DEPDC4 |
| ATP9A | CERK |
| AMD1 | XYLT2 |
| POSTN | NOL4L |
| DYRK2 | HNRNPM |
| ZNF317 | CHTOP |
| ZNF253 | SMAD5 |
| CREB1 | PSMC4 |
| TSG101 | MMD |
| SORBS3 | GTF3A |
| RBMS1 | SF3A2 |
| ZMYM6 | NCOA2 |
| PPP4R2 | ITPR1 |
| CCSAP | RBM47 |
| DSCR8 | DKK4 |
| SLFN11 | TCF20 |
| FAM122B | AP1S3 |
| C7orf73 | NEK9 |
| CHPF2 | SREK1 |
| ARL8B | BAZ2A |
| HECTD1 | SKI |
| PRTG | TGIF1 |
| APAF1 | PRDM15 |
| PAPD5 | SOS1 |
| MRPS5 | ADPRHL2 |
| MAPK1IP1L | CPEB2 |
| GREB1L | UBR2 |
| BPGM | RORA |
| ACVR2B | UMOD |
| EDEM3 | KIAA0284 |
| GABBR2 | SLC25A36 |
| RAB5A | ZNF184 |

| | |
|---|---|
| MTR | SRRM2 |
| UGT3A1 | GEMIN5 |
| PCGF5 | METTL10 |
| UBQLN2 | KLHL15 |
| KRIT1 | FXR1 |
| ZNF136 | PRODH |
| HSD17B3 | DCUN1D3 |
| GADD45G | METTL21A |
| EIF5 | SLC26A2 |
| ARHGAP35 | LYST |
| PTK2 | CUX1 |
| SLC2A3 | PTPRJ |
| ST3GAL5 | XRN1 |
| SLC25A30 | S100A10 |
| FBXO34 | C6orf62 |
| SLC9A3R1 | DUSP8 |
| PTPN13 | NOLC1 |
| IDI1 | SHOC2 |
| LCMT1 | AIMP1 |
| TRUB1 | MCUR1 |
| FOXP1 | SNAPC1 |
| SOGA1 | SH3RF1 |
| DKC1 | ANKRD46 |
| FURIN | TRIO |
| TES | PIK3C2A |
| HECTD2 | RDH10 |
| ZNF134 | C16orf72 |
| TICRR | PAWR |
| ZBTB37 | DSTN |
| PNPT1 | PUS7 |
| ETNK1 | FAM199X |
| PEG10 | TXLNG |
| STX7 | TMEM2 |
| MOB3B | PPP1R12C |
| MLEC | LCLAT1 |
| ARHGAP18 | NF1 |
| AP000350.10 | FAIM |
| ZNF232 | ZNF430 |
| CADM1 | STRN |
| LARGE | MADD |
| BRCA1 | CDK13 |
| ZNF217 | DDX55 |
| DIP2C | RANBP2 |
| KLRC4 | SUV420H1 |
| LARP4B | ZFP1 |

| | |
|---|---|
| NAA20 | CXXC5 |
| PSD3 | HIVEP1 |
| RASA1 | SP1 |
| NAA50 | DAB2IP |
| LIF | RARS |
| ZNF268 | LURAP1L |
| PTCD1 | COL1A2 |
| CPT1A | NDUFA1 |
| LDLRAP1 | RBPJ |
| ZNF697 | SYNPR |
| DMWD | PSMB1 |
| TBX4 | LRPAP1 |
| PRCP | BCL6 |
| DERL2 | GOT2 |
| DOCK10 | OSTM1 |
| CSRNP1 | FNIP1 |
| CSE1L | SMIM13 |
| FKBP7 | FAM168B |
| NA | NPTN |
| CNIH1 | CDC23 |
| PHTF2 | PTPLAD1 |
| PMAIP1 | RASSF3 |
| ANP32A | YBX1 |
| URB1 | ARRDC3 |
| YIPF4 | NFE2L2 |
| MRPS18B | IKZF2 |
| SMC4 | PSMA3 |
| DDIT4 | PGRMC1 |
| SFXN1 | SLC20A1 |
| CUL5 | FOXJ3 |
| RBM39 | MKI67 |
| TET3 | FXR2 |
| NA | PRKAR1A |
| DROSHA | BARX2 |
| ZBED6 | FAM160B1 |
| LFNG | EP300 |
| RIF1 | SUN1 |
| SF3B3 | MGEA5 |
| AGO1 | ARL5B |
| NECAP1 | ZCCHC14 |
| TMSB4X | PCDHB13 |
| EPT1 | GNAQ |
| OGDH | RASSF9 |
| SGMS2 | PUM2 |
| CRYBG3 | NFIB |

| | |
|---|---|
| PCNX | AHR |
| TTC37 | REV3L |
| C1orf52 | CELSR3 |
| KHSRP | ATF7IP |
| CLIP1 | BRD3 |
| NOTCH1 | GOLGA1 |
| GANAB | TEF |
| POLR2A | YIPF5 |
| ZNF26 | CNOT1 |
| CENPI | UBE2D3 |
| IGFBP7 | OSGIN2 |
| FBN1 | SLC9A7 |
| ENO1 | CREB3L2 |
| B4GALT1 | ATXN7 |
| ZNF227 | SCAF11 |
| AK6 | AMIGO2 |
| ALG10B | VPS41 |
| JUN | CTSB |
| ZNF711 | H3F3B |
| PIK3R3 | ARMC1 |
| PPP1R15B | CERS6 |
| RPL13A | UBE2D1 |
| WDR33 | TARBP2 |
| CCT3 | C11orf24 |
| DCP2 | G3BP2 |
| ATP6AP2 | PCDH7 |
| CLINT1 | ZNF730 |
| ZNF530 | FAM210A |
| LRCH2 | SNX13 |
| XPO6 | CDC27 |
| NCS1 | KPNA6 |
| FAM208B | IGFBP3 |
| SLC20A2 | ZNF211 |
| CCND1 | LONRF2 |
| IMPAD1 | MYO10 |
| ILF3 | ABI1 |
| N4BP2 | NTN4 |
| PFN2 | MED23 |
| LHFPL2 | MARCHF5 |
| URM1 | UBE2G1 |
| PPARA | SDC2 |
| WDR82 | CSTF2 |
| FNDC3B | MYEF2 |
| C2CD5 | PAPD7 |
| JUNB | MIS12 |

| | |
|---|---|
| **ICMT** | **PCMTD1** |
| **CTNNB1** | **AURKA** |
| **AP3B1** | **IL6ST** |
| **EFR3A** | **ATG5** |
| **HMBS** | **LEPREL2** |
| **ZNF678** | **DNAJB12** |
| **SOWAHA** | **THOC2** |
| **PUM1** | **NUP43** |
| **ACTR3** | **PLEKHA1** |
| **LPAR1** | **FAM102A** |
| **SPEN** | **AKR1A1** |
| **PTBP3** | **VEZF1** |
| **S1PR1** | **MYO5C** |
| **ZNF236** | **NSF** |
| **GFPT1** | **UBAP2L** |
| **GGPS1** | **MED19** |
| **PLEKHB2** | **CTTNBP2NL** |
| **CHRFAM7A** | **RLF** |
| **NR3C1** | **ZNRF2** |
| **E2F5** | **NDC1** |
| **SEC24C** | **SLC7A2** |
| **STRBP** | **FGD4** |
| **SBNO1** | **TULP4** |
| **NCOA6** | **AMACR** |
| **MPHOSPH6** | **FMN2** |
| **BHLHE40** | **IPO8** |
| **ARL6IP1** | **GATAD2B** |
| **IFFO2** | **CTCF** |
| **CDV3** | **COG3** |
| **DOCK1** | **RNF182** |
| **ZNF264** | **CDKN1A** |
| **C1orf131** | **KIAA0232** |
| **FAR1** | **DOCK9** |
| **TSR1** | **MDN1** |
| **SEC23B** | **SLC2A1** |
| **ST8SIA4** | **USP9X** |
| **BLOC1S6** | **WDR4** |
| **SLC19A2** | **ZFYVE21** |
| **EAPP** | **WDR43** |
| **IGFBP4** | **RP2** |
| **LRRC58** | **PDXDC1** |
| **SLC4A7** | **CNEP1R1** |
| **ADM** | **KIAA1919** |
| **TRAK1** | **MRPL44** |
| **MTA1** | **ZBTB18** |

| | |
|---|---|
| DYNC1LI2 | GFPT2 |
| CISH | SMG7 |
| GATC | SESN3 |
| HELZ | SLX4 |
| ESRP2 | RCOR3 |
| POGZ | TMEM25 |
| SACS | NOP9 |
| C2CD2 | PER2 |
| ZNF443 | ATXN3 |
| SMCHD1 | DKK1 |
| ATP5A1 | PPM1B |
| CEP41 | CCDC186 |
| ICE2 | TNPO2 |
| UBE3A | BCL11B |
| ATL2 | SUGT1 |
| ZNF101 | GAREM |
| PPAP2B | TUB |
| KTN1 | AIFM1 |
| RIT1 | ACPP |
| CSNK1A1 | GID4 |
| CREBRF | CACNA2D1 |
| DNAJC13 | ATP2A2 |
| LRRC17 | MPPED2 |
| RNF24 | HIPK1 |
| LNPEP | AHNAK |
| CEBPA | RINT1 |
| F2R | RGL1 |
| WDR48 | AP1G1 |
| MED26 | SMAD7 |
| MNDA | PAFAH1B2 |
| QKI | LEPREL1 |
| TMC5 | CEP55 |
| UBE3C | RAB8B |
| NR2C2 | UBE2W |
| CENPF | ABCA1 |
| SON | RAN |
| PIM3 | HERPUD2 |
| CD46 | DUSP5 |
| AHCTF1 | GDI2 |
| CTDSPL | JADE1 |
| ZNF626 | SLC12A5 |
| TMEM14C | OSBPL8 |
| MKLN1 | PLEKHA6 |
| PPM1A | USP11 |
| MYO9A | HIST1H1B |

| | |
|---|---|
| **HSPA14** | **NCKAP1** |
| **PRELID1** | **ANAPC13** |
| **RASSF6** | **DYRK1A** |
| **SFXN3** | **SNX25** |
| **WSB1** | **MOAP1** |
| **SRSF6** | **CTDSPL2** |
| **ELMSAN1** | **PTAR1** |
| **HIST1H3D** | **STEAP3** |
| **MLH3** | **PRDM4** |
| **FXYD6** | **DLST** |
| **LIFR** | **HIST2H2BF** |
| **TMEM9B** | **CAMK1D** |
| **ZFP36L1** | **KIRREL** |
| **COL5A1** | **ITGA6** |
| **DBNL** | **SPIN1** |
| **ARV1** | **SGPP1** |
| **SMAP1** | **PDZD8** |
| **GOLGA4** | **MAP2K4** |
| **GPBP1L1** | **SAMD9L** |
| **BLOC1S5-TXNDC5** | **RPS6KA4** |
| **KLF3** | **XRCC5** |
| **ATXN1** | **CHKA** |
| **FAM49B** | **RSBN1** |
| **CCDC82** | **RAB11FIP1** |
| **DEGS1** | **ATP8B1** |
| **DENND5B** | **GTF2A1** |
| **ZFAND1** | **TANC2** |
| **PCBP1** | **XPO1** |
| **YARS** | **EPHA4** |
| **BEND4** | **RRBP1** |
| **MSH2** | **GPBP1** |
| **ZNF132** | **PHIP** |
| **CPNE3** | **CPEB3** |
| **CENPO** | **MUT** |
| **EIF4A2** | **NFATC3** |
| **MTPN** | **DUSP16** |
| **SLC35A3** | **FDPS** |
| **ZNF460** | **PALLD** |
| **TFRC** | **SGK3** |
| **ZMIZ1** | **POLR1A** |
| **MCTP1** | **LIN54** |
| **BIRC6** | **PLEKHM1** |
| **FLNB** | **HDHD2** |
| **ADAM12** | **DEAF1** |
| **DAPK1** | **GRHL1** |

| | |
|---|---|
| TMEM241 | RAD17 |
| TMED10 | C16orf62 |
| ZAK | ATP13A3 |
| ATAD2B | RNF141 |
| SETD8 | SLAIN2 |
| KPNA1 | TLN1 |
| TSHR | GPR126 |
| AAR2 | MAPK6 |
| CEP350 | LATS2 |
| FRMD6 | PTGFRN |
| KIAA1462 | CREBBP |
| ZC3H12C | ZNF146 |
| COL1A1 | PRDX3 |
| ANKRD44 | ABL2 |
| ZMYM1 | FGF7 |
| KCTD3 | TRAF3 |
| ZNF558 | WRNIP1 |
| C6orf106 | USP33 |
| ANKLE2 | ZHX1 |
| ZMYM2 | RAD21 |
| APLP2 | TET2 |
| FLT1 | PCGF3 |
| SLC11A2 | DNAJC27 |
| BEND3 | KMT2C |
| SMNDC1 | CDK16 |
| C8A | MGAT4A |
| RPLP0 | GOLGA3 |
| SLC38A2 | SERTAD2 |
| NCOR1 | TRMT10C |
| AK3 | VAT1 |
| ZNF180 | SLFN5 |
| ARRB2 | PRSS12 |
| NIPA2 | FAM129A |
| UCHL1 | ENTPD4 |
| ULK4 | KLF6 |
| CEMIP | ARF4 |
| ZNF812 | RANBP6 |
| C8orf33 | TMEM74 |
| TMEM87B | KIAA1279 |
| SAV1 | PDK2 |
| VASH1 | RNF139 |
| CTD-2228K2.5 | ATP5J |
| JMJD1C | ZNF470 |
| RNMT | KPNB1 |
| RBM26 | CBX5 |

| | |
|---|---|
| SMAD5 | KLF4 |
| ZNF175 | TNKS |
| C5orf51 | JMY |
| NCAPG | TBL1XR1 |
| UBE2Q1 | WDR37 |
| TRIM33 | SETX |
| CREBZF | ZNF91 |
| TGOLN2 | BAZ2B |
| PTGS2 | NR2F1 |
| DENR | GATA6 |
| AGGF1 | TIMM17B |
| HECA | ZFP30 |
| PATZ1 | PIGA |
| NCOA2 | EIF1 |
| ARL6IP6 | GLYR1 |
| PABPC1 | IRF1 |
| ITGA2 | ADRB1 |
| MAP3K10 | SGK1 |
| SEC24A | PRRC2A |
| PIK3R1 | MARCHF3 |
| ZNF426 | MBOAT2 |
| NEO1 | NAA38 |
| RB1 | RBM15 |
| PNRC2 | MDM2 |
| AP1S3 | BAG5 |
| LRRC8C | GPR137C |
| AEN | PRR11 |
| BCAP29 | TNRC6C |
| BAZ2A | FANCI |
| SLC35E1 | AP5M1 |
| MARCKSL1 | VPS54 |
| CDKN2AIP | FAM110C |
| KDM5A | VKORC1L1 |
| TXNDC12 | RPS29 |
| ARL4D | PNISR |
| TMCC1 | ATF4 |
| RORA | EIF4G2 |
| SLC22A23 | BCL2L11 |
| HSP90B1 | CLOCK |
| CDC42SE2 | KIF5B |
| ZFAND6 | MTF1 |
| FARP2 | SOBP |
| EIF4ENIF1 | MYO1B |
| GLCCI1 | LIN7C |
| NRP1 | CDC42BPB |

SLC25A36
NAA30
UBE2N
MBD6
GDI1
TMPO
SRRM2
RAD18
KLHL15
ATF7IP2
NUAK2
ZNF506
TBK1
JAZF1
GNPNAT1
DDX5
EIF5B
LARP4
RIMKLB
SEC31A
MIF
BACE2
XRN1
NIF3L1
C6orf62
ANKRD13C
OTUD4
STC2
ZNF823
MAPK3
AQR
ANKFY1
PTP4A1
SDAD1
PRR4
DNMT3A
TCF21
PMPCB
KMT2D
ADAMTS1
SS18L2
MAP7
NUP50
IPO5
SESN1
NDUFB4
PAX3
CD44
KCTD20
NPTX1
ATR
MAP1B
CTGF
MIER3
CD69
MDH1
DNAJC30
COL4A1
TCEA1
NUDT19
UBR4
ZKSCAN1
LIG4
BSN
PPP1CB
FNIP2
IDH3G
KDELR2
AGPAT9
TXNIP
MRPS21
HIST1H2AH
NEFL
ANKRD18A
SRGAP2
HNF1B
MARCKS
SLC6A14
HIST1H2BH
TAF15
RNF115
MYO18A

**MCUR1**
**BCL2L2**
**TXNDC15**
**STX12**
**C6orf89**
**TMED4**
**ZNF14**
**CITED2**
**PHYH**
**DSC3**
**HILPDA**
**CARD10**
**CHD2**
**C16orf72**
**PAWR**
**INTS6**
**RAD1**
**LYRM1**
**IGF2BP3**
**P2RY1**
**ELAVL1**
**ADCK3**
**SSX2IP**
**MIOS**
**SNRNP40**
**ZBTB21**
**DHX15**
**BAG2**
**CCDC50**
**CLASP1**
**CALD1**
**ZC3H11A**
**HSPA8**
**LCLAT1**
**NF1**
**FIGN**
**CCNY**
**ZNF700**
**STRN**
**MADD**
**SMARCD2**
**KIAA0100**
**EIF2S1**
**RANBP2**
**DSG2**

**GPR83**
**RAB9A**
**YY1**
**SRP72**
**USPL1**
**INCENP**
**FAM98A**
**RBPJ**
**INPP1**
**MDM4**
**LARS**
**CD93**
**RNF146**
**SSR3**
**BLOC1S2**
**PHOX2A**
**ACVR1C**
**SLC39A6**
**SMIM13**
**FAM168B**
**PROSC**
**ONECUT2**
**GTPBP2**
**AKIRIN1**
**NPTN**
**MAT2A**
**PTPLAD1**
**ZNF451**
**TRPC4AP**
**MARK1**
**AP3D1**
**TSPYL4**
**PEAR1**
**TAF7**
**ARRDC3**
**EPC2**
**GFM1**
**ZNF788**
**SLC20A1**
**ARID4B**
**LAMC1**
**USP24**
**RAI14**
**C15orf57**

**NFE2L1**
**CNPPD1**
**FOXJ3**
**SLC22A15**
**GSTM2**
**RRP1B**
**TGFBR3**
**APEX2**
**VMA21**
**FAM160B1**
**MTRR**
**CD200R1**
**EP300**
**MIDN**
**AMOTL2**
**DDAH1**
**GHITM**
**BTG3**
**ZNF35**
**TMEM245**
**ARL5B**
**ZCCHC14**
**ARFIP1**
**GPCPD1**
**FAM73A**
**CDK17**
**DCAF17**
**PLAG1**
**PUM2**
**ZNF780A**
**NFIB**
**THUMPD1**
**TWISTNB**
**AKT3**
**RSBN1L**
**SPDL1**
**ZNF511**
**ZNF484**
**PITPNB**
**PIAS2**
**ARHGEF15**
**TEF**
**YIPF5**
**CNOT1**
**ZNF281**

**CEP135**
**UBE2D3**
**FSIP1**
**USP14**
**WHSC1**
**OSGIN2**
**TFB1M**
**SCAF11**
**BCL7A**
**KBTBD3**
**H3F3B**
**PTPRZ1**
**ZNF564**
**BRD1**
**WNT3A**
**CD4**
**GAPVD1**
**DDX39B**
**G3BP2**
**RBBP7**
**SEC22A**
**CPOX**
**MBD4**
**SMC2**
**ZFR**
**ZNF730**
**ARCN1**
**CCNE1**
**RC3H1**
**ZNF440**
**CDC27**
**KPNA6**
**GNAI2**
**TUSC1**
**FASTKD5**
**LRRN3**
**LONRF2**
**MYO10**
**SPG21**
**CEP57**
**C5orf24**
**ATP6V1C1**
**PPP1R3B**
**CYP1B1**
**SCD**

**LAPTM4B**
**ACSL4**
**UBE2G1**
**C14orf28**
**RLIM**
**ZNF644**
**SEMA7A**
**THOC2**
**NUP43**
**PLEKHA1**
**FAM102A**
**ZNF704**
**KIAA1244**
**KMT2A**
**RPN2**
**LGALSL**
**UBAP2L**
**ZBTB48**
**TRIM21**
**ALDH9A1**
**GRPEL1**
**RLF**
**IGSF3**
**ZNRF2**
**p27kip1(hsa)**
**CMC2**
**NDC1**
**SLC7A2**
**SEPT2**
**ZNF570**
**UBN2**
**CDIPT**
**CTCF**
**TMEM45A**
**PKN2**
**USP7**
**COG3**
**AKAP12**
**GSDMD**
**MCCC2**
**CALR**
**ARHGAP11B**
**TMEM30A**
**TEAD4**
**ZNF92**

**MAP2K1**
**STMN1**
**UHMK1**
**PKD2**
**CCNJ**
**OSBPL1A**
**ATP8B2**
**MFSD8**
**NANS**
**JRK**
**CHTF8**
**TNRC6A**
**MAP3K2**
**ARF6**
**RSPRY1**
**C1orf109**
**USP9X**
**MBNL1**
**WDR4**
**HEXIM1**
**NFIA**
**ZNF66**
**DNAJA4**
**ZFHX4**
**TRMT1**
**CNEP1R1**
**AVL9**
**UBR3**
**ARSJ**
**BHLHB9**
**GSG2**
**MAP3K8**
**ELOVL6**
**SSBP2**
**ZBTB18**
**PPA1**
**CARM1**
**EIF4EBP2**
**HIGD2A**
**SMG7**
**RPS14**
**SLC39A10**
**TGIF2**
**SESN3**
**COPA**

**RCOR3**
**KLHL12**
**FAT1**
**C17orf51**
**HNRNPA2B1**
**NAA25**
**PDPK1**
**PRDX6**
**RPRD1A**
**PER2**
**HK1**
**ATXN3**
**ZNF483**
**TAF15**
**ZBTB1**
**ADARB1**
**TTI1**
**ZBTB4**
**HMGCS1**
**SUGT1**
**BNIP3L**
**BARD1**
**WNT2**
**TNFRSF6B**
**NDUFS5**
**PNMA1**
**CCNI**
**MMRN1**
**BTBD7**
**FAM63B**
**RUFY2**
**FAM214A**
**CKS2**
**ATP2A2**
**PCBP2**
**STAT1**
**HIPK1**
**ENY2**
**AHNAK**
**TRPM7**
**YES1**
**YTHDF3**
**PRKDC**
**SMAD7**
**PAFAH1B2**

**ORC6**
**SRGN**
**BVES**
**ZNF717**
**CEP55**
**RAB8B**
**UBE2W**
**TCL1A**
**TMEM127**
**RAN**
**MKNK2**
**RHBDD1**
**ZIC5**
**ZNF367**
**GDI2**
**TRRAP**
**ZFP36L2**
**TMEM192**
**OSBPL8**
**NMRK2**
**ZNF106**
**KIAA1467**
**NCKAP1**
**TTC27**
**PSMD8**
**PSMA4**
**GPR157**
**ADK**
**NFYC**
**AGPAT3**
**PLXDC2**
**NKRF**
**ULK1**
**ABHD17C**
**SKA3**
**CTDSPL2**
**ATG10**
**EXOC8**
**MOB1A**
**HSPA1B**
**CLUAP1**
**VCPIP1**
**IL1A**
**SPIN1**
**SUCO**

**UBE2B**
**CNOT6L**
**TACSTD2**
**UQCRQ**
**CLSTN1**
**ZNF22**
**HOXA11**
**TMCO3**
**ZNF699**
**DNAJA2**
**B3GNT2**
**CCNC**
**ZNF618**
**ATP8B1**
**GTF2A1**
**ZNF77**
**XPO1**
**TRIM71**
**FAM171A1**
**FBXO33**
**SELK**
**TIGD2**
**PTEN**
**ZNF652**
**NUB1**
**MTF2**
**EWSR1**
**MARCHF6**
**SNX4**
**SIL1**
**SPTBN1**
**ATL3**
**LIN54**
**ISCA1**
**LATS1**
**FAM160A2**
**S100A13**
**C9orf91**
**CAP1**
**ALDH1A1**
**ADAMTS9**
**DCAF16**
**PCYOX1**
**MAPK1**
**NUP188**

NIPBL
FNDC3A
SLAIN2
MICAL3
MAPK6
MMP14
PTGFRN
ZNF561
ZNF146
PRDX3
ABL2
ZNF772
RFC3
USP33
H1F0
RTEL1
SERPINE1
USP18
FAM73B
PLAU
KMT2C
TBPL1
GRB2
ADCY9
CALCRL
CCNG2
HELLS
GCNT1
GTSE1
BTG1
RBM6
HOXA10
SERTAD2
CAPRIN2
SMAD1
VBP1
PARD6B
AVEN
ZNF567
CBX3
DVL2
MTCL1
LYSMD3
DIS3
GNB4

**ZNF766**
**ENAH**
**AKNA**
**TNPO3**
**ANKRD10**
**ZNF675**
**KNTC1**
**RBM4**
**KPNB1**
**ABHD13**
**FDX1L**
**RHOB**
**ZNF30**
**BRWD1**
**CBX5**
**TBL1XR1**
**ERO1LB**
**WDR37**
**ZNF91**
**LETM1**
**GATA6**
**ZFP30**
**ATRN**
**CCP110**
**GOLPH3**
**ASXL2**
**LYPLA1**
**TGFBR2**
**COLGALT1**
**SCN8A**
**EIF1**
**GLYR1**
**CLEC16A**
**FERMT1**
**ZNF529**
**ZFAND3**
**RPL15**
**SNRNP200**
**MARCHF3**
**MBOAT2**
**GCNT2**
**RBM15**
**PCDHB8**
**CAMK2N1**
**GLYCTK**

**DYNLRB1**
**PGK1**
**SLC30A6**
**NIN**
**LPGAT1**
**ZNF629**
**TNRC6C**
**SCML1**
**NIPA1**
**FAM98B**
**VKORC1L1**
**S100A1**
**PLA2G4C**
**BMPR2**
**KIAA1551**
**DMXL2**
**FAM169A**
**PNISR**
**ZNF565**
**ZNF703**
**KBTBD7**
**ZMYND8**
**MAP3K5**
**CCDC6**
**AJUBA**
**ZDHHC7**
**SOBP**
**EPB41L3**
**LIN7C**
**SIN3A**
**PLS3**
**BBS7**
**USP34**
**ARHGEF28**
**KCTD20**
**XIAP**
**MOSPD2**
**MAP1B**
**EPB41**
**NOL4**
**TMF1**
**CTGF**
**MIER3**
**ADH5**
**CAMTA2**

**DHX33**
**PPP1R3D**
**GSTCD**
**NUDT19**
**UBR4**
**ZKSCAN1**
**NACC2**
**FKBP10**
**PPP1CB**
**C2orf69**
**BNIP2**
**NCALD**
**SPRY2**
**MEX3C**
**YTHDC1**
**EIF3I**
**INO80D**
**PANK1**
**CUL4B**
**KDELR2**
**SIX6**
**GSPT1**
**AGPAT9**
**RMND5A**
**OR11A1**
**C7orf55-LUC7L2**
**ZCCHC3**
**TXNIP**
**GGNBP2**
**PIP4K2B**
**MT-ND2**
**ZNF229**
**PALM2-AKAP2**
**ZNF25**
**GDPGP1**
**PCGF2**
**GPR75-ASB3**
**RASL10B**
**SYNRG**
**TAF15**
**CTDSP2**
**ORAI1**

| miR-151a-5p | miR-221-3p |
|---|---|
| BMI1 | BMI1 |
| EFNB2 | EFNB2 |
| ZNF503 | ZNF503 |
| IRS2 | IRS2 |
| CDC6 | CDC6 |
| ACTB | ACTB |
| DUSP4 | DUSP4 |
| PRLR | PRLR |
| MSI2 | MSI2 |
| EGR1 | EGR1 |
| ACKR3 | ACKR3 |
| BICD2 | BICD2 |
| ADIPOR1 | ADIPOR1 |
| RNF44 | RNF44 |
| SOX4 | SOX4 |
| OLFM1 | OLFM1 |
| RIC8A | RIC8A |
| DAZAP2 | DAZAP2 |
| NFKBIZ | NFKBIZ |
| NAT1 | NAT1 |
| TXNL1 | TXNL1 |
| C18orf21 | C18orf21 |
| G3BP1 | G3BP1 |
| TCEB3 | TCEB3 |
| PLP2 | PLP2 |
| ATG13 | ATG13 |
| SNX11 | SNX11 |
| SMC1A | SMC1A |
| FSTL3 | FSTL3 |
| RP4-695O20__B.10 | RP4-695O20__B.10 |
| HNRNPR | HNRNPR |
| SOCS4 | SOCS4 |
| RIPK2 | RIPK2 |
| PLEC | PLEC |
| TM9SF1 | TM9SF1 |
| RAPGEF1 | RAPGEF1 |
| BRD2 | BRD2 |
| PHB2 | PHB2 |
| MAST4 | MAST4 |
| H2AFX | H2AFX |
| LPHN2 | LPHN2 |
| SCAF4 | SCAF4 |
| ZNF532 | ZNF532 |

| | |
|---|---|
| **MIER2** | **MIER2** |
| **ZBTB42** | **ZBTB42** |
| **SIK1** | **SIK1** |
| **GTF3C4** | **GTF3C4** |
| **FAT3** | **FAT3** |
| **WDR41** | **WDR41** |
| **VAC14** | **VAC14** |
| **PITX1** | **PITX1** |
| **BNIP1** | **BNIP1** |
| **KDM4B** | **KDM4B** |
| **RAB7A** | **RAB7A** |
| **ANXA1** | **ANXA1** |
| **PPIL1** | **PPIL1** |
| **ADSS** | **ADSS** |
| **RC3H2** | **RC3H2** |
| **MYOF** | **MYOF** |
| **HUWE1** | **HUWE1** |
| **MAP4** | **MAP4** |
| **ETFDH** | **ETFDH** |
| **TOM1L1** | **TOM1L1** |
| **STX3** | **STX3** |
| **COLEC12** | **COLEC12** |
| **IQGAP1** | **IQGAP1** |
| **CDC5L** | **CDC5L** |
| **CDH11** | **CDH11** |
| **OCLN** | **OCLN** |
| **ZNF620** | **ZNF620** |
| **APH1A** | **APH1A** |
| **DDX20** | **DDX20** |
| **OTUB1** | **OTUB1** |
| **CSDE1** | **CSDE1** |
| **LAPTM5** | **LAPTM5** |
| **SRM** | **SRM** |
| **PRPF19** | **PRPF19** |
| **ZNF292** | **ZNF292** |
| **KLHL21** | **KLHL21** |
| **CEP85** | **CEP85** |
| **BLMH** | **BLMH** |
| **IFITM3** | **IFITM3** |
| **ARFGEF2** | **ARFGEF2** |
| **FEM1C** | **FEM1C** |
| **TAB2** | **TAB2** |
| **SPRED2** | **SPRED2** |
| **SLC7A5** | **SLC7A5** |
| **RSL1D1** | **RSL1D1** |

| | |
|---|---|
| **PES1** | **PES1** |
| **CDK6** | **CDK6** |
| **EPDR1** | **EPDR1** |
| **MAP3K9** | **MAP3K9** |
| **RHEB** | **RHEB** |
| **ZC3H3** | **ZC3H3** |
| **FBL** | **FBL** |
| **CXCL2** | **CXCL2** |
| **PPIA** | **PPIA** |
| **EPC1** | **EPC1** |
| **CANX** | **CANX** |
| **DPP9** | **DPP9** |
| **ADIPOR2** | **ADIPOR2** |
| **SKP1** | **SKP1** |
| **ZNF57** | **ZNF57** |
| **TNRC6B** | **TNRC6B** |
| **PRRC2C** | **PRRC2C** |
| **HSPB1** | **HSPB1** |
| **HIVEP2** | **HIVEP2** |
| **SREBF2** | **SREBF2** |
| **MED1** | **MED1** |
| **RNF40** | **RNF40** |
| **ZNF207** | **ZNF207** |
| **TRIP12** | **TRIP12** |
| **RALGPS2** | **RALGPS2** |
| **CCNT1** | **CCNT1** |
| **SLCO5A1** | **SLCO5A1** |
| **PRR14L** | **PRR14L** |
| **RBM8A** | **RBM8A** |
| **SLC7A1** | **SLC7A1** |
| **ZNF721** | **ZNF721** |
| **SPINT2** | **SPINT2** |
| **XPO7** | **XPO7** |
| **PEG10** | **PEG10** |
| **FAM192A** | **FAM192A** |
| **NXF1** | **NXF1** |
| **LARP1** | **LARP1** |
| **NFAT5** | **NFAT5** |
| **EEF2** | **EEF2** |
| **NOL6** | **NOL6** |
| **PPP1R15B** | **PPP1R15B** |
| **BABAM1** | **BABAM1** |
| **BAG3** | **BAG3** |
| **CTNNB1** | **CTNNB1** |

| | |
|---|---|
| **TAPBP** | **TAPBP** |
| **NONO** | **NONO** |
| **WWC3** | **WWC3** |
| **NUFIP2** | **NUFIP2** |
| **TSR1** | **TSR1** |
| **SMCHD1** | **SMCHD1** |
| **ATP5A1** | **ATP5A1** |
| **PKMYT1** | **PKMYT1** |
| **MDGA1** | **MDGA1** |
| **RNASEK-C17orf49** | **RNASEK-C17orf49** |
| **MYC** | **MYC** |
| **FAM83D** | **FAM83D** |
| **CEP72** | **CEP72** |
| **CCT6B** | **CCT6B** |
| **PTGER2** | **PTGER2** |
| **LBH** | **LBH** |
| **HOXA3** | **HOXA3** |
| **RNASEK** | **RNASEK** |
| **COL5A1** | **COL5A1** |
| **ARID1A** | **ARID1A** |
| **E2F6** | **E2F6** |
| **FOXA1** | **FOXA1** |
| **PRKACA** | **PRKACA** |
| **XPR1** | **XPR1** |
| **DUT** | **DUT** |
| **HSP90AB1** | **HSP90AB1** |
| **ARHGDIA** | **ARHGDIA** |
| **SLC38A2** | **SLC38A2** |
| **GABARAPL1** | **GABARAPL1** |
| **DDB1** | **DDB1** |
| **EIF4G1** | **EIF4G1** |
| **RAB11A** | **RAB11A** |
| **MIA3** | **MIA3** |
| **COPZ1** | **COPZ1** |
| **SLC39A9** | **SLC39A9** |
| **RHOH** | **RHOH** |
| **KCMF1** | **KCMF1** |
| **ARL6IP6** | **ARL6IP6** |
| **MEPCE** | **MEPCE** |
| **TEAD1** | **TEAD1** |
| **TXNDC12** | **TXNDC12** |
| **NXN** | **NXN** |
| **GDI1** | **GDI1** |
| **ZNF592** | **ZNF592** |
| **TEX261** | **TEX261** |

| | |
|---|---|
| SMYD1 | SMYD1 |
| GPR137 | GPR137 |
| ZNF74 | ZNF74 |
| WDFY3 | WDFY3 |
| INSR | INSR |
| SNX2 | SNX2 |
| RPL26 | RPL26 |
| IPO4 | IPO4 |
| BMP2 | BMP2 |
| KDELR1 | KDELR1 |
| MPI | MPI |
| FAM199X | FAM199X |
| DHX15 | DHX15 |
| ZC3H11A | ZC3H11A |
| RRM1 | RRM1 |
| KRT80 | KRT80 |
| RAB40C | RAB40C |
| AMOTL2 | AMOTL2 |
| POT1 | POT1 |
| TAX1BP1 | TAX1BP1 |
| PIAS4 | PIAS4 |
| ARL6IP4 | ARL6IP4 |
| FBXW2 | FBXW2 |
| BRD3 | BRD3 |
| POMK | POMK |
| MPL | MPL |
| DMPK | DMPK |
| FOXP2 | FOXP2 |
| PABPN1 | PABPN1 |
| FAM210A | FAM210A |
| SEC23IP | SEC23IP |
| CAPN15 | CAPN15 |
| LONRF2 | LONRF2 |
| ATP6V1C1 | ATP6V1C1 |
| PPP1R3B | PPP1R3B |
| UBE2M | UBE2M |
| FAM102A | FAM102A |
| C6orf201 | C6orf201 |
| SLC7A2 | SLC7A2 |
| FN1 | FN1 |
| STARD7 | STARD7 |
| DIMT1 | DIMT1 |
| USP7 | USP7 |
| KIAA0232 | KIAA0232 |
| AAK1 | AAK1 |

| | |
|---|---|
| **PRKD3** | **PRKD3** |
| **R3HDM4** | **R3HDM4** |
| **HNRNPA2B1** | **HNRNPA2B1** |
| **BCOR** | **BCOR** |
| **ERGIC1** | **ERGIC1** |
| **STAT1** | **STAT1** |
| **VIMP** | **VIMP** |
| **PAPSS2** | **PAPSS2** |
| **IFITM2** | **IFITM2** |
| **AGPAT3** | **AGPAT3** |
| **DYRK1A** | **DYRK1A** |
| **XRCC5** | **XRCC5** |
| **PTEN** | **PTEN** |
| **HDHD2** | **HDHD2** |
| **MAPK1** | **MAPK1** |
| **SLAIN2** | **SLAIN2** |
| **MICAL3** | **MICAL3** |
| **ZNF561** | **ZNF561** |
| **TMPPE** | **TMPPE** |
| **GNG5** | **GNG5** |
| **RPL27A** | **RPL27A** |
| **GRB2** | **GRB2** |
| **CYB5R3** | **CYB5R3** |
| **BTG1** | **BTG1** |
| **SERTAD2** | **SERTAD2** |
| **SERBP1** | **SERBP1** |
| **SETD1A** | **SETD1A** |
| **OS9** | **OS9** |
| **RNF139** | **RNF139** |
| **RBM4** | **RBM4** |
| **RHOB** | **RHOB** |
| **NCKIPSD** | **NCKIPSD** |
| **JUP** | **JUP** |
| **CDK10** | **CDK10** |
| **DUSP1** | **DUSP1** |
| **SPOPL** | **SPOPL** |
| **BZW2** | **BZW2** |
| **GPS1** | **GPS1** |
| **SLC1A5** | **SLC1A5** |
| **ZFAND3** | **ZFAND3** |
| **USP10** | **USP10** |
| **MGAT4B** | **MGAT4B** |
| **COPS6** | **COPS6** |
| **SGK1** | **SGK1** |
| **AIG1** | **AIG1** |

| | |
|---|---|
| **N4BP1** | **N4BP1** |
| **RPS29** | **RPS29** |
| **RTN3** | **RTN3** |
| **CAMTA2** | **CAMTA2** |
| **DSP** | **DSP** |
| **KDELR2** | **KDELR2** |
| **TXNIP** | **TXNIP** |
| **NR4A1** | **NR4A1** |
| **MARCKS** | **MARCKS** |
| **ZFP91** | **ZFP91** |
| **BRD4** | **BRD4** |
