## Supplemental Table 3 for "Adenovirus 14p1 induced changes in miRNA expression increases lung immunopathogenesis"

ABCB9
ACTA2
ACTR3
ADAMTS5
ADCY3
ADGRE5
ADGRG1
ADORA1
ADORA2B
ADRB1
ADRB2
AGO4
AICDA
AIFM1
AKT2
AKT3
ALDH1A1
ANAPC1
ANKHD1/ANKHD1-EIF4EBP3
AP1S1
APAF1
APH1A
APOO
APPL1
AR
ARF3
ARF6
ARG2
ARHGEF2
ARPC5
ARRB1
ATP5MC3
ATP6V0A1
ATP6V1F
ATP6V1G1
ATPAF1
AURKB
BACH1
BAX
BBC3
BCL2
BCL2L1
BCL2L11
BCL3
BHLHE40
BLOC1S1
BMPR2
BRCC3
BTLA
CACNA2D3
CAMK2D
CAPN7

CASP3
CBL
CBX7
CCL3
CCL7
CCND1
CCND2
CCNE2
CCR7
CD1D
CD200R1
CD209
CD274
CD28
CD80
CDC25A
CDC34
CDK11A
CDK6
CDK8
CDKN1A
CDKN1B
CDKN2A
CDKN2B
CFL2
CHMP2A
CHRNA7
CHST7
CLPP
COL1A1
COL1A2
COL27A1
COL4A1
COL4A2
COL4A5
COL5A1
COL5A2
COL8A1
COPS7B
CSF1R
CSNK1D
CXCL11
CXCL12
CYB5B
CYP19A1
DCLRE1C
DDIT4
DDX19B
DDX58
DIRAS3
DKK3
DNA2

DNAJB5
DNAJB9
DNAJC1
DUSP10
E2F5
E2F6
EDN1
EEF2K
EFNB2
EGLN3
EIF1AX
EIF3J
EIF4EBP3
EIF4G2
EIF5A2
ENTPD7
ERBB3
ESR1
ETV3
EXOSC3
FADD
FANCA
FANCD2
FAS
FASLG
FGF7
FKBP1A
FLOT1
FMO2
FMO4
FNBP1
FNIP1
FOS
FOXO1
FOXO3
FOXP3
FTL
FZD3
GAB1
GAK
GALNS
GBA
GBP3
GLIS2
GLRX
GLS
GNAI3
GNAQ
GNG5
GNS
GOSR2
GPR132

GPR180
GPR183
GPR55
GPX7
GRB2
GRIN1
GRIN2B
GRIN2D
GSK3B
GSTM1
GTF2H2
GTF2H5
H3-3A/H3-3B
HACD3
HBEGF
HIF1A
HIF1AN
HMGA1
HMOX1
HNRNPA1
HSD17B4
HSD17B6
HSPA8
HTRA1
ICAM1
ICOS
IDE
IDH1
IFIT2
IGF1
IGF1R
IKZF1
IL10
IL12A
IL13
IL13RA1
IL34
IL6R
INPP5B
IRF2
IRF5
ITGA10
ITGA5
ITGAV
ITGB3
ITSN1
ITSN2
JAG1
KAT2B
KCNMB1
KCNN3
KDR

KIT
KITLG
KLF6
KLK10
KRAS
LAMC1
LATS2
LGALS1
LIMK1
LPAR6
LTB4R
MAP2K4
MAP3K13
MAP4K3
MAPK11
MAPK14
MAPK6
MARCKS
MATN3
MAX
MDM4
MEF2C
MFGE8
MLKL
MMP1
MMP10
MMP14
MMP16
MSH2
MT-ND4L
MXD1
MYC
NCOA1
NCSTN
NDST2
NDUFA1
NDUFB10
NDUFS4
NEDD4
NEFH
NF2
NGF
NLK
NLRP3
NOTCH1
NOTCH4
NRAS
NSF
NTF3
NTRK2
NUMB
NUP43

NXN
OAS1
OAS2
ODC1
PAIP1
PAIP2
PARD6B
PARP16
PCGF1
PDCD4
PDE1A
PDGFA
PDGFB
PDK4
PDPK1
PELI1
PIK3C2A
PIK3R1
PIK3R3
PLCL2
PMAIP1
POLD2
POLR2C
POLR2D
POM121/POM121C
POU2F1
PPARA
PPARG
PPARGC1B
PPIF
PPP1R12B
PPP1R3D
PPP1R7
PPP2R2A
PPP3R1
PRDM1
PRIM1
PRKAR2A
PRKCB
PRKCD
PRKCE
PRPF38A
PRPF38B
PRSS22
PSEN1
PTAFR
PTEN
PTGER4
PTGS2
PXN
RANGAP1
RAP1B

RAPGEF3
RASA1
RBBP7
RCAN2
RCOR1
RDX
RFXANK
RGS16
RHOB
RHOG
RNF168
RNF4
RPL14
RPL15
RPL34
RPL36A
RPS3
RPS6KA5
RSAD2
RUNX1
RXRA
S100A8
S1PR1
SELENOT
SERPINF2
SESN1
SH3BP4
SHE
SIGMAR1
SIRT1
SKP2
SLC1A2
SLC25A13
SLC38A1
SLC7A5
SMAD3
SMAD4
SMAD5
SMAD7
SMC1A
SMOX
SMUG1
SNAI1
SNAP25
SNRNP27
SOCS1
SOCS3
SOCS4
SOCS5
SOD2
SOD3
SRD5A1

SRF
SRM
ST14
TAF9B
TARBP2
TBK1
TCERG1
TFRC
TGFBR1
TGFBR2
TGFBR3
THBS1
TIMP3
TLR4
TLR8
TNF
TNFSF10
TNFSF15
TNFSF9
TNPO1
TOB1
TP53
TPM1
TPR
TRA
TRPM7
TSLP
TUBA1A
TXN2
TYRO3
UBA1
UBE2D1
UBE2D3
UBE2K
UBE2V1
UBE2Z
VASP
VEGFC
VIM
VPS39
WASL
WEE1
XRN1
YPEL3
YWHAQ
YWHAZ
ZFP36L1
ZNF420
